# Sampling Microbial Dynamics in the Salish Sea Estuary: Evaluating Methods to Capture Cyanobacteria and Cyanophage

**DOI:** 10.1101/2025.11.30.691463

**Authors:** Noelani R. Boise, Owen P. Leiser, Kristin Jones, Mahala Peter-Frank, Damon Leach, Stephen Crafton-Tempel, Peter Regier, Ruonan Wu, Conner Phillips, Margaret S. Cheung, Connah G. M. Johnson, Scott J. Edmundson, David D. Pollock

## Abstract

Picocyanobacteria from the genera *Prochlorococcus* and *Synechococcus* thrive across the globe in aqueous environments, have relatively small genomes, and have growth dynamics regulated by both viral interactions and abiotic conditions, making them excellent model organisms for exploring host-pathogen coevolution. The Salish Sea, located in the Western coastal waters bordering the USA and Canada, is at the current northern boundary (defined by *Prochlorococcus* versus *Synechococcus* prevalence ratios) of the range of *Prochlorococcus*. Predictions suggest that this boundary will shift northward as warmer waters move northward, providing an excellent system to study host-pathogen dynamics and coevolution in a changing environmental context. In preparation for such studies, we developed and refined methods to sample and sequence cyanobacteria, cyanophages, and their abiotic environment. In addition to basic methodological questions focused on the physical sampling, filtering, viral precipitation, DNA extraction, and technical replicability, we explored how well our filtering and extraction protocols enrich for our main target, picocyanobacteria. The protocol described herein can successfully discriminate large-cell eukaryotic organisms, but size fractionation of picocyanobacteria appears to be affected by the presence of free DNA, multicellular structures, and abundant tycheposons. Our preferred final protocol at the conclusion of these experiments based on yield and processing time is presented. We recovered substantial *Prochlorococcus, Synechococcus* and amoeba-like sequences in most samples, and preliminary exploration of relative taxon sequence read recoveries across locations, over time, and tidal conditions are also discussed. Approaches described here may be useful to other efforts such as harmful algal bloom monitoring, species isolation and enrichment, water quality assessments, anti-viral discovery, and understanding picocyanobacterial population changes over space and time.

## Introduction

The picocyanobacteria *Prochlorococcus* and *Synechococcus* are collectively a ubiquitous component of phytoplankton and play an outsized role in global primary production in both freshwater and marine systems (Callieri, 2022). They have a large geographic distribution year-round (Flombaum, et al., 2013), but are more abundant in warm summer months (Zinser, et al., 2006). Typically ranging in size from 0.2-2 µm, *Prochlorococcus* and *Synechococcus* are among the most abundant photosynthetic organisms on Earth and are key indicators of ecosystem health. Due to their prevalence in various marine and estuarine ecosystems, their relatively small genomes, and their valuable anti-viral properties, these organisms are exceptional model organisms for studying viral-bacterial dynamics (Visintini, et al., 2021; Scanlan, et al., 2009; Reynolds, et al., 2021; Edmundson, et al., 2020). Well-adapted to oligotrophic conditions (Kent, et al., 2019) they also thrive in the euphotic zone of tropical and subtropical waters and are expected to expand their distribution as ocean temperatures rise and impact oceanic communities (Flombaum, et al., 2013). Given their ubiquity, these organisms are likely present in the coastal and estuarine waters of the northern Olympic Peninsula of Washington State, USA and surrounding Salish Sea waters, but few studies specifically focus on picocyanobacteria in this region (Del Bel Belluz, et al., 2024). The Salish Sea lies along what was historically considered the northern boundary of the distribution of *Prochlorococcus*, but this boundary is expected to have shifted and continue to change as ocean temperatures increase (Ribalet, et al., 2025; Flombaum, et al., 2013; Mioduchowska, et al., 2023), making the changing interactions and relative success of *Prochlorococcus, Synechococcus*, and their viruses topics of considerable ecological and biopreparedness interest (Edmundson, et al., 2020; Reynolds, et al., 2021).

The Salish Sea is an expansive network of complex water bodies including the Strait of Juan de Fuca, the Strait of Georgia, and Puget Sound. Kathryn Sobocinski’s *The State of the Salish Sea* report showcases the dynamic currents and biogeochemistry of these waters (Sobocinski, 2021). Strong upwelling of deep, nutrient-rich water due to outflow of freshwater from the Strait of Juan de Fuca produces dynamic seasonal fluctuations. This wind-driven upwelling of deep water can also decrease dissolved oxygen (DO) levels and increase nutrient-rich water from the coast which circulates through the strait. Wastewater from the heavily populated coastal areas around the Salish Sea also has significant effects on nutrient levels and thus cyanobacterial populations in these water bodies; in the summer, reduced freshwater outflow can lead to stagnant wastewater effluent and agricultural runoff in small regions of the Salish Sea, increasing seasonal eutrophication (Sobocinski, 2021). Despite extensive research on impacts of environmental conditions on picocyanobacteria species, there is far less work documenting the impact of environmental factors on picocyanobacteria-cyanophage dynamics. Some work shows examples of environmental impacts, such as diel cycles impacting cyanobacterial gene expression and phage-host interactions (Liu, et al., 2019; Rozum, et al., 2025) and infection rates increasing in euphotic zones (Fuchsman & Hays, 2023).

Currently, there are few methods that provide relatively rapid biomass samples that target multiple size fractions from the same source water, which is critical for isolating both viruses and hosts from a discrete timeframe in addition to the environmental context these organisms are found in. By establishing consistent methods to obtain quality genomic and environmental data across locations, seasons, tidal stages, and diel cycles, this work yields holistic information to assess changing dynamics between marine cyanophages and picocyanobacteria, which can inform models that predict the behavior of higher-risk species. In a time of increasing risks in viral-host adaptations and spillover due to rapid environmental change (Carlson, et al., 2022a), it is critical that the scientific community establish consolidated methods for pairing environmental data with molecular research at a large scale to understand past, present, and future adaptation and behaviors of viruses and their hosts.

In this study, we tested and improved methods to sample and filter raw environmental water samples to extract DNA, identify picocyanobacteria and their associated phages, and describe their biotic environment, with the long-term aim to study host-pathogen interactions and coevolution. To achieve this, we collected samples from five locations at varying times of day and tidal conditions along the northern Olympic Peninsula over an initial year-long period while performing iterative method development. Sampling involved collecting surface-level samples either via in-line *in situ* serial filtration or direct surface sampling. Due to the wide variety and size distribution of phytoplankton organisms and specifically cyanobacteria (Allaf & Peerhossaini, 2022), we performed serial filtrations to remove larger debris and to reduce the amount of larger-celled non-target organisms prior to sequencing. We focused on identifying relative read counts of target organisms in the genera *Prochlorococcus* and *Synechococcus*, along with previously identified cyanophage pathogens. Based on preliminary results showing that reads similar to eukaryotic amoeba were consistently observed in our samples, we used relative amoeba read counts as a marker for removal of large-celled organisms.

In summary, the present work describes the development and evaluation of methods for enriching and analyzing picocyanobacteria and associated cyanophages in the Salish Sea for metagenomic sequencing, along with preliminary observations of select lineages under various collection conditions over time. This work highlights the efficacy and replicability of sequential filtration and DNA extraction protocols to support studies of host-pathogen dynamics and environmental impacts on microbial interactions.

## Materials and Methods

### Sampling, filtering and processing

We performed initial targeted sampling in Salish Sea waters from November 2023 to February 2025, mostly from the PNNL-Sequim floating dock and pier, and less frequently at four nearby locations: Cline Spit, south of John Wayne Marina near an estuary outflow, open waters North of Port Townsend, and in Discovery Bay (Table 1, Figure 1). Sampling events, including filtering and collection (Figures 2 and 3) occurred near-monthly at the PNNL-Sequim dock over the course of a year, with offsite sampling and multiple within-month sampling concentrated during peak summer months. A total of 30 primary sampling events were performed on 25 separate days at ebb, flood, and slack tides, with the majority occurring during flood tides. Water levels, temperature, salinity, and collection dates are shown in Figure 4.

**Table 1.** Sample location descriptions for marked sites in Figure 1. All sites were sampled only once due to logistical challenges except for the PNNL-Sequim dock site, which was sampled 26 times in year one.

|  | Sampling Location | Site Coordinates | # of Sampling Events |
| --- | --- | --- | --- |
| 1 | Cline Spit Loading Dock | 48°09'06"N 123°09'09"W | n = 1 |
| 2 | PNNL-Sequim Floating Dock | 48°04'45"N 123°02'42"W | n = 26 |
| 3 | Exterior southern side of John Wayne Marina | 48°03'41"N 123°02'20"W | n = 1 |
| 4 | Southern end of Discovery Bay | 47°59'48"N 122°52'32"W | n = 1 |
| 5 | Open water north of Port Townsend | 48°09'56"N 122°48'24"W | n = 1 |

**Figure 1.**
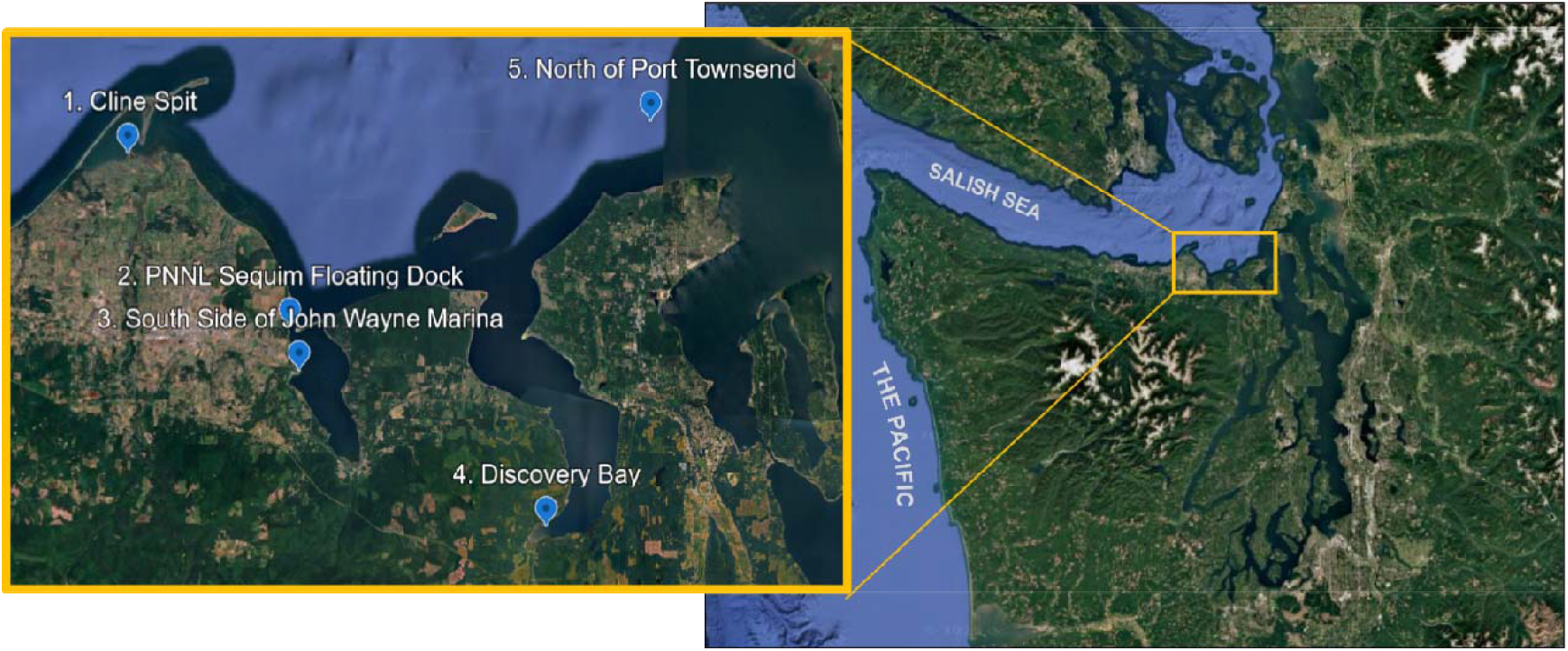
Sampling locations in the Salish Sea during the first year of sampling. 1) Cline Spit boat launch, 2) PNNL-Sequim floating dock, 3) Southern side of John Wayne Marina near estuary outflow, 4) Discovery Bay, 5) North of Port Townsend. All locations are located near the entrance of Admiralty Inlet from the Pacific Ocean.

**Figure 2.**
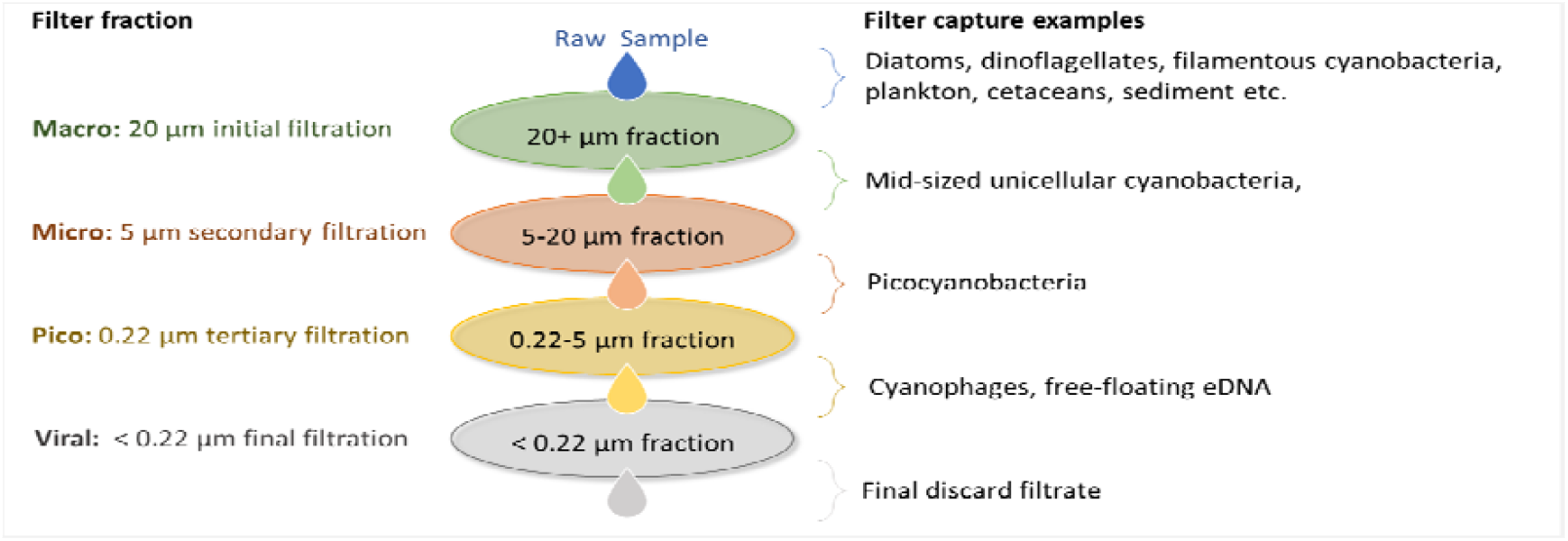
Serial filtration fractions. The projected target species enrichment goals for size fractionation are shown. The four fractions are: macro (> 20 µm); micro (5-20 µm); pico (and small micro, 0.22-5 µm); and viral (filtered through 0.22 µm, then precipitated, then collected on 0.22 µm filters). Note that the labels for these fractions are rough approximations, and the “pico” fraction in particular is generous at the upper end (5 µm) to avoid filtering out organisms with irregularly grouped cells at the larger end of the picocyanobacterial range (*Prochlorococcus* 0.5-0.7 µm, *Synechococcus* 0.8-1.5 µm).

**Figure 3.**
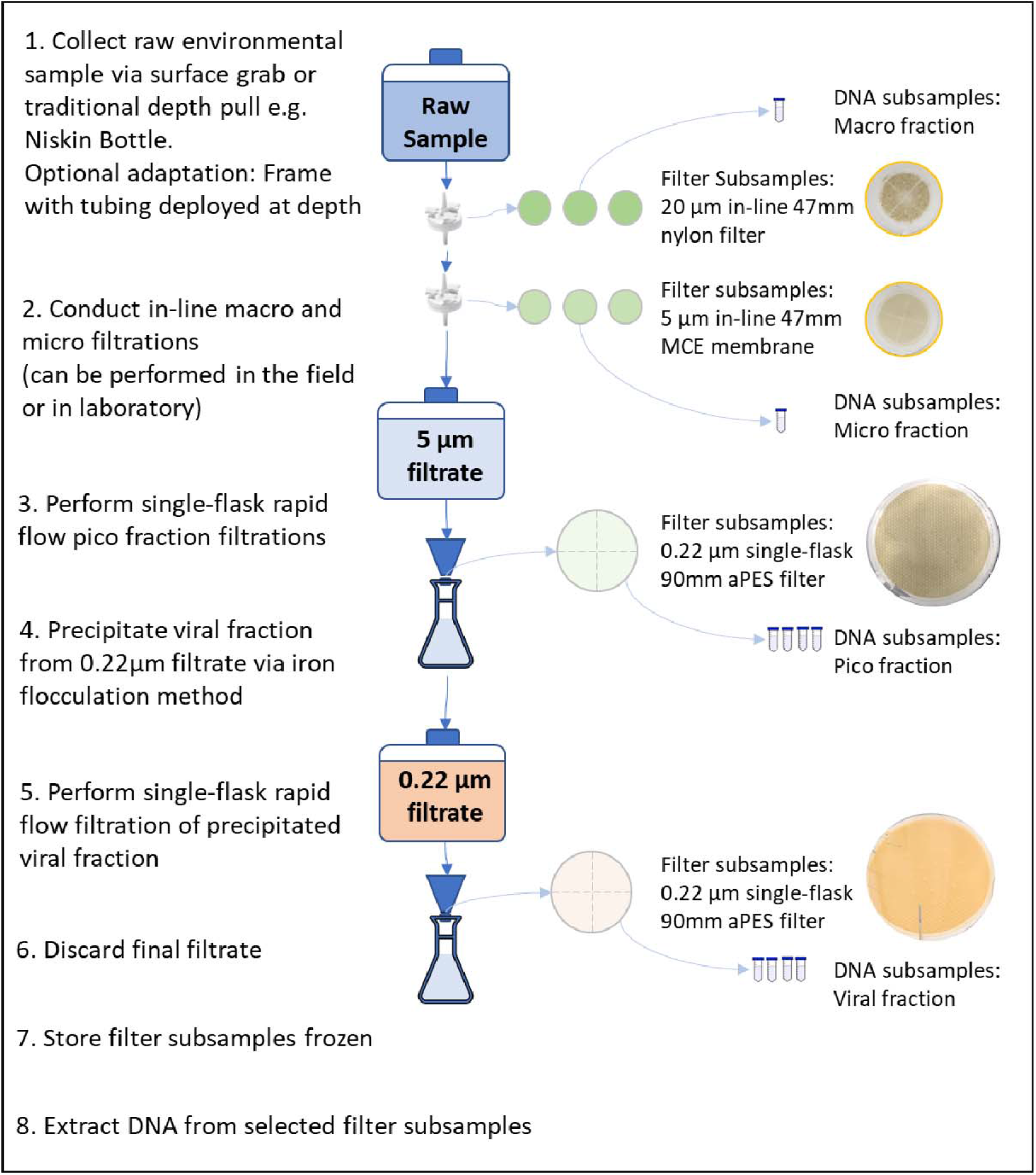
Final sampling scheme. Scheme incorporates a combination of in-line filtrations and single-flask filtrations to increase time-efficiency, minimize material waste, and improve yield of pico and viral fractions. Filters were replicated either as subsequent replacement filters in the in-line or flask filtration process, or as cut fractions through the midpoint of the filter, obtained by physically cutting individual filters after sample collection.

**Figure 4.**
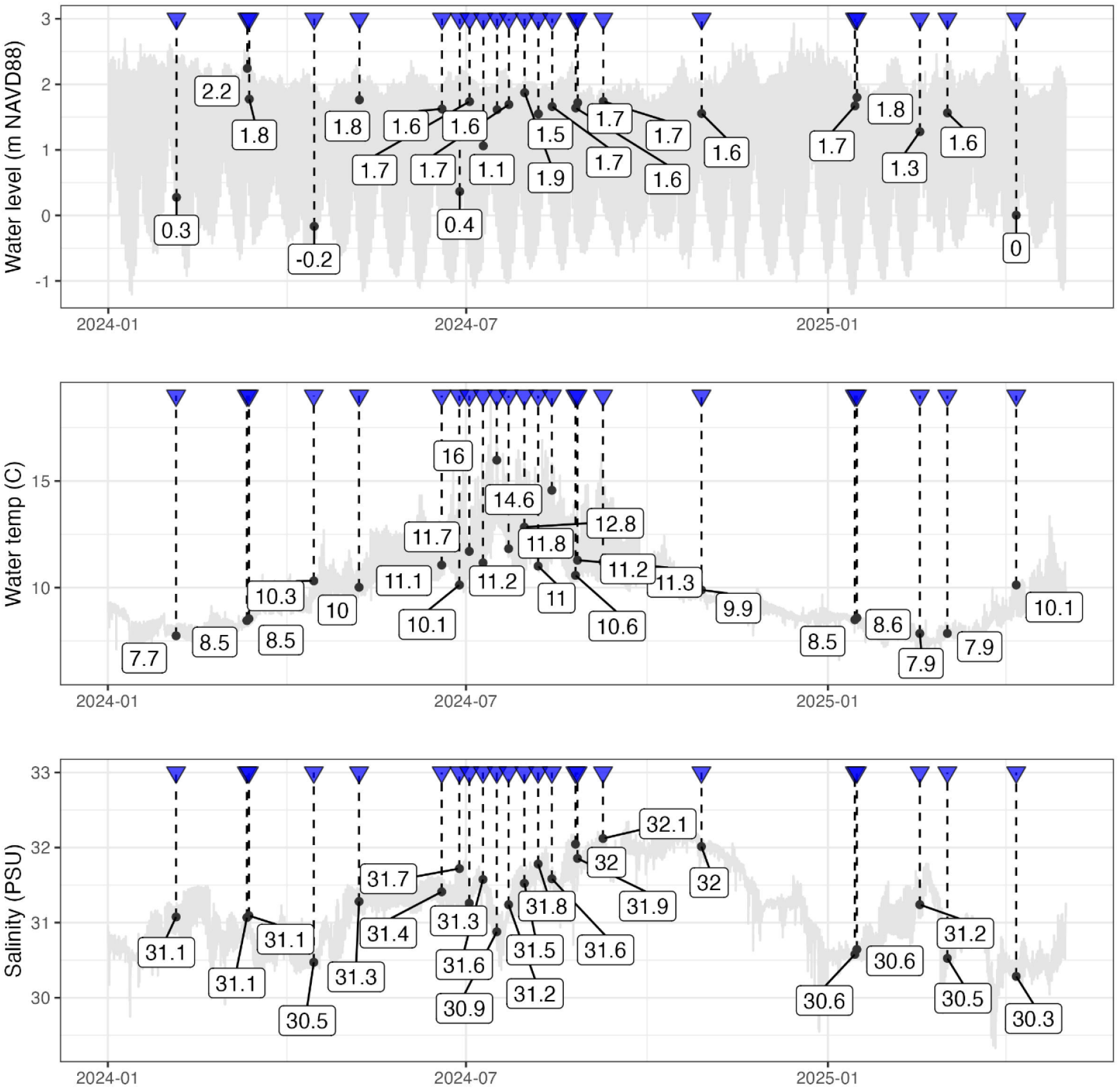
Seasonal data for water levels, temperature, and salinity. Values were measured using the MCRL Data sensor suite with denoted sampling events from 2024-2025.

Samples were collected using two pilot methods, either 1) time-discrete samples using a horizontal water sampler, Niskin bottle, or direct surface sampling using a sterilized carboy; or 2) time-extended *in situ* in-line filtrations at the sampling site. The *in-situ* apparatus consisted of a custom-designed, weighted metal frame attached to flexible tubing that we tested using both peristaltic and vacuum pumps (Supplemental Table 1). We configured the tubing as an *in-situ* mechanism for two- and three-step serial filtrations using in-line filters that could be deployed from height and immediately pump raw seawater directly through two or three sequentially smaller filter sizes to capture target species, with the final filtrate collected into a sterile vessel. Alternatively, we used a direct water sampler (a Niskin bottle, horizontal water sampler, and/or a sterilized carboy) to collect discrete samples followed by filtrations in a laboratory. The in-line system collected samples over extended periods of time (up to 3 hours with especially dense water samples through a three-step filtration), compared to sampling durations of less than one minute using a direct water sampler.

Prior to sampling events, all sample collection vessels (carboys, Niskin bottles, and horizontal water samplers) were sterilized by autoclave or disinfected using a 10% bleach solution. We primed the collection vessel, tubing, and priming filters prior to every sample collection with water from the site, then discarded the priming water away from the collection point to avoid contamination. After collecting a pre-determined target volume, we immediately recorded metadata and sampling event context and assigned a single parent sampling ID to all associated data. Sampling conditions were noted, and additional weather data from the day of sampling were downloaded from the PNNL-Sequim Marine and Coastal Research Laboratory (MCRL) data site (MCRL Data, 2025).

Depending on which fractions were already processed, we then transported samples to our laboratory in Sequim WA to process the remaining filter fractions. All tubing and vessels were rinsed with 70% ethanol and/or autoclaved prior to priming with the appropriate matrix (we used each sequential filtrate to prime the subsequent collection vessels). For most samples, this involved a combination of in-line filtrations (20µm 47 mm nylon filter and a 5 µm 47 mm mixed-cellulose ester (MCE) membrane secured in either a Millipore 47 mm polypropylene nylon filter holder or a Cytiva 47mm polycarbonate in-line filter holder using a peristaltic pump, Supplemental Tables 2 and 3) and single vacuum flask filtrations. In-line filtration was followed by mixing in a single-flask and filtration onto a sterile 0.22 µm polyethersulfone (PES) or MCE filter in a vacuum flask to capture the pico fraction. Flow-throughs from the 0.22 µm filter were precipitated using a method adapted from John et al. (2011); iron chloride hexahydrate was added to the filtrate at 1 mL of 10g/L FeCl · 6H_2_O solution in deionized water per 20 L of seawater and incubated for 1 hour, and precipitates were captured on a sterile 0.22 µm PES filter via vacuum filtration.

Additional filtration setups we tested included a 0.05 µm filter after 0.22 µm filtration, and various alternate size combinations with the three filter types: nylon, MCE membranes, and PES. Specifically, we tested 47 mm and 90 mm 0.22 or 0.2 µm MCE or PES filters, either as individual, sterile membranes or sterile vacuum units. We tested the 5 µm and 0.22 µm MCE membranes in an in-line filtration setup using either a polycarbonate filter-holder purchased from Avantor Sciences or a polypropylene filter- holder purchased from Millipore Sigma. The long-term use of 47 and 90 mm filters for differing filter fractions was driven by limited commercial availability of each material-pore size-diameter combination and for time-efficiency. A list of the tested filters, in-line filter holders, and other product information can be found in Supplemental Tables 1, 2, 3 and 4.

In the final sampling scheme, the sampling process included separating environmental samples by serial filtration into four fractions (Figure 2), labelled by their target organism size and the nominal pore sizes of the filters which they passed through and on which they were collected: macro (> 20 µm), micro (large micro, 5-20 µm), pico (small micro, 0.22-5 µm), and viral (< 0.22 µm). Further details are provided in Figure 3. Note that filter subsamples for the 20 µm and 5 µm filters were obtained by changing filters as they became clogged (as estimated qualitatively by reduced flow rate), whereas most filter subsamples for the 0.22 µm filters (both the pico and the viral fractions) were obtained by physically cutting the filters into thirds or quarters after completion of filtration (Figure 3).

#### DNA Extraction

DNA was extracted from selected filter subsamples using Zymo s Quick DNA Plant/Seed Miniprep Kit, and three Qiagen kits: DNeasy PowerWater Kit, PowerSoil Pro Kit, or Blood and Tissue Kit (Supplemental Table 4). Following extraction, DNA quality and quantity were assessed using a NanoDrop One spectrophotometer (ThermoFisher Scientific, Waltham, MA, USA) and Qubit4 fluorometer system (ThermoFisher Scientific, Waltham, MA, USA). Samples were then frozen, and select samples were later analyzed on an Agilent 4150 TapeStation for quality assessment (Supplemental Data Figure 1).

For direct comparisons, DNA from select sampling events were extracted using two separate kits (Blood & Tissue and PowerWater) and analyzed on an Agilent 4150 TapeStation using a D5000 ScreenTape and the relevant kit reagents (D5000 buffer and D5000 ladder) following the kit protocol with no modifications (Agilent Technologies, Palo Alto, CA, USA).

### Metadata collection and processing

PNNL-Sequim is uniquely positioned for marine environmental research, leveraging its bench-to-bay capabilities through direct access to Sequim Bay, raw seawater access in the facilities, and a unique array of permanently deployed environmental sensors (MCRL Data, 2025). Most sampling events took place from the PNNL-Sequim floating dock at the mouth of Sequim Bay with an extensive range of environmental parameters documented from the MCRL Data, while other locations relied solely on parameters captured from a YSI ProDSS multiparameter water quality meter (YSI Xylem, Yellow Springs OH) (Supplemental Data Table 5). For PNNL-Sequim sampling events, we acquired data post-sampling using targeted scripts (MCRLdata-Sandbox, 2025a; MCRLdata-Sandbox, 2025b) to process environmental data from an online data repository (MCRL Data, 2025). One sampling event used for a controlled membrane test was executed in PNNL’s marine mesocosm system, which circulates unfiltered seawater through a large (ca. 62,000 L) outdoor tank (Marine and Coastal Research Laboratory, 2025).

We used data collected for tidal stage, water temperature, and salinity by the sensors described above to provide environmental context for sampling events off the PNNL-Sequim dock (Figure 4). Data collected by sensors are sent to the cloud where they undergo automated initial quality control and displayed in near real-time (MCRL Data, 2025) and are provided as part of the joint metadata summary (MCRLdata-Sandbox tide gauge, 2025; MCRLdata-Sandbox CTD, 2025). Additional quality control combining automated (e.g. de-spiking) and manual visual assessment were then applied, resulting in several data gaps originating from sensor maintenance, sensor errors, or unreliable data (identified during quality control). To fill any gaps, we used the *ranger* R package to construct Random Forest models (Wright & Ziegler, 2017). First, we gap-filled water levels by building a Random Forest model using water levels from NOAA gauge #9444090 (NOAA, 2025) pulled using the *VulnToolkit* R package along with hour of day, and day of year as predictors (Hill & Anisfeld, 2021). The resulting model (R^2^ = 0.96) was then used to gap-fill water levels. Those gap-filled water levels were used (along with hour of day and day of year) to predict both water temperature and salinity measured off the MCRL dock by CTD (conductivity, temperature, and depth sensor), and those models were used to gap-fill both temperature and salinity time-series. Model goodness-of-fit was high for both temperature and salinity models (R^2^ = 0.93 and 0.95, respectively).

### Sequencing and preliminary data processing

DNA subsamples were shipped on ice to Azenta Life Sciences (South Planfield, NJ,USA) for sequencing. Sequencing libraries were prepared using NEBNext Ultra DNA Library Prep Kit (New England Biolabs, Ipswich, MA, USA) according to manufacturer’s instructions. Briefly, genomic DNA was acoustically fragmented using a Covaris S220 instrument (Covaris, Woburn, MA, USA) and fragmented DNA was cleaned and end repaired. Sequencing adapters were ligated to 3’ ends of fragments after adenylation and enriched by limited-cycle PCR. Sequence libraries were validated using a High Sensitivity D1000 ScreenTape on an Agilent TapeStation (Agilent Technologies, Palo Alto, CA, USA) and quantified using a Qubit 3.0 fluorometer (ThermoFisher Scientific, Waltham, MA, USA). Sequence libraries were also quantified using real-time PCR (Applied Biosystems, Carlsbad, CA, USA) libraries from samples that passed all quality control steps and were multiplexed and clustered onto an Illumina NovaSeq flowcell (Illumina, San Diego, CA, USA) and sequenced using 2×150 paired-end configuration. Base calling and image analysis were conducted using the NovaSeqControl Software. Raw sequence data was converted to fastq format and de-multiplexed using Illumina’s bcl2fastq software v2.20, allowing one mismatch for index sequence identification. Sequencing datasets were processed through the sequence processing pipelines described below for evaluation of the efficacy of filter fractionation methods and to refine strategies for further sampling. Raw paired-end reads were sequentially pre-processed by first interleaving paired reads into a single sequence read file preserving read pair information using the reformat functionality of BBTools v37.62 (Bushnell, et al., 2017). Interleaved reads were clustered, with optical and PCR duplicate reads removed using clumpify within BBTools. Reads derived from low-quality regions of flowcells were removed using the filterbytile functionality from BBTools. BBTools bbduk was used to sequentially remove Illumina sequencing adapters, bases from the ends of reads below a phred33 quality score of ten, any reads with an overall phred33 quality score lower than ten, and any reads mapping to PhiX spike-in. Read quality was assessed before and after preprocessing using FastQC v0.12.1 (Babraham Institute, 2023).

We generated metagenome contig assemblies using MEGAHIT v1.2.9 (Li, et al., 2015) under default settings in paired-end mode. Output contigs were filtered by length as required for downstream analyses using the reformat functionality of BBTools. Assembly qualities were assessed using QUAST v5.3.0 (Gurevich, et al., 2013). Putative coding regions and putative protein sequences were predicted within metagenome assemblies filtered to contigs larger than 2500 bases using Metaprokka v1.15.0 (Seeman, 2014).

Raw short read pairs, interleaved pre-processed reads, contig assemblies, and predicted genes and proteins are found in Supplemental Data 3 and were deposited to BioProject PRJNA1336290.

### FAIR Data Management

We assigned each sampling event a unique parent ID that was tagged to the relevant environmental data for every filter and DNA subsample, which also had its own unique subsample ID. To track sampling events, we created a standardized field form with nomenclature and categorization based on the National Microbiome Data Collaborative (NMDC) water sample template for data documentation compatibility with the other public repositories (National Microbiome Data Collaborative, 2025). The unique sample identifiers for all parent and subsamples were used to further facilitate consistent data integration. Sequences, processed sequences, analytical results and associated metadata were captured and a) stored for sharing among the team using DataFed federated data storage and Globus-enabled federated IDs to ensure that this data is searchable and well-structured (Johnson & Pollock, 2025; EcoKMER GitHub Software Repository, 2025) and b) harmonized and visualized using the concurrently-developed EcoKMER program (Johnson & Pollock, 2025) with final output stored as a CSV file (Supplemental Data 1). This integrated metadata file combined sequencing summary information with environmental and contextual data, adhering to FAIR (Findable, Accessible, Interoperable, and Reusable) principles (Wilkinson, et al., 2016) and ensures that this data is searchable and well-structured.

### Microbial sequence diversity estimation

To assess sequence diversity across environmental metagenomes, we applied Nonpareil (v3.5.5) to interleaved pre-processed reads (Rodriguez-R, et al., 2018). Reads shorter than 24 base pairs (bp) were excluded. Nonpareil estimates average community coverage and diversity using kmer redundancy, modeling the rate at which new 24 bp sequences are encountered with increasing sequencing effort. The resulting Nonpareil diversity index (Nd) used does not cleanly reflect microbial diversity, as it reports diversity inferred from identical 24mers sampled, and does not account for variable rates along genomes, or variable genome size, but it has nevertheless been demonstrated to be useful in detecting differences in community composition (Rodriguez-R, et al., 2018).

### Relative read coverage of target taxa

61 DNA subsamples produced in year one were sequenced, filtered, assembled into contigs, and 24mer sequence diversities were calculated. To quantify taxon-specific enrichment of organisms with varying sizes, we mapped metagenomic reads to curated reference genome sets using BBMap (v39.13) (Bushnell, 2014). Five reference genome sets were constructed from high-quality, complete NCBI genomes representing *Prochlorococcus* (26 reference genomes), *Synechococcus* (78 reference genomes), their phages (341 *Synechococcus* phage genomes; 37 *Prochlorococcus* phage genomes), and *Acanthamoeba spp* (5 reference genomes; accession numbers of each reference set in Supplemental Data 2). Our main reason to focus on these taxon groups was to identify the relative occurrence of the two picocyanobacterium and cyanophages that might be associated with them. In addition, we separately mapped reads to a mycetezoan amoeba to determine the effectiveness of filtering out large-celled organisms. Each set was indexed using the BBMap indexing utility. Quality-filtered reads from each sample were aligned to each reference genome set using default parameters, and coverage statistics were calculated with the BBMap pileup utility (Supplemental Data 1), which attaches each read to the reference genome in a run that matches it best, and leaves unattributed any reads that do not pass the minimum matching cutoff to any included genome(s).

Each of the six genome groups listed above were analyzed separately. During preliminary analysis, we observed that the bulk of *Acanthamoeba* reads were consistently attributed to the five major chromosomes of *Dictyostelium discoideum*, so we opted to present the sum of plus and minus reads (relative to genome orientation) most preferentially attributed to these five chromosomes as a marker for large-celled amoeboid organisms. *Dictyostelium discoideum* is nominally a soil organism, but the representation of diverse amoeboid organisms in the genome references was not strong, and these results should not be construed to positively indicate an undue degree of soil contamination in estuarine waters or to identify the actual genus or species source in these aquatic samples—relative amoeba read counts were only used as a marker for removal of large-celled organisms. For other runs, we report the sum of the plus and minus reads for all genomes in the reference set. To normalize differences in sequencing depth, total read counts per sample were used to compute reads per million (RPM). For comparison, we ran Kraken/Bracken (Lu, et al., 2022) using version 2.0.7-beta of the Kraken database using default parameters, outputting the total reads that were attributable to the groups *Prochlorococcus, Synechococcus*, and *Dictyoselium discoideum*. The output is reported in Supplemental Data 1. Read counts using Bracken are generally lower than for BBMap, likely due to use of a smaller reference database and differences in default parameters.

## Results

### Sample collection and processing

Available pumps (see Methods and Supplemental Table 1), including eDNA and peristaltic models, did not reliably generate enough power to overcome sampling challenges at height from piers or vessels, so direct surface sampling was preferred over *in-situ* filtration. Biomass accumulation quickly clogged the 0.22 µm inline filter, which inhibited consistent flow and led to eventual removal of 0.22 µm inline filters from our protocol in favor of flask filtration of the mixed 5 µm filtrate in the tertiary filtration step (Figures 2 and 3).

Millipore’s 47 mm polypropylene nylon in-line filter holders improved flow rates over Cytiva’s 47 mm polycarbonate filter holders, reducing macro and micro filtration time by over 50% and reducing sample loss due to inhibitive filtration times and insufficient gasket seals in alternative holders. The 30 primary sampling events were ultimately segregated by filtration and technical replicates into 294 filter subsamples and 424 DNA subsamples, with filters cut and used for replicate DNA extractions.

### Environmental Metadata

As expected, we found that water temperatures increased at the entrance of Sequim Bay through summer months, peaking from late July to early August 2024, with lows in February 2025. The lowest recorded water temperature of the 26 sampling events from PNNL Sequim was 7.7°C on 2 January, 2024, and the highest recorded temperature was 14.7°C on 17 July, 2024. The sampling event at Cline Spit had the highest recorded temperature of 19.8°C on 7 August, 2024. The highest salinity value derived from the MCRL dock data was 32.1 PSU on 18 August, 2024 and the lowest was 30.3 PSU on 7 April, 2025. Surface sampling events occurred across a range of tidal events, ranging from -0.2 to 2.2 meters in water depth. Of all 30 sampling events, two were sampled during a slack low tide, three during a slack high tide, 17 during a flood tide, seven during an ebb tide, and one event from the Sequim outdoor mesocosm tank did not have an assigned tidal stage.

### DNA yields under various conditions

In an exploratory controlled comparison of DNA yields, the Blood & Tissue kit outperformed the PowerWater and PowerSoil kits, yielding approximately 3900, 690, and 2100 ng DNA, respectively (Figure 5). For this controlled comparison, two pico filters (90mm PES) from a single sampling event were cut into thirds, and DNA was extracted from each third using a different kit (n = 2 extractions per kit). However, a ‘kitchen sink’ comparison (a mix of conditions and replicates, controlling only for filter pore size and location) of each filter fraction across all DNA subsamples collected at PNNL-Sequim (macro, micro, pico, and viral) that also included the Zymo Plant/Seed kit (Figure 6) resulted in highly variable yields under each condition such that there were no significant differences in the micro, pico, and viral fractions, and overlapping significance groups in the macro fraction as determined by a Tukey’s post hoc test (p < 0.05) after performing an analysis of variance (ANOVA). Yields in the three direct filter fractions (macro, micro, and pico) were on average relatively high (∼500-1000 ng or more), with a large proportion having acceptable yields for downstream sequencing (50 ng or greater; Figure 6). This supports our approach of running filters to saturation to obtain sufficient material for DNA extraction. Perhaps unsurprisingly, the viral fractions had generally lower DNA yields, averaging 450 ng compared to an average of 1210, 1272, and 924 ng for the pico, micro, and macro fractions, respectively. However, viral DNA yields were often still high enough for sequencing.

**Figure 5.**
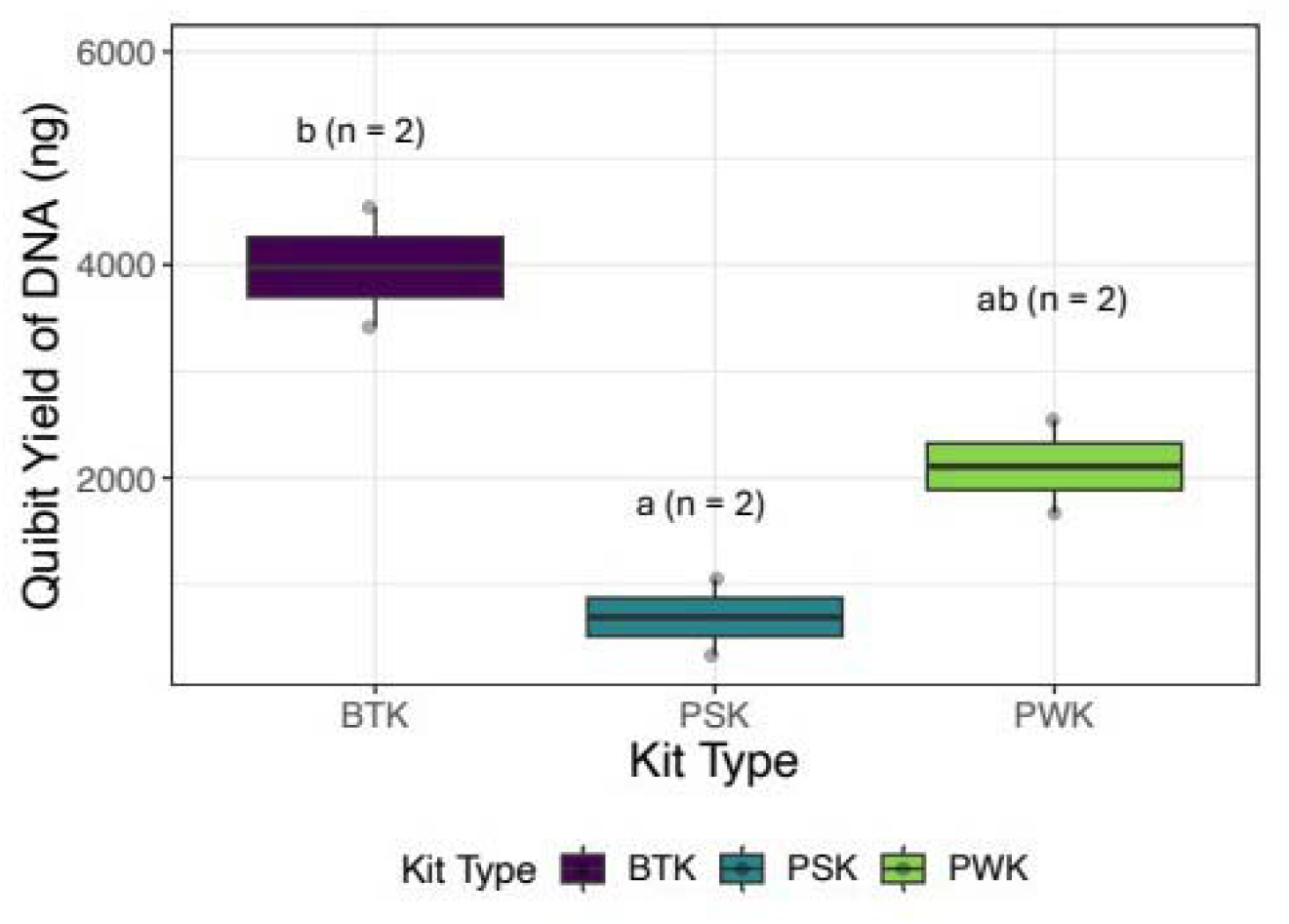
Comparative DNA yields from different extraction kits. Blood & Tissue Kit (BTK, purple), PowerSoil Kit (PSK, blue), and PowerWater Kit (PWK, green). The ng of DNA recovered (measured using a Qubit fluorometer) are shown for pico fractions (small micro) of a single sampling event on 26 August, 2024 during a high slack tide. The filters used were divided into three sections, then each third was extracted for DNA using a separate kit. Although the sample size is minimal, replicates for each extraction kit type are closely clustered, and letters indicate significance groups as denoted by a Tukey’s *post hoc* test (p < 0.05) after performing an analysis of variance (ANOVA) to determine if kit type had a significant impact on yield, while *n=2* indicates sample size in each plot. Samples labeled “ab” denote an overlap between groups a and b, indicating that DNA yields in these samples are not significantly different from either group a or group b, while groups a and b are significantly different from each other.

**Figure 6.**
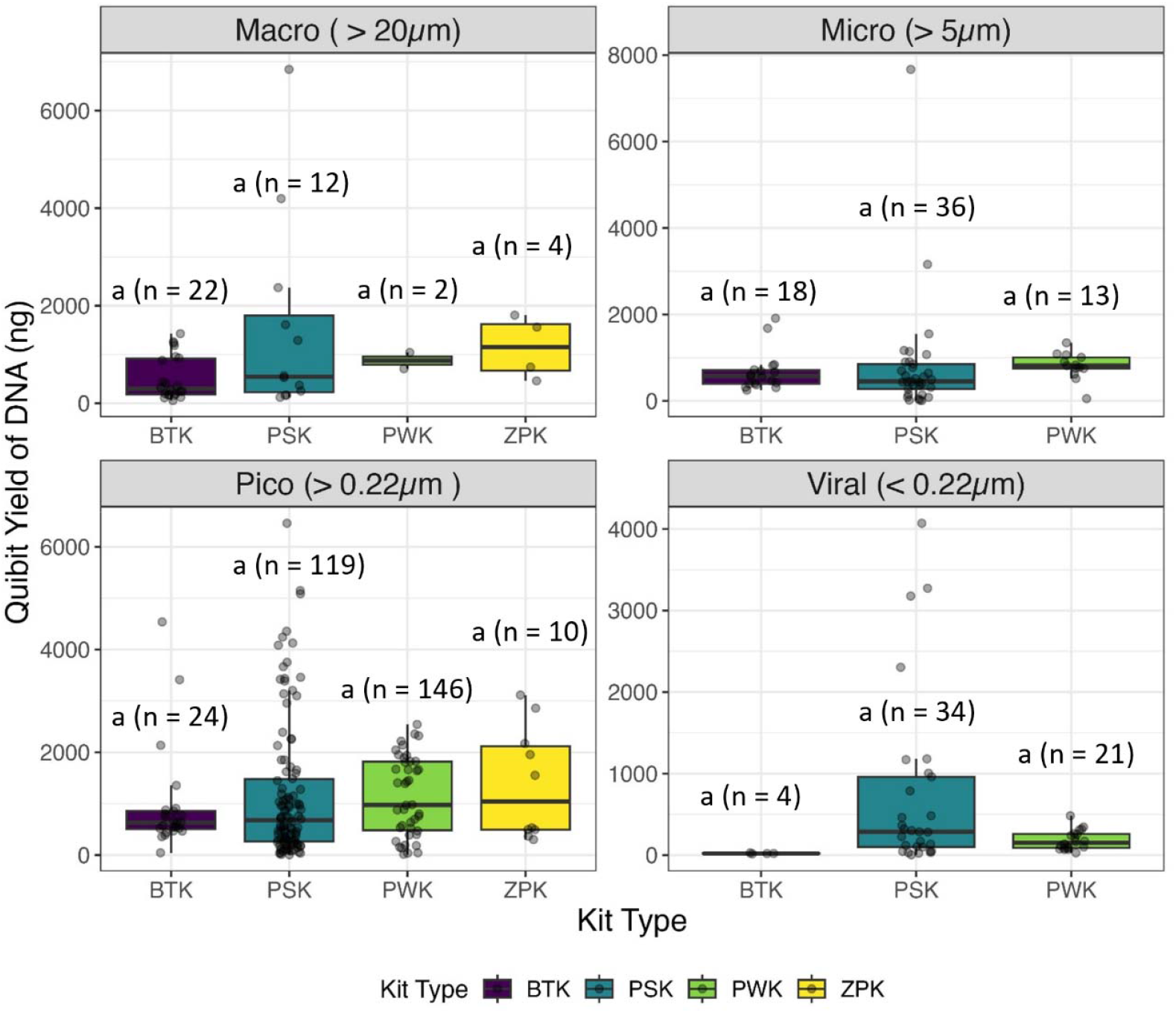
Kitchen sink comparison of DNA yields from different extraction kits for all filter fractions. The ng of DNA recovered (measured using a Qubit fluorometer) using the Blood & Tissue Kit (BTK, purple), PowerSoil Kit (PSK, blue), PowerWater Kit (PWK, green), or Zymo Plant/Seed (ZPK, yellow) across all PNNL-Sequim sampling events. Filter fractions included in this figure are: macro (top left), micro (top right), pico (bottom left) and viral (bottom right) fractions. All conditions are labelled “a” indicating that there was only one significance group as denoted by a Tukey’s *post hoc* test (p < 0.05) after performing an ANOVA to determine the extraction kit had a significant impact on yield, and n indicates sample size in each plot. These results include all samples from all PNNL-Sequim sampling events, filter types, and dates over the year. Variable conditions are not taken into account in each test other than extraction method.

A similar kitchen sink comparison (controlling only for filter pore size and location) of MCE versus PES membranes also had high variation in yield. While the MCE membranes yielded slightly more DNA than the PES membranes for the pico fraction (1260 ng average for pico MCE and 1143 ng average for pico PES), the difference was insubstantial compared to the variation among samples (Figure 7a). In a more controlled, exploratory comparison of DNA yields from samples, the MCE membranes in the pico fraction were significantly greater than the PES membranes (p < 0.05, 1818 and 1673 ng DNA, respectively), but the differences were small relative to the DNA yield (Figure 7b). Differences between PES and MCE in the viral fraction were also relatively small (324 ng difference in broad comparison, and 54 ng difference in focused comparison) and either were not significantly different or had overlapping significance groups based on a Tukey’s post hoc test (p < 0.05) after performing an ANOVA. Losses of yield when extracting DNA from filters after a long freezer storage period (over 100 days) were small and not significant (Figure 8).

**Figure 7.**
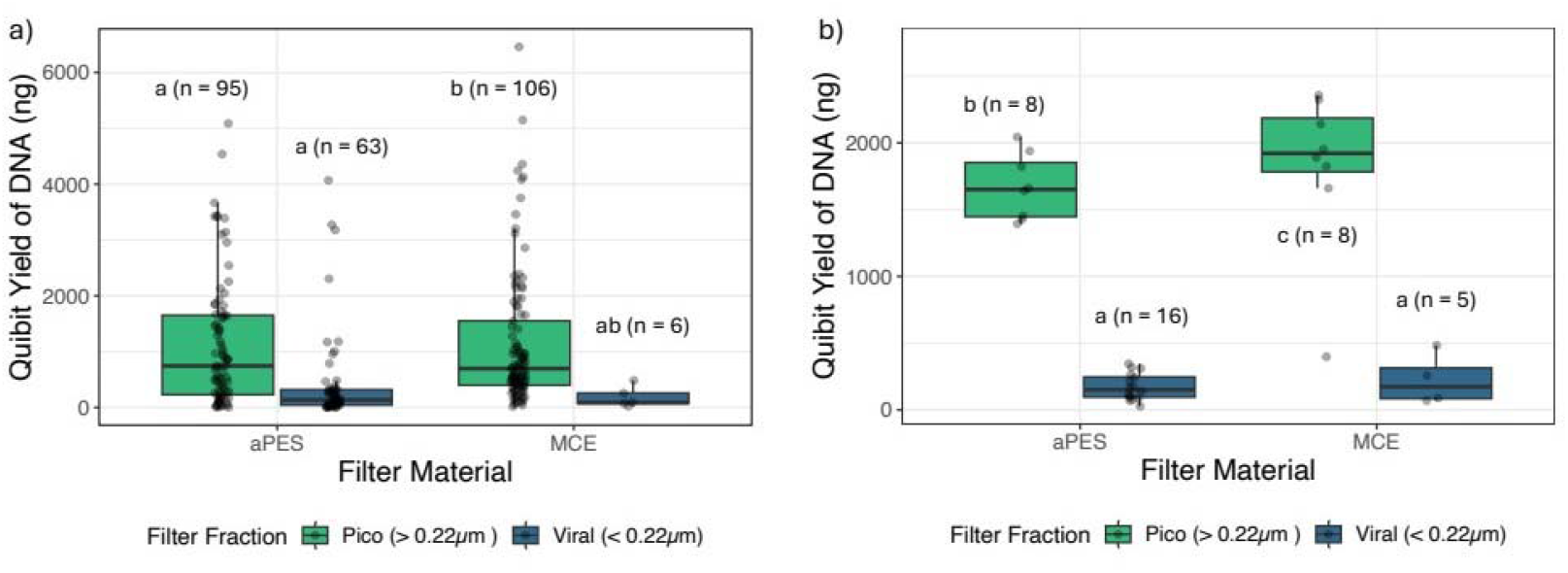
DNA yields for different filter types. The ng of DNA recovered (measured using a Qubit fluorometer) for pico (green) and viral (blue) filter fractions obtained using PES filters (left in each subfigure), and MCE membrane filters (right in each subfigure). A) Kitchen sink comparison of all filters collected from PNNL-Sequim or B) focused comparison of filters collected from a seawater mesocosm tank on 3 December, 2024. Lowercase letters inside graph area indicate significance groups as denoted by a Tukey’s post hoc test (p > 0.05) after performing an ANOVA to determine if material and fraction had a significant impact on yield. However, PNNL-Sequim dock samples included all pico and viral samples from multiple sampling events, methods, kits, and dates over the year. These variable conditions are not taken into account in this test. The samples from the mesocosm tank, in contrast, were from a singular sampling event on 3 December, 2024, and were extracted using the PowerWater kit within a week of sampling. Samples labeled ab” denote an overlap between groups a and b, indicating that DNA yields in these samples are not significantly different from either group a or group b, while groups a and b are significantly different from each other.

**Figure 8.**
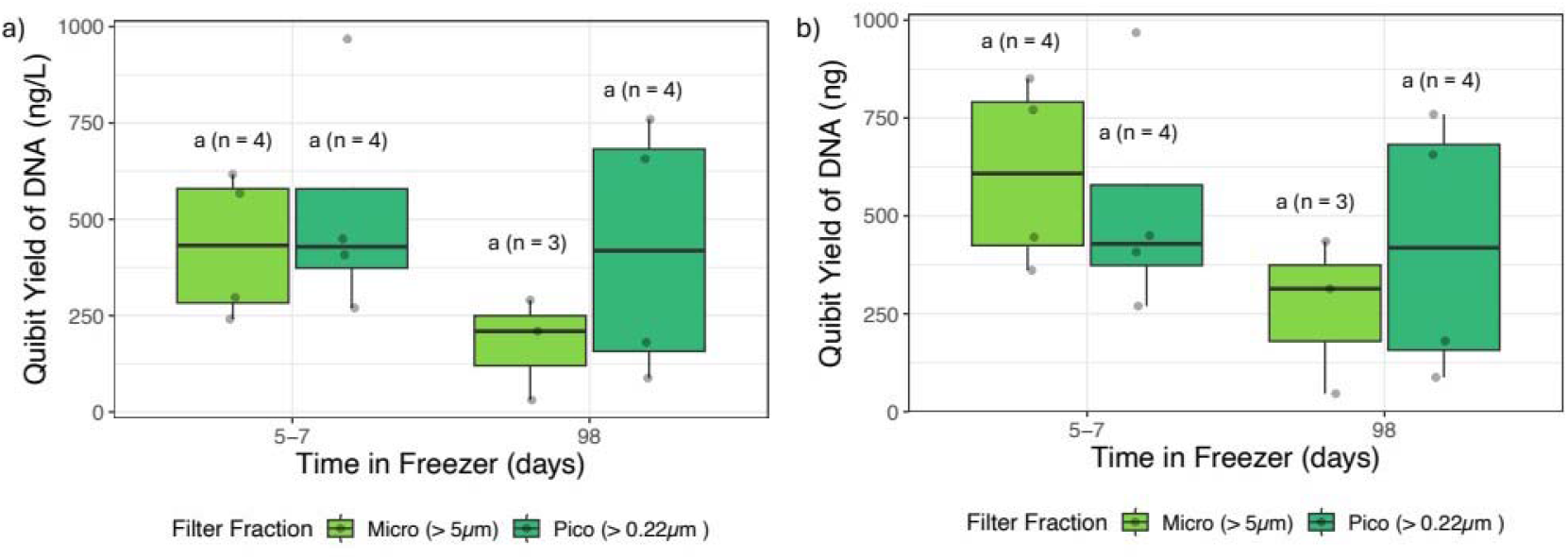
DNA yield before and after extended freezer storage. The ng of DNA recovered (measured using a Qubit fluorometer), presented as ng/L (a), and ng (b), with extractions taking place either 5-7 days after sampling (left plots within each figure) or 98 days after sampling (right plots within each figure). All data in this plot reflect DNA extracted from filters produced from a single sampling event on 31 July, 2024 extracted using the PowerSoil Kit and otherwise with the same processing methods. All conditions are labelled “a” indicating that there was only one significance group as denoted by a Tukey’s *post hoc* test (p < 0.05) after performing an ANOVA to determine if material and fraction had a significant impact on yield, and n indicates sample size in each plot.

In-line serial filtration of the macro and micro fractions onto nylon and MCE membranes, respectively, using the Millipore Sigma polypropylene in-line filter holders (see Methods and Supplemental Data Tables) was observed to be the fastest method to remove larger-sized organisms, and facilitated large-volume processing in a laboratory while monitoring and changing out 20 µm or 5 µm filters if flow lessened as they became clogged. We note that for this reason the replicates for the 5 µm filters are taken sequentially from the same sample flow through a single 20 µm filter, and may differ based on differential coating, clogging, or disruption of the 20 µm filter over that time series (see relative read counts section below). In contrast, the single-flask sterile PES filters were observed to be more time-efficient for pico and viral fractions than in-line filtration on an MCE membrane, reducing processing time per liter from over an hour to a matter of minutes in some cases. The larger (90 mm) filter size also allowed collection of quality technical replicates of DNA extraction and sequencing by cutting PES filters into equal fractions prior to freezing. A list of all yield data for samples used in these analyses can be found in Supplemental Data 3.

### Shotgun sequencing and relative read counts assigned to taxon groups

Read mapping results for four sites (PNNL-Sequim, John Wayne Marina, Cline Spit, and Discovery Bay) show that amoeba read counts were dramatically reduced between 3- and 30-fold after passage through 5 µm to 0.22 µm filters (Figure 9), with the greatest fold reduction from the highest count 5 µm filter (John Wayne Marina) and the least fold reduction from the lowest count 5 µm filter (Cline Spit). In the Cline Spit sequences, there were only 287 reads from the 5 µm filter that matched amoeba sequences. Although this indicates that large-bodied cells are effectively screened out, relative *Prochlorococcus* reads were not substantially increased or decreased in any of the 0.22 µm filter samples, while relative *Synechococcus* reads were substantially reduced in some 0.22 µm filter samples by up to a third (Figure 9 and Supplemental Data 1). *Prochlorococcus* normalized read counts were similar across sites and filters (range 718-1300 per 10 M reads), indicating that substantial numbers are trapped by the larger filters and possibly that the differential filtering of larger cells is not enough to increase the relative enrichment of small cells on the subsequent filter; other possibilities are discussed below. The coefficient of variation among *Prochlorococcus* mean-normalized read counts for the Figure 9 technical replicates (replicates of all steps from filtration onwards) on the 0.22 µm filters was also small at 6.8% (7.5%, 8.8%, and 5.0% for data in Figure 9a-c, respectively). The *Synechococcus* read counts were also similar across samples and showed slightly higher reads than those of *Prochlorococcus* with the exception of Discovery Bay, in which the *Synechococcus* read counts were approximately 10-fold higher than at Cline Spit (Figure 9c-d). The amoeba reads from the 5 µm filters at Cline Spit and Discovery Bay, were about 50%-100% greater than *Synechococcus* reads at PNNL-Sequim and John Wayne Marina. The number of reads mapping to amoeba and *Synechococcus* fluctuate much more than reads mapping to *Prochlorococcus*, but they are not in sync with each other across sites. It is also notable that the 24-mer nonpareil sequence diversity measures across all sites (Figure 9e) were consistently higher and more variable in the 5 µm filter fractions (range ∼20.5-22.5) than the 0.22 µm filter fractions (range ∼19.5-20). The 5 µm filters are thus at levels that are comparable to marine and soil samples (21-25) or freshwater and sandy soils (20-22) (Rodriguez-R, et al., 2018), while the 0.22 µm diversity is comparatively reduced.

**Figure 9.**
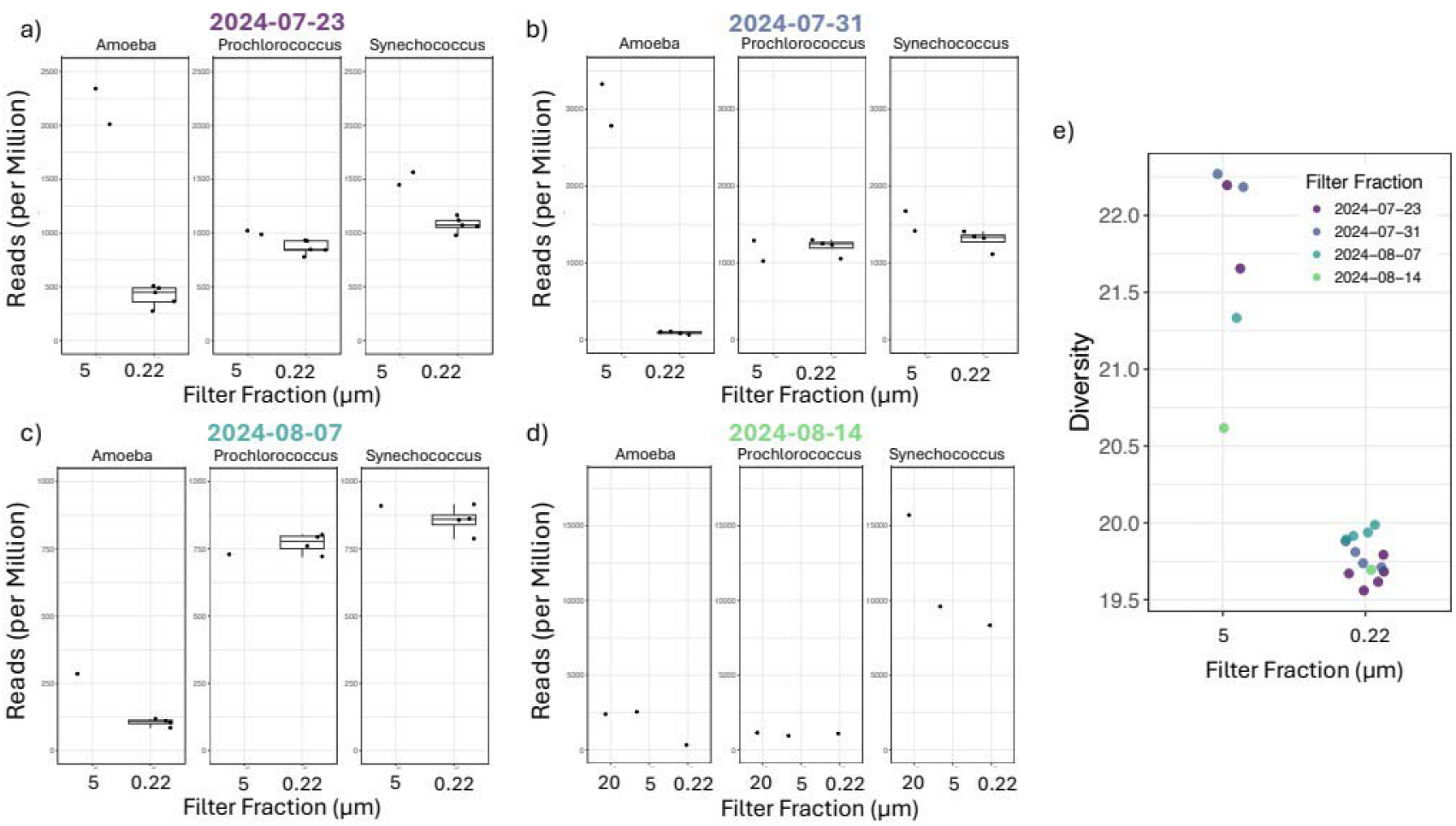
Taxon assignments and sequence diversity measures across sampling locations. Sampling occurred in mid-late Summer 2024 at four sites a) PNNL-Sequim 23 July 2024; b) John Wayne Marina 31 July 2024; c) Cline Spit 7 August 2024; and d) Discovery Bay 14 August 2024. e) Nonpareil 24-mer diversity measures (log scale) of the same samples shown in a-d, color-coded by site and sampling date as indicated in legend. Water samples were collected passing through inline 20 µm and 5 µm filters, then mixed in a flask and filtered through a 0.22 µm filter. DNA was extracted from biological material on the filter sizes labelled on the graphs. The number of reads observed to match to genomes in each taxon group (amoeba, *Prochlorococcus, and Synechococcus) were norm*alized to the total reads in each sample and shown in units of reads per million. The magnitudes of the scales vary among sample sites but are constant within sites to allow visualization of the relative read counts between taxon groups and to directly compare the change in relative read counts between different filter fractions within a sample (PNNL-Sequim scale max 2500; John Wayne Marina scale max 3500; Cline Spit scale max 1000; Discovery Bay scale max 17,500). All samples were filtered with 20 µm nylon membrane filters and 5 and 0.22 µm MCE membrane filters where applicable, except the Cline Spit and Discovery Bay sample were filtered with 5 µm nylon membranes and the Discovery Bay sample was filtered with a 0.22 µm PES membrane. All samples were collected on flood tides except the PNNL-Sequim samples, which were collected on an ebb tide. All conditions with multiple points are technical replicates of the process from sample collection and filtration through DNA extraction and sequencing process. The 5 µm filters are sequential in-line filters through the prior 20 µm filters, while the 0.22 µm filters are repeat filters of the mixed 5 µm filtrate.

*Synechococcus read counts* were remarkably similar at ebb and flood tides for both the 5 µm and 0.22 µm filters (Figure 10). In contrast, amoeba reads were about 3x less frequent at flood versus ebb tide on the 5 µm filter and greatly reduced in frequency on the 0.22 µm filters (Figure 10). *Prochlorococcus reads were* also reduced in flood versus ebb tides and were reduced by 50%-60% on the 0.22 µm filter compared to the 5 µm filter. We note that the flood tide samples were extracted considerably later than the ebb stage samples, but time stored in freezer prior to extraction was not expected to change relative frequencies based on our exploratory analysis showing no significant impact of freeze-time on yield (Figure 8).

**Figure 10.**
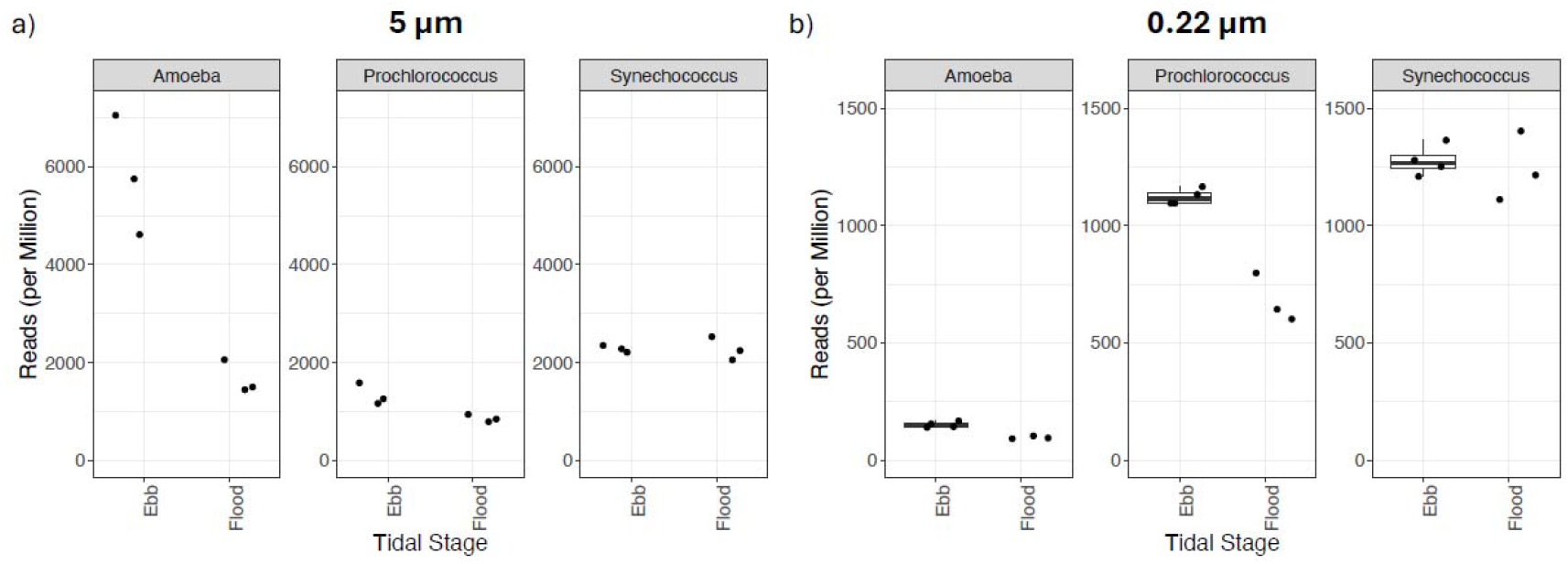
Taxon assignments of same-day samples collected at PNNL-PNNL at Ebb and Flood tides. All samples were collected on 31 July 2024 at dawn or midday (ebb and flood tides, respectively), filtered with 5 and 0.22 µm MCE membrane filters and extracted with Qiagen PowerSoil kits. Ebb samples were extracted after 5-7 days in the freezer, while flood samples were extracted after 98 days in the freezer. For readability, the magnitudes of the scales differ between the two filter pore sizes, with 5.0 µm scale max 8000 reads per million and 0.22 µm scale max 1500 reads per million. All conditions with multiple points are technical replicates of the process from sample collection and filtration through DNA extraction and sequencing process. The 5 µm filters are sequential in-line filters through the prior 20 µm filters, while the 0.22 µm filters are repeat filters of the mixed 5 µm filtrate.

Relative frequencies of microbial taxa on the 0.22 µm filters over time at PNNL-Sequim, which lasted from mid-April to late October 2024, are shown in Figure 11. These comparisons unavoidably included variation in tidal stage, and also in extraction method and time between collection and extraction, and are across a single year (see Fig 11 legend), so must be interpreted in that context. As in previous figures, the amoeba reads are generally low, indicating relative exclusion by the 5 µm filter. Amoeba and *Prochlorococcus reads had h*igh relative frequencies in early spring and late fall, indicating that they were relatively more frequent than other organisms in the filter sample). In contrast, *Synechococcus counts were* higher from April to October, with the exception of counts in July. Counts on July 17, 2024 were clustered by processing conditions for all three taxon groups, indicating that the combination of kit and time in freezer prior to extraction can affect results at nearly the same level of variation as change over time (PSK kit and short time in freezer had much higher counts). September 9, 2024 amoeba samples were strongly clustered by condition, with high relative counts for all samples with a short freezer storage duration and extracted with the PWK kit (the Prochlorococcus samples were also clustered in the same fashion). Across all three plots, DNA samples extracted within a few days of filtration had consistently clustered higher counts than samples extracted greater than 50 days after filtration (stored frozen and thawed). This suggests that extended freezer time may contribute to decreased identifiable sequence reads, yet our focused comparisons to assess the impact of freezer storage duration on DNA yield found no significant effect (Figure 8).

**Figure 11.**
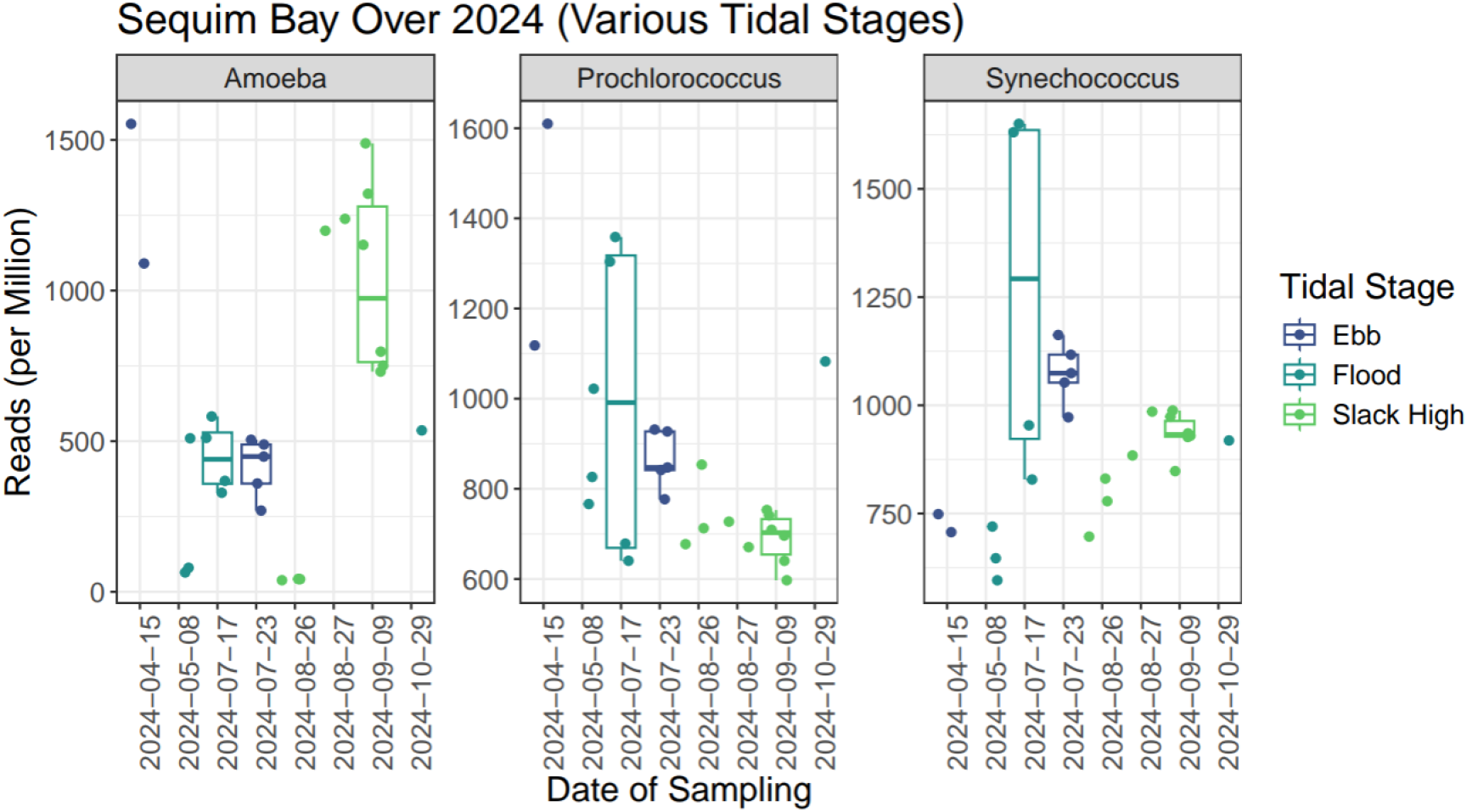
Diversity reads (per million) over six months in 2024 for all pico fraction samples from the PNNL-Sequim floating dock. PNNL-Sequim dock samples included samples from multiple sampling events, methods, kits, and dates over the year. All samples were extracted using either the Qiagen DNeasy PowerSoil Kit (PSK) or Qiagen DNeasy PowerWater Kit (PWK). 8 distinct sampling events on 8 different days are captured in this figure: **April 15** (n = 2 DNA samples extracted with PSK 3 days after filtration), **May 8** (n = 3 DNA samples extracted using PSK 5 days after filtration), **July 17** (n = 4 DNA samples; 2 extracted using PSK 1 day after filtration, and 2 extracted using PWK 124 days after filtration), **July 23** (n = 5 samples extracted using PSK 13 days after filtration), **July 31** (n = 4 DNA samples extracted using PSK 5 days after filtration), **August 26** (n = 1 DNA sample extracted using PWK 9 days after filtration), **September 9** (n = 6 DNA samples; 3 extracted using PWK 2 days after sampling, and 3 extracted using PSK 58 days after filtration), and **October 29, 2024** (n = 1 DNA sample extracted using PSK 8 days after sampling). 3 sampling events occurred during an ebb tide (purple; April 15, July 23, and July 31, 2024), 2 occurred during a flood tide (blue; May 8, July 17, and October 29, 2024), and 2 during a high slack tide (green; August 26 and September 9, 2024). Note that for the July 17 samples, the relative counts cluster by condition for all three taxon groups (higher in PSK 1 day after filtration always have the higher count), while for the September 9 samples the amoeba cluster strongly by condition (higher in PWK 2 days after sampling versus PSK 58 days after sampling); the *Prochlorococcus are all sli*ghtly higher in the September 9 PWK extracted 2 days after sampling.

## Discussion

In this paper, we present and test the efficacy of a sampling approach to enrich taxonomic fractions by size exclusion filtration prior to targeted metagenomic shotgun sequencing. Our prior expectation was that organisms with cell sizes in the given size range would be enriched in the corresponding filter fraction, but that results would vary due to variable filter performance, variable cell-cell and cell-particle attachment frequencies among organisms, cell-free DNA, and horizontal gene transfer, especially involving viral particles. We define our sample fractions as macro (greater than 20 um), micro (5 to 20 um), pico (aka small micro, 0.2 to 5 um), and viral (less than 0.2 um), and sampling occurred in conjunction with abiotic metadata collection from the environment. Our final methodology balances material recovery, filter type and size, and collection speed. It incorporates a multi-step process: inline serial filtration through a combination of filters with sequentially smaller pore sizes, followed by single-flask filtration to capture various target organism size classes, followed by precipitation of viral particles in the filtrate from the smallest 0.22 µm filters, which are expected to screen out nearly all intact cells. The final filter types were 20 µm Millipore nylon filters, 5 µm Millipore MCE membrane filters, and 0.22 µm PES filters. By increasing the filter volume processed when there was little visible biological material, we successfully produced ≥ 50 ng of DNA in the majority of extractions, our minimal cutoff for sequencing submission. This required extensive filtration processing time of up to 3 hours in some cases with higher biomass levels. All kit and filter types tested yielded sufficient DNA quantity and quality, and the more critical factors were processing time, cost, and material availability. The sequencing results presented in this paper were focused on exploratory (many conditions had small numbers or no replicates) statistical analysis to demonstrate detection of the picocyanobacteria genera *Prochlorococcus* and *Synechococcus*, the times of year they were detectable, whether we could enrich the number of reads for these genera in the pico fractions, the technical repeatability of the results, and whether we could detect enrichment for sequences homologous to viruses that might be associated with picocyanobacteria in viral precipitate fractions.

The clearest and perhaps most important sequencing result was that both *Prochlorococcus* and *Synechococcus* were detected in the Salish Sea samples, usually but not always with similar counts between them. *Synechococcus* was generally more common, especially in Discovery Bay which had dramatically higher *Synechococcus* counts per million. Detection of picocyanobacteria sequences is important because extraction of high-quality DNA from picocyanobacteria remains challenging primarily due to relatively low biomass collected in competitive and low-nutrient (oligotrophic) marine water (Wagner, et al., 2024; Hept & Greene, 2023). complicated by their potential inability to contribute proportionally as much to biomass in the more competitive and nutrient-rich estuarine waters of the Salish Sea. The relatively constant ratio of sequences from the two genera across sites and over the year is also important because of possible temperature, nutrient level, and virus interaction effects that at this latitude could have strongly affected their relative distributions (Carlson et al., 2022; Coello-Camba & Agustí, 2024; Zhang et al., 2022; Zufia et al., 2022). The validation of picocyanobacterial relative type abundance was essential because while some studies determined that picocyanobacteria are found in all regions of the euphotic zone, there is little research to validate these findings in the northern Pacific Ocean and specifically in the Salish Sea (Babić, et al., 2017). It is also notable, and important to establish, that the technical replicability (from filtering onwards) of the relative Prochlorococcus counts on the 0.22 µm filters was high (Figure 9 and throughout), allowing preliminary confidence in results differing by 10% or higher.

We note that although one viral sample had anomalously low counts, the top 5 viral samples have only 3.5x higher relative read counts (average 3417) than the remainder of the samples (average 882), indicating that a substantial amount of viral DNA is retained with the large-sized fractions. This may be due to particles or free DNA sticking to the filters, or to incorporation/infection or attachment to the cells that are stuck on those filters. We also note that phages derived from *Synechococcus* were the best matches to slightly over 50% of the cyanophage reads, although we caution against over-interpreting this result in terms of what types of organisms are infected with cyanophage, both because phages can often infect multiple species, and because the reference dataset contains many more *Synechococcus*-derived phages than cyanophages isolated from other organisms. Nevertheless, it is interesting that some of the highest read counts observed for *Synechococcus*-derived phages (2698, 2696, and 1240 respectively on the 20, 5 and 0.22 µm filters from August 14, 2024 at Discovery Bay) were from samples with some of the highest *Synechococcus* genome read counts (respectively 15,251, 9653, and 7989).

Salinity and water temperature followed similar trends to each other, with slight increases during summer months and lows during cooler seasons. The slight changes in salinity across seasons are likely due to the heavy influence of rivers and precipitation in the estuary system, with increased outflow and precipitation in colder months that decrease salinity (Walker, et al., 2022). Cline Spit’s increased water temperature compared to PNNL-Sequim may be influenced by its location within Dungeness Bay, which is protected by a long spit that reduces water volume exchange and creates shallower sites. There are also large variances in the distance from the shore between sampling sites, which may inform differences betweensamples (e.g. sampling less than one meter from shore at Cline Spit, and greater than 50 meters from shore near Port Townsend).

The trends in relative frequencies of *Prochlorococcus, Synechococcus*, and amoeba-like sequences over the course of the year at PNNL Sequim (Figure 11) were somewhat surprising. Although the numbers were small, relative numbers of *Prochlorococcus* were highest towards the beginning of the year and decreased throughout the year, while *Synechococcus* numbers were lowest early in the year and then relatively constant. One might have thought that *Prochlorococcus*, which dominates at tropical and subtropical altitudes, would do comparatively better in the warmer months. Indeed, the highest counts were expected during the late spring and summer months (Zinser, et al., 2006), so this may indicate that the estuarine dynamics of an inland sea are more unpredictable than expected. This trend was visible despite differences in tidal stage at time of sampling (which also impact relative counts, Figure 10). The trends in relative amoeba-like read counts (we remind readers that with only one amoeba sequence included, the phylogenetic definition of these sequences is poor) did not correspond to either of the picocyanobacteria, being most common in the earliest and latest months of the year sampled. Further definition of these trends will await results from planned sampling and sequencing across 2025 and 2026, and will particularly benefit from more detailed analysis of the relative frequency of sub-lineages within these broad taxonomic groups (i.e. at the species or ecotype level or below).

The relative read counts of various taxa under different filtration conditions were also unexpected. The amoeba-like sequences were more frequent on the 5 µm filters than the 0.22 µm filters, and combined with the relatively high nonpareil diversity on the 5 µm filters, this indicated that the size exclusion filtration was effective. However, *Prochlorococcus* and *Synechococcus* relative read counts were not comparatively higher on the 0.22 µm filters, and were fairly frequent in the viral fraction, which passed through the 0.22 µm filters. Given these results, we may have been overly conservative in using 5 µm filters (rather than, say, 2 µm filters) to screen out large-bodied cells, but it also appears that considerable amounts of picocyanobacterial DNA both accumulated on the larger filters and passed through the smaller filters. Although free DNA and inadequacies of the filtration process need to be considered (e.g., high variability in pore size, micro breaks in filters, DNA or cell binding to filters as they pass through), it is possible that there are alternative explanations. Tycheposons, which are numerically common DNA-containing particles from *Prochlorococcus*, are emitted into the environment as particles similar to viruses (Hackl, et al., 2023). These and other undiscovered elements in the taxa under study may strongly impact the filtration properties of DNA-containing objects. Higher-level conglomerates of cells or cell-particle mixes are now also recognized as important features of both *Prochlorococcus* and *Synechococcus*, which would contribute to their retention on larger pore-size filters. Furthermore, viruses are commonly retained with infected cells, and due to gene transfer mechanisms both viruses and host targets can contain each other’s genes. These sequencing features will also strongly benefit from more detailed analysis of the relative frequency of sub-lineages within these broad taxonomic groups, as well as detailed analysis of genomic regions that may be under- or over-sampled in different filter partitions.

### Technical Challenges

The extraction of high-quality DNA from picocyanobacteria remains challenging, likely due to relatively low biomass collected in oligotrophic marine water, and potentially due to environmental contaminants, cell lysis inefficiencies due to cyanobacterial extracellular polymeric substances, and polysaccharide production (Wagner, et al., 2024). Additionally, the consistently low biomass found at sampling sites— excluding Cline Spit, which had much higher biological material than the other sampling sites— necessitated high filtration volumes and extensive processing time.

For our preliminary estimates, we used limited high-level estimates of sequence composition, and small numbers of technical replicates from identical samples (same time, location, and water sample with same filtration protocol). We also had few replicates of biological/environmental effects such as tidal state, time of year, or sampling location. Due to continuous sampling methodology adaptations throughout year one, there was high variation in sample production, which significantly decreased the number of quality technical replicates for focused comparisons. Especially for the non-controlled comparisons, results should be validated with larger sample sets and using consistent methods; however, the results presented here provide a valuable understanding of the impact that methodological variation can have.

### Environmental Implications and Future Directions

We aimed to establish standardized methods for consistent size-fractionated sampling and metadata collection to support host-phage interaction studies influenced by environmental factors and to provide shareable, high-quality metadata for diverse applications. By analyzing community composition, species, and ecotypes at the community level, as well as variations in host and phage genes, protein structures, and interactomes at the molecular level, this approach may inform phage-host co-evolution and viral-host interactions while reducing sequencing noise. This refined sampling and metadata production method for downstream analyses may also be useful in environmental DNA pre-filtration, ecosystem health monitoring, phytoplankton carbon cycling, harmful algal bloom detection, water quality assessments, metagenomics, single-cell genomics, aquaculture, algal anti-viral research, and fisheries management. More data collection using the final sampling scheme will produce valuable, standardized biological and physiochemical datasets to feed into in-depth research applications such as evolutionary modeling of phage-host interactions.

## Supplemental Data

**Supplemental Figure 1.**
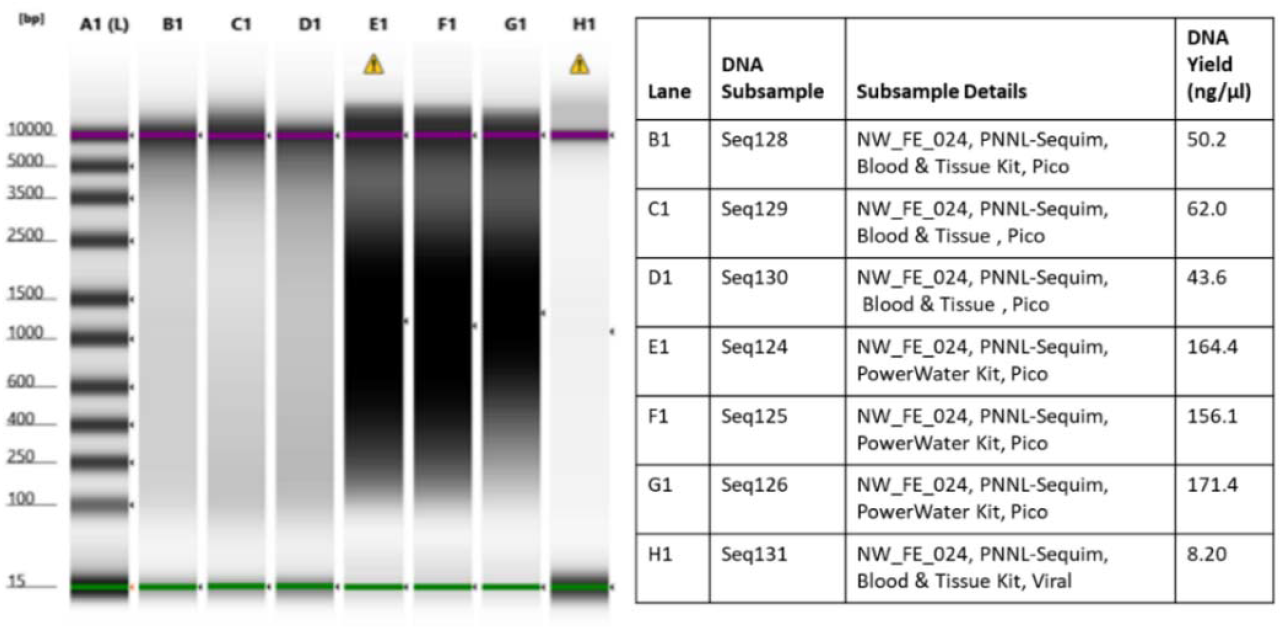
TapeStation Data: DNA fragment analysis of 6 pico fraction samples and 1 viral fraction sample from parent sample NW_FE_024, sampled from Sequim Bay during a slack tide in September of 2024.

**Supplementary Table 1.** *Peristaltic and Vacuum Pumps* used in sampling method iteration.

| Pump | Catalog Number | Purchased From |
| --- | --- | --- |
| Masterflex Standard Digital PTFE-Tubing Pump System | MFLX77912-10 | Avantor Link |
| Watson-Marlow 323S Variable-Speed Pump Drive, Manual Control, 3 to 400 rpm | UX-50130-20 | Cole-Palmer Link |
| eDNA Citizen Scientist Sampler | n/a | Smith-Root Link |

**Supplementary Table 2.** *Filter types* tested for serial filtration optimization.

| Pore Size (μm) | Material | Diameter (mm) | Purchased From | Filter Fraction(s) Used In |
| --- | --- | --- | --- | --- |
| 20 | nylon | 47 | Millipore Sigma Link | Macro (MaF) |
| 5.0 | Mixed Cellulose Ester (MC)E hydrophilic membrane | 47 | Millipore Sigma Link | Micro (MiF) |
| 0.2 | PES filter in sterile filter unit | 90 | VWR Link | Pico (PF), Viral (VF) |
| 0.22 | Mixed Cellulose Ester (MC)E hydrophilic membrane | 47 | Millipore Sigma Link | Pico (PF) |
| 0.05 | Mixed Cellulose Ester (MC)E hydrophilic membrane | 47 | Millipore Sigma Link | Viral (VF) |

**Supplementary Table 3.** *In-line filter holders* tested for serial filtration optimization.

| Description | Material | Holds Filter Diameter | Purchased From | Filter Fraction(s) |
| --- | --- | --- | --- | --- |

|  |  | er (mm) |  | Used In |
| --- | --- | --- | --- | --- |
| In-Line Filter Holder | Polycarbonate | 47 | Avantor Sciences, Cytiva Link | In-line macro, micro |
| In-Line Filter Holder | Polypropylene (nylon knobs and silicone seal) | 47 | Millipore Sigma Link | In-line macro, micro |

**Supplementary Table 4.** *DNA* Extraction kits used to test filter subsample DNA extraction optimization.

| Kit | Catalog Number | Purchased From |
| --- | --- | --- |
| DNeasy PowerSoil Pro Kit | 47014 | Qiagen Link |
| DNeasy PowerWater Kit | 14900-100-NF | Qiagen Link |
| DNeasy Blood and Tissue Kit | 69504 | Qiagen Link |
| Quick-DNA Plant/Seed Miniprep Kit | D620 | Zymo Research Link |

**Supplementary Table 5.** *Environmental metadata:* acquisition methods distinguished by data acquired from MCRL Data at PNNL-Sequim or other locations.

| Parameter | Unit | PNNL-Sequim Sampling Events | Non PNNL-Sequim Sampling Events |
| --- | --- | --- | --- |
| Water level | m | MCRL Data – Tide Guage |  |
| Water temperature | °C | MCRL Data – CTD Sensor | YSI ProDSS – Cond./Temp. Sensor |
| Dissolved oxygen | mg/L | MCRL Data – CTD Sensor | YSI ProDSS – ODO Sensor |
| Salinity | ppt | MCRL Data – CTD Sensor | YSI ProDSS – Cond./Temp. Sensor |
| Air temperature | °C | MCRL Data – CTD Sensor |  |
| Barometric pressure | Pa | MCRL Data – Met Sensor |  |
| Relative humidity | % | MCRL Data – Met Sensor |  |
| Rain | mm | MCRL Data – Met Sensor |  |
| Wind direction | degrees from | MCRL Data – Wind Station |  |
| Average windspeed | m/s | MCRL Data – Wind Station | Handheld anemometer |
| Max windspeed | m/s | MCRL Data – Wind Station | Handheld anemometer |
| Average land PAR | 1/4mol of photons m<sup>-2</sup> s<sup>-1</sup> | MCRL Data – PAR Sensor |  |
| Underwater lower PAR | 1/4mol of photons m<sup>-2</sup> s<sup>-1</sup> | MCRL Data – PAR Sensor |  |
| Underwater upper PAR | 1/4mol of photons m<sup>-2</sup> s<sup>-1</sup> | MCRL Data – PAR Sensor | LiCor Spherical Underwater Sensor |
| Chlorophyll | 1/4g/L | MCRL Data – Water Quality Sensors |  |
| Phycoerythrin | ppb | MCRL Data – Water Quality Sensors | YSI ProDSS – Total Algae Sensor, PE |
| Colored dissolved organic matter | ppb | MCRL Data – Water Quality Sensors |  |
| Turbidity | FNU | MCRL Data – Water Quality Sensors | YSI ProDSS – Turbidity Sensor |
| Cloud cover |  | Field crew observations | Field crew observations |
| Sea state |  | Field crew observations | Field crew observations |
| Secchi Depth | m | Secchi Disk | Secchi Disk |
| Tidal Stage | Ebb/Flood/Slack | NOAA Tides and Currents | NOAA Tides and Currents |

