## Supplementary Data 1 for "Sampling Microbial Dynamics in the Salish Sea Estuary: Evaluating Methods to Capture Cyanobacteria and Cyanophage"

| Parent.ID | Extraction. | Date.of.Ex | Date.of.Sa | Time.in.Fr | KitName | DNA.ID | Filter.Usec | Total.volur |
|---|---|---|---|---|---|---|---|---|
| NW_FE_005 | #011 | 45400 | 45397 | 3 | Qiagen DN | 20240418- | BF-013 | 2 |
| NW_FE_005 | #013 | 45400 | 45397 | 3 | Qiagen DN | 20240418- | BF-015 | 1 |
| NW_FE_006 | #045 | 45425 | 45420 | 5 | Qiagen DN | 20240513- | BF-022 | 2.1 |
| NW_FE_006 | #049 | 45425 | 45420 | 5 | Qiagen DN | 20240513- | BF-026 | 2 |
| NW_FE_006 | #050 | 45425 | 45420 | 5 | Qiagen DN | 20240513- | BF-027 | 1.3 |
| NW_FE_011 | #058 | 45491 | 45490 | 1 | Qiagen DN | 20240718- | BF-064 | 1.8 |
| NW_FE_011 | #059 | 45491 | 45490 | 1 | Qiagen DN | 20240718- | BF-065 | 1 |
| NW_FE_012 | #075 | 45509 | 45496 | 13 | Qiagen DN | 20240805- | BF-067 | 5 |
| NW_FE_012 | #076 | 45509 | 45496 | 13 | Qiagen DN | 20240805- | BF-069 | 5 |
| NW_FE_013 | #077 | 45509 | 45504 | 5 | Qiagen DN | 20240805- | BF-075 | 2 |
| NW_FE_013 | #078 | 45509 | 45504 | 5 | Qiagen DN | 20240805- | BF-076 | 2 |
| NW_FE_013 | #079 | 45509 | 45504 | 5 | Qiagen DN | 20240805- | BF-077 | 2 |
| NW_FE_013 | #080 | 45509 | 45504 | 5 | Qiagen DN | 20240805- | BF-078 | 2 |
| NW_FE_012 | #082 | 45509 | 45496 | 13 | Qiagen DN | 20240805- | BF-070 | 1.5 |
| NW_FE_012 | #083 | 45509 | 45496 | 13 | Qiagen DN | 20240805- | BF-071 | 1.5 |
| NW_FE_012 | #084 | 45509 | 45496 | 13 | Qiagen DN | 20240805- | BF-072 | 1.5 |
| NW_FE_012 | #085 | 45509 | 45496 | 13 | Qiagen DN | 20240805- | BF-073 | 1.5 |
| NW_FE_012 | #086 | 45509 | 45496 | 13 | Qiagen DN | 20240805- | BF-074 | 1.5 |
| NW_FE_015 | #094 | 45510 | 45504 | 6 | Qiagen DN | 20240806- | BF-087 | 2.5 |
| NW_FE_015 | #095 | 45510 | 45504 | 6 | Qiagen DN | 20240806- | BF-088 | 2.5 |
| NW_FE_015 | #102 | 45511 | 45504 | 7 | Qiagen DN | 20240807- | BF-094 | 2 |
| NW_FE_015 | #103 | 45511 | 45504 | 7 | Qiagen DN | 20240807- | BF-095 | 2 |
| NW_FE_015 | #104 | 45511 | 45504 | 7 | Qiagen DN | 20240807- | BF-096 | 2 |
| NW_FE_015 | #105 | 45511 | 45504 | 7 | Qiagen DN | 20240807- | BF-097 | 2 |
| NW_FE_013 | #106 | 45511 | 45504 | 7 | Qiagen DN | 20240807- | BF-099 | 3 |
| NW_FE_013 | #107 | 45511 | 45504 | 7 | Qiagen DN | 20240807- | BF-100 | 3.5 |
| NW_FE_013 | #108 | 45511 | 45504 | 7 | Qiagen DN | 20240807- | BF-101 | 2.5 |
| NW_FE_022 | #114 | 45533 | 45530 | 3 | Qiagen DN | 20240829- | BF-172 | 20 |
| NW_FE_022 | #119 | 45539 | 45530 | 9 | Qiagen DN | 20240829- | BF-172 | 20 |
| NW_FE_022 | #120 | 45539 | 45530 | 9 | Qiagen DN | 20240829- | BF-173 | 20 |
| NW_FE_023 | #121 | 45539 | 45531 | 8 | Qiagen DN | 20240829- | BF-178 | 20 |
| NW_FE_023 | #122 | 45539 | 45531 | 8 | Qiagen DN | 20240829- | BF-179 | 20 |
| NW_FE_024 | #124 | 45546 | 45544 | 2 | Qiagen DN | 20240911- | BF-187b | 19 |
| NW_FE_024 | #125 | 45546 | 45544 | 2 | Qiagen DN | 20240911- | BF-188b | 19 |
| NW_FE_024 | #126 | 45546 | 45544 | 2 | Qiagen DN | 20240911- | BF-189b | 19 |
| NW_FE_024 | #133 | 45602 | 45544 | 58 | Qiagen DN | 20241106- | BF-187c | 19 |
| NW_FE_024 | #134 | 45602 | 45544 | 58 | Qiagen DN | 20241106- | BF-188c | 19 |
| NW_FE_024 | #135 | 45602 | 45544 | 58 | Qiagen DN | 20241106- | BF-189c | 19 |
| NW_FE_014 | #136 | 45602 | 45504 | 98 | Qiagen DN | 20241106- | BF-081 | 3 |
| NW_FE_014 | #137 | 45602 | 45504 | 98 | Qiagen DN | 20241106- | BF-082 | 3 |
| NW_FE_014 | #138 | 45602 | 45504 | 98 | Qiagen DN | 20241106- | BF-083 | 3 |
| NW_FE_014 | #142 | 45602 | 45504 | 98 | Qiagen DN | 20241106- | BF-090 | 2 |
| NW_FE_014 | #143 | 45602 | 45504 | 98 | Qiagen DN | 20241106- | BF-091 | 2 |
| NW_FE_014 | #144 | 45602 | 45504 | 98 | Qiagen DN | 20241106- | BF-092 | 2 |
| NW_FE_025 | #148 | 45602 | 45594 | 8 | Qiagen DN | 20241106- | BF-201a | 40 |
| NW_FE_025 | #149 | 45602 | 45594 | 8 | Qiagen DN | 20241106- | BF-202a | 3 |

| | | | | | | | | |
|---|---|---|---|---|---|---|---|---|
| NW_FE_025 | #150 | 45602 | 45594 | 8 | Qiagen DN | 20241106- | BF-203a | 10 |
| NW_FE_025 | #151 | 45602 | 45594 | 8 | Qiagen DN | 20241106- | BF-204a | 14 |
| NW_FE_025 | #152 | 45602 | 45594 | 8 | Qiagen DN | 20241106- | BF-205a | 17 |
| NW_FE_025 | #153 | 45602 | 45594 | 8 | Qiagen DN | 20241106- | BF-206a | 12 |
| NW_FE_017 | #168 | 45614 | 45511 | 103 | Qiagen DN | 20241118- | BF-124 | 6 |
| NW_FE_017 | #169 | 45614 | 45511 | 103 | Qiagen DN | 20241118- | BF-125 | 0.425 |
| NW_FE_017 | #170 | 45614 | 45511 | 103 | Qiagen DN | 20241118- | BF-126 | 0.425 |
| NW_FE_017 | #171 | 45614 | 45511 | 103 | Qiagen DN | 20241118- | BF-130 | 0.45 |
| NW_FE_017 | #172 | 45614 | 45511 | 103 | Qiagen DN | 20241118- | BF-131 | 0.45 |
| NW_FE_017 | #173 | 45614 | 45511 | 103 | Qiagen DN | 20241118- | BF-132 | 0.16 |
| NW_FE_019 | #174 | 45614 | 45518 | 96 | Qiagen DN | 20241118- | BF-137 | 7.3 |
| NW_FE_019 | #175 | 45614 | 45518 | 96 | Qiagen DN | 20241118- | BF-138 | 7.3 |
| NW_FE_019 | #176 | 45614 | 45518 | 96 | Qiagen DN | 20241118- | BF-144 | 5 |

| X..Of.Who | Filtered.Vo | Sequim.Na | Sequim.Na | Sequim.A2 | Sequim.A2 | Sequim.Elu | Sequim.Sa | Sequim..Re | Sequim.Na |
|---|---|---|---|---|---|---|---|---|---|
| 0.5 | 1 | 27.6 | 0.0276 | 1.84 | 0.99 | 54 | 4 | 50 | 1.38 |
| 0.5 | 0.5 | 15.5 | 0.0155 | 1.87 | 0.9 | 54 | 4 | 50 | 0.775 |
| 0.5 | 1.05 | NA | NA | NA | NA | NA | NA | 96 | NA |
| 0.5 | 1 | NA | NA | NA | NA | NA | NA | 96 | NA |
| 0.5 | 0.65 | NA | NA | NA | NA | NA | NA | 96 | NA |
| 0.5 | 0.9 | 32.7 | 0.0327 | 1.82 | 0.23 | 100 | 4 | 96 | 3.1392 |
| 0.5 | 0.5 | 16.8 | 0.0168 | 1.77 | 0.28 | 100 | 4 | 96 | 1.6128 |
| 0.5 | 2.5 | 7.8 | 0.0078 | 1.93 | 0.15 | 100 | 6 | 94 | 0.7332 |
| 0.5 | 2.5 | 3.8 | 0.0038 | 2.45 | 0.03 | 100 | 6 | 94 | 0.3572 |
| 0.5 | 1 | 5.7 | 0.0057 | 2.54 | 0.03 | 100 | 6 | 94 | 0.5358 |
| 0.5 | 1 | 9.9 | 0.0099 | 2 | 0.13 | 100 | 6 | 94 | 0.9306 |
| 0.5 | 1 | 3.7 | 0.0037 | 3.35 | 0.05 | 100 | 6 | 94 | 0.3478 |
| 0.5 | 1 | 13.4 | 0.0134 | 2.02 | 0.04 | 100 | 6 | 94 | 1.2596 |
| 0.5 | 0.75 | 6.8 | 0.0068 | 2.09 | 0.04 | 100 | 6 | 94 | 0.6392 |
| 0.5 | 0.75 | 4 | 0.004 | 2.88 | 0.06 | 100 | 6 | 94 | 0.376 |
| 0.5 | 0.75 | 4.1 | 0.0041 | 2.52 | 0.05 | 100 | 6 | 94 | 0.3854 |
| 0.5 | 0.75 | 9.7 | 0.0097 | 1.81 | 0.1 | 100 | 6 | 94 | 0.9118 |
| 0.5 | 0.75 | 8.5 | 0.0085 | 2.18 | 0.07 | 100 | 6 | 94 | 0.799 |
| 0.5 | 1.25 | 14.8 | 0.0148 | 2.09 | 0.15 | 100 | 6 | 94 | 1.3912 |
| 0.5 | 1.25 | 20.4 | 0.0204 | 2 | 0.12 | 100 | 6 | 94 | 1.9176 |
| 0.5 | 1 | 19.7 | 0.0197 | 1.83 | 0.12 | 100 | 6 | 94 | 1.8518 |
| 0.5 | 1 | 15.1 | 0.0151 | 2.03 | 0.07 | 100 | 6 | 94 | 1.4194 |
| 0.5 | 1 | 20.8 | 0.0208 | 1.85 | 0.19 | 100 | 6 | 94 | 1.9552 |
| 0.5 | 1 | 25 | 0.025 | 1.97 | 0.08 | 100 | 6 | 94 | 2.35 |
| 0.5 | 1.5 | 18.1 | 0.0181 | 1.96 | 0.08 | 100 | 6 | 94 | 1.7014 |
| 0.5 | 1.75 | 16.2 | 0.0162 | 2.06 | 0.05 | 100 | 6 | 94 | 1.5228 |
| 0.5 | 1.25 | 7.2 | 0.0072 | 2.52 | 0.05 | 100 | 6 | 94 | 0.6768 |
| 0.33 | 6.6 | 20.8 | 0.0208 | 1.91 | 0.17 | 100 | 6 | 94 | 1.9552 |
| 0.33 | 6.6 | 36.9 | 0.0369 | 1.78 | 0.96 | 100 | 4 | 96 | 3.5424 |
| 0.33 | 6.6 | 43.4 | 0.0434 | 1.75 | 1.61 | 100 | 4 | 96 | 4.1664 |
| 1 | 20 | 12.5 | 0.0125 | 1.61 | 0.32 | 100 | 4 | 96 | 1.2 |
| 1 | 20 | 94.4 | 0.0944 | 1.79 | 0.71 | 100 | 4 | 96 | 9.0624 |
| 0.33 | 6.602366 | 164.4 | 0.1644 | 1.84 | 1.37 | 100 | 6 | 94 | 15.4536 |
| 0.33 | 5.751848 | 156.1 | 0.1561 | 1.84 | 0.65 | 100 | 6 | 94 | 14.6734 |
| 0.33 | 6.645786 | 171.4 | 0.1714 | 1.84 | 0.78 | 100 | 6 | 94 | 16.1116 |
| 0.33 | 6.27 | 158.3 | 0.1583 | 1.85 | 0.4 | 100 | 4 | 96 | 15.1968 |
| 0.33 | 6.27 | 139 | 0.139 | 1.85 | 0.52 | 100 | 4 | 96 | 13.344 |
| 0.33 | 6.27 | 116.3 | 0.1163 | 1.87 | 0.29 | 100 | 4 | 96 | 11.1648 |
| 0.5 | 1.5 | 9.7 | 0.0097 | 2.03 | 0.03 | 100 | 4 | 96 | 0.9312 |
| 0.5 | 1.5 | 11.1 | 0.0111 | 2.04 | 0.04 | 100 | 4 | 96 | 1.0656 |
| 0.5 | 1.5 | 8.4 | 0.0084 | 1.9 | 0.07 | 100 | 4 | 96 | 0.8064 |
| 0.5 | 1 | 11.4 | 0.0114 | 1.97 | 0.06 | 100 | 4 | 96 | 1.0944 |
| 0.5 | 1 | 15.1 | 0.0151 | 1.83 | 0.41 | 100 | 4 | 96 | 1.4496 |
| 0.5 | 1 | 7.7 | 0.0077 | 1.94 | 0.03 | 100 | 4 | 96 | 0.7392 |
| 0.25 | 10 | 24.4 | 0.0244 | 1.84 | 0.1 | 100 | 4 | 96 | 2.3424 |
| 0.25 | 0.75 | 6.3 | 0.0063 | 1.74 | 0.02 | 100 | 4 | 96 | 0.6048 |

| | | | | | | | | | |
|---|---|---|---|---|---|---|---|---|---|
| 0.25 | 2.5 | 9.6 | 0.0096 | 1.63 | 0.03 | 100 | 4 | 96 | 0.9216 |
| 0.25 | 3.5 | 7.1 | 0.0071 | 1.86 | 0.02 | 100 | 4 | 96 | 0.6816 |
| 0.25 | 4.25 | 10.2 | 0.0102 | 1.73 | 0.13 | 100 | 4 | 96 | 0.9792 |
| 0.25 | 3 | 10.4 | 0.0104 | 1.66 | 0.05 | 100 | 4 | 96 | 0.9984 |
| 0.5 | 3 | 11.4 | 0.0114 | 1.43 | 0.18 | 100 | 4 | 96 | 1.0944 |
| 1 | 0.425 | 45.1 | 0.0451 | 1.74 | 1.55 | 100 | 4 | 96 | 4.3296 |
| 1 | 0.425 | 49.2 | 0.0492 | 1.73 | 1.51 | 100 | 4 | 96 | 4.7232 |
| 1 | 0.45 | 54.7 | 0.0547 | 1.82 | 0.46 | 100 | 4 | 96 | 5.2512 |
| 1 | 0.45 | 48.6 | 0.0486 | 1.79 | 1.21 | 100 | 4 | 96 | 4.6656 |
| 1 | 0.16 | 6.5 | 0.0065 | 1.39 | 0.17 | 100 | 4 | 96 | 0.624 |
| 1 | 7.3 | 7.1 | 0.0071 | 1.62 | 0.48 | 100 | 4 | 96 | 0.6816 |
| 0.5 | 3.65 | 12.8 | 0.0128 | 1.69 | 0.38 | 100 | 4 | 96 | 1.2288 |
| 0.5 | 2.5 | 73.2 | 0.0732 | 1.85 | 1.86 | 100 | 4 | 96 | 7.0272 |

| Sequim.Na | Sequim.Qu | Sequim.Qu | Sequim.Qu | Sequim.Qu | Date.Shipp | Filter.Used | Experimen | Subsample | Filter.Pore |
|---|---|---|---|---|---|---|---|---|---|
| 1380 | 1X dsDNA | 2.02 | 101 | 0.101 | 45444 | BF-013 | NW_FE_0( | BF-013 | 0.2Âµm |
| 775 | 1X dsDNA | 103 | 5150 | 5.15 | 45444 | BF-015 | NW_FE_0( | BF-015 | 0.2Âµm |
| NA | 1X dsDNA | 43 | 4128 | 4.128 | 45444 | BF-022 | NW_FE_0( | BF-022 | 0.2Âµm |
| NA | 1X dsDNA | 45.4 | 4358.4 | 4.3584 | 45444 | BF-026 | NW_FE_0( | BF-026 | 0.2Âµm |
| NA | 1X dsDNA | 67.3 | 6460.8 | 6.4608 | 45444 | BF-027 | NW_FE_0( | BF-027 | 0.2Âµm |
| 3139.2 | 1X dsDNA | 24.9 | 2390.4 | 2.3904 | 45534 | BF-064 | NW_FE_01 | BF-064 | 0.2Âµm |
| 1612.8 | 1X dsDNA | 11.3 | 1084.8 | 1.0848 | 45534 | BF-065 | NW_FE_01 | BF-065 | 0.2Âµm |
| 733.2 | 1X dsDNA | 5.5 | 517 | 0.517 | 45534 | BF-067 | NW_FE_01 | BF-067 | 5Âµm |
| 357.2 | 1X dsDNA | 1.48 | 139.12 | 0.13912 | 45534 | BF-069 | NW_FE_01 | BF-069 | 5Âµm |
| 535.8 | 1X dsDNA | 3.78 | 355.32 | 0.35532 | 45534 | BF-075 | NW_FE_01 | BF-075 | 0.2Âµm |
| 930.6 | 1X dsDNA | 7.65 | 719.1 | 0.7191 | 45534 | BF-076 | NW_FE_01 | BF-076 | 0.2Âµm |
| 347.8 | 1X dsDNA | 2.74 | 257.56 | 0.25756 | 45534 | BF-077 | NW_FE_01 | BF-077 | 0.2Âµm |
| 1259.6 | 1X dsDNA | 4.86 | 456.84 | 0.45684 | 45534 | BF-078 | NW_FE_01 | BF-078 | 0.2Âµm |
| 639.2 | 1X dsDNA | 3.4 | 319.6 | 0.3196 | 45534 | BF-070 | NW_FE_01 | BF-070 | 0.2Âµm |
| 376 | 1X dsDNA | 2.31 | 217.14 | 0.21714 | 45534 | BF-071 | NW_FE_01 | BF-071 | 0.2Âµm |
| 385.4 | 1X dsDNA | 2.18 | 204.92 | 0.20492 | 45534 | BF-072 | NW_FE_01 | BF-072 | 0.2Âµm |
| 911.8 | 1X dsDNA | 3.39 | 318.66 | 0.31866 | 45534 | BF-073 | NW_FE_01 | BF-073 | 0.2Âµm |
| 799 | 1X dsDNA | 4.34 | 407.96 | 0.40796 | 45534 | BF-074 | NW_FE_01 | BF-074 | 0.2Âµm |
| 1391.2 | 1X dsDNA | 12.4 | 1165.6 | 1.1656 | 45534 | BF-087 | NW_FE_01 | BF-087 | 5Âµm |
| 1917.6 | 1X dsDNA | 7.4 | 695.6 | 0.6956 | 45534 | BF-088 | NW_FE_01 | BF-088 | 5Âµm |
| 1851.8 | 1X dsDNA | 13.6 | 1278.4 | 1.2784 | 45534 | BF-094 | NW_FE_01 | BF-094 | 0.2Âµm |
| 1419.4 | 1X dsDNA | 10.2 | 958.8 | 0.9588 | 45534 | BF-095 | NW_FE_01 | BF-095 | 0.2Âµm |
| 1955.2 | 1X dsDNA | 11 | 1034 | 1.034 | 45534 | BF-096 | NW_FE_01 | BF-096 | 0.2Âµm |
| 2350 | 1X dsDNA | 17.6 | 1654.4 | 1.6544 | 45534 | BF-097 | NW_FE_01 | BF-097 | 0.2Âµm |
| 1701.4 | 1X dsDNA | 11.4 | 1071.6 | 1.0716 | 45534 | BF-099 | NW_FE_01 | BF-099 | 5Âµm |
| 1522.8 | 1X dsDNA | 6.85 | 643.9 | 0.6439 | 45534 | BF-100 | NW_FE_01 | BF-100 | 5Âµm |
| 676.8 | 1X dsDNA | 3.84 | 360.96 | 0.36096 | 45534 | BF-101 | NW_FE_01 | BF-101 | 5Âµm |
| 1955.2 | 1X dsDNA | 11.2 | 1052.8 | 1.0528 | 45615 | BF-172 | NW_FE_02 | BF-172 | 0.2Âµm |
| 3542.4 | 1X dsDNA | 17.4 | 1670.4 | 1.6704 | 45615 | BF-172 | NW_FE_02 | BF-172 | 0.2Âµm |
| 4166.4 | 1X dsDNA | 26.5 | 2544 | 2.544 | 45615 | BF-173 | NW_FE_02 | BF-173 | 0.2Âµm |
| 1200 | 1X dsDNA | 5.55 | 532.8 | 0.5328 | 45615 | BF-178 | NW_FE_02 | BF-178 | 0.2Âµm |
| 9062.4 | 1X dsDNA | NA | NA | NA | 45615 | BF-179 | NW_FE_02 | BF-179 | 0.2Âµm |
| 15453.6 | 1X dsDNA | NA | NA | NA | 45615 | BF-187b | NW_FE_02 | BF-187a-c | 0.2Âµm |
| 14673.4 | 1X dsDNA | NA | NA | NA | 45615 | BF-188b | NW_FE_02 | BF-188a-c | 0.2Âµm |
| 16111.6 | 1X dsDNA | NA | NA | NA | 45615 | BF-189b | NW_FE_02 | BF-189a-c | 0.2Âµm |
| 15196.8 | 1X dsDNA | NA | NA | NA | 45615 | BF-187c | NW_FE_02 | BF-187a-c | 0.2Âµm |
| 13344 | 1X dsDNA | NA | NA | NA | 45615 | BF-188c | NW_FE_02 | BF-188a-c | 0.2Âµm |
| 11164.8 | 1X dsDNA | NA | NA | NA | 45615 | BF-189c | NW_FE_02 | BF-189a-c | 0.2Âµm |
| 931.2 | 1X dsDNA | 0.479 | 45.984 | 0.045984 | 45621 | BF-081 | NW_FE_01 | BF-081 | 5Âµm |
| 1065.6 | 1X dsDNA | 4.53 | 434.88 | 0.43488 | 45621 | BF-082 | NW_FE_01 | BF-082 | 5Âµm |
| 806.4 | 1X dsDNA | 3.27 | 313.92 | 0.31392 | 45621 | BF-083 | NW_FE_01 | BF-083 | 5Âµm |
| 1094.4 | 1X dsDNA | 6.84 | 656.64 | 0.65664 | 45615 | BF-090 | NW_FE_01 | BF-090 | 0.2Âµm |
| 1449.6 | 1X dsDNA | 7.91 | 759.36 | 0.75936 | 45615 | BF-091 | NW_FE_01 | BF-091 | 0.2Âµm |
| 739.2 | 1X dsDNA | 1.88 | 180.48 | 0.18048 | 45621 | BF-092 | NW_FE_01 | BF-092 | 0.2Âµm |
| 2342.4 | 1X dsDNA | 15.5 | 1488 | 1.488 | 45615 | BF-201a | NW_FE_02 | BF-201a-d | 0.2Âµm |
| 604.8 | 1X dsDNA | 0.051 | 4.896 | 0.004896 | 45615 | BF-202a | NW_FE_02 | BF-202a-d | 0.2Âµm |

| | | | | | | | | | |
|---|---|---|---|---|---|---|---|---|---|
| 921.6 | 1X dsDNA | 1.04 | 99.84 | 0.09984 | 45615 | BF-203a | NW_FE_02 | BF-203a-d | 0.45Âµm |
| 681.6 | 1X dsDNA | 1.4 | 134.4 | 0.1344 | 45615 | BF-204a | NW_FE_02 | BF-204a-d | 0.2Âµm |
| 979.2 | 1X dsDNA | 5.03 | 482.88 | 0.48288 | 45615 | BF-205a | NW_FE_02 | BF-205a-d | 0.2Âµm |
| 998.4 | 1X dsDNA | 4.82 | 462.72 | 0.46272 | 45615 | BF-206a | NW_FE_02 | BF-206a-d | 0.2Âµm |
| 1094.4 | 1X dsDNA | 4.18 | 401.28 | 0.40128 | 45615 | BF-124 | NW_FE_01 | BF-124 | 5Âµm |
| 4329.6 | 1X dsDNA | 42.7 | 4099.2 | 4.0992 | 45615 | BF-125 | NW_FE_01 | BF-125 | 0.2Âµm |
| 4723.2 | 1X dsDNA | 39.6 | 3801.6 | 3.8016 | 45615 | BF-126 | NW_FE_01 | BF-126 | 0.2Âµm |
| 5251.2 | 1X dsDNA | 46.7 | 4483.2 | 4.4832 | 45615 | BF-130 | NW_FE_01 | BF-130 | 0.2Âµm |
| 4665.6 | 1X dsDNA | 40.7 | 3907.2 | 3.9072 | 45615 | BF-131 | NW_FE_01 | BF-131 | 0.2Âµm |
| 624 | 1X dsDNA | 0.413 | 39.648 | 0.039648 | 45615 | BF-132 | NW_FE_01 | BF-132 | 0.05Âµm |
| 681.6 | 1X dsDNA | 4.35 | 417.6 | 0.4176 | 45615 | BF-137 | NW_FE_01 | BF-137 | 20Âµm |
| 1228.8 | 1X dsDNA | 8.45 | 811.2 | 0.8112 | 45615 | BF-138 | NW_FE_01 | BF-138 | 5Âµm |
| 7027.2 | 1X dsDNA | NA | NA | NA | 45615 | BF-144 | NW_FE_01 | BF-144 | 0.2Âµm |

| Filter.mate | Filtration.S | Filter.Fraction2 | Filtration.E | Time.of.Fil | Date.filtered | Date.of.Sample. | Time.betw |
|---|---|---|---|---|---|---|---|
| MCE | 20 -> 0.2Â | Pico (> 0.22Âµm ) | Single flasl | 15H 60M ( | 4/15/2024 | 4/15/2024 | 0 |
| MCE | 20 -> 0.2Â | Pico (> 0.22Âµm ) | Manifold | 15H 60M ( | 4/15/2024 | 4/15/2024 | 0 |
| MCE | 20 -> 5 -> ( | Pico (> 0.22Âµm ) | Single flasl | 15H 60M ( | 5/8/2024 | 5/8/2024 | 0 |
| MCE | 20 -> 5 -> ( | Pico (> 0.22Âµm ) | Single flasl | 16H 56M ( | 5/8/2024 | 5/8/2024 | 0 |
| MCE | 20 -> 5 -> ( | Pico (> 0.22Âµm ) | Single flasl | 12H 1M 0S | 5/9/2024 | 5/8/2024 | 1 |
| MCE | 20 -> 5 -> ( | Pico (> 0.22Âµm ) | Single flasl | 10H 35M ( | 7/17/2024 | 7/17/2024 | 0 |
| MCE | 20 -> 5 -> ( | Pico (> 0.22Âµm ) | Single flasl | 10H 35M ( | 7/17/2024 | 7/17/2024 | 0 |
| MCE | 20 -> 5 -> ( | Micro (> 5Âµm) | In-line pun | 10H 0M 0S | 7/23/2024 | 7/23/2024 | 0 |
| MCE | 20 -> 5 -> ( | Micro (> 5Âµm) | In-line pun | 10H 35M ( | 7/23/2024 | 7/23/2024 | 0 |
| MCE | 20 -> 5 -> ( | Pico (> 0.22Âµm ) | Single flasl | 8H 42M 0S | 7/31/2024 | 7/31/2024 | 0 |
| MCE | 20 -> 5 -> ( | Pico (> 0.22Âµm ) | Single flasl | 9H 35M 0S | 7/31/2024 | 7/31/2024 | 0 |
| MCE | 20 -> 5 -> ( | Pico (> 0.22Âµm ) | Single flasl | 10H 18M ( | 7/31/2024 | 7/31/2024 | 0 |
| MCE | 20 -> 5 -> ( | Pico (> 0.22Âµm ) | Single flasl | 11H 8M 0S | 7/31/2024 | 7/31/2024 | 0 |
| MCE | 20 -> 5 -> ( | Pico (> 0.22Âµm ) | Single flasl | 14H 53M ( | 7/23/2024 | 7/23/2024 | 0 |
| MCE | 20 -> 5 -> ( | Pico (> 0.22Âµm ) | Single flasl | 15H 15M ( | 7/23/2024 | 7/23/2024 | 0 |
| MCE | 20 -> 5 -> ( | Pico (> 0.22Âµm ) | Single flasl | 15H 44M ( | 7/23/2024 | 7/23/2024 | 0 |
| MCE | 20 -> 5 -> ( | Pico (> 0.22Âµm ) | Single flasl | 16H 12M ( | 7/23/2024 | 7/23/2024 | 0 |
| MCE | 20 -> 5 -> ( | Pico (> 0.22Âµm ) | Single flasl | 4H 35M 0S | 7/23/2024 | 7/23/2024 | 0 |
| MCE | 20 -> 5 -> ( | Micro (> 5Âµm) | In-line pun | 14H 5M 0S | 7/31/2024 | 7/31/2024 | 0 |
| MCE | 20 -> 5 -> ( | Micro (> 5Âµm) | In-line pun | 14H 15M ( | 7/31/2024 | 7/31/2024 | 0 |
| MCE | 20 -> 5 -> ( | Pico (> 0.22Âµm ) | Single flasl | 16H 23M ( | 7/31/2024 | 7/31/2024 | 0 |
| MCE | 20 -> 5 -> ( | Pico (> 0.22Âµm ) | Single flasl | 17H 56M ( | 7/31/2024 | 7/31/2024 | 0 |
| MCE | 20 -> 5 -> ( | Pico (> 0.22Âµm ) | Single flasl | 18H 10M ( | 7/31/2024 | 7/31/2024 | 0 |
| MCE | 20 -> 5 -> ( | Pico (> 0.22Âµm ) | Single flasl | 17H 49M ( | 7/31/2024 | 7/31/2024 | 0 |
| MCE | 20 -> 5 -> ( | Micro (> 5Âµm) | In-line pun | 6H 37M 0S | 7/31/2024 | 7/31/2024 | 0 |
| MCE | 20 -> 5 -> ( | Micro (> 5Âµm) | In-line pun | 6H 47M 0S | 7/31/2024 | 7/31/2024 | 0 |
| MCE | 20 -> 5 -> ( | Micro (> 5Âµm) | In-line pun | 6H 59M 0S | 7/31/2024 | 7/31/2024 | 0 |
| aPES | 20 -> 5 -> ( | Pico (> 0.22Âµm ) | Filter cup | 12H 0M 0S | 8/27/2024 | 8/26/2024 | 1 |
| aPES | 20 -> 5 -> ( | Pico (> 0.22Âµm ) | Filter cup | 12H 0M 0S | 8/27/2024 | 8/26/2024 | 1 |
| aPES | 20 -> 5 -> ( | Pico (> 0.22Âµm ) | Filter cup | 12H 35M ( | 8/27/2024 | 8/26/2024 | 1 |
| aPES | 20 -> 5 -> ( | Pico (> 0.22Âµm ) | Filter cup | NA | 8/28/2024 | 8/27/2024 | 1 |
| aPES | 20 -> 5 -> ( | Pico (> 0.22Âµm ) | Filter cup | NA | 8/28/2024 | 8/27/2024 | 1 |
| aPES | 20 -> 5 -> ( | Pico (> 0.22Âµm ) | Filter cup | 15H 10M ( | 9/9/2024 | 9/9/2024 | 0 |
| aPES | 20 -> 5 -> ( | Pico (> 0.22Âµm ) | Filter cup | 15H 6M 0S | 9/9/2024 | 9/9/2024 | 0 |
| aPES | 20 -> 5 -> ( | Pico (> 0.22Âµm ) | Filter cup | 16H 30M ( | 9/9/2024 | 9/9/2024 | 0 |
| aPES | 20 -> 5 -> ( | Pico (> 0.22Âµm ) | Filter cup | 15H 10M ( | 9/9/2024 | 9/9/2024 | 0 |
| aPES | 20 -> 5 -> ( | Pico (> 0.22Âµm ) | Filter cup | 15H 6M 0S | 9/9/2024 | 9/9/2024 | 0 |
| aPES | 20 -> 5 -> ( | Pico (> 0.22Âµm ) | Filter cup | 16H 30M ( | 9/9/2024 | 9/9/2024 | 0 |
| MCE | 20 -> 5 -> ( | Micro (> 5Âµm) | In-line pun | 12H 50M ( | 7/31/2024 | 7/31/2024 | 0 |
| MCE | 20 -> 5 -> ( | Micro (> 5Âµm) | In-line pun | 13H 15M ( | 7/31/2024 | 7/31/2024 | 0 |
| MCE | 20 -> 5 -> ( | Micro (> 5Âµm) | In-line pun | 13H 46M ( | 7/31/2024 | 7/31/2024 | 0 |
| MCE | 20 -> 5 -> ( | Pico (> 0.22Âµm ) | Single flasl | 14H 53M ( | 7/31/2024 | 7/31/2024 | 0 |
| MCE | 20 -> 5 -> ( | Pico (> 0.22Âµm ) | Single flasl | 15H 0M 0S | 7/31/2024 | 7/31/2024 | 0 |
| MCE | 20 -> 5 -> ( | Pico (> 0.22Âµm ) | Single flasl | 15H 50M ( | 7/31/2024 | 7/31/2024 | 0 |
| aPES | 20 -> 5 -> ( | Pico (> 0.22Âµm ) | Filter cup | 15H 60M ( | 10/29/2024 | 10/29/2024 | 0 |
| aPES | 20 -> 5 -> ( | Viral (< 0.22Âµm) | Filter cup | 15H 10M ( | 10/31/2024 | 10/29/2024 | 2 |

| | | | | | | | |
|---|---|---|---|---|---|---|---|
| aPES | 20 -> 5 -> ( | Viral (< 0.22Âµm) | Filter cup | 15H 30M ( | 10/31/2024 | 10/29/2024 | 2 |
| aPES | 20 -> 5 -> ( | Viral (< 0.22Âµm) | Filter cup | 16H 55M ( | 10/31/2024 | 10/29/2024 | 2 |
| aPES | 20 -> 5 -> ( | Viral (< 0.22Âµm) | Filter cup | 16H 55M ( | 10/31/2024 | 10/29/2024 | 2 |
| aPES | 20 -> 5 -> ( | Pico (> 0.22Âµm ) | Filter cup | 16H 40M ( | 10/29/2024 | 10/29/2024 | 0 |
| Nylon | 20 -> 5 -> ( | Micro (> 5Âµm) | Single flasl | 16H 30M ( | 8/8/2024 | 8/7/2024 | 1 |
| MCE | 20 -> 5 -> ( | Pico (> 0.22Âµm ) | Single flasl | 11H 57M ( | 8/9/2024 | 8/7/2024 | 2 |
| MCE | 20 -> 5 -> ( | Pico (> 0.22Âµm ) | Single flasl | 12H 36M ( | 8/9/2024 | 8/7/2024 | 2 |
| MCE | 20 -> 5 -> ( | Pico (> 0.22Âµm ) | Single flasl | 13H 36M ( | 8/9/2024 | 8/7/2024 | 2 |
| MCE | 20 -> 5 -> ( | Pico (> 0.22Âµm ) | Single flasl | 14H 40M ( | 8/9/2024 | 8/7/2024 | 2 |
| NA | 20 -> 5 -> ( | Viral (> 0.05Âµm) | Single flasl | 14H 36M ( | 8/9/2024 | 8/7/2024 | 2 |
| Nylon | 20 -> 5 -> ( | Macro ( > 20Âµm) | Single flasl | 15H 40M ( | 8/15/2024 | 8/14/2024 | 1 |
| Nylon | 20 -> 5 -> ( | Micro (> 5Âµm) | Single flasl | 14H 45M ( | 8/15/2024 | 8/14/2024 | 1 |
| aPES | 20 -> 5 -> ( | Pico (> 0.22Âµm ) | Filter cup | 15H 28M ( | 8/16/2024 | 8/14/2024 | 2 |

| Sample.Location | Time.of.pa | Sampling.M | Total.Samp | Volume.Fil | DNA.Samp | Sampling.D | Sampling.T |
|---|---|---|---|---|---|---|---|
| PNNL-Sequim Floating Dock | 14H 30M ( | Pump & in | 12.5 | 2 | NA | 45397 | 14H 30M ( |
| PNNL-Sequim Floating Dock | 14H 30M ( | Pump & in | 12.5 | 1 | NA | 45397 | 14H 30M ( |
| PNNL-Sequim Floating Dock | 12H 30M ( | Pump & in | 12.5 | 2.1 | 20240513- | 45420 | 12H 30M ( |
| PNNL-Sequim Floating Dock | 12H 30M ( | Pump & in | 12.5 | 2 | 20240513- | 45420 | 12H 30M ( |
| PNNL-Sequim Floating Dock | 12H 30M ( | Pump & in | 12.5 | 1.3 | 20240513- | 45420 | 12H 30M ( |
| PNNL-Sequim Floating Dock | 10H 35M ( | Surface gra | 15 | 1.8 | 20240718- | 45490 | 10H 35M ( |
| PNNL-Sequim Floating Dock | 10H 35M ( | Surface gra | 15 | 1 | 20240718- | 45490 | 10H 35M ( |
| PNNL-Sequim Floating Dock | 10H 30M ( | Surface gra | 22 | 5 | 20240805- | 45496 | 10H 30M ( |
| PNNL-Sequim Floating Dock | 10H 30M ( | Surface gra | 22 | 5 | 20240805- | 45496 | 10H 30M ( |
| PNNL-Sequim Floating Dock | 5H 35M 0S | Surface gra | 13 | 2 | 20240805- | 45504 | 5H 35M 0S |
| PNNL-Sequim Floating Dock | 5H 35M 0S | Surface gra | 13 | 2 | 20240805- | 45504 | 5H 35M 0S |
| PNNL-Sequim Floating Dock | 5H 35M 0S | Surface gra | 13 | 2 | 20240805- | 45504 | 5H 35M 0S |
| PNNL-Sequim Floating Dock | 5H 35M 0S | Surface gra | 13 | 2 | 20240805- | 45504 | 5H 35M 0S |
| PNNL-Sequim Floating Dock | 10H 30M ( | Surface gra | 22 | 1.5 | 20240805- | 45496 | 10H 30M ( |
| PNNL-Sequim Floating Dock | 10H 30M ( | Surface gra | 22 | 1.5 | 20240805- | 45496 | 10H 30M ( |
| PNNL-Sequim Floating Dock | 10H 30M ( | Surface gra | 22 | 1.5 | 20240805- | 45496 | 10H 30M ( |
| PNNL-Sequim Floating Dock | 10H 30M ( | Surface gra | 22 | 1.5 | 20240805- | 45496 | 10H 30M ( |
| PNNL-Sequim Floating Dock | 10H 30M ( | Surface gra | 22 | 1.5 | 20240805- | 45496 | 10H 30M ( |
| John Wayne Marina | 11H 52M ( | Surface gra | 18 | 2.5 | 20240806- | 45504 | 11H 52M ( |
| John Wayne Marina | 11H 52M ( | Surface gra | 18 | 2.5 | 20240806- | 45504 | 11H 52M ( |
| John Wayne Marina | 11H 52M ( | Surface gra | 18 | 2 | 20240807- | 45504 | 11H 52M ( |
| John Wayne Marina | 11H 52M ( | Surface gra | 18 | 2 | 20240807- | 45504 | 11H 52M ( |
| John Wayne Marina | 11H 52M ( | Surface gra | 18 | 2 | 20240807- | 45504 | 11H 52M ( |
| John Wayne Marina | 11H 52M ( | Surface gra | 18 | 2 | 20240807- | 45504 | 11H 52M ( |
| PNNL-Sequim Floating Dock | 5H 35M 0S | Surface gra | 13 | 3 | 20240807- | 45504 | 5H 35M 0S |
| PNNL-Sequim Floating Dock | 5H 35M 0S | Surface gra | 13 | 3.5 | 20240807- | 45504 | 5H 35M 0S |
| PNNL-Sequim Floating Dock | 5H 35M 0S | Surface gra | 13 | 2.5 | 20240807- | 45504 | 5H 35M 0S |
| PNNL-Sequim Floating Dock | 12H 10M ( | Surface gra | 47.5 | 20 | 20240828- | 45530 | 12H 10M ( |
| PNNL-Sequim Floating Dock | 12H 10M ( | Surface gra | 47.5 | 20 | 20240828- | 45530 | 12H 10M ( |
| PNNL-Sequim Floating Dock | 12H 10M ( | Surface gra | 47.5 | 20 | 20240828- | 45530 | 12H 10M ( |
| PNNL-Sequim Floating Dock | 13H 15M ( | Surface gra | 45 | 20 | 20240829- | 45531 | 13H 15M ( |
| PNNL-Sequim Floating Dock | 13H 15M ( | Surface gra | 45 | 20 | 20240829- | 45531 | 13H 15M ( |
| PNNL-Sequim Floating Dock | 9H 50M 0S | Surface gra | 70 | 19 | 20240911- | 45544 | 9H 50M 0S |
| PNNL-Sequim Floating Dock | 9H 50M 0S | Surface gra | 70 | 19 | 20240911- | 45544 | 9H 50M 0S |
| PNNL-Sequim Floating Dock | 9H 50M 0S | Surface gra | 70 | 17.5 | 20240911- | 45544 | 9H 50M 0S |
| PNNL-Sequim Floating Dock | 9H 50M 0S | Surface gra | 70 | 19 | 20240911- | 45544 | 9H 50M 0S |
| PNNL-Sequim Floating Dock | 9H 50M 0S | Surface gra | 70 | 19 | 20240911- | 45544 | 9H 50M 0S |
| PNNL-Sequim Floating Dock | 9H 50M 0S | Surface gra | 70 | 17.5 | 20240911- | 45544 | 9H 50M 0S |
| PNNL-Sequim Floating Dock | 12H 0M 0S | Surface gra | 15 | 3 | 20240806- | 45504 | 12H 0M 0S |
| PNNL-Sequim Floating Dock | 12H 0M 0S | Surface gra | 15 | 3 | 20240806- | 45504 | 12H 0M 0S |
| PNNL-Sequim Floating Dock | 12H 0M 0S | Surface gra | 15 | 3 | 20240806- | 45504 | 12H 0M 0S |
| PNNL-Sequim Floating Dock | 12H 0M 0S | Surface gra | 15 | 2 | 20240806- | 45504 | 12H 0M 0S |
| PNNL-Sequim Floating Dock | 12H 0M 0S | Surface gra | 15 | 2 | 20240806- | 45504 | 12H 0M 0S |
| PNNL-Sequim Floating Dock | 12H 0M 0S | Surface gra | 15 | 2 | 20240806- | 45504 | 12H 0M 0S |
| PNNL-Sequim Floating Dock | 9H 18M 0S | Surface gra | 63 | 40 | NA | 45594 | 9H 18M 0S |
| PNNL-Sequim Floating Dock | 9H 18M 0S | Surface gra | 63 | 3 | NA | 45594 | 9H 18M 0S |

| | | | | | | | |
|---|---|---|---|---|---|---|---|
| PNNL-Sequim Floating Dock | 9H 18M 0S | Surface gra | 63 | 10 | NA | 45594 | 9H 18M 0S |
| PNNL-Sequim Floating Dock | 9H 18M 0S | Surface gra | 63 | 14 | NA | 45594 | 9H 18M 0S |
| PNNL-Sequim Floating Dock | 9H 18M 0S | Surface gra | 63 | 17 | NA | 45594 | 9H 18M 0S |
| PNNL-Sequim Floating Dock | 9H 18M 0S | Surface gra | 63 | 12 | NA | 45594 | 9H 18M 0S |
| Cline Spit | 14H 45M ( | Surface gra | 15 | 6 | NA | 45511 | 14H 45M ( |
| Cline Spit | 14H 45M ( | Surface gra | 15 | 0.425 | NA | 45511 | 14H 45M ( |
| Cline Spit | 14H 45M ( | Surface gra | 15 | 0.425 | NA | 45511 | 14H 45M ( |
| Cline Spit | 14H 45M ( | Surface gra | 15 | 0.45 | NA | 45511 | 14H 45M ( |
| Cline Spit | 14H 45M ( | Surface gra | 15 | 0.45 | NA | 45511 | 14H 45M ( |
| Cline Spit | 14H 45M ( | Surface gra | 15 | 0.16 | NA | 45511 | 14H 45M ( |
| Discovery Bay | 11H 50M ( | Surface gra | 8 | 7.3 | 20241118- | 45518 | 11H 50M ( |
| Discovery Bay | 11H 50M ( | Surface gra | 8 | 7.3 | NA | 45518 | 11H 50M ( |
| Discovery Bay | 11H 50M ( | Surface gra | 8 | 5 | NA | 45518 | 11H 50M ( |

| Year | Month | Day | Hour | Min | Tidal.Stage | Site.Coord | Latitude | Longitude | Sampling.M |
|---|---|---|---|---|---|---|---|---|---|
| 2024 | 4 | 15 | 14 | 30 | Ebb | 48Â°04'45 | 48.07934 | -123.045 | Surface gra |
| 2024 | 4 | 15 | 14 | 30 | Ebb | 48Â°04'45 | 48.07934 | -123.045 | Surface gra |
| 2024 | 5 | 8 | 12 | 30 | Flood | 48Â°04'45 | 48.07942 | -123.045 | 20-> 5 Âµr |
| 2024 | 5 | 8 | 12 | 30 | Flood | 48Â°04'45 | 48.07942 | -123.045 | 20-> 5 Âµr |
| 2024 | 5 | 8 | 12 | 30 | Flood | 48Â°04'45 | 48.07942 | -123.045 | 20-> 5 Âµr |
| 2024 | 7 | 17 | 10 | 35 | Flood | 48Â°04'45 | 48.07934 | -123.045 | Surface gra |
| 2024 | 7 | 17 | 10 | 35 | Flood | 48Â°04'45 | 48.07934 | -123.045 | Surface gra |
| 2024 | 7 | 23 | 10 | 30 | Ebb | 48Â°04'45 | 48.07934 | -123.045 | Surface gra |
| 2024 | 7 | 23 | 10 | 30 | Ebb | 48Â°04'45 | 48.07934 | -123.045 | Surface gra |
| 2024 | 7 | 31 | 5 | 35 | Ebb | 48Â°04'45 | 48.07942 | -123.045 | Surface gra |
| 2024 | 7 | 31 | 5 | 35 | Ebb | 48Â°04'45 | 48.07942 | -123.045 | Surface gra |
| 2024 | 7 | 31 | 5 | 35 | Ebb | 48Â°04'45 | 48.07942 | -123.045 | Surface gra |
| 2024 | 7 | 31 | 5 | 35 | Ebb | 48Â°04'45 | 48.07942 | -123.045 | Surface gra |
| 2024 | 7 | 23 | 10 | 30 | Ebb | 48Â°04'45 | 48.07934 | -123.045 | Surface gra |
| 2024 | 7 | 23 | 10 | 30 | Ebb | 48Â°04'45 | 48.07934 | -123.045 | Surface gra |
| 2024 | 7 | 23 | 10 | 30 | Ebb | 48Â°04'45 | 48.07934 | -123.045 | Surface gra |
| 2024 | 7 | 23 | 10 | 30 | Ebb | 48Â°04'45 | 48.07934 | -123.045 | Surface gra |
| 2024 | 7 | 23 | 10 | 30 | Ebb | 48Â°04'45 | 48.07934 | -123.045 | Surface gra |
| 2024 | 7 | 31 | 11 | 52 | Flood | 48Â°03'41 | 48.06144 | -123.039 | Surface gra |
| 2024 | 7 | 31 | 11 | 52 | Flood | 48Â°03'41 | 48.06144 | -123.039 | Surface gra |
| 2024 | 7 | 31 | 11 | 52 | Flood | 48Â°03'41 | 48.06144 | -123.039 | Surface gra |
| 2024 | 7 | 31 | 11 | 52 | Flood | 48Â°03'41 | 48.06144 | -123.039 | Surface gra |
| 2024 | 7 | 31 | 11 | 52 | Flood | 48Â°03'41 | 48.06144 | -123.039 | Surface gra |
| 2024 | 7 | 31 | 11 | 52 | Flood | 48Â°03'41 | 48.06144 | -123.039 | Surface gra |
| 2024 | 7 | 31 | 5 | 35 | Ebb | 48Â°04'45 | 48.07942 | -123.045 | Surface gra |
| 2024 | 7 | 31 | 5 | 35 | Ebb | 48Â°04'45 | 48.07942 | -123.045 | Surface gra |
| 2024 | 7 | 31 | 5 | 35 | Ebb | 48Â°04'45 | 48.07942 | -123.045 | Surface gra |
| 2024 | 8 | 26 | 12 | 10 | Slack High | 48Â°04'45 | 48.07942 | -123.045 | Surface gra |
| 2024 | 8 | 26 | 12 | 10 | Slack High | 48Â°04'45 | 48.07942 | -123.045 | Surface gra |
| 2024 | 8 | 26 | 12 | 10 | Slack High | 48Â°04'45 | 48.07942 | -123.045 | Surface gra |
| 2024 | 8 | 27 | 13 | 15 | Slack High | 48Â°04'45 | 48.07942 | -123.045 | Surface gra |
| 2024 | 8 | 27 | 13 | 15 | Slack High | 48Â°04'45 | 48.07942 | -123.045 | Surface gra |
| 2024 | 9 | 9 | 9 | 50 | Slack High | 48Â°04'45 | 48.07942 | -123.045 | Surface gra |
| 2024 | 9 | 9 | 9 | 50 | Slack High | 48Â°04'45 | 48.07942 | -123.045 | Surface gra |
| 2024 | 9 | 9 | 9 | 50 | Slack High | 48Â°04'45 | 48.07942 | -123.045 | Surface gra |
| 2024 | 9 | 9 | 9 | 50 | Slack High | 48Â°04'45 | 48.07942 | -123.045 | Surface gra |
| 2024 | 9 | 9 | 9 | 50 | Slack High | 48Â°04'45 | 48.07942 | -123.045 | Surface gra |
| 2024 | 9 | 9 | 9 | 50 | Slack High | 48Â°04'45 | 48.07942 | -123.045 | Surface gra |
| 2024 | 7 | 31 | 12 | 0 | Flood | 48Â°04'45 | 48.07942 | -123.045 | Surface gra |
| 2024 | 7 | 31 | 12 | 0 | Flood | 48Â°04'45 | 48.07942 | -123.045 | Surface gra |
| 2024 | 7 | 31 | 12 | 0 | Flood | 48Â°04'45 | 48.07942 | -123.045 | Surface gra |
| 2024 | 7 | 31 | 12 | 0 | Flood | 48Â°04'45 | 48.07942 | -123.045 | Surface gra |
| 2024 | 7 | 31 | 12 | 0 | Flood | 48Â°04'45 | 48.07942 | -123.045 | Surface gra |
| 2024 | 7 | 31 | 12 | 0 | Flood | 48Â°04'45 | 48.07942 | -123.045 | Surface gra |
| 2024 | 10 | 29 | 9 | 18 | Flood | 48Â°04'45 | 48.07942 | -123.045 | Surface gra |
| 2024 | 10 | 29 | 9 | 18 | Flood | 48Â°04'45 | 48.07942 | -123.045 | Surface gra |

| | | | | | | | | | |
|---|---|---|---|---|---|---|---|---|---|
| 2024 | 10 | 29 | 9 | 18 | Flood | 48Â°04'45 | 48.07942 | -123.045 | Surface gra |
| 2024 | 10 | 29 | 9 | 18 | Flood | 48Â°04'45 | 48.07942 | -123.045 | Surface gra |
| 2024 | 10 | 29 | 9 | 18 | Flood | 48Â°04'45 | 48.07942 | -123.045 | Surface gra |
| 2024 | 10 | 29 | 9 | 18 | Flood | 48Â°04'45 | 48.07942 | -123.045 | Surface gra |
| 2024 | NA | NA | NA | NA | Flood | 48Â°09'06 | 48.15181 | -123.153 | Surface Gr |
| 2024 | NA | NA | NA | NA | Flood | 48Â°09'06 | 48.15181 | -123.153 | Surface Gr |
| 2024 | NA | NA | NA | NA | Flood | 48Â°09'06 | 48.15181 | -123.153 | Surface Gr |
| 2024 | NA | NA | NA | NA | Flood | 48Â°09'06 | 48.15181 | -123.153 | Surface Gr |
| 2024 | NA | NA | NA | NA | Flood | 48Â°09'06 | 48.15181 | -123.153 | Surface Gr |
| 2024 | NA | NA | NA | NA | Flood | 48Â°09'06 | 48.15181 | -123.153 | Surface Gr |
| 2024 | NA | NA | NA | NA | Flood | 47Â°59'48 | 47.99678 | -122.876 | Surface gra |
| 2024 | NA | NA | NA | NA | Flood | 47Â°59'48 | 47.99678 | -122.876 | Surface gra |
| 2024 | NA | NA | NA | NA | Flood | 47Â°59'48 | 47.99678 | -122.876 | Surface gra |

| Volume.Co | Secchi.Dep | Time.of.Se | Cloud.Cov | Sea.State.. | Wind.spee | Air.Temp.. | Water.tem | Dissolved.( | Salinity..pp |
|---|---|---|---|---|---|---|---|---|---|
| 12.5 | 5 | 14H 35M ( | mostly clo | cats paws, | NA | NA | NA | NA | NA |
| 12.5 | 5 | 14H 35M ( | mostly clo | cats paws, | NA | NA | NA | NA | NA |
| 12.5 | 4 | 12H 40M ( | Clear | moslty cal | 12.2 | NA | NA | NA | NA |
| 12.5 | 4 | 12H 40M ( | Clear | moslty cal | 12.2 | NA | NA | NA | NA |
| 12.5 | 4 | 12H 40M ( | Clear | moslty cal | 12.2 | NA | NA | NA | NA |
| 15 | 5 | 10H 40M ( | partially cl | calm | NA | NA | NA | NA | NA |
| 15 | 5 | 10H 40M ( | partially cl | calm | NA | NA | NA | NA | NA |
| 22 | NA | NA | partially cl | fast surfac | NA | NA | NA | NA | NA |
| 22 | NA | NA | partially cl | fast surfac | NA | NA | NA | NA | NA |
| 13 | 7.5 | NA | overcast | calm | NA | NA | NA | NA | NA |
| 13 | 7.5 | NA | overcast | calm | NA | NA | NA | NA | NA |
| 13 | 7.5 | NA | overcast | calm | NA | NA | NA | NA | NA |
| 13 | 7.5 | NA | overcast | calm | NA | NA | NA | NA | NA |
| 22 | NA | NA | partially cl | fast surfac | NA | NA | NA | NA | NA |
| 22 | NA | NA | partially cl | fast surfac | NA | NA | NA | NA | NA |
| 22 | NA | NA | partially cl | fast surfac | NA | NA | NA | NA | NA |
| 22 | NA | NA | partially cl | fast surfac | NA | NA | NA | NA | NA |
| 22 | NA | NA | partially cl | fast surfac | NA | NA | NA | NA | NA |
| 18 | NA | NA | fully overc | calm, cats | NA | NA | 16.1 | 13.42 | 31.61 |
| 18 | NA | NA | fully overc | calm, cats | NA | NA | 16.1 | 13.42 | 31.61 |
| 18 | NA | NA | fully overc | calm, cats | NA | NA | 16.1 | 13.42 | 31.61 |
| 18 | NA | NA | fully overc | calm, cats | NA | NA | 16.1 | 13.42 | 31.61 |
| 18 | NA | NA | fully overc | calm, cats | NA | NA | 16.1 | 13.42 | 31.61 |
| 18 | NA | NA | fully overc | calm, cats | NA | NA | 16.1 | 13.42 | 31.61 |
| 13 | 7.5 | NA | overcast | calm | NA | NA | NA | NA | NA |
| 13 | 7.5 | NA | overcast | calm | NA | NA | NA | NA | NA |
| 13 | 7.5 | NA | overcast | calm | NA | NA | NA | NA | NA |
| 47.5 | 5.5 | 12H 10M ( | partially cl | very calm | NA | NA | NA | NA | NA |
| 47.5 | 5.5 | 12H 10M ( | partially cl | very calm | NA | NA | NA | NA | NA |
| 47.5 | 5.5 | 12H 10M ( | partially cl | very calm | NA | NA | NA | NA | NA |
| 45 | 6 | 13H 15M ( | Clear | fairly calm, | NA | NA | NA | NA | NA |
| 45 | 6 | 13H 15M ( | Clear | fairly calm, | NA | NA | NA | NA | NA |
| 70 | 6 | NA | overcast | calm | NA | NA | NA | NA | NA |
| 70 | 6 | NA | overcast | calm | NA | NA | NA | NA | NA |
| 70 | 6 | NA | overcast | calm | NA | NA | NA | NA | NA |
| 70 | 6 | NA | overcast | calm | NA | NA | NA | NA | NA |
| 70 | 6 | NA | overcast | calm | NA | NA | NA | NA | NA |
| 70 | 6 | NA | overcast | calm | NA | NA | NA | NA | NA |
| 15 | 5 | 12H 0M 0S | overcast | calm | NA | NA | NA | NA | NA |
| 15 | 5 | 12H 0M 0S | overcast | calm | NA | NA | NA | NA | NA |
| 15 | 5 | 12H 0M 0S | overcast | calm | NA | NA | NA | NA | NA |
| 15 | 5 | 12H 0M 0S | overcast | calm | NA | NA | NA | NA | NA |
| 15 | 5 | 12H 0M 0S | overcast | calm | NA | NA | NA | NA | NA |
| 15 | 5 | 12H 0M 0S | overcast | calm | NA | NA | NA | NA | NA |
| 60 | 7.5 | 9H 21M 0S | mostly sur | calm | NA | NA | NA | NA | NA |
| 60 | 7.5 | 9H 21M 0S | mostly sur | calm | NA | NA | NA | NA | NA |

| | | | | | | | | | |
|---|---|---|---|---|---|---|---|---|---|
| 60 | 7.5 | 9H 21M 0S | mostly sur | calm | NA | NA | NA | NA | NA |
| 60 | 7.5 | 9H 21M 0S | mostly sur | calm | NA | NA | NA | NA | NA |
| 60 | 7.5 | 9H 21M 0S | mostly sur | calm | NA | NA | NA | NA | NA |
| 60 | 7.5 | 9H 21M 0S | mostly sur | calm | NA | NA | NA | NA | NA |
| 15 | NA | NA | clear besid | minor ripp | 3.2 | 18.4 | 19.8 | 9.34 | 30.28 |
| 15 | NA | NA | clear besid | minor ripp | 3.2 | 18.4 | 19.8 | 9.34 | 30.28 |
| 15 | NA | NA | clear besid | minor ripp | 3.2 | 18.4 | 19.8 | 9.34 | 30.28 |
| 15 | NA | NA | clear besid | minor ripp | 3.2 | 18.4 | 19.8 | 9.34 | 30.28 |
| 15 | NA | NA | clear besid | minor ripp | 3.2 | 18.4 | 19.8 | 9.34 | 30.28 |
| 15 | NA | NA | clear besid | minor ripp | 3.2 | 18.4 | 19.8 | 9.34 | 30.28 |
| 8 | 2 | 11H 46M 0 | partially cl | calm | 2 | 20.1 | NA | NA | NA |
| 8 | 2 | 11H 46M 0 | partially cl | calm | 2 | 20.1 | NA | NA | NA |
| 8 | 2 | 11H 46M 0 | partially cl | calm | 2 | 20.1 | NA | NA | NA |

| Turbidity... | pH...YSI.Se | Chlorophy | P.E.........YS | water_leve | temp..deg | do..mg.L... | salinity..pp | airtemp_a | baro_press |
|---|---|---|---|---|---|---|---|---|---|
| NA | NA | NA | NA | 1.425 | NA | NA | NA | 9.45 | 1023.458 |
| NA | NA | NA | NA | 1.425 | NA | NA | NA | 9.45 | 1023.458 |
| NA | NA | NA | NA | 1.511 | NA | NA | NA | 7.896 | 1030.355 |
| NA | NA | NA | NA | 1.511 | NA | NA | NA | 7.896 | 1030.355 |
| NA | NA | NA | NA | 1.511 | NA | NA | NA | 7.896 | 1030.355 |
| NA | NA | NA | NA | 0.919074 | 14.7057 | 11.945 | 31.0226 | 14.68 | 1013.207 |
| NA | NA | NA | NA | 0.919074 | 14.7057 | 11.945 | 31.0226 | 14.68 | 1013.207 |
| NA | NA | NA | NA | 1.910263 | 11.4591 | 7.265 | 31.3504 | 13.29 | 1023.117 |
| NA | NA | NA | NA | 1.910263 | 11.4591 | 7.265 | 31.3504 | 13.29 | 1023.117 |
| NA | NA | NA | NA | 2.104993 | 12.5199 | 8.178 | 31.5527 | 16.72 | 1017.2 |
| NA | NA | NA | NA | 2.104993 | 12.5199 | 8.178 | 31.5527 | 16.72 | 1017.2 |
| NA | NA | NA | NA | 2.104993 | 12.5199 | 8.178 | 31.5527 | 16.72 | 1017.2 |
| NA | NA | NA | NA | 2.104993 | 12.5199 | 8.178 | 31.5527 | 16.72 | 1017.2 |
| NA | NA | NA | NA | 1.910263 | 11.4591 | 7.265 | 31.3504 | 13.29 | 1023.117 |
| NA | NA | NA | NA | 1.910263 | 11.4591 | 7.265 | 31.3504 | 13.29 | 1023.117 |
| NA | NA | NA | NA | 1.910263 | 11.4591 | 7.265 | 31.3504 | 13.29 | 1023.117 |
| NA | NA | NA | NA | 1.910263 | 11.4591 | 7.265 | 31.3504 | 13.29 | 1023.117 |
| NA | NA | NA | NA | 1.910263 | 11.4591 | 7.265 | 31.3504 | 13.29 | 1023.117 |
| 1.1 | 8.14 | NA | NA | NA | NA | NA | NA | NA | NA |
| 1.1 | 8.14 | NA | NA | NA | NA | NA | NA | NA | NA |
| 1.1 | 8.14 | NA | NA | NA | NA | NA | NA | NA | NA |
| 1.1 | 8.14 | NA | NA | NA | NA | NA | NA | NA | NA |
| 1.1 | 8.14 | NA | NA | NA | NA | NA | NA | NA | NA |
| 1.1 | 8.14 | NA | NA | NA | NA | NA | NA | NA | NA |
| NA | NA | NA | NA | 2.104993 | 12.5199 | 8.178 | 31.5527 | 16.72 | 1017.2 |
| NA | NA | NA | NA | 2.104993 | 12.5199 | 8.178 | 31.5527 | 16.72 | 1017.2 |
| NA | NA | NA | NA | 2.104993 | 12.5199 | 8.178 | 31.5527 | 16.72 | 1017.2 |
| NA | NA | NA | NA | -0.23541 | 11.6738 | 6.489 | 31.808 | 11.84 | 1022.228 |
| NA | NA | NA | NA | -0.23541 | 11.6738 | 6.489 | 31.808 | 11.84 | 1022.228 |
| NA | NA | NA | NA | -0.23541 | 11.6738 | 6.489 | 31.808 | 11.84 | 1022.228 |
| NA | NA | NA | NA | -0.27953 | 11.182 | 5.823 | 31.8526 | 10.27 | 1020.315 |
| NA | NA | NA | NA | -0.27953 | 11.182 | 5.823 | 31.8526 | 10.27 | 1020.315 |
| NA | NA | NA | NA | 0.066413 | -9999 | -9999 | -9999 | 13.84 | 1014.2 |
| NA | NA | NA | NA | 0.066413 | -9999 | -9999 | -9999 | 13.84 | 1014.2 |
| NA | NA | NA | NA | 0.066413 | -9999 | -9999 | -9999 | 13.84 | 1014.2 |
| NA | NA | NA | NA | 0.066413 | -9999 | -9999 | -9999 | 13.84 | 1014.2 |
| NA | NA | NA | NA | 0.066413 | -9999 | -9999 | -9999 | 13.84 | 1014.2 |
| NA | NA | NA | NA | 0.066413 | -9999 | -9999 | -9999 | 13.84 | 1014.2 |
| NA | NA | NA | NA | 0.083453 | 12.7594 | 7.342 | 31.4584 | 14.23 | 1017 |
| NA | NA | NA | NA | 0.083453 | 12.7594 | 7.342 | 31.4584 | 14.23 | 1017 |
| NA | NA | NA | NA | 0.083453 | 12.7594 | 7.342 | 31.4584 | 14.23 | 1017 |
| NA | NA | NA | NA | 0.083453 | 12.7594 | 7.342 | 31.4584 | 14.23 | 1017 |
| NA | NA | NA | NA | 0.083453 | 12.7594 | 7.342 | 31.4584 | 14.23 | 1017 |
| NA | NA | NA | NA | 0.083453 | 12.7594 | 7.342 | 31.4584 | 14.23 | 1017 |
| NA | NA | NA | NA | 1.408433 | 9.6607 | 5.47 | 31.9367 | 8.05 | 1017.667 |
| NA | NA | NA | NA | 1.408433 | 9.6607 | 5.47 | 31.9367 | 8.05 | 1017.667 |

| | | | | | | | | | |
|---|---|---|---|---|---|---|---|---|---|
| NA | NA | NA | NA | 1.408433 | 9.6607 | 5.47 | 31.9367 | 8.05 | 1017.667 |
| NA | NA | NA | NA | 1.408433 | 9.6607 | 5.47 | 31.9367 | 8.05 | 1017.667 |
| NA | NA | NA | NA | 1.408433 | 9.6607 | 5.47 | 31.9367 | 8.05 | 1017.667 |
| NA | NA | NA | NA | 1.408433 | 9.6607 | 5.47 | 31.9367 | 8.05 | 1017.667 |
| 55.22 | 8.2 | NA | NA | NA | NA | NA | NA | NA | NA |
| 55.22 | 8.2 | NA | NA | NA | NA | NA | NA | NA | NA |
| 55.22 | 8.2 | NA | NA | NA | NA | NA | NA | NA | NA |
| 55.22 | 8.2 | NA | NA | NA | NA | NA | NA | NA | NA |
| 55.22 | 8.2 | NA | NA | NA | NA | NA | NA | NA | NA |
| 55.22 | 8.2 | NA | NA | NA | NA | NA | NA | NA | NA |
| NA | NA | NA | NA | NA | NA | NA | NA | NA | NA |
| NA | NA | NA | NA | NA | NA | NA | NA | NA | NA |
| NA | NA | NA | NA | NA | NA | NA | NA | NA | NA |

| rh......Sequ | rain..mm.. | winddir..de | windspeec | windspeec | land.par_a | underwate | underwate | chlorophyl | phycoerytl |
|---|---|---|---|---|---|---|---|---|---|
| 62.42 | 0 | 353.2 | 1.213 | 3.199 | 505.7 | 3558 | 7999 | 0.52 | 7.924 |
| 62.42 | 0 | 353.2 | 1.213 | 3.199 | 505.7 | 3558 | 7999 | 0.52 | 7.924 |
| 71.3 | 0 | 171.4 | 1.537 | 2.566 | 1.579 | 3313 | 7999 | 0.8954 | -0.1278 |
| 71.3 | 0 | 171.4 | 1.537 | 2.566 | 1.579 | 3313 | 7999 | 0.8954 | -0.1278 |
| 71.3 | 0 | 171.4 | 1.537 | 2.566 | 1.579 | 3313 | 7999 | 0.8954 | -0.1278 |
| 88.8 | 0 | 317.9 | 0.445 | 1.1 | 0 | 3137 | 7999 | 1.4036 | -0.0426 |
| 88.8 | 0 | 317.9 | 0.445 | 1.1 | 0 | 3137 | 7999 | 1.4036 | -0.0426 |
| 74.94 | 0 | 347.8 | 1.654 | 3.132 | 0 | 3204 | 7999 | 2.057 | 0.0426 |
| 74.94 | 0 | 347.8 | 1.654 | 3.132 | 0 | 3204 | 7999 | 2.057 | 0.0426 |
| 89.6 | 0 | 25.24 | 0.633 | 2.299 | 0 | 3030 | 7999 | 22.385 | 0.426 |
| 89.6 | 0 | 25.24 | 0.633 | 2.299 | 0 | 3030 | 7999 | 22.385 | 0.426 |
| 89.6 | 0 | 25.24 | 0.633 | 2.299 | 0 | 3030 | 7999 | 22.385 | 0.426 |
| 89.6 | 0 | 25.24 | 0.633 | 2.299 | 0 | 3030 | 7999 | 22.385 | 0.426 |
| 74.94 | 0 | 347.8 | 1.654 | 3.132 | 0 | 3204 | 7999 | 2.057 | 0.0426 |
| 74.94 | 0 | 347.8 | 1.654 | 3.132 | 0 | 3204 | 7999 | 2.057 | 0.0426 |
| 74.94 | 0 | 347.8 | 1.654 | 3.132 | 0 | 3204 | 7999 | 2.057 | 0.0426 |
| 74.94 | 0 | 347.8 | 1.654 | 3.132 | 0 | 3204 | 7999 | 2.057 | 0.0426 |
| 74.94 | 0 | 347.8 | 1.654 | 3.132 | 0 | 3204 | 7999 | 2.057 | 0.0426 |
| NA | NA | NA | NA | NA | NA | NA | NA | NA | NA |
| NA | NA | NA | NA | NA | NA | NA | NA | NA | NA |
| NA | NA | NA | NA | NA | NA | NA | NA | NA | NA |
| NA | NA | NA | NA | NA | NA | NA | NA | NA | NA |
| NA | NA | NA | NA | NA | NA | NA | NA | NA | NA |
| NA | NA | NA | NA | NA | NA | NA | NA | NA | NA |
| 89.6 | 0 | 25.24 | 0.633 | 2.299 | 0 | 3030 | 7999 | 22.385 | 0.426 |
| 89.6 | 0 | 25.24 | 0.633 | 2.299 | 0 | 3030 | 7999 | 22.385 | 0.426 |
| 89.6 | 0 | 25.24 | 0.633 | 2.299 | 0 | 3030 | 7999 | 22.385 | 0.426 |
| 93.1 | 0 | 344.5 | 0 | 0 | 0 | 167.5 | 0 | 0.8591 | 0 |
| 93.1 | 0 | 344.5 | 0 | 0 | 0 | 167.5 | 0 | 0.8591 | 0 |
| 93.1 | 0 | 344.5 | 0 | 0 | 0 | 167.5 | 0 | 0.8591 | 0 |
| 86.3 | 0 | 172.9 | 0.744 | 1.299 | 2.002 | 614.4 | 0 | 0.8349 | 0 |
| 86.3 | 0 | 172.9 | 0.744 | 1.299 | 2.002 | 614.4 | 0 | 0.8349 | 0 |
| 94.1 | 0 | 321.4 | 0.964 | 1.466 | 0 | 497.4 | 0 | 4.9731 | 0.1278 |
| 94.1 | 0 | 321.4 | 0.964 | 1.466 | 0 | 497.4 | 0 | 4.9731 | 0.1278 |
| 94.1 | 0 | 321.4 | 0.964 | 1.466 | 0 | 497.4 | 0 | 4.9731 | 0.1278 |
| 94.1 | 0 | 321.4 | 0.964 | 1.466 | 0 | 497.4 | 0 | 4.9731 | 0.1278 |
| 94.1 | 0 | 321.4 | 0.964 | 1.466 | 0 | 497.4 | 0 | 4.9731 | 0.1278 |
| 94.1 | 0 | 321.4 | 0.964 | 1.466 | 0 | 497.4 | 0 | 4.9731 | 0.1278 |
| 96.4 | 0 | 9.54 | 0 | 0 | 0 | 3066 | 7999 | 14.3869 | 0.9372 |
| 96.4 | 0 | 9.54 | 0 | 0 | 0 | 3066 | 7999 | 14.3869 | 0.9372 |
| 96.4 | 0 | 9.54 | 0 | 0 | 0 | 3066 | 7999 | 14.3869 | 0.9372 |
| 96.4 | 0 | 9.54 | 0 | 0 | 0 | 3066 | 7999 | 14.3869 | 0.9372 |
| 96.4 | 0 | 9.54 | 0 | 0 | 0 | 3066 | 7999 | 14.3869 | 0.9372 |
| 96.4 | 0 | 9.54 | 0 | 0 | 0 | 3066 | 7999 | 14.3869 | 0.9372 |
| 90.2 | 0 | 176.3 | 0.084 | 0.833 | 0 | 0 | 0 | 0.1815 | -0.0426 |
| 90.2 | 0 | 176.3 | 0.084 | 0.833 | 0 | 0 | 0 | 0.1815 | -0.0426 |

| | | | | | | | | | |
|---|---|---|---|---|---|---|---|---|---|
| 90.2 | 0 | 176.3 | 0.084 | 0.833 | 0 | 0 | 0 | 0.1815 | -0.0426 |
| 90.2 | 0 | 176.3 | 0.084 | 0.833 | 0 | 0 | 0 | 0.1815 | -0.0426 |
| 90.2 | 0 | 176.3 | 0.084 | 0.833 | 0 | 0 | 0 | 0.1815 | -0.0426 |
| 90.2 | 0 | 176.3 | 0.084 | 0.833 | 0 | 0 | 0 | 0.1815 | -0.0426 |
| NA | NA | NA | NA | NA | NA | NA | NA | NA | NA |
| NA | NA | NA | NA | NA | NA | NA | NA | NA | NA |
| NA | NA | NA | NA | NA | NA | NA | NA | NA | NA |
| NA | NA | NA | NA | NA | NA | NA | NA | NA | NA |
| NA | NA | NA | NA | NA | NA | NA | NA | NA | NA |
| NA | NA | NA | NA | NA | NA | NA | NA | NA | NA |
| NA | NA | NA | NA | NA | NA | NA | NA | NA | NA |
| NA | NA | NA | NA | NA | NA | NA | NA | NA | NA |
| NA | NA | NA | NA | NA | NA | NA | NA | NA | NA |

| cdom..ppb | pco2_wate | pco2_air..S | o2_water. | o2_air | pH..Sequin | Environme | Processing | X..of.Macr | X..of.Micro |
|---|---|---|---|---|---|---|---|---|---|
| 0.546 | 494.47 | 431.467 | 20.679 | 21.496 | NA | 1. Incomin | NA | NA | NA |
| 0.546 | 494.47 | 431.467 | 20.679 | 21.496 | NA | 1. Incomin | NA | NA | NA |
| 0.91 | NA | NA | NA | NA | 7.772737 | 1. Slight bl | NA | NA | NA |
| 0.91 | NA | NA | NA | NA | 7.772737 | 1. Slight bl | NA | NA | NA |
| 0.91 | NA | NA | NA | NA | 7.772737 | 1. Slight bl | NA | NA | NA |
| 1.274 | NA | NA | NA | NA | 8.250541 | NA | 1. In-line fi | NA | NA |
| 1.274 | NA | NA | NA | NA | 8.250541 | NA | 1. In-line fi | NA | NA |
| 1.092 | 461.5607 | 430.988 | 20.52696 | 21.13006 | 7.974967 | 1. Secchi h | NA | NA | NA |
| 1.092 | 461.5607 | 430.988 | 20.52696 | 21.13006 | 7.974967 | 1. Secchi h | NA | NA | NA |
| 0.728 | 491.8286 | 424.7019 | 20.19966 | 21.16328 | 7.989702 | Dawn Sam | 1.Primed a | NA | NA |
| 0.728 | 491.8286 | 424.7019 | 20.19966 | 21.16328 | 7.989702 | Dawn Sam | 1.Primed a | NA | NA |
| 0.728 | 491.8286 | 424.7019 | 20.19966 | 21.16328 | 7.989702 | Dawn Sam | 1.Primed a | NA | NA |
| 0.728 | 491.8286 | 424.7019 | 20.19966 | 21.16328 | 7.989702 | Dawn Sam | 1.Primed a | NA | NA |
| 1.092 | 461.5607 | 430.988 | 20.52696 | 21.13006 | 7.974967 | 1. Secchi h | NA | NA | NA |
| 1.092 | 461.5607 | 430.988 | 20.52696 | 21.13006 | 7.974967 | 1. Secchi h | NA | NA | NA |
| 1.092 | 461.5607 | 430.988 | 20.52696 | 21.13006 | 7.974967 | 1. Secchi h | NA | NA | NA |
| 1.092 | 461.5607 | 430.988 | 20.52696 | 21.13006 | 7.974967 | 1. Secchi h | NA | NA | NA |
| 1.092 | 461.5607 | 430.988 | 20.52696 | 21.13006 | 7.974967 | 1. Secchi h | NA | NA | NA |
| NA | NA | NA | NA | NA | NA | NA | NA | NA | NA |
| NA | NA | NA | NA | NA | NA | NA | NA | NA | NA |
| NA | NA | NA | NA | NA | NA | NA | NA | NA | NA |
| NA | NA | NA | NA | NA | NA | NA | NA | NA | NA |
| NA | NA | NA | NA | NA | NA | NA | NA | NA | NA |
| NA | NA | NA | NA | NA | NA | NA | NA | NA | NA |
| 0.728 | 491.8286 | 424.7019 | 20.19966 | 21.16328 | 7.989702 | Dawn Sam | 1.Primed a | NA | NA |
| 0.728 | 491.8286 | 424.7019 | 20.19966 | 21.16328 | 7.989702 | Dawn Sam | 1.Primed a | NA | NA |
| 0.728 | 491.8286 | 424.7019 | 20.19966 | 21.16328 | 7.989702 | Dawn Sam | 1.Primed a | NA | NA |
| 0.182 | 722.0337 | 486.2953 | 18.22477 | 21.02195 | 7.679168 | 1. Secchi h | NA | NA | NA |
| 0.182 | 722.0337 | 486.2953 | 18.22477 | 21.02195 | 7.679168 | 1. Secchi h | NA | NA | NA |
| 0.182 | 722.0337 | 486.2953 | 18.22477 | 21.02195 | 7.679168 | 1. Secchi h | NA | NA | NA |
| 0.182 | 772.2938 | 424.3737 | 18.09052 | 21.16397 | 7.828396 | 1. Secchi h | NA | NA | NA |
| 0.182 | 772.2938 | 424.3737 | 18.09052 | 21.16397 | 7.828396 | 1. Secchi h | NA | NA | NA |
| 6.552 | 673.0436 | 428.9513 | 19.84983 | 21.14954 | NA | NA | NA | NA | NA |
| 6.552 | 673.0436 | 428.9513 | 19.84983 | 21.14954 | NA | NA | NA | NA | NA |
| 6.552 | 673.0436 | 428.9513 | 19.84983 | 21.14954 | NA | NA | NA | NA | NA |
| 6.552 | 673.0436 | 428.9513 | 19.84983 | 21.14954 | NA | NA | NA | NA | NA |
| 6.552 | 673.0436 | 428.9513 | 19.84983 | 21.14954 | NA | NA | NA | NA | NA |
| 6.552 | 673.0436 | 428.9513 | 19.84983 | 21.14954 | NA | NA | NA | NA | NA |
| 0.819 | 542.5183 | 427.9448 | 20.03581 | 21.13724 | 7.893584 | NA | NA | NA | NA |
| 0.819 | 542.5183 | 427.9448 | 20.03581 | 21.13724 | 7.893584 | NA | NA | NA | NA |
| 0.819 | 542.5183 | 427.9448 | 20.03581 | 21.13724 | 7.893584 | NA | NA | NA | NA |
| 0.819 | 542.5183 | 427.9448 | 20.03581 | 21.13724 | 7.893584 | NA | NA | NA | NA |
| 0.819 | 542.5183 | 427.9448 | 20.03581 | 21.13724 | 7.893584 | NA | NA | NA | NA |
| 0.819 | 542.5183 | 427.9448 | 20.03581 | 21.13724 | 7.893584 | NA | NA | NA | NA |
| 0.091 | 949.7529 | 437.998 | 17.82908 | 21.41684 | -9999 | Calm wate | NA | NA | NA |
| 0.091 | 949.7529 | 437.998 | 17.82908 | 21.41684 | -9999 | Calm wate | NA | NA | NA |

| | | | | | | | | | |
|---|---|---|---|---|---|---|---|---|---|
| 0.091 | 949.7529 | 437.998 | 17.82908 | 21.41684 | -9999 | Calm wate | NA | NA | NA |
| 0.091 | 949.7529 | 437.998 | 17.82908 | 21.41684 | -9999 | Calm wate | NA | NA | NA |
| 0.091 | 949.7529 | 437.998 | 17.82908 | 21.41684 | -9999 | Calm wate | NA | NA | NA |
| 0.091 | 949.7529 | 437.998 | 17.82908 | 21.41684 | -9999 | Calm wate | NA | NA | NA |
| NA | NA | NA | NA | NA | NA | 1. Too sha | NA | NA | NA |
| NA | NA | NA | NA | NA | NA | 1. Too sha | NA | NA | NA |
| NA | NA | NA | NA | NA | NA | 1. Too sha | NA | NA | NA |
| NA | NA | NA | NA | NA | NA | 1. Too sha | NA | NA | NA |
| NA | NA | NA | NA | NA | NA | 1. Too sha | NA | NA | NA |
| NA | NA | NA | NA | NA | NA | 1. Too sha | NA | NA | NA |
| NA | NA | NA | NA | NA | NA | 1. Secchi h | 1. Two 4L | NA | NA |
| NA | NA | NA | NA | NA | NA | 1. Secchi h | 1. Two 4L | NA | NA |
| NA | NA | NA | NA | NA | NA | 1. Secchi h | 1. Two 4L | NA | NA |

| X..of.PIco.f | X..of.viral.f | DNA.Extrac | Sequenced | TargetSequ | Grouping | kappa | C | LR | modelR |
|---|---|---|---|---|---|---|---|---|---|
| NA | NA | NA | NA | 10 | Pump & in | 0.71179 | 0.732878 | 6.55E+09 | 0.999773 |
| NA | NA | NA | NA | 10 | Pump & in | 0.65915 | 0.683172 | 5.11E+09 | 0.999829 |
| NA | NA | NA | NA | 10 | Pump & in | 0.69095 | 0.713239 | 6.1E+09 | 0.999711 |
| NA | NA | NA | NA | 10 | Pump & in | 0.78202 | 0.798709 | 6.15E+09 | 0.997466 |
| NA | NA | NA | NA | 10 | Pump & in | 0.81727 | 0.831556 | 6.32E+09 | 0.998761 |
| NA | NA | NA | NA | 10 | Surface gra | 0.82898 | 0.842441 | 7.69E+09 | 0.999317 |
| NA | NA | NA | NA | 10 | Surface gra | 0.79011 | 0.806258 | 5.63E+09 | 0.999435 |
| NA | NA | NA | NA | 10 | Surface gra | 0.47858 | 0.509847 | 7.08E+09 | 0.999516 |
| NA | NA | NA | NA | 10 | Surface gra | 0.56245 | 0.590945 | 8.6E+09 | 0.999222 |
| NA | NA | NA | NA | 10 | Surface gra | 0.84258 | 0.855066 | 7.05E+09 | 0.999453 |
| NA | NA | NA | NA | 10 | Surface gra | 0.8394 | 0.852116 | 7.19E+09 | 0.99938 |
| NA | NA | NA | NA | 10 | Surface gra | 0.83426 | 0.847345 | 6.26E+09 | 0.999393 |
| NA | NA | NA | NA | 10 | Surface gra | 0.83614 | 0.84909 | 7.47E+09 | 0.999492 |
| NA | NA | NA | NA | 10 | Surface gra | 0.81706 | 0.831361 | 8.77E+09 | 0.999118 |
| NA | NA | NA | NA | 10 | Surface gra | 0.81227 | 0.826905 | 7E+09 | 0.999491 |
| NA | NA | NA | NA | 10 | Surface gra | 0.81602 | 0.830394 | 8.41E+09 | 0.99929 |
| NA | NA | NA | NA | 10 | Surface gra | 0.8023 | 0.817622 | 7.05E+09 | 0.999436 |
| NA | NA | NA | NA | 10 | Surface gra | 0.78573 | 0.802172 | 6.89E+09 | 0.999234 |
| NA | NA | NA | NA | 10 | Surface gra | 0.4893 | 0.520277 | 7.5E+09 | 0.999657 |
| NA | NA | NA | NA | 10 | Surface gra | 0.47568 | 0.507022 | 6.83E+09 | 0.999678 |
| NA | NA | NA | NA | 10 | Surface gra | 0.75176 | 0.770409 | 6.42E+09 | 0.998985 |
| NA | NA | NA | NA | 10 | Surface gra | 0.77626 | 0.793329 | 7.03E+09 | 0.999047 |
| NA | NA | NA | NA | 10 | Surface gra | 0.76822 | 0.785815 | 7.04E+09 | 0.999163 |
| NA | NA | NA | NA | 10 | Surface gra | 0.76004 | 0.778162 | 6.02E+09 | 0.999169 |
| NA | NA | NA | NA | 10 | Surface gra | 0.52966 | 0.559372 | 8.08E+09 | 0.999037 |
| NA | NA | NA | NA | 10 | Surface gra | 0.56853 | 0.596781 | 6.89E+09 | 0.998849 |
| NA | NA | NA | NA | 10 | Surface gra | 0.56708 | 0.59539 | 7.4E+09 | 0.998701 |
| NA | NA | NA | NA | 10 | Surface gra | 0.84774 | 0.859852 | 6.68E+09 | 0.998861 |
| NA | NA | NA | NA | 10 | Surface gra | 0.84136 | 0.853934 | 6.56E+09 | 0.999016 |
| NA | NA | NA | NA | 10 | Surface gra | 0.81233 | 0.82696 | 5.78E+09 | 0.999182 |
| NA | NA | NA | NA | 10 | Surface gra | 0.61033 | 0.636767 | 5.9E+09 | 0.999204 |
| NA | NA | NA | NA | 10 | Surface gra | 0.66064 | 0.684583 | 6.49E+09 | 0.999422 |
| NA | NA | NA | NA | 10 | Surface gra | 0.79413 | 0.810007 | 6.19E+09 | 0.999672 |
| NA | NA | NA | NA | 10 | Surface gra | 0.77458 | 0.79176 | 5.91E+09 | 0.999622 |
| NA | NA | NA | NA | 10 | Surface gra | 0.75584 | 0.774231 | 5.55E+09 | 0.999457 |
| NA | NA | NA | NA | 10 | Surface gra | 0.7975 | 0.813149 | 5.16E+09 | 0.999631 |
| NA | NA | NA | NA | 10 | Surface gra | 0.78922 | 0.805428 | 5.19E+09 | 0.999588 |
| NA | NA | NA | NA | 10 | Surface gra | 0.79835 | 0.813941 | 5.5E+09 | 0.999628 |
| NA | NA | NA | NA | 10 | Surface gra | 0.63695 | 0.662108 | 5.6E+09 | 0.998968 |
| NA | NA | NA | NA | 10 | Surface gra | 0.6238 | 0.649601 | 5.12E+09 | 0.998916 |
| NA | NA | NA | NA | 10 | Surface gra | 0.62845 | 0.654026 | 5.41E+09 | 0.999118 |
| NA | NA | NA | NA | 10 | Surface gra | 0.77219 | 0.789526 | 6.28E+09 | 0.999253 |
| NA | NA | NA | NA | 10 | Surface gra | 0.77474 | 0.791909 | 6.22E+09 | 0.999461 |
| NA | NA | NA | NA | 10 | Surface gra | 0.77398 | 0.791199 | 5.14E+09 | 0.999474 |
| NA | NA | NA | NA | 10 | Surface gra | 0.70023 | 0.721991 | 6.28E+09 | 0.998504 |
| NA | NA | NA | NA | 10 | Surface gra | 0.66822 | 0.69176 | 5.73E+09 | 0.982211 |

| | | | | | | | | | |
|---|---|---|---|---|---|---|---|---|---|
| NA | NA | NA | NA | 10 | Surface gra | 0.4311 | 0.463405 | 5.48E+09 | 0.993833 |
| NA | NA | NA | NA | 10 | Surface gra | 0.43705 | 0.469248 | 6E+09 | 0.992807 |
| NA | NA | NA | NA | 10 | Surface gra | 0.52485 | 0.554726 | 5.85E+09 | 0.992971 |
| NA | NA | NA | NA | 10 | Surface gra | 0.61614 | 0.642306 | 6.17E+09 | 0.99893 |
| NA | NA | NA | NA | 10 | Surface gra | 0.58003 | 0.607807 | 5.72E+09 | 0.999399 |
| NA | NA | NA | NA | 10 | Surface gra | 0.75432 | 0.772807 | 6.5E+09 | 0.999025 |
| NA | NA | NA | NA | 10 | Surface gra | 0.76139 | 0.779426 | 6.16E+09 | 0.999231 |
| NA | NA | NA | NA | 10 | Surface gra | 0.74232 | 0.761561 | 5.31E+09 | 0.999078 |
| NA | NA | NA | NA | 10 | Surface gra | 0.75673 | 0.775064 | 6.16E+09 | 0.999096 |
| NA | NA | NA | NA | 10 | Surface gra | 0.75534 | 0.773762 | 6.14E+09 | 0.957673 |
| NA | NA | NA | NA | 10 | Surface gra | 0.63339 | 0.658724 | 6.57E+09 | 0.999338 |
| NA | NA | NA | NA | 10 | Surface gra | 0.65264 | 0.677001 | 5.94E+09 | 0.999276 |
| NA | NA | NA | NA | 10 | Surface gra | 0.832 | 0.845246 | 6.69E+09 | 0.999582 |

| LRstar | diversity | No.OfViral | No.OfAsse | No.Of10kb | No.OfViral | Read.Coun | Subset_hi | ProSyn_cy | Synechoco |
|---|---|---|---|---|---|---|---|---|---|
| 1.8E+11 | 20.25358 | 449 | 766243 | 3026 | 476 | 68421186 | 116384 | 62971 | 12246 |
| 2.66E+11 | 20.50224 | 354 | 617993 | 2174 | 391 | 53659150 | 74134 | 32109 | 8677 |
| 1.22E+11 | 20.66698 | 505 | 914820 | 3322 | 559 | 64122460 | 103653 | 51603 | 13172 |
| 9E+10 | 19.87276 | 597 | 873694 | 3638 | 664 | 61823054 | 87218 | 39065 | 14476 |
| 6.13E+10 | 19.37486 | 418 | 802116 | 3495 | 455 | 63814346 | 69658 | 22579 | 13160 |
| 2.87E+10 | 19.63308 | 690 | 745248 | 4653 | 852 | 89474936 | 92432 | 37108 | 30168 |
| 3.44E+10 | 19.69769 | 461 | 656816 | 3976 | 585 | 64480204 | 68970 | 29119 | 24181 |
| 1.56E+12 | 22.19936 | 669 | 747803 | 2399 | 818 | 80266668 | 95495 | 32779 | 27063 |
| 9.09E+11 | 21.65551 | 819 | 906191 | 3126 | 989 | 98345030 | 114323 | 37145 | 30290 |
| 2.29E+10 | 19.42459 | 534 | 676536 | 4484 | 651 | 80004360 | 78769 | 25298 | 19440 |
| 2.32E+10 | 19.54061 | 665 | 699960 | 4377 | 773 | 84230206 | 82773 | 27988 | 21884 |
| 2.22E+10 | 19.4383 | 492 | 643315 | 4034 | 605 | 71119856 | 71340 | 23579 | 18258 |
| 2.68E+10 | 19.60767 | 716 | 817705 | 4742 | 859 | 82680594 | 85987 | 30206 | 23579 |
| 3.96E+10 | 19.67298 | 1479 | 914753 | 4875 | 1730 | 99725298 | 106491 | 33614 | 25807 |
| 3.84E+10 | 19.55858 | 1131 | 743062 | 3667 | 1337 | 78266386 | 83506 | 26036 | 20439 |
| 4.13E+10 | 19.61555 | 1220 | 814369 | 4093 | 1428 | 93722742 | 96106 | 28958 | 22140 |
| 4.18E+10 | 19.68476 | 1166 | 723712 | 3549 | 1399 | 78905840 | 82598 | 25618 | 19753 |
| 4.52E+10 | 19.7893 | 1102 | 678302 | 3372 | 1315 | 79065000 | 80042 | 24160 | 18458 |
| 1.88E+12 | 22.18722 | 421 | 863694 | 2136 | 523 | 84172220 | 114802 | 26242 | 20160 |
| 1.42E+12 | 22.26722 | 351 | 821132 | 2032 | 437 | 74918418 | 102208 | 24519 | 19156 |
| 7.04E+10 | 19.88077 | 464 | 705986 | 3858 | 556 | 70802418 | 66308 | 20619 | 15528 |
| 5.87E+10 | 19.71258 | 314 | 700318 | 3738 | 376 | 76780930 | 67901 | 20864 | 15884 |
| 6.58E+10 | 19.81187 | 431 | 710813 | 3738 | 536 | 78068024 | 68941 | 20875 | 15518 |
| 6.39E+10 | 19.73512 | 314 | 628797 | 3387 | 369 | 65818308 | 59087 | 18198 | 13836 |
| 2.32E+12 | 21.74449 | 516 | 771851 | 2500 | 630 | 95652504 | 98861 | 32806 | 26738 |
| 8.09E+11 | 21.38517 | 481 | 770859 | 2669 | 601 | 78211238 | 105965 | 48072 | 42824 |
| 7.99E+11 | 21.57274 | 628 | 919265 | 3011 | 778 | 84221020 | 122183 | 57996 | 52148 |
| 1.76E+10 | 19.06389 | 70 | 533243 | 3456 | 84 | 77821164 | 68859 | 18709 | 11803 |
| 1.95E+10 | 19.3138 | 138 | 586290 | 4465 | 180 | 75860140 | 70354 | 21120 | 13866 |
| 2.69E+10 | 19.45614 | 129 | 593389 | 4158 | 154 | 66207316 | 68367 | 20786 | 14005 |
| 2.33E+11 | 21.16654 | 293 | 862392 | 3190 | 331 | 66873880 | 81221 | 28821 | 22234 |
| 1.82E+11 | 20.8676 | 150 | 893786 | 3743 | 174 | 73335592 | 84544 | 28101 | 20191 |
| 5.33E+10 | 19.42236 | 407 | 649421 | 3718 | 498 | 70790496 | 83018 | 30690 | 22199 |
| 7.03E+10 | 19.52226 | 374 | 641709 | 3534 | 460 | 67522430 | 75674 | 26050 | 18024 |
| 7.14E+10 | 19.66378 | 435 | 650335 | 3584 | 532 | 63151524 | 75646 | 27858 | 20371 |
| 3.99E+10 | 19.33591 | 294 | 558959 | 3248 | 359 | 58852098 | 70550 | 28069 | 20875 |
| 4.56E+10 | 19.34558 | 246 | 552322 | 3480 | 313 | 58962712 | 68606 | 25836 | 18464 |
| 4.2E+10 | 19.40676 | 327 | 575996 | 3473 | 413 | 62737418 | 74146 | 28917 | 21490 |
| 1.48E+11 | 20.93425 | 802 | 675558 | 4214 | 966 | 63439676 | 112301 | 62216 | 56117 |
| 1.49E+11 | 20.97207 | 708 | 657103 | 3920 | 865 | 57823428 | 98069 | 52390 | 47040 |
| 1.58E+11 | 20.97964 | 660 | 708153 | 4062 | 807 | 61278090 | 113376 | 65756 | 60229 |
| 4.4E+10 | 20.10432 | 840 | 849011 | 4685 | 982 | 71677280 | 101557 | 47679 | 40301 |
| 4.14E+10 | 20.11243 | 950 | 862814 | 4743 | 1116 | 70995036 | 95855 | 41660 | 34451 |
| 3.46E+10 | 19.93909 | 424 | 665905 | 3969 | 517 | 58519028 | 70001 | 27174 | 21231 |
| 1.09E+11 | 20.32016 | 41 | 736238 | 3285 | 51 | 72560182 | 72418 | 18835 | 11196 |
| 2.24E+14 | 19.21328 | 519 | 555571 | 726 | 554 | 47040264 | 43548 | 9302 | 6359 |

| | | | | | | | | | |
|---|---|---|---|---|---|---|---|---|---|
| 4.2E+13 | 22.58586 | 238 | 823425 | 627 | 270 | 63449906 | 62089 | 13194 | 8825 |
| 8.32E+13 | 22.61081 | 233 | 908238 | 655 | 270 | 69588592 | 68657 | 14415 | 9539 |
| 8.67E+12 | 21.73781 | 744 | 975349 | 1257 | 805 | 66282366 | 67337 | 14302 | 9219 |
| 3.03E+11 | 20.99941 | 40 | 789252 | 2833 | 53 | 70685112 | 71339 | 19402 | 12852 |
| 3.28E+11 | 21.33024 | 157 | 881023 | 2972 | 197 | 64991932 | 68592 | 21178 | 16100 |
| 6.39E+10 | 19.98437 | 277 | 712413 | 4091 | 347 | 72976832 | 67224 | 20031 | 13876 |
| 5.75E+10 | 19.8937 | 231 | 674083 | 4030 | 282 | 68934248 | 63050 | 19338 | 13981 |
| 5.79E+10 | 19.93805 | 244 | 615821 | 3718 | 301 | 59934932 | 55707 | 16742 | 11924 |
| 6.05E+10 | 19.91353 | 253 | 677013 | 3973 | 318 | 69399102 | 62397 | 18422 | 13004 |
| 1.62E+13 | 16.06732 | 37 | 458889 | 124 | 39 | 33381606 | 31454 | 8049 | 6179 |
| 4.05E+11 | 20.87281 | 985 | 806716 | 3017 | 1142 | 73854868 | 608525 | 548923 | 542978 |
| 2.73E+11 | 20.61465 | 1030 | 770614 | 2707 | 1180 | 67233578 | 364781 | 310847 | 305762 |
| 2.51E+10 | 19.69256 | 510 | 675406 | 5187 | 587 | 75426304 | 336959 | 282362 | 274607 |

| Prochloroc | Subset_hig | Subset_hig | Subset_hig | Dictyosteli | X2024101 | X2024101 | X2024101 | X2024102 | X2024102 |
|---|---|---|---|---|---|---|---|---|---|
| 50725 | 54 | 390 | 1168 | 2742 | 179837 | 110161 | 48386 | 12977 | 13846 |
| 23432 | 37 | 2184 | 788 | 2138 | 122526 | 59997 | 40186 | 10436 | 8926 |
| 38431 | 49 | 346 | 279 | 3250 | 123851 | 65538 | 38206 | 6486 | 6763 |
| 24589 | 57 | 385 | 634 | 2980 | 103975 | 51089 | 39999 | 4874 | 5105 |
| 9419 | 43 | 0 | 595 | 3210 | 114501 | 48900 | 45941 | 3218 | 3291 |
| 6940 | 113 | 0 | 518 | 2408 | 317082 | 121564 | 147662 | 22105 | 17029 |
| 4938 | 65 | 0 | 227 | 1727 | 220443 | 84090 | 105163 | 11943 | 9282 |
| 5716 | 99 | 0 | 612 | 2261 | 274938 | 79217 | 125602 | 32321 | 24978 |
| 6855 | 130 | 0 | 719 | 3025 | 318140 | 100583 | 142635 | 35648 | 27614 |
| 5858 | 77 | 102 | 207 | 2012 | 212776 | 87590 | 96806 | 13292 | 9773 |
| 6104 | 92 | 0 | 269 | 2139 | 249831 | 98164 | 114970 | 17727 | 12956 |
| 5321 | 56 | 118 | 155 | 1835 | 194479 | 80495 | 89088 | 11463 | 8284 |
| 6627 | 98 | 0 | 569 | 2333 | 224729 | 90495 | 105651 | 14017 | 10059 |
| 7807 | 238 | 0 | 1294 | 2927 | 198251 | 77473 | 96966 | 32352 | 24616 |
| 5597 | 186 | 661 | 528 | 2063 | 168716 | 65871 | 82366 | 24957 | 18263 |
| 6818 | 213 | 494 | 566 | 2535 | 219674 | 86909 | 104694 | 31736 | 23675 |
| 5865 | 185 | 414 | 1002 | 2250 | 177650 | 66862 | 84760 | 28611 | 21778 |
| 5702 | 182 | 0 | 1131 | 2088 | 197144 | 73650 | 91903 | 31576 | 23637 |
| 6082 | 65 | 47 | 226 | 2327 | 399474 | 108335 | 140691 | 19539 | 12941 |
| 5363 | 70 | 56 | 129 | 2036 | 304875 | 76992 | 106331 | 13435 | 8956 |
| 5091 | 53 | 152 | 115 | 1712 | 184716 | 74842 | 78950 | 5953 | 4381 |
| 4980 | 57 | 0 | 227 | 1680 | 245497 | 99856 | 108289 | 7147 | 4767 |
| 5357 | 75 | 660 | 261 | 1779 | 242809 | 97920 | 105035 | 8214 | 5939 |
| 4362 | 59 | 173 | 120 | 1383 | 199803 | 81546 | 87316 | 5610 | 4031 |
| 6068 | 74 | 0 | 309 | 2953 | 646460 | 151105 | 224358 | 43363 | 26462 |
| 5248 | 57 | 0 | 271 | 2337 | 427323 | 97365 | 172021 | 25482 | 17049 |
| 5848 | 83 | 99 | 278 | 2609 | 457353 | 98527 | 192467 | 37108 | 20203 |
| 6906 | 16 | 65 | 40 | 3011 | 163561 | 66433 | 64658 | 1779 | 1255 |
| 7254 | 25 | 150 | 149 | 3205 | 127337 | 51377 | 52840 | 2667 | 2097 |
| 6781 | 31 | 0 | 71 | 2675 | 116975 | 47179 | 51527 | 2507 | 1837 |
| 6587 | 51 | 0 | 141 | 2674 | 146955 | 44853 | 65916 | 19369 | 14650 |
| 7910 | 40 | 0 | 109 | 3370 | 158092 | 53321 | 64811 | 11914 | 8312 |
| 8491 | 22 | 0 | 375 | 3588 | 167179 | 53306 | 68904 | 4726 | 2907 |
| 8026 | 29 | 0 | 292 | 3511 | 158056 | 47888 | 57276 | 5141 | 3086 |
| 7487 | 35 | 0 | 141 | 3229 | 157421 | 46786 | 62375 | 5548 | 3494 |
| 7194 | 19 | 0 | 204 | 3086 | 119612 | 35142 | 54708 | 3212 | 2190 |
| 7372 | 26 | 0 | 163 | 3113 | 129614 | 41047 | 54642 | 3286 | 2218 |
| 7427 | 22 | 0 | 101 | 3045 | 139444 | 40168 | 58667 | 4234 | 2756 |
| 6099 | 79 | 253 | 384 | 2368 | 231211 | 52796 | 141745 | 21518 | 16853 |
| 5350 | 59 | 0 | 306 | 2303 | 192196 | 46056 | 119129 | 17832 | 13348 |
| 5527 | 92 | 0 | 286 | 2291 | 247100 | 57839 | 155188 | 19638 | 14615 |
| 7378 | 84 | 0 | 204 | 2855 | 157625 | 46105 | 100629 | 13042 | 9685 |
| 7209 | 103 | 0 | 246 | 2926 | 138947 | 42699 | 86311 | 14171 | 10445 |
| 5943 | 36 | 0 | 137 | 2295 | 123149 | 46662 | 64989 | 9403 | 6736 |
| 7639 | 17 | 0 | 131 | 2863 | 170871 | 78546 | 66635 | 5592 | 4000 |
| 2943 | 118 | 0 | 206 | 1650 | 323415 | 140161 | 226337 | 62908 | 25413 |

| | | | | | | | | | |
|---|---|---|---|---|---|---|---|---|---|
| 4369 | 75 | 0 | 213 | 2690 | 263532 | 115451 | 96627 | 38051 | 29794 |
| 4876 | 78 | 0 | 435 | 2904 | 288338 | 127446 | 106257 | 40669 | 32053 |
| 5083 | 155 | 1 | 520 | 3169 | 204112 | 91449 | 78380 | 75083 | 60965 |
| 6550 | 23 | 70 | 52 | 2650 | 217824 | 98153 | 91751 | 9316 | 6341 |
| 5078 | 38 | 0 | 68 | 2005 | 123837 | 47642 | 59245 | 4338 | 3314 |
| 6155 | 34 | 0 | 77 | 2754 | 128420 | 52454 | 57400 | 3779 | 2771 |
| 5357 | 25 | 0 | 410 | 2318 | 136223 | 55423 | 63154 | 3653 | 2341 |
| 4818 | 29 | 0 | 58 | 2093 | 113122 | 45623 | 51396 | 3380 | 2417 |
| 5418 | 25 | 184 | 258 | 2412 | 133880 | 55126 | 59829 | 3643 | 2479 |
| 1870 | 18 | 0 | 834 | 739 | 534752 | 242387 | 466416 | 79220 | 4669 |
| 5945 | 166 | 117 | 583 | 2742 | 1239541 | 83758 | 1126331 | 43808 | 32369 |
| 5085 | 182 | 0 | 918 | 2452 | 751184 | 63353 | 648973 | 44985 | 34019 |
| 7755 | 44 | 0 | 427 | 2597 | 680102 | 81218 | 602611 | 24352 | 18198 |

| X2024102: | amoeba_g | Subset_hig | ProSyn_cy | Synechoco | Prochloroc | Subset_hig | Subset_hig | Subset_hig | Dictyosteli |
|---|---|---|---|---|---|---|---|---|---|
| 23343 | 74614 | 1700.994 | 920.3436 | 178.9797 | 741.3639 | 0.789229 | 5.699989 | 17.07074 | 40.07531 |
| 18392 | 83375 | 1381.572 | 598.3882 | 161.7059 | 436.6823 | 0.689538 | 40.70135 | 14.68529 | 39.84409 |
| 11090 | 32693 | 1616.485 | 804.757 | 205.4194 | 599.3376 | 0.764163 | 5.395925 | 4.35105 | 50.68427 |
| 9432 | 4960 | 1410.768 | 631.884 | 234.1521 | 397.7319 | 0.921986 | 6.22745 | 10.25507 | 48.20208 |
| 6798 | 4101 | 1091.573 | 353.8233 | 206.2232 | 147.6 | 0.67383 | 0 | 9.323922 | 50.30217 |
| 48303 | 52088 | 1033.049 | 414.7307 | 337.167 | 77.56362 | 1.262924 | 0 | 5.789331 | 26.91256 |
| 26277 | 32970 | 1069.631 | 451.596 | 375.0143 | 76.58164 | 1.008061 | 0 | 3.52046 | 26.78341 |
| 64356 | 161442 | 1189.722 | 408.3762 | 337.1636 | 71.21262 | 1.233389 | 0 | 7.624585 | 28.1686 |
| 66779 | 230546 | 1162.469 | 377.7008 | 307.9973 | 69.70357 | 1.321877 | 0 | 7.310995 | 30.75905 |
| 28590 | 12435 | 984.5588 | 316.2078 | 242.9868 | 73.22101 | 0.962448 | 1.274931 | 2.587359 | 25.14863 |
| 36972 | 14157 | 982.6997 | 332.2798 | 259.8118 | 72.46806 | 1.092245 | 0 | 3.193629 | 25.39469 |
| 23904 | 10229 | 1003.095 | 331.5389 | 256.7216 | 74.81736 | 0.787403 | 1.659171 | 2.179419 | 25.80151 |
| 30161 | 11505 | 1039.99 | 365.3336 | 285.1818 | 80.15182 | 1.185284 | 0 | 6.881905 | 28.21702 |
| 73673 | 26916 | 1067.843 | 337.0659 | 258.7809 | 78.28505 | 2.386556 | 0 | 12.97564 | 29.35063 |
| 59761 | 28125 | 1066.946 | 332.6588 | 261.1466 | 71.51218 | 2.376499 | 8.445516 | 6.746191 | 26.3587 |
| 71234 | 42099 | 1025.429 | 308.9752 | 236.2287 | 72.74648 | 2.272661 | 5.270866 | 6.039089 | 27.04786 |
| 64491 | 38583 | 1046.792 | 324.6654 | 250.3364 | 74.3291 | 2.344567 | 5.24676 | 12.69868 | 28.515 |
| 69264 | 39908 | 1012.357 | 305.5714 | 233.4535 | 72.11788 | 2.301903 | 0 | 14.30469 | 26.40865 |
| 37215 | 279989 | 1363.894 | 311.7656 | 239.5089 | 72.25662 | 0.772226 | 0.558379 | 2.684971 | 27.6457 |
| 24303 | 208681 | 1364.257 | 327.276 | 255.6915 | 71.58453 | 0.93435 | 0.74748 | 1.721873 | 27.17623 |
| 11547 | 4661 | 936.5217 | 291.2189 | 219.3145 | 71.90432 | 0.748562 | 2.146819 | 1.624238 | 24.17997 |
| 14608 | 8011 | 884.3472 | 271.7341 | 206.8743 | 64.85986 | 0.742372 | 0 | 2.956463 | 21.88043 |
| 16090 | 8470 | 883.0888 | 267.395 | 198.7754 | 68.61964 | 0.960701 | 8.454166 | 3.343238 | 22.78782 |
| 11140 | 5656 | 897.7289 | 276.4884 | 210.2151 | 66.27335 | 0.896407 | 2.628448 | 1.823201 | 21.01239 |
| 95515 | 674561 | 1033.543 | 342.9706 | 279.5327 | 63.43796 | 0.773634 | 0 | 3.230443 | 30.87217 |
| 50336 | 360257 | 1354.856 | 614.6431 | 547.5428 | 67.10033 | 0.728796 | 0 | 3.464975 | 29.88062 |
| 85753 | 485462 | 1450.742 | 688.6167 | 619.1803 | 69.43635 | 0.985502 | 1.175479 | 3.300839 | 30.97801 |
| 3203 | 3343 | 884.8364 | 240.4102 | 151.6683 | 88.74193 | 0.2056 | 0.835248 | 0.513999 | 38.69127 |
| 4778 | 2910 | 927.4172 | 278.4071 | 182.7837 | 95.62334 | 0.329554 | 1.977323 | 1.964141 | 42.2488 |
| 4396 | 2762 | 1032.62 | 313.9532 | 211.5325 | 102.4207 | 0.468226 | 0 | 1.072389 | 40.40339 |
| 51099 | 82819 | 1214.54 | 430.9754 | 332.4766 | 98.49885 | 0.76263 | 0 | 2.108447 | 39.98572 |
| 27279 | 87871 | 1152.837 | 383.1837 | 275.3233 | 107.8603 | 0.545438 | 0 | 1.486318 | 45.95313 |
| 11351 | 81495 | 1172.728 | 433.5328 | 313.5873 | 119.9455 | 0.310776 | 0 | 5.297321 | 50.68477 |
| 12093 | 89258 | 1120.724 | 385.7977 | 266.9335 | 118.8642 | 0.429487 | 0 | 4.324489 | 51.99754 |
| 13518 | 93995 | 1197.849 | 441.1295 | 322.5734 | 118.5561 | 0.554223 | 0 | 2.232725 | 51.13099 |
| 6434 | 44186 | 1198.768 | 476.9414 | 354.7027 | 122.2386 | 0.322843 | 0 | 3.466317 | 52.43653 |
| 6008 | 43106 | 1163.549 | 438.1752 | 313.1471 | 125.0282 | 0.440957 | 0 | 2.764459 | 52.79608 |
| 7801 | 50021 | 1181.847 | 460.9211 | 342.5388 | 118.3823 | 0.350668 | 0 | 1.609885 | 48.53563 |
| 52889 | 94402 | 1770.201 | 980.7112 | 884.5726 | 96.13857 | 1.245277 | 3.988041 | 6.052994 | 37.3268 |
| 44020 | 83872 | 1696.008 | 906.0341 | 813.5111 | 92.52305 | 1.020348 | 0 | 5.291973 | 39.82815 |
| 48787 | 126420 | 1850.188 | 1073.075 | 982.8799 | 90.19537 | 1.501352 | 0 | 4.667247 | 37.38694 |
| 35952 | 7458 | 1416.865 | 665.1899 | 562.2563 | 102.9336 | 1.171919 | 0 | 2.84609 | 39.83131 |
| 40569 | 6761 | 1350.165 | 586.8016 | 485.2593 | 101.5423 | 1.450806 | 0 | 3.465031 | 41.21415 |
| 24211 | 5356 | 1196.209 | 464.3618 | 362.8051 | 101.5567 | 0.615185 | 0 | 2.341119 | 39.21801 |
| 11496 | 38852 | 998.0405 | 259.5776 | 154.2995 | 105.2781 | 0.234288 | 0 | 1.805398 | 39.4569 |
| 145849 | 95755 | 925.7601 | 197.7455 | 135.1821 | 62.56342 | 2.508489 | 0 | 4.379227 | 35.07633 |

| | | | | | | | | | |
|---|---|---|---|---|---|---|---|---|---|
| 89103 | 165088 | 978.5515 | 207.9436 | 139.0861 | 68.85747 | 1.182035 | 0 | 3.356979 | 42.39565 |
| 94542 | 184941 | 986.6129 | 207.146 | 137.0771 | 70.06896 | 1.120873 | 0 | 6.251025 | 41.73098 |
| 234678 | 105391 | 1015.911 | 215.7738 | 139.0868 | 76.68706 | 2.33848 | 0.015087 | 7.845224 | 47.8106 |
| 17623 | 87266 | 1009.251 | 274.485 | 181.8205 | 92.66449 | 0.325387 | 0.990308 | 0.735657 | 37.49021 |
| 7989 | 18665 | 1055.393 | 325.8558 | 247.7231 | 78.13278 | 0.584688 | 0 | 1.046284 | 30.84998 |
| 8283 | 6079 | 921.1691 | 274.4844 | 190.1425 | 84.34184 | 0.465901 | 0 | 1.055129 | 37.738 |
| 7682 | 7290 | 914.6397 | 280.5282 | 202.8165 | 77.71173 | 0.362664 | 0 | 5.947697 | 33.62625 |
| 7140 | 7106 | 929.458 | 279.3363 | 198.9491 | 80.38718 | 0.483858 | 0 | 0.967716 | 34.9212 |
| 7624 | 7663 | 899.1039 | 265.4501 | 187.3799 | 78.07017 | 0.360235 | 2.651331 | 3.717627 | 34.75549 |
| 139568 | 54132 | 942.2554 | 241.1208 | 185.1019 | 56.01887 | 0.539219 | 0 | 24.98382 | 22.13794 |
| 199239 | 171685 | 8239.47 | 7432.455 | 7351.96 | 80.49571 | 2.247651 | 1.584188 | 7.89386 | 37.12687 |
| 181252 | 171975 | 5425.578 | 4623.389 | 4547.757 | 75.63185 | 2.706981 | 0 | 13.65389 | 36.46987 |
| 93577 | 21061 | 4467.394 | 3743.548 | 3640.733 | 102.8156 | 0.583351 | 0 | 5.661155 | 34.43096 |

| X2024101 | X20241018_pro_all_genomes | X20241018_syn_all_genomes_ | X20241021_cyanophage_all_ge |
|---|---|---|---|
| 2628.382 | 1610.042246 | 707.1786215 | 189.6634765 |
| 2283.413 | 1118.113127 | 748.9123477 | 194.4868676 |
| 1931.476 | 1022.075572 | 595.8286691 | 101.1502054 |
| 1681.816 | 826.3745754 | 646.9916546 | 78.83790406 |
| 1794.283 | 766.2853741 | 719.9164903 | 50.42753239 |
| 3543.808 | 1358.637462 | 1650.316911 | 247.0524259 |
| 3418.77 | 1304.121184 | 1630.934666 | 185.2196373 |
| 3425.307 | 986.9227411 | 1564.808944 | 402.6702591 |
| 3234.937 | 1022.75631 | 1450.352905 | 362.4789173 |
| 2659.555 | 1094.815333 | 1210.009055 | 166.1409453 |
| 2966.05 | 1165.425145 | 1364.949766 | 210.4589415 |
| 2734.525 | 1131.821752 | 1252.645956 | 161.1786166 |
| 2718.038 | 1094.513182 | 1277.821008 | 169.5319218 |
| 1987.971 | 776.8640611 | 972.3310127 | 324.411164 |
| 2155.664 | 841.6256757 | 1052.380264 | 318.8725234 |
| 2343.871 | 927.2989474 | 1117.060788 | 338.6157866 |
| 2251.418 | 847.3644029 | 1074.19172 | 362.5967356 |
| 2493.442 | 931.512047 | 1162.372731 | 399.3676089 |
| 4745.913 | 1287.063594 | 1671.465954 | 232.1312186 |
| 4069.427 | 1027.677867 | 1419.290514 | 179.3283996 |
| 2608.894 | 1057.054294 | 1115.074912 | 84.07904939 |
| 3197.369 | 1300.531265 | 1410.363224 | 93.08300902 |
| 3110.223 | 1254.290745 | 1345.429212 | 105.2159332 |
| 3035.675 | 1238.956188 | 1326.621766 | 85.23464322 |
| 6758.422 | 1579.728639 | 2345.552815 | 453.3388901 |
| 5463.703 | 1244.897824 | 2199.440955 | 325.8099559 |
| 5430.39 | 1169.862346 | 2285.260853 | 440.6025954 |
| 2101.755 | 853.662379 | 830.8536737 | 22.86010525 |
| 1678.576 | 677.2594936 | 696.544984 | 35.15680303 |
| 1766.799 | 712.5949646 | 778.2674652 | 37.86590594 |
| 2197.495 | 670.7102983 | 985.6763209 | 289.6347572 |
| 2155.734 | 727.0821513 | 883.7591439 | 162.4586326 |
| 2361.602 | 753.0106866 | 973.350999 | 66.76037416 |
| 2340.793 | 709.2161819 | 848.2514625 | 76.1376627 |
| 2492.751 | 740.8530632 | 987.7037963 | 87.85219498 |
| 2032.417 | 597.1239972 | 929.5845324 | 54.57749357 |
| 2198.237 | 696.1518324 | 926.7212811 | 55.7301367 |
| 2222.661 | 640.2558677 | 935.1197717 | 67.48763553 |
| 3644.58 | 832.2236702 | 2234.3273 | 339.1883653 |
| 3323.843 | 796.4937672 | 2060.220297 | 308.3871126 |
| 4032.436 | 943.8773304 | 2532.520188 | 320.4734351 |
| 2199.093 | 643.2303235 | 1403.917671 | 181.9544492 |
| 1957.137 | 601.4364159 | 1215.732886 | 199.6055048 |
| 2104.427 | 797.3816653 | 1110.56185 | 160.6827783 |
| 2354.887 | 1082.494528 | 918.3411365 | 77.06706138 |
| 6875.28 | 2979.596373 | 4811.558881 | 1337.322427 |

| | | | |
|---|---|---|---|
| 4153.387 | 1819.561403 | 1522.886417 | 599.7014401 |
| 4143.467 | 1831.420874 | 1526.931311 | 584.4205039 |
| 3079.431 | 1379.688227 | 1182.516629 | 1132.774892 |
| 3081.611 | 1388.595098 | 1298.024399 | 131.7957875 |
| 1905.421 | 733.0448339 | 911.5746859 | 66.74674635 |
| 1759.737 | 718.7760631 | 786.5509974 | 51.78355783 |
| 1976.129 | 803.9980359 | 916.1483853 | 52.99252702 |
| 1887.414 | 761.2088389 | 857.5299627 | 56.39449128 |
| 1929.132 | 794.3330448 | 862.1004923 | 52.49347463 |
| 16019.36 | 7261.094628 | 13972.24567 | 2373.163232 |
| 16783.47 | 1134.089089 | 15250.59932 | 593.1633376 |
| 11172.75 | 942.2821436 | 9652.513213 | 669.0853192 |
| 9016.775 | 1076.786157 | 7989.401151 | 322.8581902 |

| X20241021_proc_phage_all_ge | X20241021_syn_phage_all_gen | amoeba_genome_5_all_rpm |
|---|---|---|
| 202.3642209 | 341.1662581 | 1090.510182 |
| 166.3462802 | 342.7560817 | 1553.789056 |
| 105.4700646 | 172.9503204 | 509.852554 |
| 82.57437428 | 152.5644463 | 80.22897089 |
| 51.57147579 | 106.5277704 | 64.26454641 |
| 190.3214549 | 539.849506 | 582.1518554 |
| 143.9511575 | 407.5204229 | 511.3197223 |
| 311.1877025 | 801.7773953 | 2011.320565 |
| 280.7869396 | 679.0277048 | 2344.256746 |
| 122.1558425 | 357.3555241 | 155.4290291 |
| 153.8165536 | 438.9399214 | 168.0750965 |
| 116.4794259 | 336.108667 | 143.8276253 |
| 121.6609547 | 364.7893483 | 139.1499437 |
| 246.8380691 | 738.7593868 | 269.9014246 |
| 233.3441076 | 763.5589562 | 359.3496702 |
| 252.6067793 | 760.0503195 | 449.1866019 |
| 275.9998499 | 817.31593 | 488.975214 |
| 298.9565547 | 876.0387023 | 504.7492569 |
| 153.7443114 | 442.1292441 | 3326.382505 |
| 119.5433678 | 324.3928616 | 2785.443227 |
| 61.87641784 | 163.0876505 | 65.83108503 |
| 62.08572884 | 190.2555752 | 104.3358032 |
| 76.07468072 | 206.1023089 | 108.4951247 |
| 61.24435772 | 169.2538192 | 85.93353691 |
| 276.6472271 | 998.5624631 | 7052.2043 |
| 217.9865763 | 643.5903751 | 4606.205057 |
| 239.8807329 | 1018.189996 | 5764.142966 |
| 16.12671843 | 41.15846944 | 42.95746592 |
| 27.64297561 | 62.98432879 | 38.3600663 |
| 27.74617838 | 66.39749601 | 41.71744404 |
| 219.0690895 | 764.1099933 | 1238.435694 |
| 113.3419636 | 371.9749068 | 1198.204004 |
| 41.06483447 | 160.3463832 | 1151.213858 |
| 45.70333147 | 179.0960426 | 1321.901478 |
| 55.32724753 | 214.0565919 | 1488.404302 |
| 37.21192743 | 109.3249046 | 750.7973632 |
| 37.61699428 | 101.8949061 | 731.0722071 |
| 43.92912695 | 124.3436572 | 797.3072784 |
| 265.6539419 | 833.6896298 | 1488.059302 |
| 230.8406897 | 761.2831256 | 1450.484741 |
| 238.5028646 | 796.1573215 | 2063.053858 |
| 135.1195246 | 501.5815332 | 104.0497072 |
| 147.1229622 | 571.4343183 | 95.23201031 |
| 115.1078586 | 413.7286764 | 91.52578542 |
| 55.12665335 | 158.4340017 | 535.445184 |
| 540.239315 | 3100.514062 | 2035.596569 |

| | | |
|---|---|---|
| 469.5672835 | 1404.304681 | 2601.863587 |
| 460.6071064 | 1358.584752 | 2657.633883 |
| 919.7770641 | 3540.579707 | 1590.030748 |
| 89.70771667 | 249.316999 | 1234.573979 |
| 50.99094454 | 122.9229499 | 287.189493 |
| 37.97095495 | 113.5017755 | 83.30040964 |
| 33.95989755 | 111.4395271 | 105.7529488 |
| 40.32706669 | 119.1291916 | 118.5619098 |
| 35.72092331 | 109.8573293 | 110.4192962 |
| 139.8674468 | 4180.985181 | 1621.611614 |
| 438.2784896 | 2697.709784 | 2324.626726 |
| 505.9822935 | 2695.855336 | 2557.873686 |
| 241.2686163 | 1240.641461 | 279.2261967 |
