## Supplementary Data 2 for "Sampling Microbial Dynamics in the Salish Sea Estuary: Evaluating Methods to Capture Cyanobacteria and Cyanophage"

Prochlorococcus reference genomes:

GCA_000007925.1_ASM792v1

GCA_000011465.1_ASM1146v1

GCA_000011485.1_ASM1148v1

GCA_000012465.1_ASM1246v1

GCA_000012645.1_ASM1264v1

GCA_000015645.1_ASM1564v1

GCA_000015665.1_ASM1566v1

GCA_000015685.1_ASM1568v1

GCA_000015705.1_ASM1570v1

GCA_000015965.1_ASM1596v1

GCA_000018065.1_ASM1806v1

GCA_000757845.1_ASM75784v1

GCA_000757865.1_ASM75786v1

GCA_027359355.1_ASM2735935v1

GCA_027359375.1_ASM2735937v1

GCA_027359395.1_ASM2735939v1

GCA_027359415.1_ASM2735941v1

GCA_027359475.1_ASM2735947v1

GCA_027359525.1_ASM2735952v1

GCA_027359575.1_ASM2735957v1

GCA_027359595.1_ASM2735959v1

GCA_034092375.1_ASM3409237v1

GCA_034092395.1_ASM3409239v1

GCA_034092415.1_ASM3409241v1

GCA_034092465.1_ASM3409246v1

GCA_034093315.1_ASM3409331v1

Synechococcus reference genomes:

GCA_000010065.1_ASM1006v1

GCA_000012505.1_ASM1250v1

GCA_000012525.1_ASM1252v1

GCA_000012625.1_ASM1262v1

GCA_000013205.1_ASM1320v1

GCA_000013225.1_ASM1322v1

GCA_000014585.1_ASM1458v1

GCA_000019485.1_ASM1948v1

GCA_000063505.1_ASM6350v1

GCA_000063525.1_ASM6352v1

GCA_000161795.2_ASM16179v2

GCA_000179235.2_ASM17923v2

GCA_000316685.1_ASM31668v1

GCA_000317085.1_ASM31708v1

GCA_000737535.1_ASM73753v1

GCA_000737575.1_ASM73757v1

GCA_000737595.1_ASM73759v1

GCA_000817325.1_ASM81732v1

GCA_001182765.1_WH8103.1

GCA_001521855.1_ASM152185v1

GCA_001693255.1_ASM169325v1

GCA_001693275.1_ASM169327v1

GCA_001693295.1_ASM169329v1

GCA_001885215.1_ASM188521v1

GCA_002356215.1_ASM235621v1

GCA_003846445.2_ASM384644v2

GCA_003957805.1_ASM395780v1

GCA_004209775.1_ASM420977v1

GCA_008807075.1_ASM880707v1

GCA_009498715.2_ASM949871v2

GCA_014217875.1_ASM1421787v1

GCA_014279535.1_ASM1427953v1

GCA_014279555.1_ASM1427955v1

GCA_014279595.1_ASM1427959v1

GCA_014279615.1_ASM1427961v1

GCA_014279635.1_ASM1427963v1

GCA_014279655.1_ASM1427965v1

GCA_014279755.1_ASM1427975v1

GCA_014279775.1_ASM1427977v1

GCA_014279795.1_ASM1427979v1

GCA_014279815.1_ASM1427981v1

GCA_014279835.1_ASM1427983v1

GCA_014279855.1_ASM1427985v1

GCA_014279875.1_ASM1427987v1

GCA_014279895.1_ASM1427989v1

GCA_014279955.1_ASM1427995v1

GCA_014279975.1_ASM1427997v1

GCA_014280015.1_ASM1428001v1

GCA_014280035.1_ASM1428003v1

GCA_014280055.1_ASM1428005v1

GCA_014280075.1_ASM1428007v1

GCA_014280095.1_ASM1428009v1

GCA_014280115.1_ASM1428011v1

GCA_014280175.1_ASM1428017v1

GCA_014280195.1_ASM1428019v1

GCA_014280215.1_ASM1428021v1

GCA_014304795.1_ASM1430479v1

GCA_014304815.1_ASM1430481v1

GCA_015840335.1_ASM1584033v1

GCA_015840525.1_ASM1584052v1

GCA_015840715.1_ASM1584071v1

GCA_015840915.1_ASM1584091v1

GCA_015841355.1_ASM1584135v1

GCA_022984195.1_ASM2298419v1

GCA_023078835.1_ASM2307883v1

GCA_023115425.1_ASM2311542v1

GCA_030544905.1_ASM3054490v1

GCA_036630065.1_ASM3663006v1

GCA_036630075.1_ASM3663007v1

GCA_036630085.1_ASM3663008v1

GCA_036630095.1_ASM3663009v1

GCA_036630105.1_ASM3663010v1

GCA_036630115.1_ASM3663011v1

GCA_040371545.1_ASM4037154v1

GCA_963920465.1_KAC_114

GCA_963920475.1_KAC_105

GCA_963920495.1_KAC_102

GCA_963920505.1_KAC_106

Prochlorococcus phage reference genomes:

GCA_000857985.1_ViralProj15136

GCA_000858745.1_ViralProj15134

GCA_000859585.1_ViralProj15135

GCA_000890835.1_ViralProj64703

GCA_000891815.1_ViralProj64707

GCA_000892455.1_ViralProj64697

GCA_000892475.1_ViralProj64705

GCA_000892495.1_ViralProj64713

GCA_000893395.1_ViralProj64717

GCA_000904555.1_ViralProj195517

GCA_000905515.1_ViralProj195499

GCA_000906175.1_ViralProj195518

GCA_000907775.1_ViralProj209210

GCA_001503075.1_ViralProj307841

GCA_002630285.1_ASM263028v1

GCA_002745975.1_ASM274597v1

GCA_002991095.1_ASM299109v1

GCA_009674125.1_ASM967412v1

GCA_013348255.1_ASM1334825v1

GCA_024499585.1_ASM2449958v1

GCA_024499595.1_ASM2449959v1

GCA_024499605.1_ASM2449960v1

GCA_024499615.1_ASM2449961v1

GCA_024499625.1_ASM2449962v1

GCA_024499635.1_ASM2449963v1

GCA_024499645.1_ASM2449964v1

GCA_024499655.1_ASM2449965v1

GCA_024499665.1_ASM2449966v1

GCA_024499675.1_ASM2449967v1

GCA_024499685.1_ASM2449968v1

GCA_024499695.1_ASM2449969v1

GCA_024499705.1_ASM2449970v1

GCA_024499715.1_ASM2449971v1

GCA_024499725.1_ASM2449972v1

GCA_024499735.1_ASM2449973v1

GCA_024499745.1_ASM2449974v1

GCA_024499755.1_ASM2449975v1

Synechococcus phage reference genomes:

GCA_000849825.2_ViralProj14628

GCA_000870005.1_ViralProj17541

GCA_000873225.1_ViralProj19763

GCA_000886695.1_ViralProj39923

GCA_000890815.1_ViralProj64695

GCA_000890855.1_ViralProj64711

GCA_000891035.1_ViralProj66397

GCA_000891795.1_ViralProj64699

GCA_000891835.1_ViralProj64715

GCA_000891995.1_ViralProj66393

GCA_000892775.1_ViralProj67251

GCA_000893355.1_ViralProj64701

GCA_000893375.1_ViralProj64709

GCA_000894995.1_ViralProj76741

GCA_000896775.1_ViralProj82651

GCA_000901695.1_ViralProj181073

GCA_000902315.1_ViralProj181072

GCA_000904235.1_ViralProj192853

GCA_000904455.1_ViralProj195489

GCA_000905455.1_ViralProj195487

GCA_000906055.1_ViralProj195486

GCA_000906915.1_ViralProj195484

GCA_000906935.1_ViralProj195488

GCA_000907715.1_ViralProj209065

GCA_000908735.1_ViralProj209067

GCA_000917315.1_ViralProj240044

GCA_000918375.1_ViralProj240047

GCA_000925075.1_ViralProj266790

GCA_000925935.1_ViralProj266791

GCA_000929515.1_ViralProj266784

GCA_000929555.1_ViralProj266792

GCA_000982305.1_ViralProj282477

GCA_000982325.1_ViralProj282474

GCA_000982345.1_ViralProj282478

GCA_000982365.1_ViralProj282475

GCA_000982385.1_ViralProj282479

GCA_001019915.1_ViralProj284772

GCA_001019995.1_ViralProj285603

GCA_001503235.1_ViralProj307875

GCA_001881915.1_ViralProj353683

GCA_001881955.1_ViralProj353680

GCA_001882035.1_ViralProj353681

GCA_001882115.1_ViralProj353686

GCA_001882155.1_ViralProj353682

GCA_001885345.1_ViralProj353687

GCA_002366065.1_ViralProj353501

GCA_002412125.1_ASM241212v1

GCA_002534635.1_ASM253463v1

GCA_002534655.1_ASM253465v1

GCA_002546055.1_ASM254605v1

GCA_002546075.1_ASM254607v1

GCA_002546105.1_ASM254610v1

GCA_002546145.1_ASM254614v1

GCA_002546175.1_ASM254617v1

GCA_002546205.1_ASM254620v1

GCA_002546255.1_ASM254625v1

GCA_002546285.1_ASM254628v1

GCA_002546315.1_ASM254631v1

GCA_002546355.1_ASM254635v1

GCA_002546415.1_ASM254641v1

GCA_002546445.1_ASM254644v1

GCA_002546485.1_ASM254648v1

GCA_002546525.1_ASM254652v1

GCA_002546555.1_ASM254655v1

GCA_002546595.1_ASM254659v1

GCA_002546645.1_ASM254664v1

GCA_002546675.1_ASM254667v1

GCA_002546705.1_ASM254670v1

GCA_002546735.1_ASM254673v1

GCA_002546755.1_ASM254675v1

GCA_002546775.1_ASM254677v1

GCA_002546825.1_ASM254682v1

GCA_002546885.1_ASM254688v1

GCA_002546915.1_ASM254691v1

GCA_002546945.1_ASM254694v1

GCA_002546985.1_ASM254698v1

GCA_002547015.1_ASM254701v1

GCA_002547045.1_ASM254704v1

GCA_002547095.1_ASM254709v1

GCA_002547125.1_ASM254712v1

GCA_002547175.1_ASM254717v1

GCA_002547225.1_ASM254722v1

GCA_002547255.1_ASM254725v1

GCA_002547275.1_ASM254727v1

GCA_002547325.1_ASM254732v1

GCA_002547385.1_ASM254738v1

GCA_002547405.1_ASM254740v1

GCA_002547435.1_ASM254743v1

GCA_002547465.1_ASM254746v1

GCA_002547515.1_ASM254751v1

GCA_002547575.1_ASM254757v1

GCA_002547595.1_ASM254759v1

GCA_002547625.1_ASM254762v1

GCA_002547655.1_ASM254765v1

GCA_002547705.1_ASM254770v1

GCA_002547745.1_ASM254774v1

GCA_002547775.1_ASM254777v1

GCA_002547795.1_ASM254779v1

GCA_002547815.1_ASM254781v1

GCA_002547875.1_ASM254787v1

GCA_002547895.1_ASM254789v1

GCA_002547955.1_ASM254795v1

GCA_002547975.1_ASM254797v1

GCA_002547995.1_ASM254799v1

GCA_002548035.1_ASM254803v1

GCA_002548055.1_ASM254805v1

GCA_002548105.1_ASM254810v1

GCA_002548135.1_ASM254813v1

GCA_002548155.1_ASM254815v1

GCA_002548185.1_ASM254818v1

GCA_002548215.1_ASM254821v1

GCA_002548235.1_ASM254823v1

GCA_002548275.1_ASM254827v1

GCA_002548325.1_ASM254832v1

GCA_002548365.1_ASM254836v1

GCA_002548405.1_ASM254840v1

GCA_002548435.1_ASM254843v1

GCA_002548475.1_ASM254847v1

GCA_002548515.1_ASM254851v1

GCA_002548555.1_ASM254855v1

GCA_002548575.1_ASM254857v1

GCA_002548615.1_ASM254861v1

GCA_002548655.1_ASM254865v1

GCA_002548685.1_ASM254868v1

GCA_002548725.1_ASM254872v1

GCA_002548765.1_ASM254876v1

GCA_002548785.1_ASM254878v1

GCA_002548815.1_ASM254881v1

GCA_002548835.1_ASM254883v1

GCA_002548855.1_ASM254885v1

GCA_002548875.1_ASM254887v1

GCA_002548905.1_ASM254890v1

GCA_002548965.1_ASM254896v1

GCA_002549015.1_ASM254901v1

GCA_002549035.1_ASM254903v1

GCA_002549055.1_ASM254905v1

GCA_002549075.1_ASM254907v1

GCA_002549095.1_ASM254909v1

GCA_002549155.1_ASM254915v1

GCA_002549215.1_ASM254921v1

GCA_002549235.1_ASM254923v1

GCA_002549255.1_ASM254925v1

GCA_002549335.1_ASM254933v1

GCA_002549375.1_ASM254937v1

GCA_002549405.1_ASM254940v1

GCA_002549445.1_ASM254944v1

GCA_002549465.1_ASM254946v1

GCA_002549515.1_ASM254951v1

GCA_002549535.1_ASM254953v1

GCA_002593785.1_ASM259378v1

GCA_002593805.1_ASM259380v1

GCA_002593825.1_ASM259382v1

GCA_002593845.1_ASM259384v1

GCA_002593865.1_ASM259386v1

GCA_002593885.1_ASM259388v1

GCA_002593905.1_ASM259390v1

GCA_002593925.1_ASM259392v1

GCA_002593945.1_ASM259394v1

GCA_002593965.1_ASM259396v1

GCA_002593985.1_ASM259398v1

GCA_002594005.1_ASM259400v1

GCA_002594025.1_ASM259402v1

GCA_002594045.1_ASM259404v1

GCA_002594065.1_ASM259406v1

GCA_002594085.1_ASM259408v1

GCA_002594105.1_ASM259410v1

GCA_002594125.1_ASM259412v1

GCA_002594145.1_ASM259414v1

GCA_002594165.1_ASM259416v1

GCA_002594185.1_ASM259418v1

GCA_002594205.1_ASM259420v1

GCA_002594225.1_ASM259422v1

GCA_002594245.1_ASM259424v1

GCA_002594265.1_ASM259426v1

GCA_002594285.1_ASM259428v1

GCA_002594305.1_ASM259430v1

GCA_002594325.1_ASM259432v1

GCA_002594345.1_ASM259434v1

GCA_002594365.1_ASM259436v1

GCA_002594385.1_ASM259438v1

GCA_002594405.1_ASM259440v1

GCA_002594425.1_ASM259442v1

GCA_002594445.1_ASM259444v1

GCA_002594465.1_ASM259446v1

GCA_002594485.1_ASM259448v1

GCA_002594505.1_ASM259450v1

GCA_002594525.1_ASM259452v1

GCA_002594545.1_ASM259454v1

GCA_002594565.1_ASM259456v1

GCA_002595605.1_ASM259560v1

GCA_002595625.1_ASM259562v1

GCA_002595645.1_ASM259564v1

GCA_002595665.1_ASM259566v1

GCA_002595685.1_ASM259568v1

GCA_002595705.1_ASM259570v1

GCA_002595725.1_ASM259572v1

GCA_002595745.1_ASM259574v1

GCA_002595765.1_ASM259576v1

GCA_002595785.1_ASM259578v1

GCA_002595805.1_ASM259580v1

GCA_002595825.1_ASM259582v1

GCA_002595845.1_ASM259584v1

GCA_002595865.1_ASM259586v1

GCA_002595885.1_ASM259588v1

GCA_002595905.1_ASM259590v1

GCA_002595925.1_ASM259592v1

GCA_002595945.1_ASM259594v1

GCA_002595965.1_ASM259596v1

GCA_002595985.1_ASM259598v1

GCA_002596005.1_ASM259600v1

GCA_002596025.1_ASM259602v1

GCA_002596045.1_ASM259604v1

GCA_002596865.1_ASM259686v1

GCA_002596885.1_ASM259688v1

GCA_002596905.1_ASM259690v1

GCA_002596925.1_ASM259692v1

GCA_002596945.1_ASM259694v1

GCA_002596965.1_ASM259696v1

GCA_002596985.1_ASM259698v1

GCA_002597005.1_ASM259700v1

GCA_002597025.1_ASM259702v1

GCA_002597045.1_ASM259704v1

GCA_002597065.1_ASM259706v1

GCA_002597085.1_ASM259708v1

GCA_002597105.1_ASM259710v1

GCA_002597125.1_ASM259712v1

GCA_002597145.1_ASM259714v1

GCA_002597165.1_ASM259716v1

GCA_002597185.1_ASM259718v1

GCA_002598245.1_ASM259824v1

GCA_002598265.1_ASM259826v1

GCA_002598285.1_ASM259828v1

GCA_002598305.1_ASM259830v1

GCA_002598325.1_ASM259832v1

GCA_002598345.1_ASM259834v1

GCA_002598365.1_ASM259836v1

GCA_002598385.1_ASM259838v1

GCA_002603765.1_ASM260376v1

GCA_002604485.1_ASM260448v1

GCA_002604645.1_ASM260464v1

GCA_002612085.1_ASM261208v1

GCA_002619805.1_ASM261980v1

GCA_002623685.1_ASM262368v1

GCA_002627525.1_ASM262752v1

GCA_002630745.1_ASM263074v1

GCA_002723435.1_ASM272343v1

GCA_002745335.1_ASM274533v1

GCA_002755835.1_ASM275583v1

GCA_002921745.1_ASM292174v1

GCA_002921755.1_ASM292175v1

GCA_002921765.1_ASM292176v1

GCA_002921775.1_ASM292177v1

GCA_002921785.1_ASM292178v1

GCA_002921795.1_ASM292179v1

GCA_002922075.1_ASM292207v1

GCA_002922085.1_ASM292208v1

GCA_002922095.1_ASM292209v1

GCA_002922105.1_ASM292210v1

GCA_002922115.1_ASM292211v1

GCA_002922125.1_ASM292212v1

GCA_002922135.1_ASM292213v1

GCA_002922145.1_ASM292214v1

GCA_002922155.1_ASM292215v1

GCA_002922165.1_ASM292216v1

GCA_002922175.1_ASM292217v1

GCA_002922185.1_ASM292218v1

GCA_002922195.1_ASM292219v1

GCA_002955335.1_ASM295533v1

GCA_003093875.1_ASM309387v1

GCA_003147225.1_ASM314722v1

GCA_003369305.1_ASM336930v1

GCA_003428315.1_ASM342831v1

GCA_003443335.1_ASM344333v1

GCA_003723215.1_ASM372321v1

GCA_003723235.1_ASM372323v1

GCA_003723255.1_ASM372325v1

GCA_003723455.1_ASM372345v1

GCA_003861695.1_ASM386169v1

GCA_003862135.1_ASM386213v1

GCA_004521515.1_ASM452151v1

GCA_004521535.1_ASM452153v1

GCA_004521555.1_ASM452155v1

GCA_004800935.1_ASM480093v1

GCA_004875765.2_ASM487576v1

GCA_005394265.1_ASM539426v1

GCA_005394285.1_ASM539428v1

GCA_005394325.1_ASM539432v1

GCA_005394345.1_ASM539434v1

GCA_006083695.1_ASM608369v1

GCA_006869725.1_ASM686972v1

GCA_009745195.1_ASM974519v1

GCA_009745215.1_ASM974521v1

GCA_011757285.2_ASM1175728v2

GCA_011757295.2_ASM1175729v2

GCA_011757305.1_ASM1175730v1

GCA_013086035.1_ASM1308603v1

GCA_013187275.1_ASM1318727v1

GCA_013426285.1_ASM1342628v1

GCA_014190395.2_ASM1419039v1

GCA_015502395.1_ASM1550239v1

GCA_015502405.1_ASM1550240v1

GCA_015502415.1_ASM1550241v1

GCA_017654335.1_ASM1765433v1

GCA_017654345.1_ASM1765434v1

GCA_020473815.1_ASM2047381v1

GCA_020490015.1_ASM2049001v1

GCA_022694625.1_ASM2269462v1

GCA_024426065.1_ASM2442606v1

GCA_025630435.1_ASM2563043v1

GCA_025727455.1_ASM2572745v1

GCA_026568195.1_ASM2656819v1

GCA_027574075.1_ASM2757407v1

GCA_028974265.1_ASM2897426v1

GCA_028974415.1_ASM2897441v1

GCA_029875735.1_ASM2987573v1

GCA_029948225.1_ASM2994822v1

GCA_029948235.1_ASM2994823v1

GCA_029948265.1_ASM2994826v1

GCA_029948275.1_ASM2994827v1

GCA_029948305.1_ASM2994830v1

GCA_030463335.1_ASM3046333v1

GCA_030463365.1_ASM3046336v1

GCA_030463415.1_ASM3046341v1

GCA_030875095.1_ASM3087509v1

GCA_030875105.1_ASM3087510v1

GCA_030875115.1_ASM3087511v1

GCA_030875125.1_ASM3087512v1

GCA_030875135.1_ASM3087513v1

GCA_032255685.1_ASM3225568v1

GCA_032255735.1_ASM3225573v1

GCA_032255765.1_ASM3225576v1

GCA_032255945.1_ASM3225594v1

GCA_036584255.1_ASM3658425v1

GCA_036584265.1_ASM3658426v1

GCA_036584275.1_ASM3658427v1

GCA_036584285.1_ASM3658428v1

GCA_039568115.1_ASM3956811v1

GCA_039568125.1_ASM3956812v1

GCA_039568135.1_ASM3956813v1

GCA_039568145.1_ASM3956814v1

GCA_039568165.1_ASM3956816v1

Acanthamoeba reference genomes:

CM000150.2 Dictyostelium discoideum AX4 chromosome 1, whole genome shotgun sequence

CM000151.3 Dictyostelium discoideum AX4 chromosome 2, whole genome shotgun sequence

CM000152.2 Dictyostelium discoideum AX4 chromosome 3, whole genome shotgun sequence

CM000153.2 Dictyostelium discoideum AX4 chromosome 4, whole genome shotgun sequence

CM000154.2 Dictyostelium discoideum AX4 chromosome 5, whole genome shotgun sequence

CM000155.2 Dictyostelium discoideum AX4 chromosome 6, whole genome shotgun sequence

CH709156.1 Dictyostelium discoideum AX4 chrUn_00010 genomic scaffold, whole genome shotgun sequence

CH709157.1 Dictyostelium discoideum AX4 chrUn_00011 genomic scaffold, whole genome shotgun sequence

CH709158.1 Dictyostelium discoideum AX4 chrUn_00012 genomic scaffold, whole genome shotgun sequence

CH709159.1 Dictyostelium discoideum AX4 chrUn_00013 genomic scaffold, whole genome shotgun sequence

CH709160.1 Dictyostelium discoideum AX4 chrUn_00014 genomic scaffold, whole genome shotgun sequence

CH709161.1 Dictyostelium discoideum AX4 chrUn_00015 genomic scaffold, whole genome shotgun sequence

CH709162.1 Dictyostelium discoideum AX4 chrUn_00016 genomic scaffold, whole genome shotgun sequence

CH709164.1 Dictyostelium discoideum AX4 chrUn_00018 genomic scaffold, whole genome shotgun sequence

CH709148.1 Dictyostelium discoideum AX4 chrUn_0002 genomic scaffold, whole genome shotgun sequence

CH709166.1 Dictyostelium discoideum AX4 chrUn_00020 genomic scaffold, whole genome shotgun sequence

CH709167.1 Dictyostelium discoideum AX4 chrUn_00021 genomic scaffold, whole genome shotgun sequence

CH709168.1 Dictyostelium discoideum AX4 chrUn_00022 genomic scaffold, whole genome shotgun sequence

CH709169.1 Dictyostelium discoideum AX4 chrUn_00023 genomic scaffold, whole genome shotgun sequence

CH709170.1 Dictyostelium discoideum AX4 chrUn_00024 genomic scaffold, whole genome shotgun sequence

CH709171.1 Dictyostelium discoideum AX4 chrUn_00025 genomic scaffold, whole genome shotgun sequence

CH709172.1 Dictyostelium discoideum AX4 chrUn_00026 genomic scaffold, whole genome shotgun sequence

CH709173.1 Dictyostelium discoideum AX4 chrUn_00027 genomic scaffold, whole genome shotgun sequence

CH709174.1 Dictyostelium discoideum AX4 chrUn_00028 genomic scaffold, whole genome shotgun sequence

CH709175.1 Dictyostelium discoideum AX4 chrUn_00029 genomic scaffold, whole genome shotgun sequence

CH709149.1 Dictyostelium discoideum AX4 chrUn_0003 genomic scaffold, whole genome shotgun sequence

CH709176.1 Dictyostelium discoideum AX4 chrUn_00030 genomic scaffold, whole genome shotgun sequence

CH709177.1 Dictyostelium discoideum AX4 chrUn_00031 genomic scaffold, whole genome shotgun sequence

CH709178.1 Dictyostelium discoideum AX4 chrUn_00032 genomic scaffold, whole genome shotgun sequence

CH709179.1 Dictyostelium discoideum AX4 chrUn_00033 genomic scaffold, whole genome shotgun sequence

CH709180.1 Dictyostelium discoideum AX4 chrUn_00034 genomic scaffold, whole genome shotgun sequence

CH709181.1 Dictyostelium discoideum AX4 chrUn_00035 genomic scaffold, whole genome shotgun sequence

CH709182.1 Dictyostelium discoideum AX4 chrUn_00036 genomic scaffold, whole genome shotgun sequence

CH709150.1 Dictyostelium discoideum AX4 chrUn_0004 genomic scaffold, whole genome shotgun sequence

CH709151.1 Dictyostelium discoideum AX4 chrUn_0005 genomic scaffold, whole genome shotgun sequence

CH709152.1 Dictyostelium discoideum AX4 chrUn_0006 genomic scaffold, whole genome shotgun sequence

CH709153.1 Dictyostelium discoideum AX4 chrUn_0007 genomic scaffold, whole genome shotgun sequence

CH709154.1 Dictyostelium discoideum AX4 chrUn_0008 genomic scaffold, whole genome shotgun sequence

CH709155.1 Dictyostelium discoideum AX4 chrUn_0009 genomic scaffold, whole genome shotgun sequence

JH723757.1 Dictyostelium firmibasis strain TNS-C-0014 unplaced genomic scaffold scaffold00001, whole genome shotgun sequence

JH723758.1 Dictyostelium firmibasis strain TNS-C-0014 unplaced genomic scaffold scaffold00002, whole genome shotgun sequence

JH723759.1 Dictyostelium firmibasis strain TNS-C-0014 unplaced genomic scaffold scaffold00003, whole genome shotgun sequence

JH723760.1 Dictyostelium firmibasis strain TNS-C-0014 unplaced genomic scaffold scaffold00004, whole genome shotgun sequence

JH723761.1 Dictyostelium firmibasis strain TNS-C-0014 unplaced genomic scaffold scaffold00005, whole genome shotgun sequence

JH723762.1 Dictyostelium firmibasis strain TNS-C-0014 unplaced genomic scaffold scaffold00006, whole genome shotgun sequence

JH723763.1 Dictyostelium firmibasis strain TNS-C-0014 unplaced genomic scaffold scaffold00007, whole genome shotgun sequence

AJWH01000865.1 Dictyostelium firmibasis strain TNS-C-0014 Contig2007, whole genome shotgun sequence

JH723764.1 Dictyostelium firmibasis strain TNS-C-0014 unplaced genomic scaffold scaffold00009, whole genome shotgun sequence

JH723765.1 Dictyostelium firmibasis strain TNS-C-0014 unplaced genomic scaffold scaffold00010, whole genome shotgun sequence

JH723766.1 Dictyostelium firmibasis strain TNS-C-0014 unplaced genomic scaffold scaffold00011, whole genome shotgun sequence

JH723767.1 Dictyostelium firmibasis strain TNS-C-0014 unplaced genomic scaffold scaffold00012, whole genome shotgun sequence

JH723768.1 Dictyostelium firmibasis strain TNS-C-0014 unplaced genomic scaffold scaffold00013, whole genome shotgun sequence

JH723769.1 Dictyostelium firmibasis strain TNS-C-0014 unplaced genomic scaffold scaffold00014, whole genome shotgun sequence

JH723770.1 Dictyostelium firmibasis strain TNS-C-0014 unplaced genomic scaffold scaffold00015, whole genome shotgun sequence

JH723771.1 Dictyostelium firmibasis strain TNS-C-0014 unplaced genomic scaffold scaffold00016, whole genome shotgun sequence

JH723772.1 Dictyostelium firmibasis strain TNS-C-0014 unplaced genomic scaffold scaffold00017, whole genome shotgun sequence

JH723773.1 Dictyostelium firmibasis strain TNS-C-0014 unplaced genomic scaffold scaffold00018, whole genome shotgun sequence

AJWH01001718.1 Dictyostelium firmibasis strain TNS-C-0014 Contig3430, whole genome shotgun sequence

JH723774.1 Dictyostelium firmibasis strain TNS-C-0014 unplaced genomic scaffold scaffold00020, whole genome shotgun sequence

JH723775.1 Dictyostelium firmibasis strain TNS-C-0014 unplaced genomic scaffold scaffold00021, whole genome shotgun sequence

JH723776.1 Dictyostelium firmibasis strain TNS-C-0014 unplaced genomic scaffold scaffold00022, whole genome shotgun sequence

JH723777.1 Dictyostelium firmibasis strain TNS-C-0014 unplaced genomic scaffold scaffold00023, whole genome shotgun sequence

JH723778.1 Dictyostelium firmibasis strain TNS-C-0014 unplaced genomic scaffold scaffold00024, whole genome shotgun sequence

JH723779.1 Dictyostelium firmibasis strain TNS-C-0014 unplaced genomic scaffold scaffold00025, whole genome shotgun sequence

JH723780.1 Dictyostelium firmibasis strain TNS-C-0014 unplaced genomic scaffold scaffold00026, whole genome shotgun sequence

JH723781.1 Dictyostelium firmibasis strain TNS-C-0014 unplaced genomic scaffold scaffold00027, whole genome shotgun sequence

JH723782.1 Dictyostelium firmibasis strain TNS-C-0014 unplaced genomic scaffold scaffold00028, whole genome shotgun sequence

AJWH01002397.1 Dictyostelium firmibasis strain TNS-C-0014 Contig5179, whole genome shotgun sequence

JH723783.1 Dictyostelium firmibasis strain TNS-C-0014 unplaced genomic scaffold scaffold00030, whole genome shotgun sequence

JH723784.1 Dictyostelium firmibasis strain TNS-C-0014 unplaced genomic scaffold scaffold00031, whole genome shotgun sequence

JH723785.1 Dictyostelium firmibasis strain TNS-C-0014 unplaced genomic scaffold scaffold00032, whole genome shotgun sequence

JH723786.1 Dictyostelium firmibasis strain TNS-C-0014 unplaced genomic scaffold scaffold00033, whole genome shotgun sequence

JH723787.1 Dictyostelium firmibasis strain TNS-C-0014 unplaced genomic scaffold scaffold00034, whole genome shotgun sequence

JH723788.1 Dictyostelium firmibasis strain TNS-C-0014 unplaced genomic scaffold scaffold00035, whole genome shotgun sequence

JH723789.1 Dictyostelium firmibasis strain TNS-C-0014 unplaced genomic scaffold scaffold00036, whole genome shotgun sequence

AJWH01002792.1 Dictyostelium firmibasis strain TNS-C-0014 Contig5944, whole genome shotgun sequence

JH723790.1 Dictyostelium firmibasis strain TNS-C-0014 unplaced genomic scaffold scaffold00038, whole genome shotgun sequence

JH723791.1 Dictyostelium firmibasis strain TNS-C-0014 unplaced genomic scaffold scaffold00039, whole genome shotgun sequence

AJWH01002900.1 Dictyostelium firmibasis strain TNS-C-0014 Contig6161, whole genome shotgun sequence

JH723792.1 Dictyostelium firmibasis strain TNS-C-0014 unplaced genomic scaffold scaffold00041, whole genome shotgun sequence

AJWH01002948.1 Dictyostelium firmibasis strain TNS-C-0014 Contig6230, whole genome shotgun sequence

AJWH01002949.1 Dictyostelium firmibasis strain TNS-C-0014 Contig6338, whole genome shotgun sequence

JH723793.1 Dictyostelium firmibasis strain TNS-C-0014 unplaced genomic scaffold scaffold00044, whole genome shotgun sequence

JH723794.1 Dictyostelium firmibasis strain TNS-C-0014 unplaced genomic scaffold scaffold00045, whole genome shotgun sequence

JH723795.1 Dictyostelium firmibasis strain TNS-C-0014 unplaced genomic scaffold scaffold00046, whole genome shotgun sequence

JH723796.1 Dictyostelium firmibasis strain TNS-C-0014 unplaced genomic scaffold scaffold00047, whole genome shotgun sequence

JH723797.1 Dictyostelium firmibasis strain TNS-C-0014 unplaced genomic scaffold scaffold00048, whole genome shotgun sequence

JH723798.1 Dictyostelium firmibasis strain TNS-C-0014 unplaced genomic scaffold scaffold00049, whole genome shotgun sequence

AJWH01003260.1 Dictyostelium firmibasis strain TNS-C-0014 Contig6845, whole genome shotgun sequence

JH723799.1 Dictyostelium firmibasis strain TNS-C-0014 unplaced genomic scaffold scaffold00051, whole genome shotgun sequence

JH723800.1 Dictyostelium firmibasis strain TNS-C-0014 unplaced genomic scaffold scaffold00052, whole genome shotgun sequence

JH723801.1 Dictyostelium firmibasis strain TNS-C-0014 unplaced genomic scaffold scaffold00053, whole genome shotgun sequence

JH723802.1 Dictyostelium firmibasis strain TNS-C-0014 unplaced genomic scaffold scaffold00054, whole genome shotgun sequence

JH723803.1 Dictyostelium firmibasis strain TNS-C-0014 unplaced genomic scaffold scaffold00055, whole genome shotgun sequence

AJWH01003518.1 Dictyostelium firmibasis strain TNS-C-0014 Contig7425, whole genome shotgun sequence

JH723804.1 Dictyostelium firmibasis strain TNS-C-0014 unplaced genomic scaffold scaffold00057, whole genome shotgun sequence

JH723805.1 Dictyostelium firmibasis strain TNS-C-0014 unplaced genomic scaffold scaffold00058, whole genome shotgun sequence

JH723806.1 Dictyostelium firmibasis strain TNS-C-0014 unplaced genomic scaffold scaffold00059, whole genome shotgun sequence

AJWH01003649.1 Dictyostelium firmibasis strain TNS-C-0014 Contig7593, whole genome shotgun sequence

JH723807.1 Dictyostelium firmibasis strain TNS-C-0014 unplaced genomic scaffold scaffold00061, whole genome shotgun sequence

JH723808.1 Dictyostelium firmibasis strain TNS-C-0014 unplaced genomic scaffold scaffold00062, whole genome shotgun sequence

JH723809.1 Dictyostelium firmibasis strain TNS-C-0014 unplaced genomic scaffold scaffold00063, whole genome shotgun sequence

JH723810.1 Dictyostelium firmibasis strain TNS-C-0014 unplaced genomic scaffold scaffold00064, whole genome shotgun sequence

AJWH01003821.1 Dictyostelium firmibasis strain TNS-C-0014 Contig7894, whole genome shotgun sequence

JH723811.1 Dictyostelium firmibasis strain TNS-C-0014 unplaced genomic scaffold scaffold00066, whole genome shotgun sequence

JH723812.1 Dictyostelium firmibasis strain TNS-C-0014 unplaced genomic scaffold scaffold00067, whole genome shotgun sequence

JH723813.1 Dictyostelium firmibasis strain TNS-C-0014 unplaced genomic scaffold scaffold00068, whole genome shotgun sequence

JH723814.1 Dictyostelium firmibasis strain TNS-C-0014 unplaced genomic scaffold scaffold00069, whole genome shotgun sequence

JH723815.1 Dictyostelium firmibasis strain TNS-C-0014 unplaced genomic scaffold scaffold00070, whole genome shotgun sequence

JH723816.1 Dictyostelium firmibasis strain TNS-C-0014 unplaced genomic scaffold scaffold00071, whole genome shotgun sequence

AJWH01003991.1 Dictyostelium firmibasis strain TNS-C-0014 Contig8265, whole genome shotgun sequence

JH723817.1 Dictyostelium firmibasis strain TNS-C-0014 unplaced genomic scaffold scaffold00073, whole genome shotgun sequence

JH723818.1 Dictyostelium firmibasis strain TNS-C-0014 unplaced genomic scaffold scaffold00074, whole genome shotgun sequence

JH723819.1 Dictyostelium firmibasis strain TNS-C-0014 unplaced genomic scaffold scaffold00075, whole genome shotgun sequence

AJWH01004091.1 Dictyostelium firmibasis strain TNS-C-0014 Contig8399, whole genome shotgun sequence

JH723820.1 Dictyostelium firmibasis strain TNS-C-0014 unplaced genomic scaffold scaffold00077, whole genome shotgun sequence

JH723821.1 Dictyostelium firmibasis strain TNS-C-0014 unplaced genomic scaffold scaffold00078, whole genome shotgun sequence

AJWH01004158.1 Dictyostelium firmibasis strain TNS-C-0014 Contig8528, whole genome shotgun sequence

JH723822.1 Dictyostelium firmibasis strain TNS-C-0014 unplaced genomic scaffold scaffold00080, whole genome shotgun sequence

AJWH01004196.1 Dictyostelium firmibasis strain TNS-C-0014 Contig8682, whole genome shotgun sequence

JH723823.1 Dictyostelium firmibasis strain TNS-C-0014 unplaced genomic scaffold scaffold00082, whole genome shotgun sequence

JH723824.1 Dictyostelium firmibasis strain TNS-C-0014 unplaced genomic scaffold scaffold00083, whole genome shotgun sequence

JH723825.1 Dictyostelium firmibasis strain TNS-C-0014 unplaced genomic scaffold scaffold00084, whole genome shotgun sequence

JH723826.1 Dictyostelium firmibasis strain TNS-C-0014 unplaced genomic scaffold scaffold00085, whole genome shotgun sequence

JH723827.1 Dictyostelium firmibasis strain TNS-C-0014 unplaced genomic scaffold scaffold00086, whole genome shotgun sequence

AJWH01004349.1 Dictyostelium firmibasis strain TNS-C-0014 Contig9194, whole genome shotgun sequence

JH723828.1 Dictyostelium firmibasis strain TNS-C-0014 unplaced genomic scaffold scaffold00088, whole genome shotgun sequence

JH723829.1 Dictyostelium firmibasis strain TNS-C-0014 unplaced genomic scaffold scaffold00089, whole genome shotgun sequence

JH723830.1 Dictyostelium firmibasis strain TNS-C-0014 unplaced genomic scaffold scaffold00090, whole genome shotgun sequence

JH723831.1 Dictyostelium firmibasis strain TNS-C-0014 unplaced genomic scaffold scaffold00091, whole genome shotgun sequence

JH723832.1 Dictyostelium firmibasis strain TNS-C-0014 unplaced genomic scaffold scaffold00092, whole genome shotgun sequence

JH723833.1 Dictyostelium firmibasis strain TNS-C-0014 unplaced genomic scaffold scaffold00093, whole genome shotgun sequence

AJWH01004558.1 Dictyostelium firmibasis strain TNS-C-0014 Contig9764, whole genome shotgun sequence

JH723834.1 Dictyostelium firmibasis strain TNS-C-0014 unplaced genomic scaffold scaffold00095, whole genome shotgun sequence

JH723835.1 Dictyostelium firmibasis strain TNS-C-0014 unplaced genomic scaffold scaffold00096, whole genome shotgun sequence

JH723836.1 Dictyostelium firmibasis strain TNS-C-0014 unplaced genomic scaffold scaffold00097, whole genome shotgun sequence

JH723837.1 Dictyostelium firmibasis strain TNS-C-0014 unplaced genomic scaffold scaffold00098, whole genome shotgun sequence

JH723838.1 Dictyostelium firmibasis strain TNS-C-0014 unplaced genomic scaffold scaffold00099, whole genome shotgun sequence

JH723839.1 Dictyostelium firmibasis strain TNS-C-0014 unplaced genomic scaffold scaffold00100, whole genome shotgun sequence

JH723840.1 Dictyostelium firmibasis strain TNS-C-0014 unplaced genomic scaffold scaffold00101, whole genome shotgun sequence

AJWH01004744.1 Dictyostelium firmibasis strain TNS-C-0014 Contig10123, whole genome shotgun sequence

JH723841.1 Dictyostelium firmibasis strain TNS-C-0014 unplaced genomic scaffold scaffold00103, whole genome shotgun sequence

JH723842.1 Dictyostelium firmibasis strain TNS-C-0014 unplaced genomic scaffold scaffold00104, whole genome shotgun sequence

JH723843.1 Dictyostelium firmibasis strain TNS-C-0014 unplaced genomic scaffold scaffold00105, whole genome shotgun sequence

AJWH01004814.1 Dictyostelium firmibasis strain TNS-C-0014 Contig10305, whole genome shotgun sequence

JH723844.1 Dictyostelium firmibasis strain TNS-C-0014 unplaced genomic scaffold scaffold00107, whole genome shotgun sequence

JH723845.1 Dictyostelium firmibasis strain TNS-C-0014 unplaced genomic scaffold scaffold00108, whole genome shotgun sequence

JH723846.1 Dictyostelium firmibasis strain TNS-C-0014 unplaced genomic scaffold scaffold00109, whole genome shotgun sequence

AJWH01004899.1 Dictyostelium firmibasis strain TNS-C-0014 Contig10450, whole genome shotgun sequence

AJWH01004900.1 Dictyostelium firmibasis strain TNS-C-0014 Contig10464, whole genome shotgun sequence

JH723847.1 Dictyostelium firmibasis strain TNS-C-0014 unplaced genomic scaffold scaffold00112, whole genome shotgun sequence

JH723848.1 Dictyostelium firmibasis strain TNS-C-0014 unplaced genomic scaffold scaffold00113, whole genome shotgun sequence

JH723849.1 Dictyostelium firmibasis strain TNS-C-0014 unplaced genomic scaffold scaffold00114, whole genome shotgun sequence

JH723850.1 Dictyostelium firmibasis strain TNS-C-0014 unplaced genomic scaffold scaffold00115, whole genome shotgun sequence

JH723851.1 Dictyostelium firmibasis strain TNS-C-0014 unplaced genomic scaffold scaffold00116, whole genome shotgun sequence

JH723852.1 Dictyostelium firmibasis strain TNS-C-0014 unplaced genomic scaffold scaffold00117, whole genome shotgun sequence

JH723853.1 Dictyostelium firmibasis strain TNS-C-0014 unplaced genomic scaffold scaffold00118, whole genome shotgun sequence

AJWH01005024.1 Dictyostelium firmibasis strain TNS-C-0014 Contig10741, whole genome shotgun sequence

JH723854.1 Dictyostelium firmibasis strain TNS-C-0014 unplaced genomic scaffold scaffold00120, whole genome shotgun sequence

JH723855.1 Dictyostelium firmibasis strain TNS-C-0014 unplaced genomic scaffold scaffold00121, whole genome shotgun sequence

JH723856.1 Dictyostelium firmibasis strain TNS-C-0014 unplaced genomic scaffold scaffold00122, whole genome shotgun sequence

AJWH01005081.1 Dictyostelium firmibasis strain TNS-C-0014 Contig11013, whole genome shotgun sequence

AJWH01005084.1 Dictyostelium firmibasis strain TNS-C-0014 Contig11078, whole genome shotgun sequence

JH723857.1 Dictyostelium firmibasis strain TNS-C-0014 unplaced genomic scaffold scaffold00125, whole genome shotgun sequence

JH723858.1 Dictyostelium firmibasis strain TNS-C-0014 unplaced genomic scaffold scaffold00126, whole genome shotgun sequence

JH723859.1 Dictyostelium firmibasis strain TNS-C-0014 unplaced genomic scaffold scaffold00127, whole genome shotgun sequence

JH723860.1 Dictyostelium firmibasis strain TNS-C-0014 unplaced genomic scaffold scaffold00128, whole genome shotgun sequence

JH723861.1 Dictyostelium firmibasis strain TNS-C-0014 unplaced genomic scaffold scaffold00129, whole genome shotgun sequence

JH723862.1 Dictyostelium firmibasis strain TNS-C-0014 unplaced genomic scaffold scaffold00130, whole genome shotgun sequence

JH723863.1 Dictyostelium firmibasis strain TNS-C-0014 unplaced genomic scaffold scaffold00131, whole genome shotgun sequence

JH723864.1 Dictyostelium firmibasis strain TNS-C-0014 unplaced genomic scaffold scaffold00132, whole genome shotgun sequence

JH723865.1 Dictyostelium firmibasis strain TNS-C-0014 unplaced genomic scaffold scaffold00133, whole genome shotgun sequence

JH723866.1 Dictyostelium firmibasis strain TNS-C-0014 unplaced genomic scaffold scaffold00134, whole genome shotgun sequence

JH723867.1 Dictyostelium firmibasis strain TNS-C-0014 unplaced genomic scaffold scaffold00135, whole genome shotgun sequence

JH723868.1 Dictyostelium firmibasis strain TNS-C-0014 unplaced genomic scaffold scaffold00136, whole genome shotgun sequence

JH723869.1 Dictyostelium firmibasis strain TNS-C-0014 unplaced genomic scaffold scaffold00137, whole genome shotgun sequence

AJWH01005405.1 Dictyostelium firmibasis strain TNS-C-0014 Contig11569, whole genome shotgun sequence

JH723870.1 Dictyostelium firmibasis strain TNS-C-0014 unplaced genomic scaffold scaffold00139, whole genome shotgun sequence

AJWH01005428.1 Dictyostelium firmibasis strain TNS-C-0014 Contig11661, whole genome shotgun sequence

AJWH01005429.1 Dictyostelium firmibasis strain TNS-C-0014 Contig11688, whole genome shotgun sequence

JH723871.1 Dictyostelium firmibasis strain TNS-C-0014 unplaced genomic scaffold scaffold00142, whole genome shotgun sequence

JH723872.1 Dictyostelium firmibasis strain TNS-C-0014 unplaced genomic scaffold scaffold00143, whole genome shotgun sequence

JH723873.1 Dictyostelium firmibasis strain TNS-C-0014 unplaced genomic scaffold scaffold00144, whole genome shotgun sequence

AJWH01005496.1 Dictyostelium firmibasis strain TNS-C-0014 Contig11934, whole genome shotgun sequence

JH723874.1 Dictyostelium firmibasis strain TNS-C-0014 unplaced genomic scaffold scaffold00146, whole genome shotgun sequence

JH723875.1 Dictyostelium firmibasis strain TNS-C-0014 unplaced genomic scaffold scaffold00147, whole genome shotgun sequence

JH723876.1 Dictyostelium firmibasis strain TNS-C-0014 unplaced genomic scaffold scaffold00148, whole genome shotgun sequence

JH723877.1 Dictyostelium firmibasis strain TNS-C-0014 unplaced genomic scaffold scaffold00149, whole genome shotgun sequence

AJWH01005562.1 Dictyostelium firmibasis strain TNS-C-0014 Contig12107, whole genome shotgun sequence

JH723878.1 Dictyostelium firmibasis strain TNS-C-0014 unplaced genomic scaffold scaffold00151, whole genome shotgun sequence

JH723879.1 Dictyostelium firmibasis strain TNS-C-0014 unplaced genomic scaffold scaffold00152, whole genome shotgun sequence

JH723880.1 Dictyostelium firmibasis strain TNS-C-0014 unplaced genomic scaffold scaffold00153, whole genome shotgun sequence

AJWH01005621.1 Dictyostelium firmibasis strain TNS-C-0014 Contig12260, whole genome shotgun sequence

JH723881.1 Dictyostelium firmibasis strain TNS-C-0014 unplaced genomic scaffold scaffold00155, whole genome shotgun sequence

JH723882.1 Dictyostelium firmibasis strain TNS-C-0014 unplaced genomic scaffold scaffold00156, whole genome shotgun sequence

JH723883.1 Dictyostelium firmibasis strain TNS-C-0014 unplaced genomic scaffold scaffold00157, whole genome shotgun sequence

JH723884.1 Dictyostelium firmibasis strain TNS-C-0014 unplaced genomic scaffold scaffold00158, whole genome shotgun sequence

AJWH01005709.1 Dictyostelium firmibasis strain TNS-C-0014 Contig12440, whole genome shotgun sequence

JH723885.1 Dictyostelium firmibasis strain TNS-C-0014 unplaced genomic scaffold scaffold00160, whole genome shotgun sequence

AJWH01005728.1 Dictyostelium firmibasis strain TNS-C-0014 Contig12477, whole genome shotgun sequence

JH723886.1 Dictyostelium firmibasis strain TNS-C-0014 unplaced genomic scaffold scaffold00162, whole genome shotgun sequence

AJWH01005731.1 Dictyostelium firmibasis strain TNS-C-0014 Contig12551, whole genome shotgun sequence

JH723887.1 Dictyostelium firmibasis strain TNS-C-0014 unplaced genomic scaffold scaffold00164, whole genome shotgun sequence

JH723888.1 Dictyostelium firmibasis strain TNS-C-0014 unplaced genomic scaffold scaffold00165, whole genome shotgun sequence

AJWH01005767.1 Dictyostelium firmibasis strain TNS-C-0014 Contig12613, whole genome shotgun sequence

AJWH01005768.1 Dictyostelium firmibasis strain TNS-C-0014 Contig12657, whole genome shotgun sequence

AJWH01005769.1 Dictyostelium firmibasis strain TNS-C-0014 Contig12700, whole genome shotgun sequence

JH723889.1 Dictyostelium firmibasis strain TNS-C-0014 unplaced genomic scaffold scaffold00169, whole genome shotgun sequence

AJWH01005790.1 Dictyostelium firmibasis strain TNS-C-0014 Contig12737, whole genome shotgun sequence

JH723890.1 Dictyostelium firmibasis strain TNS-C-0014 unplaced genomic scaffold scaffold00171, whole genome shotgun sequence

AJWH01005809.1 Dictyostelium firmibasis strain TNS-C-0014 Contig12842, whole genome shotgun sequence

JH723891.1 Dictyostelium firmibasis strain TNS-C-0014 unplaced genomic scaffold scaffold00173, whole genome shotgun sequence

JH723892.1 Dictyostelium firmibasis strain TNS-C-0014 unplaced genomic scaffold scaffold00174, whole genome shotgun sequence

JH723893.1 Dictyostelium firmibasis strain TNS-C-0014 unplaced genomic scaffold scaffold00175, whole genome shotgun sequence

JH723894.1 Dictyostelium firmibasis strain TNS-C-0014 unplaced genomic scaffold scaffold00176, whole genome shotgun sequence

AJWH01005887.1 Dictyostelium firmibasis strain TNS-C-0014 Contig12950, whole genome shotgun sequence

JH723895.1 Dictyostelium firmibasis strain TNS-C-0014 unplaced genomic scaffold scaffold00178, whole genome shotgun sequence

AJWH01005906.1 Dictyostelium firmibasis strain TNS-C-0014 Contig13078, whole genome shotgun sequence

JH723896.1 Dictyostelium firmibasis strain TNS-C-0014 unplaced genomic scaffold scaffold00180, whole genome shotgun sequence

AJWH01005923.1 Dictyostelium firmibasis strain TNS-C-0014 Contig13116, whole genome shotgun sequence

JH723897.1 Dictyostelium firmibasis strain TNS-C-0014 unplaced genomic scaffold scaffold00182, whole genome shotgun sequence

JH723898.1 Dictyostelium firmibasis strain TNS-C-0014 unplaced genomic scaffold scaffold00183, whole genome shotgun sequence

JH723899.1 Dictyostelium firmibasis strain TNS-C-0014 unplaced genomic scaffold scaffold00184, whole genome shotgun sequence

AJWH01005951.1 Dictyostelium firmibasis strain TNS-C-0014 Contig13187, whole genome shotgun sequence

AJWH01005952.1 Dictyostelium firmibasis strain TNS-C-0014 Contig13220, whole genome shotgun sequence

AJWH01005953.1 Dictyostelium firmibasis strain TNS-C-0014 Contig13262, whole genome shotgun sequence

JH723900.1 Dictyostelium firmibasis strain TNS-C-0014 unplaced genomic scaffold scaffold00188, whole genome shotgun sequence

JH723901.1 Dictyostelium firmibasis strain TNS-C-0014 unplaced genomic scaffold scaffold00189, whole genome shotgun sequence

JH723902.1 Dictyostelium firmibasis strain TNS-C-0014 unplaced genomic scaffold scaffold00190, whole genome shotgun sequence

JH723903.1 Dictyostelium firmibasis strain TNS-C-0014 unplaced genomic scaffold scaffold00191, whole genome shotgun sequence

JH723904.1 Dictyostelium firmibasis strain TNS-C-0014 unplaced genomic scaffold scaffold00192, whole genome shotgun sequence

JH723905.1 Dictyostelium firmibasis strain TNS-C-0014 unplaced genomic scaffold scaffold00193, whole genome shotgun sequence

AJWH01006049.1 Dictyostelium firmibasis strain TNS-C-0014 Contig13466, whole genome shotgun sequence

JH723906.1 Dictyostelium firmibasis strain TNS-C-0014 unplaced genomic scaffold scaffold00195, whole genome shotgun sequence

AJWH01006068.1 Dictyostelium firmibasis strain TNS-C-0014 Contig13544, whole genome shotgun sequence

JH723907.1 Dictyostelium firmibasis strain TNS-C-0014 unplaced genomic scaffold scaffold00197, whole genome shotgun sequence

JH723908.1 Dictyostelium firmibasis strain TNS-C-0014 unplaced genomic scaffold scaffold00198, whole genome shotgun sequence

JH723909.1 Dictyostelium firmibasis strain TNS-C-0014 unplaced genomic scaffold scaffold00199, whole genome shotgun sequence

JH723910.1 Dictyostelium firmibasis strain TNS-C-0014 unplaced genomic scaffold scaffold00200, whole genome shotgun sequence

JH723911.1 Dictyostelium firmibasis strain TNS-C-0014 unplaced genomic scaffold scaffold00201, whole genome shotgun sequence

JH723912.1 Dictyostelium firmibasis strain TNS-C-0014 unplaced genomic scaffold scaffold00202, whole genome shotgun sequence

JH723913.1 Dictyostelium firmibasis strain TNS-C-0014 unplaced genomic scaffold scaffold00203, whole genome shotgun sequence

JH723914.1 Dictyostelium firmibasis strain TNS-C-0014 unplaced genomic scaffold scaffold00204, whole genome shotgun sequence

JH723915.1 Dictyostelium firmibasis strain TNS-C-0014 unplaced genomic scaffold scaffold00205, whole genome shotgun sequence

JH723916.1 Dictyostelium firmibasis strain TNS-C-0014 unplaced genomic scaffold scaffold00206, whole genome shotgun sequence

AJWH01006179.1 Dictyostelium firmibasis strain TNS-C-0014 Contig13729, whole genome shotgun sequence

JH723917.1 Dictyostelium firmibasis strain TNS-C-0014 unplaced genomic scaffold scaffold00208, whole genome shotgun sequence

JH723918.1 Dictyostelium firmibasis strain TNS-C-0014 unplaced genomic scaffold scaffold00209, whole genome shotgun sequence

JH723919.1 Dictyostelium firmibasis strain TNS-C-0014 unplaced genomic scaffold scaffold00210, whole genome shotgun sequence

JH723920.1 Dictyostelium firmibasis strain TNS-C-0014 unplaced genomic scaffold scaffold00211, whole genome shotgun sequence

JH723921.1 Dictyostelium firmibasis strain TNS-C-0014 unplaced genomic scaffold scaffold00212, whole genome shotgun sequence

JH723922.1 Dictyostelium firmibasis strain TNS-C-0014 unplaced genomic scaffold scaffold00213, whole genome shotgun sequence

JH723923.1 Dictyostelium firmibasis strain TNS-C-0014 unplaced genomic scaffold scaffold00214, whole genome shotgun sequence

JH723924.1 Dictyostelium firmibasis strain TNS-C-0014 unplaced genomic scaffold scaffold00215, whole genome shotgun sequence

JH723925.1 Dictyostelium firmibasis strain TNS-C-0014 unplaced genomic scaffold scaffold00216, whole genome shotgun sequence

JH723926.1 Dictyostelium firmibasis strain TNS-C-0014 unplaced genomic scaffold scaffold00217, whole genome shotgun sequence

AJWH01006299.1 Dictyostelium firmibasis strain TNS-C-0014 Contig13947, whole genome shotgun sequence

JH723927.1 Dictyostelium firmibasis strain TNS-C-0014 unplaced genomic scaffold scaffold00219, whole genome shotgun sequence

AJWH01006310.1 Dictyostelium firmibasis strain TNS-C-0014 Contig13986, whole genome shotgun sequence

JH723928.1 Dictyostelium firmibasis strain TNS-C-0014 unplaced genomic scaffold scaffold00221, whole genome shotgun sequence

JH723929.1 Dictyostelium firmibasis strain TNS-C-0014 unplaced genomic scaffold scaffold00222, whole genome shotgun sequence

JH723930.1 Dictyostelium firmibasis strain TNS-C-0014 unplaced genomic scaffold scaffold00223, whole genome shotgun sequence

JH723931.1 Dictyostelium firmibasis strain TNS-C-0014 unplaced genomic scaffold scaffold00224, whole genome shotgun sequence

JH723932.1 Dictyostelium firmibasis strain TNS-C-0014 unplaced genomic scaffold scaffold00225, whole genome shotgun sequence

JH723933.1 Dictyostelium firmibasis strain TNS-C-0014 unplaced genomic scaffold scaffold00226, whole genome shotgun sequence

JH723934.1 Dictyostelium firmibasis strain TNS-C-0014 unplaced genomic scaffold scaffold00227, whole genome shotgun sequence

JH723935.1 Dictyostelium firmibasis strain TNS-C-0014 unplaced genomic scaffold scaffold00228, whole genome shotgun sequence

AJWH01006383.1 Dictyostelium firmibasis strain TNS-C-0014 Contig14111, whole genome shotgun sequence

JH723936.1 Dictyostelium firmibasis strain TNS-C-0014 unplaced genomic scaffold scaffold00230, whole genome shotgun sequence

JH723937.1 Dictyostelium firmibasis strain TNS-C-0014 unplaced genomic scaffold scaffold00231, whole genome shotgun sequence

JH723938.1 Dictyostelium firmibasis strain TNS-C-0014 unplaced genomic scaffold scaffold00232, whole genome shotgun sequence

JH723939.1 Dictyostelium firmibasis strain TNS-C-0014 unplaced genomic scaffold scaffold00233, whole genome shotgun sequence

JH723940.1 Dictyostelium firmibasis strain TNS-C-0014 unplaced genomic scaffold scaffold00234, whole genome shotgun sequence

JH723941.1 Dictyostelium firmibasis strain TNS-C-0014 unplaced genomic scaffold scaffold00235, whole genome shotgun sequence

JH723942.1 Dictyostelium firmibasis strain TNS-C-0014 unplaced genomic scaffold scaffold00236, whole genome shotgun sequence

AJWH01006447.1 Dictyostelium firmibasis strain TNS-C-0014 Contig14293, whole genome shotgun sequence

AJWH01006448.1 Dictyostelium firmibasis strain TNS-C-0014 Contig14302, whole genome shotgun sequence

JH723943.1 Dictyostelium firmibasis strain TNS-C-0014 unplaced genomic scaffold scaffold00239, whole genome shotgun sequence

JH723944.1 Dictyostelium firmibasis strain TNS-C-0014 unplaced genomic scaffold scaffold00240, whole genome shotgun sequence

JH723945.1 Dictyostelium firmibasis strain TNS-C-0014 unplaced genomic scaffold scaffold00241, whole genome shotgun sequence

JH723946.1 Dictyostelium firmibasis strain TNS-C-0014 unplaced genomic scaffold scaffold00242, whole genome shotgun sequence

JH723947.1 Dictyostelium firmibasis strain TNS-C-0014 unplaced genomic scaffold scaffold00243, whole genome shotgun sequence

JH723948.1 Dictyostelium firmibasis strain TNS-C-0014 unplaced genomic scaffold scaffold00244, whole genome shotgun sequence

JH723949.1 Dictyostelium firmibasis strain TNS-C-0014 unplaced genomic scaffold scaffold00245, whole genome shotgun sequence

JH723950.1 Dictyostelium firmibasis strain TNS-C-0014 unplaced genomic scaffold scaffold00246, whole genome shotgun sequence

AJWH01006501.1 Dictyostelium firmibasis strain TNS-C-0014 Contig14430, whole genome shotgun sequence

JH723951.1 Dictyostelium firmibasis strain TNS-C-0014 unplaced genomic scaffold scaffold00248, whole genome shotgun sequence

AJWH01006512.1 Dictyostelium firmibasis strain TNS-C-0014 Contig14450, whole genome shotgun sequence

JH723952.1 Dictyostelium firmibasis strain TNS-C-0014 unplaced genomic scaffold scaffold00250, whole genome shotgun sequence

JH723953.1 Dictyostelium firmibasis strain TNS-C-0014 unplaced genomic scaffold scaffold00251, whole genome shotgun sequence

JH723954.1 Dictyostelium firmibasis strain TNS-C-0014 unplaced genomic scaffold scaffold00252, whole genome shotgun sequence

JH723955.1 Dictyostelium firmibasis strain TNS-C-0014 unplaced genomic scaffold scaffold00253, whole genome shotgun sequence

JH723956.1 Dictyostelium firmibasis strain TNS-C-0014 unplaced genomic scaffold scaffold00254, whole genome shotgun sequence

JH723957.1 Dictyostelium firmibasis strain TNS-C-0014 unplaced genomic scaffold scaffold00255, whole genome shotgun sequence

JH723958.1 Dictyostelium firmibasis strain TNS-C-0014 unplaced genomic scaffold scaffold00256, whole genome shotgun sequence

JH723959.1 Dictyostelium firmibasis strain TNS-C-0014 unplaced genomic scaffold scaffold00257, whole genome shotgun sequence

JH723960.1 Dictyostelium firmibasis strain TNS-C-0014 unplaced genomic scaffold scaffold00258, whole genome shotgun sequence

JH723961.1 Dictyostelium firmibasis strain TNS-C-0014 unplaced genomic scaffold scaffold00259, whole genome shotgun sequence

AJWH01006563.1 Dictyostelium firmibasis strain TNS-C-0014 Contig14570, whole genome shotgun sequence

JH723962.1 Dictyostelium firmibasis strain TNS-C-0014 unplaced genomic scaffold scaffold00261, whole genome shotgun sequence

JH723963.1 Dictyostelium firmibasis strain TNS-C-0014 unplaced genomic scaffold scaffold00262, whole genome shotgun sequence

JH723964.1 Dictyostelium firmibasis strain TNS-C-0014 unplaced genomic scaffold scaffold00263, whole genome shotgun sequence

AJWH01006579.1 Dictyostelium firmibasis strain TNS-C-0014 Contig14597, whole genome shotgun sequence

JH723965.1 Dictyostelium firmibasis strain TNS-C-0014 unplaced genomic scaffold scaffold00265, whole genome shotgun sequence

AJWH01006584.1 Dictyostelium firmibasis strain TNS-C-0014 Contig14603, whole genome shotgun sequence

JH723966.1 Dictyostelium firmibasis strain TNS-C-0014 unplaced genomic scaffold scaffold00267, whole genome shotgun sequence

JH723967.1 Dictyostelium firmibasis strain TNS-C-0014 unplaced genomic scaffold scaffold00268, whole genome shotgun sequence

JH723968.1 Dictyostelium firmibasis strain TNS-C-0014 unplaced genomic scaffold scaffold00269, whole genome shotgun sequence

AJWH01006599.1 Dictyostelium firmibasis strain TNS-C-0014 Contig14624, whole genome shotgun sequence

JH723969.1 Dictyostelium firmibasis strain TNS-C-0014 unplaced genomic scaffold scaffold00271, whole genome shotgun sequence

JH723970.1 Dictyostelium firmibasis strain TNS-C-0014 unplaced genomic scaffold scaffold00272, whole genome shotgun sequence

JH723971.1 Dictyostelium firmibasis strain TNS-C-0014 unplaced genomic scaffold scaffold00273, whole genome shotgun sequence

JH723972.1 Dictyostelium firmibasis strain TNS-C-0014 unplaced genomic scaffold scaffold00274, whole genome shotgun sequence

AJWH01006618.1 Dictyostelium firmibasis strain TNS-C-0014 Contig14680, whole genome shotgun sequence

JH723973.1 Dictyostelium firmibasis strain TNS-C-0014 unplaced genomic scaffold scaffold00276, whole genome shotgun sequence

JH723974.1 Dictyostelium firmibasis strain TNS-C-0014 unplaced genomic scaffold scaffold00277, whole genome shotgun sequence

AJWH01006626.1 Dictyostelium firmibasis strain TNS-C-0014 Contig14692, whole genome shotgun sequence

JH723975.1 Dictyostelium firmibasis strain TNS-C-0014 unplaced genomic scaffold scaffold00279, whole genome shotgun sequence

JH723976.1 Dictyostelium firmibasis strain TNS-C-0014 unplaced genomic scaffold scaffold00280, whole genome shotgun sequence

AJWH01006634.1 Dictyostelium firmibasis strain TNS-C-0014 Contig14704, whole genome shotgun sequence

AJWH01006635.1 Dictyostelium firmibasis strain TNS-C-0014 Contig14708, whole genome shotgun sequence

AJWH01006636.1 Dictyostelium firmibasis strain TNS-C-0014 Contig14709, whole genome shotgun sequence

JH723977.1 Dictyostelium firmibasis strain TNS-C-0014 unplaced genomic scaffold scaffold00284, whole genome shotgun sequence

AJWH01006641.1 Dictyostelium firmibasis strain TNS-C-0014 Contig14714, whole genome shotgun sequence

JH723978.1 Dictyostelium firmibasis strain TNS-C-0014 unplaced genomic scaffold scaffold00286, whole genome shotgun sequence

JH723979.1 Dictyostelium firmibasis strain TNS-C-0014 unplaced genomic scaffold scaffold00287, whole genome shotgun sequence

JH723980.1 Dictyostelium firmibasis strain TNS-C-0014 unplaced genomic scaffold scaffold00288, whole genome shotgun sequence

JH723981.1 Dictyostelium firmibasis strain TNS-C-0014 unplaced genomic scaffold scaffold00289, whole genome shotgun sequence

AJWH01006656.1 Dictyostelium firmibasis strain TNS-C-0014 Contig14729, whole genome shotgun sequence

JH723982.1 Dictyostelium firmibasis strain TNS-C-0014 unplaced genomic scaffold scaffold00291, whole genome shotgun sequence

AJWH01006661.1 Dictyostelium firmibasis strain TNS-C-0014 Contig14734, whole genome shotgun sequence

JH723983.1 Dictyostelium firmibasis strain TNS-C-0014 unplaced genomic scaffold scaffold00293, whole genome shotgun sequence

JH723984.1 Dictyostelium firmibasis strain TNS-C-0014 unplaced genomic scaffold scaffold00294, whole genome shotgun sequence

JH723985.1 Dictyostelium firmibasis strain TNS-C-0014 unplaced genomic scaffold scaffold00295, whole genome shotgun sequence

JH723986.1 Dictyostelium firmibasis strain TNS-C-0014 unplaced genomic scaffold scaffold00296, whole genome shotgun sequence

AJWH01006675.1 Dictyostelium firmibasis strain TNS-C-0014 Contig14751, whole genome shotgun sequence

AJWH01006676.1 Dictyostelium firmibasis strain TNS-C-0014 Contig14752, whole genome shotgun sequence

AJWH01006677.1 Dictyostelium firmibasis strain TNS-C-0014 Contig14759, whole genome shotgun sequence

JH723987.1 Dictyostelium firmibasis strain TNS-C-0014 unplaced genomic scaffold scaffold00300, whole genome shotgun sequence

JH723988.1 Dictyostelium firmibasis strain TNS-C-0014 unplaced genomic scaffold scaffold00301, whole genome shotgun sequence

JH723989.1 Dictyostelium firmibasis strain TNS-C-0014 unplaced genomic scaffold scaffold00302, whole genome shotgun sequence

JH723990.1 Dictyostelium firmibasis strain TNS-C-0014 unplaced genomic scaffold scaffold00303, whole genome shotgun sequence

JH723991.1 Dictyostelium firmibasis strain TNS-C-0014 unplaced genomic scaffold scaffold00304, whole genome shotgun sequence

AJWH01006691.1 Dictyostelium firmibasis strain TNS-C-0014 Contig14774, whole genome shotgun sequence

JH723992.1 Dictyostelium firmibasis strain TNS-C-0014 unplaced genomic scaffold scaffold00306, whole genome shotgun sequence

AJWH01006694.1 Dictyostelium firmibasis strain TNS-C-0014 Contig14777, whole genome shotgun sequence

AJWH01006695.1 Dictyostelium firmibasis strain TNS-C-0014 Contig14781, whole genome shotgun sequence

AJWH01006696.1 Dictyostelium firmibasis strain TNS-C-0014 Contig14789, whole genome shotgun sequence

JH723993.1 Dictyostelium firmibasis strain TNS-C-0014 unplaced genomic scaffold scaffold00310, whole genome shotgun sequence

AJWH01006699.1 Dictyostelium firmibasis strain TNS-C-0014 Contig14794, whole genome shotgun sequence

AJWH01006700.1 Dictyostelium firmibasis strain TNS-C-0014 Contig14795, whole genome shotgun sequence

AJWH01006701.1 Dictyostelium firmibasis strain TNS-C-0014 Contig14796, whole genome shotgun sequence

JH723994.1 Dictyostelium firmibasis strain TNS-C-0014 unplaced genomic scaffold scaffold00314, whole genome shotgun sequence

AJWH01006704.1 Dictyostelium firmibasis strain TNS-C-0014 Contig14799, whole genome shotgun sequence

AJWH01006705.1 Dictyostelium firmibasis strain TNS-C-0014 Contig14800, whole genome shotgun sequence

JH723995.1 Dictyostelium firmibasis strain TNS-C-0014 unplaced genomic scaffold scaffold00317, whole genome shotgun sequence

AJWH01006709.1 Dictyostelium firmibasis strain TNS-C-0014 Contig14813, whole genome shotgun sequence

AJWH01006710.1 Dictyostelium firmibasis strain TNS-C-0014 Contig14814, whole genome shotgun sequence

AJWH01006711.1 Dictyostelium firmibasis strain TNS-C-0014 Contig14820, whole genome shotgun sequence

AJWH01006712.1 Dictyostelium firmibasis strain TNS-C-0014 Contig14821, whole genome shotgun sequence

JH723996.1 Dictyostelium firmibasis strain TNS-C-0014 unplaced genomic scaffold scaffold00322, whole genome shotgun sequence

AJWH01006717.1 Dictyostelium firmibasis strain TNS-C-0014 Contig14826, whole genome shotgun sequence

AJWH01006718.1 Dictyostelium firmibasis strain TNS-C-0014 Contig14827, whole genome shotgun sequence

AJWH01006719.1 Dictyostelium firmibasis strain TNS-C-0014 Contig14830, whole genome shotgun sequence

JH723997.1 Dictyostelium firmibasis strain TNS-C-0014 unplaced genomic scaffold scaffold00326, whole genome shotgun sequence

AJWH01006725.1 Dictyostelium firmibasis strain TNS-C-0014 Contig14840, whole genome shotgun sequence

AJWH01006726.1 Dictyostelium firmibasis strain TNS-C-0014 Contig14841, whole genome shotgun sequence

JH723998.1 Dictyostelium firmibasis strain TNS-C-0014 unplaced genomic scaffold scaffold00329, whole genome shotgun sequence

JH723999.1 Dictyostelium firmibasis strain TNS-C-0014 unplaced genomic scaffold scaffold00330, whole genome shotgun sequence

JH724000.1 Dictyostelium firmibasis strain TNS-C-0014 unplaced genomic scaffold scaffold00331, whole genome shotgun sequence

AJWH01006735.1 Dictyostelium firmibasis strain TNS-C-0014 Contig14850, whole genome shotgun sequence

AJWH01006736.1 Dictyostelium firmibasis strain TNS-C-0014 Contig14852, whole genome shotgun sequence

JH724001.1 Dictyostelium firmibasis strain TNS-C-0014 unplaced genomic scaffold scaffold00334, whole genome shotgun sequence

AJWH01006740.1 Dictyostelium firmibasis strain TNS-C-0014 Contig14857, whole genome shotgun sequence

AJWH01006741.1 Dictyostelium firmibasis strain TNS-C-0014 Contig14858, whole genome shotgun sequence

AJWH01006742.1 Dictyostelium firmibasis strain TNS-C-0014 Contig14859, whole genome shotgun sequence

JH724002.1 Dictyostelium firmibasis strain TNS-C-0014 unplaced genomic scaffold scaffold00338, whole genome shotgun sequence

JH724003.1 Dictyostelium firmibasis strain TNS-C-0014 unplaced genomic scaffold scaffold00339, whole genome shotgun sequence

AJWH01006749.1 Dictyostelium firmibasis strain TNS-C-0014 Contig14866, whole genome shotgun sequence

AJWH01006750.1 Dictyostelium firmibasis strain TNS-C-0014 Contig14867, whole genome shotgun sequence

AJWH01006751.1 Dictyostelium firmibasis strain TNS-C-0014 Contig14868, whole genome shotgun sequence

JH724004.1 Dictyostelium firmibasis strain TNS-C-0014 unplaced genomic scaffold scaffold00343, whole genome shotgun sequence

AJWH01006754.1 Dictyostelium firmibasis strain TNS-C-0014 Contig14871, whole genome shotgun sequence

AJWH01006755.1 Dictyostelium firmibasis strain TNS-C-0014 Contig14878, whole genome shotgun sequence

AJWH01006756.1 Dictyostelium firmibasis strain TNS-C-0014 Contig14879, whole genome shotgun sequence

AJWH01006757.1 Dictyostelium firmibasis strain TNS-C-0014 Contig14880, whole genome shotgun sequence

AJWH01006758.1 Dictyostelium firmibasis strain TNS-C-0014 Contig14881, whole genome shotgun sequence

AJWH01006759.1 Dictyostelium firmibasis strain TNS-C-0014 Contig14882, whole genome shotgun sequence

AJWH01006760.1 Dictyostelium firmibasis strain TNS-C-0014 Contig14883, whole genome shotgun sequence

AJWH01006761.1 Dictyostelium firmibasis strain TNS-C-0014 Contig14884, whole genome shotgun sequence

AJWH01006762.1 Dictyostelium firmibasis strain TNS-C-0014 Contig14885, whole genome shotgun sequence

AJWH01006763.1 Dictyostelium firmibasis strain TNS-C-0014 Contig14886, whole genome shotgun sequence

JH724005.1 Dictyostelium firmibasis strain TNS-C-0014 unplaced genomic scaffold scaffold00354, whole genome shotgun sequence

AJWH01006767.1 Dictyostelium firmibasis strain TNS-C-0014 Contig14892, whole genome shotgun sequence

JH724006.1 Dictyostelium firmibasis strain TNS-C-0014 unplaced genomic scaffold scaffold00356, whole genome shotgun sequence

JH724007.1 Dictyostelium firmibasis strain TNS-C-0014 unplaced genomic scaffold scaffold00357, whole genome shotgun sequence

JH724008.1 Dictyostelium firmibasis strain TNS-C-0014 unplaced genomic scaffold scaffold00358, whole genome shotgun sequence

AJWH01006775.1 Dictyostelium firmibasis strain TNS-C-0014 Contig14900, whole genome shotgun sequence

AJWH01006776.1 Dictyostelium firmibasis strain TNS-C-0014 Contig14901, whole genome shotgun sequence

AJWH01006777.1 Dictyostelium firmibasis strain TNS-C-0014 Contig14903, whole genome shotgun sequence

AJWH01006778.1 Dictyostelium firmibasis strain TNS-C-0014 Contig14906, whole genome shotgun sequence

AJWH01006779.1 Dictyostelium firmibasis strain TNS-C-0014 Contig14907, whole genome shotgun sequence

AJWH01006780.1 Dictyostelium firmibasis strain TNS-C-0014 Contig14908, whole genome shotgun sequence

AJWH01006781.1 Dictyostelium firmibasis strain TNS-C-0014 Contig14909, whole genome shotgun sequence

AJWH01006782.1 Dictyostelium firmibasis strain TNS-C-0014 Contig14910, whole genome shotgun sequence

JH724009.1 Dictyostelium firmibasis strain TNS-C-0014 unplaced genomic scaffold scaffold00367, whole genome shotgun sequence

JH724010.1 Dictyostelium firmibasis strain TNS-C-0014 unplaced genomic scaffold scaffold00368, whole genome shotgun sequence

AJWH01006787.1 Dictyostelium firmibasis strain TNS-C-0014 Contig14915, whole genome shotgun sequence

JH724011.1 Dictyostelium firmibasis strain TNS-C-0014 unplaced genomic scaffold scaffold00370, whole genome shotgun sequence

JH724012.1 Dictyostelium firmibasis strain TNS-C-0014 unplaced genomic scaffold scaffold00371, whole genome shotgun sequence

AJWH01006794.1 Dictyostelium firmibasis strain TNS-C-0014 Contig14925, whole genome shotgun sequence

AJWH01006795.1 Dictyostelium firmibasis strain TNS-C-0014 Contig14926, whole genome shotgun sequence

AJWH01006796.1 Dictyostelium firmibasis strain TNS-C-0014 Contig14931, whole genome shotgun sequence

AJWH01006797.1 Dictyostelium firmibasis strain TNS-C-0014 Contig14933, whole genome shotgun sequence

JH724013.1 Dictyostelium firmibasis strain TNS-C-0014 unplaced genomic scaffold scaffold00376, whole genome shotgun sequence

JH724014.1 Dictyostelium firmibasis strain TNS-C-0014 unplaced genomic scaffold scaffold00377, whole genome shotgun sequence

AJWH01006804.1 Dictyostelium firmibasis strain TNS-C-0014 Contig14940, whole genome shotgun sequence

AJWH01006805.1 Dictyostelium firmibasis strain TNS-C-0014 Contig14941, whole genome shotgun sequence

AJWH01006806.1 Dictyostelium firmibasis strain TNS-C-0014 Contig14942, whole genome shotgun sequence

AJWH01006807.1 Dictyostelium firmibasis strain TNS-C-0014 Contig14944, whole genome shotgun sequence

AJWH01006808.1 Dictyostelium firmibasis strain TNS-C-0014 Contig14945, whole genome shotgun sequence

AJWH01006809.1 Dictyostelium firmibasis strain TNS-C-0014 Contig14946, whole genome shotgun sequence

JH724015.1 Dictyostelium firmibasis strain TNS-C-0014 unplaced genomic scaffold scaffold00384, whole genome shotgun sequence

JH724016.1 Dictyostelium firmibasis strain TNS-C-0014 unplaced genomic scaffold scaffold00385, whole genome shotgun sequence

AJWH01006817.1 Dictyostelium firmibasis strain TNS-C-0014 Contig14956, whole genome shotgun sequence

AJWH01006818.1 Dictyostelium firmibasis strain TNS-C-0014 Contig14957, whole genome shotgun sequence

JH724017.1 Dictyostelium firmibasis strain TNS-C-0014 unplaced genomic scaffold scaffold00388, whole genome shotgun sequence

JH724018.1 Dictyostelium firmibasis strain TNS-C-0014 unplaced genomic scaffold scaffold00389, whole genome shotgun sequence

JH724019.1 Dictyostelium firmibasis strain TNS-C-0014 unplaced genomic scaffold scaffold00390, whole genome shotgun sequence

AJWH01006827.1 Dictyostelium firmibasis strain TNS-C-0014 Contig14969, whole genome shotgun sequence

JH724020.1 Dictyostelium firmibasis strain TNS-C-0014 unplaced genomic scaffold scaffold00392, whole genome shotgun sequence

JH724021.1 Dictyostelium firmibasis strain TNS-C-0014 unplaced genomic scaffold scaffold00393, whole genome shotgun sequence

AJWH01006833.1 Dictyostelium firmibasis strain TNS-C-0014 Contig14977, whole genome shotgun sequence

AJWH01006834.1 Dictyostelium firmibasis strain TNS-C-0014 Contig14979, whole genome shotgun sequence

AJWH01006835.1 Dictyostelium firmibasis strain TNS-C-0014 Contig14980, whole genome shotgun sequence

JH724022.1 Dictyostelium firmibasis strain TNS-C-0014 unplaced genomic scaffold scaffold00397, whole genome shotgun sequence

AJWH01006839.1 Dictyostelium firmibasis strain TNS-C-0014 Contig14984, whole genome shotgun sequence

JH724023.1 Dictyostelium firmibasis strain TNS-C-0014 unplaced genomic scaffold scaffold00399, whole genome shotgun sequence

JH724024.1 Dictyostelium firmibasis strain TNS-C-0014 unplaced genomic scaffold scaffold00400, whole genome shotgun sequence

JH724025.1 Dictyostelium firmibasis strain TNS-C-0014 unplaced genomic scaffold scaffold00401, whole genome shotgun sequence

AJWH01006848.1 Dictyostelium firmibasis strain TNS-C-0014 Contig14993, whole genome shotgun sequence

JH724026.1 Dictyostelium firmibasis strain TNS-C-0014 unplaced genomic scaffold scaffold00403, whole genome shotgun sequence

JH724027.1 Dictyostelium firmibasis strain TNS-C-0014 unplaced genomic scaffold scaffold00404, whole genome shotgun sequence

AJWH01006855.1 Dictyostelium firmibasis strain TNS-C-0014 Contig15002, whole genome shotgun sequence

AJWH01006856.1 Dictyostelium firmibasis strain TNS-C-0014 Contig15003, whole genome shotgun sequence

JH724028.1 Dictyostelium firmibasis strain TNS-C-0014 unplaced genomic scaffold scaffold00407, whole genome shotgun sequence

AJWH01006859.1 Dictyostelium firmibasis strain TNS-C-0014 Contig15007, whole genome shotgun sequence

AJWH01006860.1 Dictyostelium firmibasis strain TNS-C-0014 Contig15009, whole genome shotgun sequence

AJWH01006861.1 Dictyostelium firmibasis strain TNS-C-0014 Contig15010, whole genome shotgun sequence

AJWH01006862.1 Dictyostelium firmibasis strain TNS-C-0014 Contig15011, whole genome shotgun sequence

AJWH01006863.1 Dictyostelium firmibasis strain TNS-C-0014 Contig15012, whole genome shotgun sequence

AJWH01006864.1 Dictyostelium firmibasis strain TNS-C-0014 Contig15013, whole genome shotgun sequence

AJWH01006865.1 Dictyostelium firmibasis strain TNS-C-0014 Contig15014, whole genome shotgun sequence

JH724029.1 Dictyostelium firmibasis strain TNS-C-0014 unplaced genomic scaffold scaffold00415, whole genome shotgun sequence

AJWH01006869.1 Dictyostelium firmibasis strain TNS-C-0014 Contig15021, whole genome shotgun sequence

AJWH01006870.1 Dictyostelium firmibasis strain TNS-C-0014 Contig15022, whole genome shotgun sequence

AJWH01006871.1 Dictyostelium firmibasis strain TNS-C-0014 Contig15023, whole genome shotgun sequence

AJWH01006872.1 Dictyostelium firmibasis strain TNS-C-0014 Contig15025, whole genome shotgun sequence

AJWH01006873.1 Dictyostelium firmibasis strain TNS-C-0014 Contig15026, whole genome shotgun sequence

JH724030.1 Dictyostelium firmibasis strain TNS-C-0014 unplaced genomic scaffold scaffold00421, whole genome shotgun sequence

JH724031.1 Dictyostelium firmibasis strain TNS-C-0014 unplaced genomic scaffold scaffold00422, whole genome shotgun sequence

AJWH01006879.1 Dictyostelium firmibasis strain TNS-C-0014 Contig15032, whole genome shotgun sequence

JH724032.1 Dictyostelium firmibasis strain TNS-C-0014 unplaced genomic scaffold scaffold00424, whole genome shotgun sequence

AJWH01006883.1 Dictyostelium firmibasis strain TNS-C-0014 Contig15039, whole genome shotgun sequence

JH724033.1 Dictyostelium firmibasis strain TNS-C-0014 unplaced genomic scaffold scaffold00426, whole genome shotgun sequence

AJWH01006887.1 Dictyostelium firmibasis strain TNS-C-0014 Contig15043, whole genome shotgun sequence

AJWH01006888.1 Dictyostelium firmibasis strain TNS-C-0014 Contig15045, whole genome shotgun sequence

AJWH01006889.1 Dictyostelium firmibasis strain TNS-C-0014 Contig15046, whole genome shotgun sequence

JH724034.1 Dictyostelium firmibasis strain TNS-C-0014 unplaced genomic scaffold scaffold00430, whole genome shotgun sequence

JH724035.1 Dictyostelium firmibasis strain TNS-C-0014 unplaced genomic scaffold scaffold00431, whole genome shotgun sequence

AJWH01006896.1 Dictyostelium firmibasis strain TNS-C-0014 Contig15053, whole genome shotgun sequence

JH724036.1 Dictyostelium firmibasis strain TNS-C-0014 unplaced genomic scaffold scaffold00433, whole genome shotgun sequence

AJWH01006899.1 Dictyostelium firmibasis strain TNS-C-0014 Contig15056, whole genome shotgun sequence

AJWH01006900.1 Dictyostelium firmibasis strain TNS-C-0014 Contig15057, whole genome shotgun sequence

AJWH01006901.1 Dictyostelium firmibasis strain TNS-C-0014 Contig15058, whole genome shotgun sequence

AJWH01006902.1 Dictyostelium firmibasis strain TNS-C-0014 Contig15059, whole genome shotgun sequence

AJWH01006903.1 Dictyostelium firmibasis strain TNS-C-0014 Contig15060, whole genome shotgun sequence

AJWH01006904.1 Dictyostelium firmibasis strain TNS-C-0014 Contig15061, whole genome shotgun sequence

AJWH01006905.1 Dictyostelium firmibasis strain TNS-C-0014 Contig15062, whole genome shotgun sequence

AJWH01006906.1 Dictyostelium firmibasis strain TNS-C-0014 Contig15063, whole genome shotgun sequence

AJWH01006907.1 Dictyostelium firmibasis strain TNS-C-0014 Contig15064, whole genome shotgun sequence

AJWH01006908.1 Dictyostelium firmibasis strain TNS-C-0014 Contig15065, whole genome shotgun sequence

AJWH01006909.1 Dictyostelium firmibasis strain TNS-C-0014 Contig15066, whole genome shotgun sequence

AJWH01006910.1 Dictyostelium firmibasis strain TNS-C-0014 Contig15067, whole genome shotgun sequence

AJWH01006911.1 Dictyostelium firmibasis strain TNS-C-0014 Contig15068, whole genome shotgun sequence

JH724037.1 Dictyostelium firmibasis strain TNS-C-0014 unplaced genomic scaffold scaffold00447, whole genome shotgun sequence

JH724038.1 Dictyostelium firmibasis strain TNS-C-0014 unplaced genomic scaffold scaffold00448, whole genome shotgun sequence

AJWH01006917.1 Dictyostelium firmibasis strain TNS-C-0014 Contig15074, whole genome shotgun sequence

AJWH01006918.1 Dictyostelium firmibasis strain TNS-C-0014 Contig15078, whole genome shotgun sequence

AJWH01006919.1 Dictyostelium firmibasis strain TNS-C-0014 Contig15079, whole genome shotgun sequence

AJWH01006920.1 Dictyostelium firmibasis strain TNS-C-0014 Contig15080, whole genome shotgun sequence

AJWH01006921.1 Dictyostelium firmibasis strain TNS-C-0014 Contig15081, whole genome shotgun sequence

JH724039.1 Dictyostelium firmibasis strain TNS-C-0014 unplaced genomic scaffold scaffold00454, whole genome shotgun sequence

JH724040.1 Dictyostelium firmibasis strain TNS-C-0014 unplaced genomic scaffold scaffold00455, whole genome shotgun sequence

AJWH01006926.1 Dictyostelium firmibasis strain TNS-C-0014 Contig15086, whole genome shotgun sequence

AJWH01006927.1 Dictyostelium firmibasis strain TNS-C-0014 Contig15087, whole genome shotgun sequence

AJWH01006928.1 Dictyostelium firmibasis strain TNS-C-0014 Contig15088, whole genome shotgun sequence

AJWH01006929.1 Dictyostelium firmibasis strain TNS-C-0014 Contig15089, whole genome shotgun sequence

AJWH01006930.1 Dictyostelium firmibasis strain TNS-C-0014 Contig15090, whole genome shotgun sequence

AJWH01006931.1 Dictyostelium firmibasis strain TNS-C-0014 Contig15093, whole genome shotgun sequence

AJWH01006932.1 Dictyostelium firmibasis strain TNS-C-0014 Contig15094, whole genome shotgun sequence

AJWH01006933.1 Dictyostelium firmibasis strain TNS-C-0014 Contig15096, whole genome shotgun sequence

AJWH01006934.1 Dictyostelium firmibasis strain TNS-C-0014 Contig15097, whole genome shotgun sequence

AJWH01006935.1 Dictyostelium firmibasis strain TNS-C-0014 Contig15098, whole genome shotgun sequence

AJWH01006936.1 Dictyostelium firmibasis strain TNS-C-0014 Contig15099, whole genome shotgun sequence

AJWH01006937.1 Dictyostelium firmibasis strain TNS-C-0014 Contig15100, whole genome shotgun sequence

AJWH01006938.1 Dictyostelium firmibasis strain TNS-C-0014 Contig15102, whole genome shotgun sequence

AJWH01006939.1 Dictyostelium firmibasis strain TNS-C-0014 Contig15103, whole genome shotgun sequence

JH724041.1 Dictyostelium firmibasis strain TNS-C-0014 unplaced genomic scaffold scaffold00470, whole genome shotgun sequence

AJWH01006942.1 Dictyostelium firmibasis strain TNS-C-0014 Contig15106, whole genome shotgun sequence

AJWH01006943.1 Dictyostelium firmibasis strain TNS-C-0014 Contig15107, whole genome shotgun sequence

AJWH01006944.1 Dictyostelium firmibasis strain TNS-C-0014 Contig15108, whole genome shotgun sequence

AJWH01006945.1 Dictyostelium firmibasis strain TNS-C-0014 Contig15109, whole genome shotgun sequence

AJWH01006946.1 Dictyostelium firmibasis strain TNS-C-0014 Contig15110, whole genome shotgun sequence

JH724042.1 Dictyostelium firmibasis strain TNS-C-0014 unplaced genomic scaffold scaffold00476, whole genome shotgun sequence

AJWH01006949.1 Dictyostelium firmibasis strain TNS-C-0014 Contig15113, whole genome shotgun sequence

AJWH01006950.1 Dictyostelium firmibasis strain TNS-C-0014 Contig15114, whole genome shotgun sequence

AJWH01006951.1 Dictyostelium firmibasis strain TNS-C-0014 Contig15115, whole genome shotgun sequence

AJWH01006952.1 Dictyostelium firmibasis strain TNS-C-0014 Contig15116, whole genome shotgun sequence

AJWH01006953.1 Dictyostelium firmibasis strain TNS-C-0014 Contig15117, whole genome shotgun sequence

AJWH01006954.1 Dictyostelium firmibasis strain TNS-C-0014 Contig15118, whole genome shotgun sequence

AJWH01006955.1 Dictyostelium firmibasis strain TNS-C-0014 Contig15119, whole genome shotgun sequence

AJWH01006956.1 Dictyostelium firmibasis strain TNS-C-0014 Contig15120, whole genome shotgun sequence

AJWH01006957.1 Dictyostelium firmibasis strain TNS-C-0014 Contig15121, whole genome shotgun sequence

JH724043.1 Dictyostelium firmibasis strain TNS-C-0014 unplaced genomic scaffold scaffold00486, whole genome shotgun sequence

JH724044.1 Dictyostelium firmibasis strain TNS-C-0014 unplaced genomic scaffold scaffold00487, whole genome shotgun sequence

AJWH01006962.1 Dictyostelium firmibasis strain TNS-C-0014 Contig15126, whole genome shotgun sequence

AJWH01006963.1 Dictyostelium firmibasis strain TNS-C-0014 Contig15128, whole genome shotgun sequence

JH724045.1 Dictyostelium firmibasis strain TNS-C-0014 unplaced genomic scaffold scaffold00490, whole genome shotgun sequence

AJWH01006966.1 Dictyostelium firmibasis strain TNS-C-0014 Contig15131, whole genome shotgun sequence

AJWH01006967.1 Dictyostelium firmibasis strain TNS-C-0014 Contig15132, whole genome shotgun sequence

AJWH01006968.1 Dictyostelium firmibasis strain TNS-C-0014 Contig15133, whole genome shotgun sequence

AJWH01006969.1 Dictyostelium firmibasis strain TNS-C-0014 Contig15134, whole genome shotgun sequence

AJWH01006970.1 Dictyostelium firmibasis strain TNS-C-0014 Contig15135, whole genome shotgun sequence

AJWH01006971.1 Dictyostelium firmibasis strain TNS-C-0014 Contig15136, whole genome shotgun sequence

AJWH01006972.1 Dictyostelium firmibasis strain TNS-C-0014 Contig15137, whole genome shotgun sequence

AJWH01006973.1 Dictyostelium firmibasis strain TNS-C-0014 Contig15138, whole genome shotgun sequence

JH724046.1 Dictyostelium firmibasis strain TNS-C-0014 unplaced genomic scaffold scaffold00499, whole genome shotgun sequence

AJWH01006976.1 Dictyostelium firmibasis strain TNS-C-0014 Contig15141, whole genome shotgun sequence

AJWH01006977.1 Dictyostelium firmibasis strain TNS-C-0014 Contig15142, whole genome shotgun sequence

AJWH01006978.1 Dictyostelium firmibasis strain TNS-C-0014 Contig15143, whole genome shotgun sequence

AJWH01006979.1 Dictyostelium firmibasis strain TNS-C-0014 Contig15144, whole genome shotgun sequence

AJWH01006980.1 Dictyostelium firmibasis strain TNS-C-0014 Contig15145, whole genome shotgun sequence

AJWH01006981.1 Dictyostelium firmibasis strain TNS-C-0014 Contig15146, whole genome shotgun sequence

AJWH01006982.1 Dictyostelium firmibasis strain TNS-C-0014 Contig15147, whole genome shotgun sequence

AJWH01006983.1 Dictyostelium firmibasis strain TNS-C-0014 Contig15148, whole genome shotgun sequence

JH724047.1 Dictyostelium firmibasis strain TNS-C-0014 unplaced genomic scaffold scaffold00508, whole genome shotgun sequence

AJWH01006987.1 Dictyostelium firmibasis strain TNS-C-0014 Contig15152, whole genome shotgun sequence

AJWH01006988.1 Dictyostelium firmibasis strain TNS-C-0014 Contig15153, whole genome shotgun sequence

AJWH01006989.1 Dictyostelium firmibasis strain TNS-C-0014 Contig15154, whole genome shotgun sequence

AJWH01006990.1 Dictyostelium firmibasis strain TNS-C-0014 Contig15155, whole genome shotgun sequence

AJWH01006991.1 Dictyostelium firmibasis strain TNS-C-0014 Contig15156, whole genome shotgun sequence

AJWH01006992.1 Dictyostelium firmibasis strain TNS-C-0014 Contig15157, whole genome shotgun sequence

AJWH01006993.1 Dictyostelium firmibasis strain TNS-C-0014 Contig15158, whole genome shotgun sequence

JH724048.1 Dictyostelium firmibasis strain TNS-C-0014 unplaced genomic scaffold scaffold00516, whole genome shotgun sequence

AJWH01006996.1 Dictyostelium firmibasis strain TNS-C-0014 Contig15161, whole genome shotgun sequence

AJWH01006997.1 Dictyostelium firmibasis strain TNS-C-0014 Contig15162, whole genome shotgun sequence

AJWH01006998.1 Dictyostelium firmibasis strain TNS-C-0014 Contig15163, whole genome shotgun sequence

JH724049.1 Dictyostelium firmibasis strain TNS-C-0014 unplaced genomic scaffold scaffold00520, whole genome shotgun sequence

AJWH01007001.1 Dictyostelium firmibasis strain TNS-C-0014 Contig15166, whole genome shotgun sequence

AJWH01007002.1 Dictyostelium firmibasis strain TNS-C-0014 Contig15167, whole genome shotgun sequence

AJWH01007003.1 Dictyostelium firmibasis strain TNS-C-0014 Contig15168, whole genome shotgun sequence

AJWH01007004.1 Dictyostelium firmibasis strain TNS-C-0014 Contig15169, whole genome shotgun sequence

JH724050.1 Dictyostelium firmibasis strain TNS-C-0014 unplaced genomic scaffold scaffold00525, whole genome shotgun sequence

AJWH01007007.1 Dictyostelium firmibasis strain TNS-C-0014 Contig15174, whole genome shotgun sequence

AJWH01007008.1 Dictyostelium firmibasis strain TNS-C-0014 Contig15175, whole genome shotgun sequence

AJWH01007009.1 Dictyostelium firmibasis strain TNS-C-0014 Contig15176, whole genome shotgun sequence

AJWH01007010.1 Dictyostelium firmibasis strain TNS-C-0014 Contig15177, whole genome shotgun sequence

JH724051.1 Dictyostelium firmibasis strain TNS-C-0014 unplaced genomic scaffold scaffold00530, whole genome shotgun sequence

JH724052.1 Dictyostelium firmibasis strain TNS-C-0014 unplaced genomic scaffold scaffold00531, whole genome shotgun sequence

AJWH01007015.1 Dictyostelium firmibasis strain TNS-C-0014 Contig15182, whole genome shotgun sequence

JH724053.1 Dictyostelium firmibasis strain TNS-C-0014 unplaced genomic scaffold scaffold00533, whole genome shotgun sequence

AJWH01007018.1 Dictyostelium firmibasis strain TNS-C-0014 Contig15188, whole genome shotgun sequence

AJWH01007019.1 Dictyostelium firmibasis strain TNS-C-0014 Contig15189, whole genome shotgun sequence

AJWH01007020.1 Dictyostelium firmibasis strain TNS-C-0014 Contig15190, whole genome shotgun sequence

AJWH01007021.1 Dictyostelium firmibasis strain TNS-C-0014 Contig15191, whole genome shotgun sequence

AJWH01007022.1 Dictyostelium firmibasis strain TNS-C-0014 Contig15192, whole genome shotgun sequence

AJWH01007023.1 Dictyostelium firmibasis strain TNS-C-0014 Contig15193, whole genome shotgun sequence

AJWH01007024.1 Dictyostelium firmibasis strain TNS-C-0014 Contig15194, whole genome shotgun sequence

JH724054.1 Dictyostelium firmibasis strain TNS-C-0014 unplaced genomic scaffold scaffold00541, whole genome shotgun sequence

AJWH01007027.1 Dictyostelium firmibasis strain TNS-C-0014 Contig15197, whole genome shotgun sequence

AJWH01007028.1 Dictyostelium firmibasis strain TNS-C-0014 Contig15198, whole genome shotgun sequence

AJWH01007029.1 Dictyostelium firmibasis strain TNS-C-0014 Contig15199, whole genome shotgun sequence

AJWH01007030.1 Dictyostelium firmibasis strain TNS-C-0014 Contig15200, whole genome shotgun sequence

JH724055.1 Dictyostelium firmibasis strain TNS-C-0014 unplaced genomic scaffold scaffold00546, whole genome shotgun sequence

AJWH01007033.1 Dictyostelium firmibasis strain TNS-C-0014 Contig15203, whole genome shotgun sequence

AJWH01007034.1 Dictyostelium firmibasis strain TNS-C-0014 Contig15204, whole genome shotgun sequence

AJWH01007035.1 Dictyostelium firmibasis strain TNS-C-0014 Contig15205, whole genome shotgun sequence

AJWH01007036.1 Dictyostelium firmibasis strain TNS-C-0014 Contig15206, whole genome shotgun sequence

AJWH01007037.1 Dictyostelium firmibasis strain TNS-C-0014 Contig15208, whole genome shotgun sequence

JH724056.1 Dictyostelium firmibasis strain TNS-C-0014 unplaced genomic scaffold scaffold00552, whole genome shotgun sequence

AJWH01007040.1 Dictyostelium firmibasis strain TNS-C-0014 Contig15211, whole genome shotgun sequence

AJWH01007041.1 Dictyostelium firmibasis strain TNS-C-0014 Contig15213, whole genome shotgun sequence

AJWH01007042.1 Dictyostelium firmibasis strain TNS-C-0014 Contig15214, whole genome shotgun sequence

AJWH01007043.1 Dictyostelium firmibasis strain TNS-C-0014 Contig15215, whole genome shotgun sequence

AJWH01007044.1 Dictyostelium firmibasis strain TNS-C-0014 Contig15218, whole genome shotgun sequence

AJWH01007045.1 Dictyostelium firmibasis strain TNS-C-0014 Contig15219, whole genome shotgun sequence

AJWH01007046.1 Dictyostelium firmibasis strain TNS-C-0014 Contig15221, whole genome shotgun sequence

AJWH01007047.1 Dictyostelium firmibasis strain TNS-C-0014 Contig15222, whole genome shotgun sequence

JH724057.1 Dictyostelium firmibasis strain TNS-C-0014 unplaced genomic scaffold scaffold00561, whole genome shotgun sequence

JH724058.1 Dictyostelium firmibasis strain TNS-C-0014 unplaced genomic scaffold scaffold00562, whole genome shotgun sequence

AJWH01007052.1 Dictyostelium firmibasis strain TNS-C-0014 Contig15227, whole genome shotgun sequence

AJWH01007053.1 Dictyostelium firmibasis strain TNS-C-0014 Contig15230, whole genome shotgun sequence

JH724059.1 Dictyostelium firmibasis strain TNS-C-0014 unplaced genomic scaffold scaffold00565, whole genome shotgun sequence

AJWH01007056.1 Dictyostelium firmibasis strain TNS-C-0014 Contig15233, whole genome shotgun sequence

AJWH01007057.1 Dictyostelium firmibasis strain TNS-C-0014 Contig15234, whole genome shotgun sequence

JH724060.1 Dictyostelium firmibasis strain TNS-C-0014 unplaced genomic scaffold scaffold00568, whole genome shotgun sequence

JH724061.1 Dictyostelium firmibasis strain TNS-C-0014 unplaced genomic scaffold scaffold00569, whole genome shotgun sequence

AJWH01007062.1 Dictyostelium firmibasis strain TNS-C-0014 Contig15241, whole genome shotgun sequence

AJWH01007063.1 Dictyostelium firmibasis strain TNS-C-0014 Contig15242, whole genome shotgun sequence

AJWH01007064.1 Dictyostelium firmibasis strain TNS-C-0014 Contig15243, whole genome shotgun sequence

AJWH01007065.1 Dictyostelium firmibasis strain TNS-C-0014 Contig15244, whole genome shotgun sequence

AJWH01007066.1 Dictyostelium firmibasis strain TNS-C-0014 Contig15245, whole genome shotgun sequence

AJWH01007067.1 Dictyostelium firmibasis strain TNS-C-0014 Contig15246, whole genome shotgun sequence

AJWH01007068.1 Dictyostelium firmibasis strain TNS-C-0014 Contig15247, whole genome shotgun sequence

AJWH01007069.1 Dictyostelium firmibasis strain TNS-C-0014 Contig15248, whole genome shotgun sequence

AJWH01007070.1 Dictyostelium firmibasis strain TNS-C-0014 Contig15249, whole genome shotgun sequence

AJWH01007071.1 Dictyostelium firmibasis strain TNS-C-0014 Contig15250, whole genome shotgun sequence

AJWH01007072.1 Dictyostelium firmibasis strain TNS-C-0014 Contig15251, whole genome shotgun sequence

JH724062.1 Dictyostelium firmibasis strain TNS-C-0014 unplaced genomic scaffold scaffold00581, whole genome shotgun sequence

AJWH01007075.1 Dictyostelium firmibasis strain TNS-C-0014 Contig15254, whole genome shotgun sequence

AJWH01007076.1 Dictyostelium firmibasis strain TNS-C-0014 Contig15255, whole genome shotgun sequence

AJWH01007077.1 Dictyostelium firmibasis strain TNS-C-0014 Contig15256, whole genome shotgun sequence

AJWH01007078.1 Dictyostelium firmibasis strain TNS-C-0014 Contig15258, whole genome shotgun sequence

AJWH01007079.1 Dictyostelium firmibasis strain TNS-C-0014 Contig15259, whole genome shotgun sequence

AJWH01007080.1 Dictyostelium firmibasis strain TNS-C-0014 Contig15260, whole genome shotgun sequence

JH724063.1 Dictyostelium firmibasis strain TNS-C-0014 unplaced genomic scaffold scaffold00588, whole genome shotgun sequence

JH724064.1 Dictyostelium firmibasis strain TNS-C-0014 unplaced genomic scaffold scaffold00589, whole genome shotgun sequence

JH724065.1 Dictyostelium firmibasis strain TNS-C-0014 unplaced genomic scaffold scaffold00590, whole genome shotgun sequence

AJWH01007087.1 Dictyostelium firmibasis strain TNS-C-0014 Contig15267, whole genome shotgun sequence

JH724066.1 Dictyostelium firmibasis strain TNS-C-0014 unplaced genomic scaffold scaffold00592, whole genome shotgun sequence

AJWH01007090.1 Dictyostelium firmibasis strain TNS-C-0014 Contig15270, whole genome shotgun sequence

AJWH01007091.1 Dictyostelium firmibasis strain TNS-C-0014 Contig15271, whole genome shotgun sequence

JH724067.1 Dictyostelium firmibasis strain TNS-C-0014 unplaced genomic scaffold scaffold00595, whole genome shotgun sequence

AJWH01007094.1 Dictyostelium firmibasis strain TNS-C-0014 Contig15274, whole genome shotgun sequence

AJWH01007095.1 Dictyostelium firmibasis strain TNS-C-0014 Contig15275, whole genome shotgun sequence

AJWH01007096.1 Dictyostelium firmibasis strain TNS-C-0014 Contig15276, whole genome shotgun sequence

AJWH01007097.1 Dictyostelium firmibasis strain TNS-C-0014 Contig15277, whole genome shotgun sequence

JH724068.1 Dictyostelium firmibasis strain TNS-C-0014 unplaced genomic scaffold scaffold00600, whole genome shotgun sequence

JH724069.1 Dictyostelium firmibasis strain TNS-C-0014 unplaced genomic scaffold scaffold00601, whole genome shotgun sequence

AJWH01007102.1 Dictyostelium firmibasis strain TNS-C-0014 Contig15282, whole genome shotgun sequence

AJWH01007103.1 Dictyostelium firmibasis strain TNS-C-0014 Contig15283, whole genome shotgun sequence

AJWH01007104.1 Dictyostelium firmibasis strain TNS-C-0014 Contig15284, whole genome shotgun sequence

AJWH01007105.1 Dictyostelium firmibasis strain TNS-C-0014 Contig15285, whole genome shotgun sequence

AJWH01007106.1 Dictyostelium firmibasis strain TNS-C-0014 Contig15286, whole genome shotgun sequence

AJWH01007107.1 Dictyostelium firmibasis strain TNS-C-0014 Contig15287, whole genome shotgun sequence

AJWH01007108.1 Dictyostelium firmibasis strain TNS-C-0014 Contig15288, whole genome shotgun sequence

AJWH01007109.1 Dictyostelium firmibasis strain TNS-C-0014 Contig15289, whole genome shotgun sequence

AJWH01007110.1 Dictyostelium firmibasis strain TNS-C-0014 Contig15290, whole genome shotgun sequence

AJWH01007111.1 Dictyostelium firmibasis strain TNS-C-0014 Contig15291, whole genome shotgun sequence

AJWH01007112.1 Dictyostelium firmibasis strain TNS-C-0014 Contig15292, whole genome shotgun sequence

AJWH01007113.1 Dictyostelium firmibasis strain TNS-C-0014 Contig15293, whole genome shotgun sequence

AJWH01007114.1 Dictyostelium firmibasis strain TNS-C-0014 Contig15294, whole genome shotgun sequence

AJWH01007115.1 Dictyostelium firmibasis strain TNS-C-0014 Contig15297, whole genome shotgun sequence

JH724070.1 Dictyostelium firmibasis strain TNS-C-0014 unplaced genomic scaffold scaffold00616, whole genome shotgun sequence

JH724071.1 Dictyostelium firmibasis strain TNS-C-0014 unplaced genomic scaffold scaffold00617, whole genome shotgun sequence

AJWH01007120.1 Dictyostelium firmibasis strain TNS-C-0014 Contig15302, whole genome shotgun sequence

AJWH01007121.1 Dictyostelium firmibasis strain TNS-C-0014 Contig15303, whole genome shotgun sequence

AJWH01007122.1 Dictyostelium firmibasis strain TNS-C-0014 Contig15308, whole genome shotgun sequence

AJWH01007123.1 Dictyostelium firmibasis strain TNS-C-0014 Contig15309, whole genome shotgun sequence

AJWH01007124.1 Dictyostelium firmibasis strain TNS-C-0014 Contig15310, whole genome shotgun sequence

AJWH01007125.1 Dictyostelium firmibasis strain TNS-C-0014 Contig15313, whole genome shotgun sequence

JH724072.1 Dictyostelium firmibasis strain TNS-C-0014 unplaced genomic scaffold scaffold00624, whole genome shotgun sequence

AJWH01007128.1 Dictyostelium firmibasis strain TNS-C-0014 Contig15316, whole genome shotgun sequence

AJWH01007129.1 Dictyostelium firmibasis strain TNS-C-0014 Contig15317, whole genome shotgun sequence

JH724073.1 Dictyostelium firmibasis strain TNS-C-0014 unplaced genomic scaffold scaffold00627, whole genome shotgun sequence

AJWH01007132.1 Dictyostelium firmibasis strain TNS-C-0014 Contig15320, whole genome shotgun sequence

JH724074.1 Dictyostelium firmibasis strain TNS-C-0014 unplaced genomic scaffold scaffold00629, whole genome shotgun sequence

AJWH01007135.1 Dictyostelium firmibasis strain TNS-C-0014 Contig15323, whole genome shotgun sequence

AJWH01007136.1 Dictyostelium firmibasis strain TNS-C-0014 Contig15324, whole genome shotgun sequence

AJWH01007137.1 Dictyostelium firmibasis strain TNS-C-0014 Contig15326, whole genome shotgun sequence

AJWH01007138.1 Dictyostelium firmibasis strain TNS-C-0014 Contig15327, whole genome shotgun sequence

AJWH01007139.1 Dictyostelium firmibasis strain TNS-C-0014 Contig15328, whole genome shotgun sequence

AJWH01007140.1 Dictyostelium firmibasis strain TNS-C-0014 Contig15329, whole genome shotgun sequence

AJWH01007141.1 Dictyostelium firmibasis strain TNS-C-0014 Contig15330, whole genome shotgun sequence

AJWH01007142.1 Dictyostelium firmibasis strain TNS-C-0014 Contig15332, whole genome shotgun sequence

JH724075.1 Dictyostelium firmibasis strain TNS-C-0014 unplaced genomic scaffold scaffold00638, whole genome shotgun sequence

AJWH01007145.1 Dictyostelium firmibasis strain TNS-C-0014 Contig15336, whole genome shotgun sequence

AJWH01007146.1 Dictyostelium firmibasis strain TNS-C-0014 Contig15337, whole genome shotgun sequence

AJWH01007147.1 Dictyostelium firmibasis strain TNS-C-0014 Contig15338, whole genome shotgun sequence

AJWH01007148.1 Dictyostelium firmibasis strain TNS-C-0014 Contig15339, whole genome shotgun sequence

AJWH01007149.1 Dictyostelium firmibasis strain TNS-C-0014 Contig15340, whole genome shotgun sequence

AJWH01007150.1 Dictyostelium firmibasis strain TNS-C-0014 Contig15341, whole genome shotgun sequence

JH724076.1 Dictyostelium firmibasis strain TNS-C-0014 unplaced genomic scaffold scaffold00645, whole genome shotgun sequence

AJWH01007153.1 Dictyostelium firmibasis strain TNS-C-0014 Contig15345, whole genome shotgun sequence

JH724077.1 Dictyostelium firmibasis strain TNS-C-0014 unplaced genomic scaffold scaffold00647, whole genome shotgun sequence

AJWH01007156.1 Dictyostelium firmibasis strain TNS-C-0014 Contig15348, whole genome shotgun sequence

AJWH01007157.1 Dictyostelium firmibasis strain TNS-C-0014 Contig15349, whole genome shotgun sequence

AJWH01007158.1 Dictyostelium firmibasis strain TNS-C-0014 Contig15350, whole genome shotgun sequence

AJWH01007159.1 Dictyostelium firmibasis strain TNS-C-0014 Contig15351, whole genome shotgun sequence

AJWH01007160.1 Dictyostelium firmibasis strain TNS-C-0014 Contig15352, whole genome shotgun sequence

AJWH01007161.1 Dictyostelium firmibasis strain TNS-C-0014 Contig15353, whole genome shotgun sequence

AJWH01007162.1 Dictyostelium firmibasis strain TNS-C-0014 Contig15354, whole genome shotgun sequence

AJWH01007163.1 Dictyostelium firmibasis strain TNS-C-0014 Contig15355, whole genome shotgun sequence

AJWH01007164.1 Dictyostelium firmibasis strain TNS-C-0014 Contig15356, whole genome shotgun sequence

AJWH01007165.1 Dictyostelium firmibasis strain TNS-C-0014 Contig15357, whole genome shotgun sequence

AJWH01007166.1 Dictyostelium firmibasis strain TNS-C-0014 Contig15358, whole genome shotgun sequence

AJWH01007167.1 Dictyostelium firmibasis strain TNS-C-0014 Contig15359, whole genome shotgun sequence

AJWH01007168.1 Dictyostelium firmibasis strain TNS-C-0014 Contig15360, whole genome shotgun sequence

AJWH01007169.1 Dictyostelium firmibasis strain TNS-C-0014 Contig15361, whole genome shotgun sequence

AJWH01007170.1 Dictyostelium firmibasis strain TNS-C-0014 Contig15362, whole genome shotgun sequence

AJWH01007171.1 Dictyostelium firmibasis strain TNS-C-0014 Contig15363, whole genome shotgun sequence

AJWH01007172.1 Dictyostelium firmibasis strain TNS-C-0014 Contig15364, whole genome shotgun sequence

AJWH01007173.1 Dictyostelium firmibasis strain TNS-C-0014 Contig15365, whole genome shotgun sequence

AJWH01007174.1 Dictyostelium firmibasis strain TNS-C-0014 Contig15366, whole genome shotgun sequence

AJWH01007175.1 Dictyostelium firmibasis strain TNS-C-0014 Contig15367, whole genome shotgun sequence

AJWH01007176.1 Dictyostelium firmibasis strain TNS-C-0014 Contig15368, whole genome shotgun sequence

AJWH01007177.1 Dictyostelium firmibasis strain TNS-C-0014 Contig15369, whole genome shotgun sequence

AJWH01007178.1 Dictyostelium firmibasis strain TNS-C-0014 Contig15370, whole genome shotgun sequence

AJWH01007179.1 Dictyostelium firmibasis strain TNS-C-0014 Contig15371, whole genome shotgun sequence

AJWH01007180.1 Dictyostelium firmibasis strain TNS-C-0014 Contig15372, whole genome shotgun sequence

AJWH01007181.1 Dictyostelium firmibasis strain TNS-C-0014 Contig15373, whole genome shotgun sequence

AJWH01007182.1 Dictyostelium firmibasis strain TNS-C-0014 Contig15374, whole genome shotgun sequence

AJWH01007183.1 Dictyostelium firmibasis strain TNS-C-0014 Contig15375, whole genome shotgun sequence

AJWH01007184.1 Dictyostelium firmibasis strain TNS-C-0014 Contig15376, whole genome shotgun sequence

AJWH01007185.1 Dictyostelium firmibasis strain TNS-C-0014 Contig15377, whole genome shotgun sequence

AJWH01007186.1 Dictyostelium firmibasis strain TNS-C-0014 Contig15378, whole genome shotgun sequence

AJWH01007187.1 Dictyostelium firmibasis strain TNS-C-0014 Contig15379, whole genome shotgun sequence

AJWH01007188.1 Dictyostelium firmibasis strain TNS-C-0014 Contig15380, whole genome shotgun sequence

JH724078.1 Dictyostelium firmibasis strain TNS-C-0014 unplaced genomic scaffold scaffold00681, whole genome shotgun sequence

AJWH01007191.1 Dictyostelium firmibasis strain TNS-C-0014 Contig15383, whole genome shotgun sequence

AJWH01007192.1 Dictyostelium firmibasis strain TNS-C-0014 Contig15384, whole genome shotgun sequence

AJWH01007193.1 Dictyostelium firmibasis strain TNS-C-0014 Contig15385, whole genome shotgun sequence

AJWH01007194.1 Dictyostelium firmibasis strain TNS-C-0014 Contig15386, whole genome shotgun sequence

AJWH01007195.1 Dictyostelium firmibasis strain TNS-C-0014 Contig15387, whole genome shotgun sequence

AJWH01007196.1 Dictyostelium firmibasis strain TNS-C-0014 Contig15388, whole genome shotgun sequence

AJWH01007197.1 Dictyostelium firmibasis strain TNS-C-0014 Contig15389, whole genome shotgun sequence

AJWH01007198.1 Dictyostelium firmibasis strain TNS-C-0014 Contig15390, whole genome shotgun sequence

AJWH01007199.1 Dictyostelium firmibasis strain TNS-C-0014 Contig15391, whole genome shotgun sequence

AJWH01007200.1 Dictyostelium firmibasis strain TNS-C-0014 Contig15392, whole genome shotgun sequence

AJWH01007201.1 Dictyostelium firmibasis strain TNS-C-0014 Contig15393, whole genome shotgun sequence

AJWH01007202.1 Dictyostelium firmibasis strain TNS-C-0014 Contig15394, whole genome shotgun sequence

AJWH01007203.1 Dictyostelium firmibasis strain TNS-C-0014 Contig15395, whole genome shotgun sequence

AJWH01007204.1 Dictyostelium firmibasis strain TNS-C-0014 Contig15396, whole genome shotgun sequence

AJWH01007205.1 Dictyostelium firmibasis strain TNS-C-0014 Contig15397, whole genome shotgun sequence

AJWH01007206.1 Dictyostelium firmibasis strain TNS-C-0014 Contig15398, whole genome shotgun sequence

AJWH01007207.1 Dictyostelium firmibasis strain TNS-C-0014 Contig15399, whole genome shotgun sequence

AJWH01007208.1 Dictyostelium firmibasis strain TNS-C-0014 Contig15400, whole genome shotgun sequence

AJWH01007209.1 Dictyostelium firmibasis strain TNS-C-0014 Contig15401, whole genome shotgun sequence

AJWH01007210.1 Dictyostelium firmibasis strain TNS-C-0014 Contig15402, whole genome shotgun sequence

AJWH01007211.1 Dictyostelium firmibasis strain TNS-C-0014 Contig15403, whole genome shotgun sequence

AJWH01007212.1 Dictyostelium firmibasis strain TNS-C-0014 Contig15404, whole genome shotgun sequence

AJWH01007213.1 Dictyostelium firmibasis strain TNS-C-0014 Contig15405, whole genome shotgun sequence

AJWH01007214.1 Dictyostelium firmibasis strain TNS-C-0014 Contig15406, whole genome shotgun sequence

AJWH01007215.1 Dictyostelium firmibasis strain TNS-C-0014 Contig15407, whole genome shotgun sequence

AJWH01007216.1 Dictyostelium firmibasis strain TNS-C-0014 Contig15409, whole genome shotgun sequence

AJWH01007217.1 Dictyostelium firmibasis strain TNS-C-0014 Contig15410, whole genome shotgun sequence

AJWH01007218.1 Dictyostelium firmibasis strain TNS-C-0014 Contig15411, whole genome shotgun sequence

AJWH01007219.1 Dictyostelium firmibasis strain TNS-C-0014 Contig15412, whole genome shotgun sequence

AJWH01007220.1 Dictyostelium firmibasis strain TNS-C-0014 Contig15413, whole genome shotgun sequence

AJWH01007221.1 Dictyostelium firmibasis strain TNS-C-0014 Contig15414, whole genome shotgun sequence

AJWH01007222.1 Dictyostelium firmibasis strain TNS-C-0014 Contig15415, whole genome shotgun sequence

AJWH01007223.1 Dictyostelium firmibasis strain TNS-C-0014 Contig15416, whole genome shotgun sequence

AJWH01007224.1 Dictyostelium firmibasis strain TNS-C-0014 Contig15417, whole genome shotgun sequence

AJWH01007225.1 Dictyostelium firmibasis strain TNS-C-0014 Contig15418, whole genome shotgun sequence

AJWH01007226.1 Dictyostelium firmibasis strain TNS-C-0014 Contig15419, whole genome shotgun sequence

AJWH01007227.1 Dictyostelium firmibasis strain TNS-C-0014 Contig15420, whole genome shotgun sequence

AJWH01007228.1 Dictyostelium firmibasis strain TNS-C-0014 Contig15421, whole genome shotgun sequence

AJWH01007229.1 Dictyostelium firmibasis strain TNS-C-0014 Contig15422, whole genome shotgun sequence

AJWH01007230.1 Dictyostelium firmibasis strain TNS-C-0014 Contig15423, whole genome shotgun sequence

AJWH01007231.1 Dictyostelium firmibasis strain TNS-C-0014 Contig15424, whole genome shotgun sequence

AJWH01007232.1 Dictyostelium firmibasis strain TNS-C-0014 Contig15425, whole genome shotgun sequence

AJWH01007233.1 Dictyostelium firmibasis strain TNS-C-0014 Contig15426, whole genome shotgun sequence

AJWH01007234.1 Dictyostelium firmibasis strain TNS-C-0014 Contig15427, whole genome shotgun sequence

AJWH01007235.1 Dictyostelium firmibasis strain TNS-C-0014 Contig15428, whole genome shotgun sequence

AJWH01007236.1 Dictyostelium firmibasis strain TNS-C-0014 Contig15430, whole genome shotgun sequence

AJWH01007237.1 Dictyostelium firmibasis strain TNS-C-0014 Contig15431, whole genome shotgun sequence

AJWH01007238.1 Dictyostelium firmibasis strain TNS-C-0014 Contig15432, whole genome shotgun sequence

AJWH01007239.1 Dictyostelium firmibasis strain TNS-C-0014 Contig15433, whole genome shotgun sequence

AJWH01007240.1 Dictyostelium firmibasis strain TNS-C-0014 Contig15434, whole genome shotgun sequence

AJWH01007241.1 Dictyostelium firmibasis strain TNS-C-0014 Contig15435, whole genome shotgun sequence

AJWH01007242.1 Dictyostelium firmibasis strain TNS-C-0014 Contig15436, whole genome shotgun sequence

AJWH01007243.1 Dictyostelium firmibasis strain TNS-C-0014 Contig15437, whole genome shotgun sequence

AJWH01007244.1 Dictyostelium firmibasis strain TNS-C-0014 Contig15438, whole genome shotgun sequence

AJWH01007245.1 Dictyostelium firmibasis strain TNS-C-0014 Contig15439, whole genome shotgun sequence

AJWH01007246.1 Dictyostelium firmibasis strain TNS-C-0014 Contig15440, whole genome shotgun sequence

AJWH01007247.1 Dictyostelium firmibasis strain TNS-C-0014 Contig15441, whole genome shotgun sequence

AJWH01007248.1 Dictyostelium firmibasis strain TNS-C-0014 Contig15442, whole genome shotgun sequence

AJWH01007249.1 Dictyostelium firmibasis strain TNS-C-0014 Contig15443, whole genome shotgun sequence

AJWH01007250.1 Dictyostelium firmibasis strain TNS-C-0014 Contig15444, whole genome shotgun sequence

AJWH01007251.1 Dictyostelium firmibasis strain TNS-C-0014 Contig15445, whole genome shotgun sequence

AJWH01007252.1 Dictyostelium firmibasis strain TNS-C-0014 Contig15446, whole genome shotgun sequence

AJWH01007253.1 Dictyostelium firmibasis strain TNS-C-0014 Contig15447, whole genome shotgun sequence

AJWH01007254.1 Dictyostelium firmibasis strain TNS-C-0014 Contig15448, whole genome shotgun sequence

AJWH01007255.1 Dictyostelium firmibasis strain TNS-C-0014 Contig15450, whole genome shotgun sequence

AJWH01007256.1 Dictyostelium firmibasis strain TNS-C-0014 Contig15451, whole genome shotgun sequence

AJWH01007257.1 Dictyostelium firmibasis strain TNS-C-0014 Contig15452, whole genome shotgun sequence

AJWH01007258.1 Dictyostelium firmibasis strain TNS-C-0014 Contig15453, whole genome shotgun sequence

AJWH01007259.1 Dictyostelium firmibasis strain TNS-C-0014 Contig15454, whole genome shotgun sequence

AJWH01007260.1 Dictyostelium firmibasis strain TNS-C-0014 Contig15455, whole genome shotgun sequence

AJWH01007261.1 Dictyostelium firmibasis strain TNS-C-0014 Contig15456, whole genome shotgun sequence

AJWH01007262.1 Dictyostelium firmibasis strain TNS-C-0014 Contig15457, whole genome shotgun sequence

AJWH01007263.1 Dictyostelium firmibasis strain TNS-C-0014 Contig15458, whole genome shotgun sequence

AJWH01007264.1 Dictyostelium firmibasis strain TNS-C-0014 Contig15459, whole genome shotgun sequence

AJWH01007265.1 Dictyostelium firmibasis strain TNS-C-0014 Contig15460, whole genome shotgun sequence

AJWH01007266.1 Dictyostelium firmibasis strain TNS-C-0014 Contig15461, whole genome shotgun sequence

AJWH01007267.1 Dictyostelium firmibasis strain TNS-C-0014 Contig15462, whole genome shotgun sequence

AJWH01007268.1 Dictyostelium firmibasis strain TNS-C-0014 Contig15463, whole genome shotgun sequence

AJWH01007269.1 Dictyostelium firmibasis strain TNS-C-0014 Contig15465, whole genome shotgun sequence

AJWH01007270.1 Dictyostelium firmibasis strain TNS-C-0014 Contig15466, whole genome shotgun sequence

AJWH01007271.1 Dictyostelium firmibasis strain TNS-C-0014 Contig15467, whole genome shotgun sequence

AJWH01007272.1 Dictyostelium firmibasis strain TNS-C-0014 Contig15468, whole genome shotgun sequence

AJWH01007273.1 Dictyostelium firmibasis strain TNS-C-0014 Contig15469, whole genome shotgun sequence

AJWH01007274.1 Dictyostelium firmibasis strain TNS-C-0014 Contig15470, whole genome shotgun sequence

AJWH01007275.1 Dictyostelium firmibasis strain TNS-C-0014 Contig15471, whole genome shotgun sequence

AJWH01007276.1 Dictyostelium firmibasis strain TNS-C-0014 Contig15472, whole genome shotgun sequence

AJWH01007277.1 Dictyostelium firmibasis strain TNS-C-0014 Contig15473, whole genome shotgun sequence

AJWH01007278.1 Dictyostelium firmibasis strain TNS-C-0014 Contig15474, whole genome shotgun sequence

AJWH01007279.1 Dictyostelium firmibasis strain TNS-C-0014 Contig15475, whole genome shotgun sequence

AJWH01007280.1 Dictyostelium firmibasis strain TNS-C-0014 Contig15476, whole genome shotgun sequence

AJWH01007281.1 Dictyostelium firmibasis strain TNS-C-0014 Contig15477, whole genome shotgun sequence

AJWH01007282.1 Dictyostelium firmibasis strain TNS-C-0014 Contig15478, whole genome shotgun sequence

AJWH01007283.1 Dictyostelium firmibasis strain TNS-C-0014 Contig15479, whole genome shotgun sequence

AJWH01007284.1 Dictyostelium firmibasis strain TNS-C-0014 Contig15480, whole genome shotgun sequence

AJWH01007285.1 Dictyostelium firmibasis strain TNS-C-0014 Contig15481, whole genome shotgun sequence

AJWH01007286.1 Dictyostelium firmibasis strain TNS-C-0014 Contig15483, whole genome shotgun sequence

AJWH01007287.1 Dictyostelium firmibasis strain TNS-C-0014 Contig15484, whole genome shotgun sequence

AJWH01007288.1 Dictyostelium firmibasis strain TNS-C-0014 Contig15485, whole genome shotgun sequence

AJWH01007289.1 Dictyostelium firmibasis strain TNS-C-0014 Contig15486, whole genome shotgun sequence

AJWH01007290.1 Dictyostelium firmibasis strain TNS-C-0014 Contig15487, whole genome shotgun sequence

AJWH01007291.1 Dictyostelium firmibasis strain TNS-C-0014 Contig15488, whole genome shotgun sequence

AJWH01007292.1 Dictyostelium firmibasis strain TNS-C-0014 Contig15489, whole genome shotgun sequence

AJWH01007293.1 Dictyostelium firmibasis strain TNS-C-0014 Contig15490, whole genome shotgun sequence

AJWH01007294.1 Dictyostelium firmibasis strain TNS-C-0014 Contig15491, whole genome shotgun sequence

AJWH01007295.1 Dictyostelium firmibasis strain TNS-C-0014 Contig15492, whole genome shotgun sequence

AJWH01007296.1 Dictyostelium firmibasis strain TNS-C-0014 Contig15493, whole genome shotgun sequence

AJWH01007297.1 Dictyostelium firmibasis strain TNS-C-0014 Contig15494, whole genome shotgun sequence

AJWH01007298.1 Dictyostelium firmibasis strain TNS-C-0014 Contig15495, whole genome shotgun sequence

AJWH01007299.1 Dictyostelium firmibasis strain TNS-C-0014 Contig15496, whole genome shotgun sequence

AJWH01007300.1 Dictyostelium firmibasis strain TNS-C-0014 Contig15497, whole genome shotgun sequence

AJWH01007301.1 Dictyostelium firmibasis strain TNS-C-0014 Contig15498, whole genome shotgun sequence

AJWH01007302.1 Dictyostelium firmibasis strain TNS-C-0014 Contig15499, whole genome shotgun sequence

AJWH01007303.1 Dictyostelium firmibasis strain TNS-C-0014 Contig15500, whole genome shotgun sequence

AJWH01007304.1 Dictyostelium firmibasis strain TNS-C-0014 Contig15502, whole genome shotgun sequence

AJWH01007305.1 Dictyostelium firmibasis strain TNS-C-0014 Contig15505, whole genome shotgun sequence

AJWH01007306.1 Dictyostelium firmibasis strain TNS-C-0014 Contig15506, whole genome shotgun sequence

AJWH01007307.1 Dictyostelium firmibasis strain TNS-C-0014 Contig15508, whole genome shotgun sequence

AJWH01007308.1 Dictyostelium firmibasis strain TNS-C-0014 Contig15509, whole genome shotgun sequence

AJWH01007309.1 Dictyostelium firmibasis strain TNS-C-0014 Contig15510, whole genome shotgun sequence

AJWH01000273.1 Dictyostelium firmibasis strain TNS-C-0014 Contig294, whole genome shotgun sequence

AJWH01000274.1 Dictyostelium firmibasis strain TNS-C-0014 Contig411, whole genome shotgun sequence

AJWH01000275.1 Dictyostelium firmibasis strain TNS-C-0014 Contig567, whole genome shotgun sequence

AJWH01000276.1 Dictyostelium firmibasis strain TNS-C-0014 Contig603, whole genome shotgun sequence

AJWH01000277.1 Dictyostelium firmibasis strain TNS-C-0014 Contig604, whole genome shotgun sequence

AJWH01000278.1 Dictyostelium firmibasis strain TNS-C-0014 Contig611, whole genome shotgun sequence

AJWH01000279.1 Dictyostelium firmibasis strain TNS-C-0014 Contig635, whole genome shotgun sequence

AJWH01000401.1 Dictyostelium firmibasis strain TNS-C-0014 Contig829, whole genome shotgun sequence

AJWH01000402.1 Dictyostelium firmibasis strain TNS-C-0014 Contig837, whole genome shotgun sequence

AJWH01000403.1 Dictyostelium firmibasis strain TNS-C-0014 Contig838, whole genome shotgun sequence

AJWH01000404.1 Dictyostelium firmibasis strain TNS-C-0014 Contig839, whole genome shotgun sequence

AJWH01000405.1 Dictyostelium firmibasis strain TNS-C-0014 Contig840, whole genome shotgun sequence

AJWH01000406.1 Dictyostelium firmibasis strain TNS-C-0014 Contig841, whole genome shotgun sequence

AJWH01000407.1 Dictyostelium firmibasis strain TNS-C-0014 Contig842, whole genome shotgun sequence

AJWH01000408.1 Dictyostelium firmibasis strain TNS-C-0014 Contig843, whole genome shotgun sequence

AJWH01000409.1 Dictyostelium firmibasis strain TNS-C-0014 Contig896, whole genome shotgun sequence

AJWH01000410.1 Dictyostelium firmibasis strain TNS-C-0014 Contig915, whole genome shotgun sequence

AJWH01000830.1 Dictyostelium firmibasis strain TNS-C-0014 Contig1469, whole genome shotgun sequence

AJWH01000831.1 Dictyostelium firmibasis strain TNS-C-0014 Contig1470, whole genome shotgun sequence

AJWH01000832.1 Dictyostelium firmibasis strain TNS-C-0014 Contig1471, whole genome shotgun sequence

AJWH01000833.1 Dictyostelium firmibasis strain TNS-C-0014 Contig1472, whole genome shotgun sequence

AJWH01000834.1 Dictyostelium firmibasis strain TNS-C-0014 Contig1473, whole genome shotgun sequence

AJWH01000835.1 Dictyostelium firmibasis strain TNS-C-0014 Contig1474, whole genome shotgun sequence

AJWH01000836.1 Dictyostelium firmibasis strain TNS-C-0014 Contig1475, whole genome shotgun sequence

AJWH01000837.1 Dictyostelium firmibasis strain TNS-C-0014 Contig1476, whole genome shotgun sequence

AJWH01000838.1 Dictyostelium firmibasis strain TNS-C-0014 Contig1477, whole genome shotgun sequence

AJWH01000839.1 Dictyostelium firmibasis strain TNS-C-0014 Contig1478, whole genome shotgun sequence

AJWH01000840.1 Dictyostelium firmibasis strain TNS-C-0014 Contig1479, whole genome shotgun sequence

AJWH01000841.1 Dictyostelium firmibasis strain TNS-C-0014 Contig1480, whole genome shotgun sequence

AJWH01000842.1 Dictyostelium firmibasis strain TNS-C-0014 Contig1481, whole genome shotgun sequence

AJWH01000843.1 Dictyostelium firmibasis strain TNS-C-0014 Contig1533, whole genome shotgun sequence

AJWH01000844.1 Dictyostelium firmibasis strain TNS-C-0014 Contig1534, whole genome shotgun sequence

AJWH01000845.1 Dictyostelium firmibasis strain TNS-C-0014 Contig1551, whole genome shotgun sequence

AJWH01000846.1 Dictyostelium firmibasis strain TNS-C-0014 Contig1563, whole genome shotgun sequence

AJWH01000847.1 Dictyostelium firmibasis strain TNS-C-0014 Contig1649, whole genome shotgun sequence

AJWH01000848.1 Dictyostelium firmibasis strain TNS-C-0014 Contig1659, whole genome shotgun sequence

AJWH01000849.1 Dictyostelium firmibasis strain TNS-C-0014 Contig1660, whole genome shotgun sequence

AJWH01000850.1 Dictyostelium firmibasis strain TNS-C-0014 Contig1661, whole genome shotgun sequence

AJWH01000851.1 Dictyostelium firmibasis strain TNS-C-0014 Contig1662, whole genome shotgun sequence

AJWH01000852.1 Dictyostelium firmibasis strain TNS-C-0014 Contig1663, whole genome shotgun sequence

AJWH01000853.1 Dictyostelium firmibasis strain TNS-C-0014 Contig1694, whole genome shotgun sequence

AJWH01000854.1 Dictyostelium firmibasis strain TNS-C-0014 Contig1729, whole genome shotgun sequence

AJWH01000855.1 Dictyostelium firmibasis strain TNS-C-0014 Contig1743, whole genome shotgun sequence

AJWH01000856.1 Dictyostelium firmibasis strain TNS-C-0014 Contig1761, whole genome shotgun sequence

AJWH01000857.1 Dictyostelium firmibasis strain TNS-C-0014 Contig1811, whole genome shotgun sequence

AJWH01000858.1 Dictyostelium firmibasis strain TNS-C-0014 Contig1814, whole genome shotgun sequence

AJWH01000859.1 Dictyostelium firmibasis strain TNS-C-0014 Contig1843, whole genome shotgun sequence

AJWH01000860.1 Dictyostelium firmibasis strain TNS-C-0014 Contig1879, whole genome shotgun sequence

AJWH01000861.1 Dictyostelium firmibasis strain TNS-C-0014 Contig1890, whole genome shotgun sequence

AJWH01000862.1 Dictyostelium firmibasis strain TNS-C-0014 Contig1897, whole genome shotgun sequence

AJWH01000863.1 Dictyostelium firmibasis strain TNS-C-0014 Contig1920, whole genome shotgun sequence

AJWH01000864.1 Dictyostelium firmibasis strain TNS-C-0014 Contig1959, whole genome shotgun sequence

AJWH01000959.1 Dictyostelium firmibasis strain TNS-C-0014 Contig2217, whole genome shotgun sequence

AJWH01000960.1 Dictyostelium firmibasis strain TNS-C-0014 Contig2256, whole genome shotgun sequence

AJWH01001137.1 Dictyostelium firmibasis strain TNS-C-0014 Contig2595, whole genome shotgun sequence

AJWH01001138.1 Dictyostelium firmibasis strain TNS-C-0014 Contig2615, whole genome shotgun sequence

AJWH01001139.1 Dictyostelium firmibasis strain TNS-C-0014 Contig2651, whole genome shotgun sequence

AJWH01001140.1 Dictyostelium firmibasis strain TNS-C-0014 Contig2669, whole genome shotgun sequence

AJWH01001641.1 Dictyostelium firmibasis strain TNS-C-0014 Contig3179, whole genome shotgun sequence

AJWH01001642.1 Dictyostelium firmibasis strain TNS-C-0014 Contig3229, whole genome shotgun sequence

AJWH01001643.1 Dictyostelium firmibasis strain TNS-C-0014 Contig3255, whole genome shotgun sequence

AJWH01001644.1 Dictyostelium firmibasis strain TNS-C-0014 Contig3319, whole genome shotgun sequence

AJWH01001645.1 Dictyostelium firmibasis strain TNS-C-0014 Contig3322, whole genome shotgun sequence

AJWH01001646.1 Dictyostelium firmibasis strain TNS-C-0014 Contig3323, whole genome shotgun sequence

AJWH01001647.1 Dictyostelium firmibasis strain TNS-C-0014 Contig3333, whole genome shotgun sequence

AJWH01001802.1 Dictyostelium firmibasis strain TNS-C-0014 Contig3592, whole genome shotgun sequence

AJWH01001803.1 Dictyostelium firmibasis strain TNS-C-0014 Contig3629, whole genome shotgun sequence

AJWH01002102.1 Dictyostelium firmibasis strain TNS-C-0014 Contig4013, whole genome shotgun sequence

AJWH01002103.1 Dictyostelium firmibasis strain TNS-C-0014 Contig4023, whole genome shotgun sequence

AJWH01002241.1 Dictyostelium firmibasis strain TNS-C-0014 Contig4237, whole genome shotgun sequence

AJWH01002242.1 Dictyostelium firmibasis strain TNS-C-0014 Contig4267, whole genome shotgun sequence

AJWH01002243.1 Dictyostelium firmibasis strain TNS-C-0014 Contig4270, whole genome shotgun sequence

AJWH01002244.1 Dictyostelium firmibasis strain TNS-C-0014 Contig4272, whole genome shotgun sequence

AJWH01002245.1 Dictyostelium firmibasis strain TNS-C-0014 Contig4273, whole genome shotgun sequence

AJWH01002246.1 Dictyostelium firmibasis strain TNS-C-0014 Contig4274, whole genome shotgun sequence

AJWH01002247.1 Dictyostelium firmibasis strain TNS-C-0014 Contig4284, whole genome shotgun sequence

AJWH01002248.1 Dictyostelium firmibasis strain TNS-C-0014 Contig4285, whole genome shotgun sequence

AJWH01002249.1 Dictyostelium firmibasis strain TNS-C-0014 Contig4286, whole genome shotgun sequence

AJWH01002250.1 Dictyostelium firmibasis strain TNS-C-0014 Contig4309, whole genome shotgun sequence

AJWH01002251.1 Dictyostelium firmibasis strain TNS-C-0014 Contig4311, whole genome shotgun sequence

AJWH01002252.1 Dictyostelium firmibasis strain TNS-C-0014 Contig4334, whole genome shotgun sequence

AJWH01002253.1 Dictyostelium firmibasis strain TNS-C-0014 Contig4335, whole genome shotgun sequence

AJWH01002254.1 Dictyostelium firmibasis strain TNS-C-0014 Contig4342, whole genome shotgun sequence

AJWH01002255.1 Dictyostelium firmibasis strain TNS-C-0014 Contig4351, whole genome shotgun sequence

AJWH01002256.1 Dictyostelium firmibasis strain TNS-C-0014 Contig4385, whole genome shotgun sequence

AJWH01002257.1 Dictyostelium firmibasis strain TNS-C-0014 Contig4405, whole genome shotgun sequence

AJWH01002258.1 Dictyostelium firmibasis strain TNS-C-0014 Contig4456, whole genome shotgun sequence

AJWH01002259.1 Dictyostelium firmibasis strain TNS-C-0014 Contig4488, whole genome shotgun sequence

AJWH01002325.1 Dictyostelium firmibasis strain TNS-C-0014 Contig4650, whole genome shotgun sequence

AJWH01002326.1 Dictyostelium firmibasis strain TNS-C-0014 Contig4675, whole genome shotgun sequence

AJWH01002327.1 Dictyostelium firmibasis strain TNS-C-0014 Contig4846, whole genome shotgun sequence

AJWH01002328.1 Dictyostelium firmibasis strain TNS-C-0014 Contig4883, whole genome shotgun sequence

AJWH01002329.1 Dictyostelium firmibasis strain TNS-C-0014 Contig4889, whole genome shotgun sequence

AJWH01002330.1 Dictyostelium firmibasis strain TNS-C-0014 Contig4916, whole genome shotgun sequence

AJWH01002331.1 Dictyostelium firmibasis strain TNS-C-0014 Contig4939, whole genome shotgun sequence

AJWH01002332.1 Dictyostelium firmibasis strain TNS-C-0014 Contig4955, whole genome shotgun sequence

AJWH01002333.1 Dictyostelium firmibasis strain TNS-C-0014 Contig4956, whole genome shotgun sequence

AJWH01002334.1 Dictyostelium firmibasis strain TNS-C-0014 Contig4968, whole genome shotgun sequence

AJWH01002394.1 Dictyostelium firmibasis strain TNS-C-0014 Contig5053, whole genome shotgun sequence

AJWH01002395.1 Dictyostelium firmibasis strain TNS-C-0014 Contig5060, whole genome shotgun sequence

AJWH01002396.1 Dictyostelium firmibasis strain TNS-C-0014 Contig5070, whole genome shotgun sequence

AJWH01002607.1 Dictyostelium firmibasis strain TNS-C-0014 Contig5451, whole genome shotgun sequence

AJWH01002608.1 Dictyostelium firmibasis strain TNS-C-0014 Contig5476, whole genome shotgun sequence

AJWH01002609.1 Dictyostelium firmibasis strain TNS-C-0014 Contig5501, whole genome shotgun sequence

AJWH01002610.1 Dictyostelium firmibasis strain TNS-C-0014 Contig5561, whole genome shotgun sequence

AJWH01002611.1 Dictyostelium firmibasis strain TNS-C-0014 Contig5562, whole genome shotgun sequence

AJWH01002612.1 Dictyostelium firmibasis strain TNS-C-0014 Contig5582, whole genome shotgun sequence

AJWH01002613.1 Dictyostelium firmibasis strain TNS-C-0014 Contig5608, whole genome shotgun sequence

AJWH01002614.1 Dictyostelium firmibasis strain TNS-C-0014 Contig5661, whole genome shotgun sequence

AJWH01002615.1 Dictyostelium firmibasis strain TNS-C-0014 Contig5669, whole genome shotgun sequence

AJWH01003052.1 Dictyostelium firmibasis strain TNS-C-0014 Contig6517, whole genome shotgun sequence

AJWH01003053.1 Dictyostelium firmibasis strain TNS-C-0014 Contig6518, whole genome shotgun sequence

AJWH01003054.1 Dictyostelium firmibasis strain TNS-C-0014 Contig6529, whole genome shotgun sequence

AJWH01003055.1 Dictyostelium firmibasis strain TNS-C-0014 Contig6551, whole genome shotgun sequence

AJWH01003056.1 Dictyostelium firmibasis strain TNS-C-0014 Contig6552, whole genome shotgun sequence

AJWH01003408.1 Dictyostelium firmibasis strain TNS-C-0014 Contig7118, whole genome shotgun sequence

AJWH01003409.1 Dictyostelium firmibasis strain TNS-C-0014 Contig7119, whole genome shotgun sequence

AJWH01003410.1 Dictyostelium firmibasis strain TNS-C-0014 Contig7122, whole genome shotgun sequence

AJWH01003411.1 Dictyostelium firmibasis strain TNS-C-0014 Contig7142, whole genome shotgun sequence

AJWH01003412.1 Dictyostelium firmibasis strain TNS-C-0014 Contig7163, whole genome shotgun sequence

AJWH01003413.1 Dictyostelium firmibasis strain TNS-C-0014 Contig7174, whole genome shotgun sequence

AJWH01003414.1 Dictyostelium firmibasis strain TNS-C-0014 Contig7198, whole genome shotgun sequence

AJWH01003465.1 Dictyostelium firmibasis strain TNS-C-0014 Contig7262, whole genome shotgun sequence

AJWH01003466.1 Dictyostelium firmibasis strain TNS-C-0014 Contig7263, whole genome shotgun sequence

AJWH01003467.1 Dictyostelium firmibasis strain TNS-C-0014 Contig7273, whole genome shotgun sequence

AJWH01003468.1 Dictyostelium firmibasis strain TNS-C-0014 Contig7285, whole genome shotgun sequence

AJWH01003816.1 Dictyostelium firmibasis strain TNS-C-0014 Contig7849, whole genome shotgun sequence

AJWH01003817.1 Dictyostelium firmibasis strain TNS-C-0014 Contig7850, whole genome shotgun sequence

AJWH01003818.1 Dictyostelium firmibasis strain TNS-C-0014 Contig7851, whole genome shotgun sequence

AJWH01003819.1 Dictyostelium firmibasis strain TNS-C-0014 Contig7863, whole genome shotgun sequence

AJWH01003820.1 Dictyostelium firmibasis strain TNS-C-0014 Contig7868, whole genome shotgun sequence

AJWH01004244.1 Dictyostelium firmibasis strain TNS-C-0014 Contig8829, whole genome shotgun sequence

AJWH01004245.1 Dictyostelium firmibasis strain TNS-C-0014 Contig8831, whole genome shotgun sequence

AJWH01004246.1 Dictyostelium firmibasis strain TNS-C-0014 Contig8845, whole genome shotgun sequence

AJWH01004247.1 Dictyostelium firmibasis strain TNS-C-0014 Contig8846, whole genome shotgun sequence

AJWH01004248.1 Dictyostelium firmibasis strain TNS-C-0014 Contig8847, whole genome shotgun sequence

AJWH01004249.1 Dictyostelium firmibasis strain TNS-C-0014 Contig8850, whole genome shotgun sequence

AJWH01004250.1 Dictyostelium firmibasis strain TNS-C-0014 Contig8851, whole genome shotgun sequence

AJWH01004251.1 Dictyostelium firmibasis strain TNS-C-0014 Contig8867, whole genome shotgun sequence

AJWH01004252.1 Dictyostelium firmibasis strain TNS-C-0014 Contig8871, whole genome shotgun sequence

AJWH01004386.1 Dictyostelium firmibasis strain TNS-C-0014 Contig9264, whole genome shotgun sequence

AJWH01004387.1 Dictyostelium firmibasis strain TNS-C-0014 Contig9280, whole genome shotgun sequence

AJWH01004460.1 Dictyostelium firmibasis strain TNS-C-0014 Contig9393, whole genome shotgun sequence

AJWH01004461.1 Dictyostelium firmibasis strain TNS-C-0014 Contig9402, whole genome shotgun sequence

AJWH01004462.1 Dictyostelium firmibasis strain TNS-C-0014 Contig9405, whole genome shotgun sequence

AJWH01004498.1 Dictyostelium firmibasis strain TNS-C-0014 Contig9595, whole genome shotgun sequence

AJWH01004499.1 Dictyostelium firmibasis strain TNS-C-0014 Contig9596, whole genome shotgun sequence

AJWH01004500.1 Dictyostelium firmibasis strain TNS-C-0014 Contig9605, whole genome shotgun sequence

AJWH01004501.1 Dictyostelium firmibasis strain TNS-C-0014 Contig9609, whole genome shotgun sequence

AJWH01004502.1 Dictyostelium firmibasis strain TNS-C-0014 Contig9628, whole genome shotgun sequence

AJWH01004776.1 Dictyostelium firmibasis strain TNS-C-0014 Contig10166, whole genome shotgun sequence

AJWH01004777.1 Dictyostelium firmibasis strain TNS-C-0014 Contig10180, whole genome shotgun sequence

AJWH01004778.1 Dictyostelium firmibasis strain TNS-C-0014 Contig10189, whole genome shotgun sequence

AJWH01004943.1 Dictyostelium firmibasis strain TNS-C-0014 Contig10563, whole genome shotgun sequence

AJWH01004944.1 Dictyostelium firmibasis strain TNS-C-0014 Contig10564, whole genome shotgun sequence

AJWH01004945.1 Dictyostelium firmibasis strain TNS-C-0014 Contig10579, whole genome shotgun sequence

AJWH01005047.1 Dictyostelium firmibasis strain TNS-C-0014 Contig10860, whole genome shotgun sequence

AJWH01005048.1 Dictyostelium firmibasis strain TNS-C-0014 Contig10861, whole genome shotgun sequence

AJWH01005049.1 Dictyostelium firmibasis strain TNS-C-0014 Contig10862, whole genome shotgun sequence

AJWH01005050.1 Dictyostelium firmibasis strain TNS-C-0014 Contig10863, whole genome shotgun sequence

AJWH01005051.1 Dictyostelium firmibasis strain TNS-C-0014 Contig10874, whole genome shotgun sequence

AJWH01005078.1 Dictyostelium firmibasis strain TNS-C-0014 Contig10994, whole genome shotgun sequence

AJWH01005079.1 Dictyostelium firmibasis strain TNS-C-0014 Contig10996, whole genome shotgun sequence

AJWH01005080.1 Dictyostelium firmibasis strain TNS-C-0014 Contig10999, whole genome shotgun sequence

AJWH01005082.1 Dictyostelium firmibasis strain TNS-C-0014 Contig11045, whole genome shotgun sequence

AJWH01005083.1 Dictyostelium firmibasis strain TNS-C-0014 Contig11048, whole genome shotgun sequence

AJWH01005401.1 Dictyostelium firmibasis strain TNS-C-0014 Contig11528, whole genome shotgun sequence

AJWH01005402.1 Dictyostelium firmibasis strain TNS-C-0014 Contig11531, whole genome shotgun sequence

AJWH01005403.1 Dictyostelium firmibasis strain TNS-C-0014 Contig11535, whole genome shotgun sequence

AJWH01005404.1 Dictyostelium firmibasis strain TNS-C-0014 Contig11539, whole genome shotgun sequence

AJWH01005560.1 Dictyostelium firmibasis strain TNS-C-0014 Contig12083, whole genome shotgun sequence

AJWH01005561.1 Dictyostelium firmibasis strain TNS-C-0014 Contig12085, whole genome shotgun sequence

AJWH01005596.1 Dictyostelium firmibasis strain TNS-C-0014 Contig12176, whole genome shotgun sequence

AJWH01005597.1 Dictyostelium firmibasis strain TNS-C-0014 Contig12196, whole genome shotgun sequence

AJWH01005619.1 Dictyostelium firmibasis strain TNS-C-0014 Contig12219, whole genome shotgun sequence

AJWH01005620.1 Dictyostelium firmibasis strain TNS-C-0014 Contig12227, whole genome shotgun sequence

AJWH01005904.1 Dictyostelium firmibasis strain TNS-C-0014 Contig12981, whole genome shotgun sequence

AJWH01005905.1 Dictyostelium firmibasis strain TNS-C-0014 Contig12998, whole genome shotgun sequence

AJWH01006064.1 Dictyostelium firmibasis strain TNS-C-0014 Contig13522, whole genome shotgun sequence

AJWH01006065.1 Dictyostelium firmibasis strain TNS-C-0014 Contig13524, whole genome shotgun sequence

AJWH01006066.1 Dictyostelium firmibasis strain TNS-C-0014 Contig13525, whole genome shotgun sequence

AJWH01006067.1 Dictyostelium firmibasis strain TNS-C-0014 Contig13526, whole genome shotgun sequence

AJWH01006199.1 Dictyostelium firmibasis strain TNS-C-0014 Contig13762, whole genome shotgun sequence

AJWH01006200.1 Dictyostelium firmibasis strain TNS-C-0014 Contig13767, whole genome shotgun sequence

AJWH01006201.1 Dictyostelium firmibasis strain TNS-C-0014 Contig13768, whole genome shotgun sequence

AJWH01006247.1 Dictyostelium firmibasis strain TNS-C-0014 Contig13832, whole genome shotgun sequence

AJWH01006248.1 Dictyostelium firmibasis strain TNS-C-0014 Contig13833, whole genome shotgun sequence

AJWH01006249.1 Dictyostelium firmibasis strain TNS-C-0014 Contig13834, whole genome shotgun sequence

AJWH01006250.1 Dictyostelium firmibasis strain TNS-C-0014 Contig13838, whole genome shotgun sequence

AJWH01006331.1 Dictyostelium firmibasis strain TNS-C-0014 Contig14034, whole genome shotgun sequence

AJWH01006332.1 Dictyostelium firmibasis strain TNS-C-0014 Contig14044, whole genome shotgun sequence

AJWH01006430.1 Dictyostelium firmibasis strain TNS-C-0014 Contig14211, whole genome shotgun sequence

AJWH01006431.1 Dictyostelium firmibasis strain TNS-C-0014 Contig14214, whole genome shotgun sequence

AJWH01006535.1 Dictyostelium firmibasis strain TNS-C-0014 Contig14483, whole genome shotgun sequence

AJWH01006536.1 Dictyostelium firmibasis strain TNS-C-0014 Contig14485, whole genome shotgun sequence

AJWH01006720.1 Dictyostelium firmibasis strain TNS-C-0014 Contig14832, whole genome shotgun sequence

AJWH01006721.1 Dictyostelium firmibasis strain TNS-C-0014 Contig14836, whole genome shotgun sequence

AP023109.1 Entamoeba histolytica HM-1:IMSS Clone 6 2001 DNA, chromosome 1, nearly complete sequence

AP023110.1 Entamoeba histolytica HM-1:IMSS Clone 6 2001 DNA, chromosome 2, nearly complete sequence

AP023111.1 Entamoeba histolytica HM-1:IMSS Clone 6 2001 DNA, chromosome 3, nearly complete sequence

AP023112.1 Entamoeba histolytica HM-1:IMSS Clone 6 2001 DNA, chromosome 39, nearly complete sequence

AP023113.1 Entamoeba histolytica HM-1:IMSS Clone 6 2001 DNA, chromosome 5, nearly complete sequence

AP023114.1 Entamoeba histolytica HM-1:IMSS Clone 6 2001 DNA, chromosome 6, nearly complete sequence

AP023115.1 Entamoeba histolytica HM-1:IMSS Clone 6 2001 DNA, chromosome 7, nearly complete sequence

AP023116.1 Entamoeba histolytica HM-1:IMSS Clone 6 2001 DNA, chromosome 8, nearly complete sequence

AP023117.1 Entamoeba histolytica HM-1:IMSS Clone 6 2001 DNA, chromosome 9, nearly complete sequence

AP023118.1 Entamoeba histolytica HM-1:IMSS Clone 6 2001 DNA, chromosome 10, nearly complete sequence

AP023119.1 Entamoeba histolytica HM-1:IMSS Clone 6 2001 DNA, chromosome 11, nearly complete sequence

AP023120.1 Entamoeba histolytica HM-1:IMSS Clone 6 2001 DNA, chromosome 12, nearly complete sequence

AP023121.1 Entamoeba histolytica HM-1:IMSS Clone 6 2001 DNA, chromosome 13, nearly complete sequence

AP023122.1 Entamoeba histolytica HM-1:IMSS Clone 6 2001 DNA, chromosome 14, nearly complete sequence

AP023123.1 Entamoeba histolytica HM-1:IMSS Clone 6 2001 DNA, chromosome 15, nearly complete sequence

AP023124.1 Entamoeba histolytica HM-1:IMSS Clone 6 2001 DNA, chromosome 16, nearly complete sequence

AP023125.1 Entamoeba histolytica HM-1:IMSS Clone 6 2001 DNA, chromosome 17, nearly complete sequence

AP023126.1 Entamoeba histolytica HM-1:IMSS Clone 6 2001 DNA, chromosome 18, nearly complete sequence

AP023127.1 Entamoeba histolytica HM-1:IMSS Clone 6 2001 DNA, chromosome 19, nearly complete sequence

AP023128.1 Entamoeba histolytica HM-1:IMSS Clone 6 2001 DNA, chromosome 20, nearly complete sequence

AP023129.1 Entamoeba histolytica HM-1:IMSS Clone 6 2001 DNA, chromosome 21, nearly complete sequence

AP023130.1 Entamoeba histolytica HM-1:IMSS Clone 6 2001 DNA, chromosome 22, nearly complete sequence

AP023131.1 Entamoeba histolytica HM-1:IMSS Clone 6 2001 DNA, chromosome 23, nearly complete sequence

AP023132.1 Entamoeba histolytica HM-1:IMSS Clone 6 2001 DNA, chromosome 24, nearly complete sequence

AP023133.1 Entamoeba histolytica HM-1:IMSS Clone 6 2001 DNA, chromosome 25, nearly complete sequence

AP023134.1 Entamoeba histolytica HM-1:IMSS Clone 6 2001 DNA, chromosome 26, nearly complete sequence

AP023135.1 Entamoeba histolytica HM-1:IMSS Clone 6 2001 DNA, chromosome 27, nearly complete sequence

AP023136.1 Entamoeba histolytica HM-1:IMSS Clone 6 2001 DNA, chromosome 28, nearly complete sequence

AP023137.1 Entamoeba histolytica HM-1:IMSS Clone 6 2001 DNA, chromosome 29, nearly complete sequence

AP023138.1 Entamoeba histolytica HM-1:IMSS Clone 6 2001 DNA, chromosome 30, nearly complete sequence

AP023139.1 Entamoeba histolytica HM-1:IMSS Clone 6 2001 DNA, chromosome 31, nearly complete sequence

AP023140.1 Entamoeba histolytica HM-1:IMSS Clone 6 2001 DNA, chromosome 32, nearly complete sequence

AP023141.1 Entamoeba histolytica HM-1:IMSS Clone 6 2001 DNA, chromosome 33, nearly complete sequence

AP023142.1 Entamoeba histolytica HM-1:IMSS Clone 6 2001 DNA, chromosome 34, nearly complete sequence

AP023143.1 Entamoeba histolytica HM-1:IMSS Clone 6 2001 DNA, chromosome 35, nearly complete sequence

AP023144.1 Entamoeba histolytica HM-1:IMSS Clone 6 2001 DNA, chromosome 36, nearly complete sequence

AP023145.1 Entamoeba histolytica HM-1:IMSS Clone 6 2001 DNA, chromosome 37, nearly complete sequence

AP023146.1 Entamoeba histolytica HM-1:IMSS Clone 6 2001 DNA, chromosome 38, nearly complete sequence

AP023147.1 Entamoeba histolytica HM-1:IMSS Clone 6 2001 plasmid pEhCl6-2001 DNA, complete sequence

CM069765.1 Dictyostelium firmibasis strain TNS-C-14 chromosome 1, whole genome shotgun sequence

CM069766.1 Dictyostelium firmibasis strain TNS-C-14 chromosome 2, whole genome shotgun sequence

CM069767.1 Dictyostelium firmibasis strain TNS-C-14 chromosome 3, whole genome shotgun sequence

CM069768.1 Dictyostelium firmibasis strain TNS-C-14 chromosome 4, whole genome shotgun sequence

CM069769.1 Dictyostelium firmibasis strain TNS-C-14 chromosome 5, whole genome shotgun sequence

CM069770.1 Dictyostelium firmibasis strain TNS-C-14 chromosome 6, whole genome shotgun sequence

JAVFKY010000007.1 Dictyostelium firmibasis strain TNS-C-14 contig_33_np1212, whole genome shotgun sequence

JAVFKY010000008.1 Dictyostelium firmibasis strain TNS-C-14 contig_23_np1212, whole genome shotgun sequence

JAVFKY010000010.1 Dictyostelium firmibasis strain TNS-C-14 contig_9_np1212, whole genome shotgun sequence

JAVFKY010000011.1 Dictyostelium firmibasis strain TNS-C-14 contig_16_np1212, whole genome shotgun sequence

JAVFKY010000012.1 Dictyostelium firmibasis strain TNS-C-14 contig_63_np1212, whole genome shotgun sequence

CM069771.1 Dictyostelium firmibasis strain TNS-C-14 mitochondrion, complete sequence, whole genome shotgun sequence

NC_007087.3 Dictyostelium discoideum AX4 chromosome 1 chromosome, whole genome shotgun sequence

NC_007088.5 Dictyostelium discoideum AX4 chromosome 2 chromosome, whole genome shotgun sequence

NC_007089.4 Dictyostelium discoideum AX4 chromosome 3 chromosome, whole genome shotgun sequence

NC_007090.3 Dictyostelium discoideum AX4 chromosome 4 chromosome, whole genome shotgun sequence

NC_007091.3 Dictyostelium discoideum AX4 chromosome 5 chromosome, whole genome shotgun sequence

NC_007092.3 Dictyostelium discoideum AX4 chromosome 6 chromosome, whole genome shotgun sequence

NC_001889.1 Dictyostelium discoideum WS2162 plasmid Ddp5, complete sequence

NW_003102053.1 Dictyostelium discoideum AX4 chromosome Un chrUn_00010, whole genome shotgun sequence

NW_003102052.1 Dictyostelium discoideum AX4 chromosome Un chrUn_00011, whole genome shotgun sequence

NW_003102051.1 Dictyostelium discoideum AX4 chromosome Un chrUn_00012, whole genome shotgun sequence

NW_003102050.1 Dictyostelium discoideum AX4 chromosome Un chrUn_00013, whole genome shotgun sequence

NW_003102049.1 Dictyostelium discoideum AX4 chromosome Un chrUn_00014, whole genome shotgun sequence

NW_003102048.1 Dictyostelium discoideum AX4 chromosome Un chrUn_00015, whole genome shotgun sequence

NW_003102047.1 Dictyostelium discoideum AX4 chromosome Un chrUn_00016, whole genome shotgun sequence

NW_003102046.1 Dictyostelium discoideum AX4 chromosome Un chrUn_00018, whole genome shotgun sequence

NW_003102061.1 Dictyostelium discoideum AX4 chromosome Un chrUn_0002, whole genome shotgun sequence

NW_003102045.1 Dictyostelium discoideum AX4 chromosome Un chrUn_00020, whole genome shotgun sequence

NW_003102044.1 Dictyostelium discoideum AX4 chromosome Un chrUn_00021, whole genome shotgun sequence

NW_003102043.1 Dictyostelium discoideum AX4 chromosome Un chrUn_00022, whole genome shotgun sequence

NW_003102042.1 Dictyostelium discoideum AX4 chromosome Un chrUn_00023, whole genome shotgun sequence

NW_003102041.1 Dictyostelium discoideum AX4 chromosome Un chrUn_00024, whole genome shotgun sequence

NW_003102040.1 Dictyostelium discoideum AX4 chromosome Un chrUn_00025, whole genome shotgun sequence

NW_003102039.1 Dictyostelium discoideum AX4 chromosome Un chrUn_00026, whole genome shotgun sequence

NW_003102038.1 Dictyostelium discoideum AX4 chrUn_00027 genomic scaffold, whole genome shotgun sequence

NW_003102037.1 Dictyostelium discoideum AX4 chromosome Un chrUn_00028, whole genome shotgun sequence

NW_003102036.1 Dictyostelium discoideum AX4 chromosome Un chrUn_00029, whole genome shotgun sequence

NW_003102060.1 Dictyostelium discoideum AX4 chromosome Un chrUn_0003, whole genome shotgun sequence

NW_003102035.1 Dictyostelium discoideum AX4 chrUn_00030 genomic scaffold, whole genome shotgun sequence

NW_003102034.1 Dictyostelium discoideum AX4 chromosome Un chrUn_00031, whole genome shotgun sequence

NW_003102033.1 Dictyostelium discoideum AX4 chromosome Un chrUn_00032, whole genome shotgun sequence

NW_003102032.1 Dictyostelium discoideum AX4 chromosome Un chrUn_00033, whole genome shotgun sequence

NW_003102031.1 Dictyostelium discoideum AX4 chromosome Un chrUn_00034, whole genome shotgun sequence

NW_003102030.1 Dictyostelium discoideum AX4 chromosome Un chrUn_00035, whole genome shotgun sequence

NW_003102029.1 Dictyostelium discoideum AX4 chromosome Un chrUn_00036, whole genome shotgun sequence

NW_003102059.1 Dictyostelium discoideum AX4 chromosome Un chrUn_0004, whole genome shotgun sequence

NW_003102058.1 Dictyostelium discoideum AX4 chromosome Un chrUn_0005, whole genome shotgun sequence

NW_003102057.1 Dictyostelium discoideum AX4 chromosome Un chrUn_0006, whole genome shotgun sequence

NW_003102056.1 Dictyostelium discoideum AX4 chromosome Un chrUn_0007, whole genome shotgun sequence

NW_003102055.1 Dictyostelium discoideum AX4 chromosome Un chrUn_0008, whole genome shotgun sequence

NW_003102054.1 Dictyostelium discoideum AX4 chromosome Un chrUn_0009, whole genome shotgun sequence

NC_000895.1 Dictyostelium discoideum mitochondrion, complete genome
