## Supplementary Data 3 for "Sampling Microbial Dynamics in the Salish Sea Estuary: Evaluating Methods to Capture Cyanobacteria and Cyanophage"

| Parent ID | Date of Extraction | Date of Sampling | Time in Freezer (Day | KitName |
|---|---|---|---|---|
| NW_FE_003 | 3/26/2024 | 3/12/2024 | 14 | PSK |
| NW_FE_003 | 3/26/2024 | 3/12/2024 | 14 | PSK |
| NW_FE_003 | 3/26/2024 | 3/12/2024 | 14 | PSK |
| NW_FE_004 | 3/26/2024 | 3/13/2024 | 13 | PSK |
| NW_FE_004 | 3/26/2024 | 3/13/2024 | 13 | PSK |
| NW_FE_004 | 3/26/2024 | 3/13/2024 | 13 | PSK |
| NW_FE_004 | 3/26/2024 | 3/13/2024 | 13 | PSK |
| NW_FE_005 | 4/18/2024 | 4/15/2024 | 3 | PSK |
| NW_FE_005 | 4/18/2024 | 4/15/2024 | 3 | PSK |
| NW_FE_005 | 4/18/2024 | 4/15/2024 | 3 | PSK |
| NW_FE_005 | 4/18/2024 | 4/15/2024 | 3 | PSK |
| NW_FE_005 | 4/18/2024 | 4/15/2024 | 3 | PSK |
| NW_FE_005 | 4/18/2024 | 4/15/2024 | 3 | PSK |
| NW_FE_005 | 4/18/2024 | 4/15/2024 | 3 | PSK |
| NW_FE_005 | 4/18/2024 | 4/15/2024 | 3 | PSK |
| NW_FE_005 | 4/18/2024 | 4/15/2024 | 3 | PSK |
| NW_FE_005 | 4/18/2024 | 4/15/2024 | 3 | PSK |
| NW_FE_005 | 4/18/2024 | 4/15/2024 | 3 | PSK |
| NW_FE_005 | 4/18/2024 | 4/15/2024 | 3 | PSK |
| NW_FE_005 | 4/18/2024 | 4/15/2024 | 3 | PSK |
| NW_FE_005 | 4/18/2024 | 4/15/2024 | 3 | ZPK |
| NW_FE_005 | 4/18/2024 | 4/15/2024 | 3 | ZPK |
| NW_FE_005 | 4/18/2024 | 4/15/2024 | 3 | ZPK |
| NW_FE_005 | 4/18/2024 | 4/15/2024 | 3 | ZPK |
| NW_FE_005 | 4/18/2024 | 4/15/2024 | 3 | ZPK |
| NW_FE_005 | 4/18/2024 | 4/15/2024 | 3 | ZPK |
| NW_FE_005 | 4/18/2024 | 4/15/2024 | 3 | ZPK |
| NW_FE_005 | 4/18/2024 | 4/15/2024 | 3 | ZPK |
| NW_FE_005 | 4/18/2024 | 4/15/2024 | 3 | ZPK |
| NW_FE_005 | 4/18/2024 | 4/15/2024 | 3 | ZPK |
| NW_FE_005 | 4/18/2024 | 4/15/2024 | 3 | ZPK |
| NW_FE_005 | 4/18/2024 | 4/15/2024 | 3 | ZPK |
| NW_FE_005 | 4/18/2024 | 4/15/2024 | 3 | ZPK |
| NW_FE_005 | 4/18/2024 | 4/15/2024 | 3 | ZPK |
| NW_FE_005 | 4/18/2024 | 4/15/2024 | 3 | PSK |
| NW_FE_006 | 5/13/2024 | 5/8/2024 | 5 | PSK |
| NW_FE_006 | 5/13/2024 | 5/8/2024 | 5 | PSK |
| NW_FE_006 | 5/13/2024 | 5/8/2024 | 5 | PSK |
| NW_FE_006 | 5/13/2024 | 5/8/2024 | 5 | PSK |
| NW_FE_006 | 5/13/2024 | 5/8/2024 | 5 | PSK |
| NW_FE_006 | 5/13/2024 | 5/8/2024 | 5 | PSK |
| NW_FE_006 | 5/13/2024 | 5/8/2024 | 5 | PSK |
| NW_FE_006 | 5/13/2024 | 5/8/2024 | 5 | PSK |
| NW_FE_006 | 5/13/2024 | 5/8/2024 | 5 | PSK |
| NW_FE_006 | 5/13/2024 | 5/8/2024 | 5 | PSK |
| NW_FE_006 | 5/13/2024 | 5/8/2024 | 5 | PSK |

| | | | | |
|---|---|---|---|---|
| NW_FE_006 | 5/13/2024 | 5/8/2024 | 5 | PSK |
| NW_FE_006 | 5/13/2024 | 5/8/2024 | 5 | PSK |
| NW_FE_010 | 7/18/2024 | 7/10/2024 | 8 | PSK |
| NW_FE_010 | 7/18/2024 | 7/10/2024 | 8 | PSK |
| NW_FE_010 | 7/18/2024 | 7/10/2024 | 8 | PSK |
| NW_FE_011 | 7/18/2024 | 7/17/2024 | 1 | PSK |
| NW_FE_011 | 7/18/2024 | 7/17/2024 | 1 | PSK |
| NW_FE_008 | 7/19/2024 | 6/28/2024 | 21 | PSK |
| NW_FE_008 | 7/19/2024 | 6/28/2024 | 21 | PSK |
| NW_FE_009 | 7/19/2024 | 7/3/2024 | 16 | PSK |
| NW_FE_010 | 7/19/2024 | 7/10/2024 | 9 | PSK |
| NW_FE_010 | 7/19/2024 | 7/10/2024 | 9 | PSK |
| NW_FE_007 | 7/24/2024 | 6/19/2024 | 35 | PSK |
| NW_FE_007 | 7/24/2024 | 6/19/2024 | 35 | PSK |
| NW_FE_007 | 7/24/2024 | 6/19/2024 | 35 | PSK |
| NW_FE_007 | 7/25/2024 | 6/19/2024 | 36 | BTK |
| NW_FE_007 | 7/25/2024 | 6/19/2024 | 36 | BTK |
| NW_FE_007 | 7/25/2024 | 6/19/2024 | 36 | BTK |
| NW_FE_012 | 8/5/2024 | 7/23/2024 | 13 | PSK |
| NW_FE_012 | 8/5/2024 | 7/23/2024 | 13 | PSK |
| NW_FE_013 | 8/5/2024 | 7/31/2024 | 5 | PSK |
| NW_FE_013 | 8/5/2024 | 7/31/2024 | 5 | PSK |
| NW_FE_013 | 8/5/2024 | 7/31/2024 | 5 | PSK |
| NW_FE_013 | 8/5/2024 | 7/31/2024 | 5 | PSK |
| NW_FE_014 | 8/5/2024 | 7/31/2024 | 5 | PSK |
| NW_FE_012 | 8/5/2024 | 7/23/2024 | 13 | PSK |
| NW_FE_012 | 8/5/2024 | 7/23/2024 | 13 | PSK |
| NW_FE_012 | 8/5/2024 | 7/23/2024 | 13 | PSK |
| NW_FE_012 | 8/5/2024 | 7/23/2024 | 13 | PSK |
| NW_FE_012 | 8/5/2024 | 7/23/2024 | 13 | PSK |
| NW_FE_014 | 8/6/2024 | 7/31/2024 | 6 | PSK |
| NW_FE_014 | 8/6/2024 | 7/31/2024 | 6 | PSK |
| NW_FE_014 | 8/6/2024 | 7/31/2024 | 6 | PSK |
| NW_FE_015 | 8/6/2024 | 7/31/2024 | 6 | PSK |
| NW_FE_015 | 8/6/2024 | 7/31/2024 | 6 | PSK |
| NW_FE_015 | 8/6/2024 | 7/31/2024 | 6 | PSK |
| NW_FE_015 | 8/6/2024 | 7/31/2024 | 6 | PSK |
| NW_FE_015 | 8/6/2024 | 7/31/2024 | 6 | PSK |
| NW_FE_014 | 8/6/2024 | 7/31/2024 | 6 | PSK |
| NW_FE_014 | 8/6/2024 | 7/31/2024 | 6 | PSK |
| NW_FE_014 | 8/6/2024 | 7/31/2024 | 6 | PSK |
| NW_FE_011 | 8/7/2024 | 7/17/2024 | 21 | PSK |
| NW_FE_014 | 8/7/2024 | 7/31/2024 | 7 | PSK |
| NW_FE_015 | 8/7/2024 | 7/31/2024 | 7 | PSK |
| NW_FE_015 | 8/7/2024 | 7/31/2024 | 7 | PSK |
| NW_FE_015 | 8/7/2024 | 7/31/2024 | 7 | PSK |
| NW_FE_015 | 8/7/2024 | 7/31/2024 | 7 | PSK |

| | | | | |
|---|---|---|---|---|
| NW_FE_013 | 8/7/2024 | 7/31/2024 | 7 | PSK |
| NW_FE_013 | 8/7/2024 | 7/31/2024 | 7 | PSK |
| NW_FE_013 | 8/7/2024 | 7/31/2024 | 7 | PSK |
| NW_FE_017 | 8/28/2024 | 8/7/2024 | 21 | PSK |
| NW_FE_022 | 8/28/2024 | 8/26/2024 | 2 | BTK |
| NW_FE_022 | 8/28/2024 | 8/26/2024 | 2 | BTK |
| NW_FE_022 | 8/29/2024 | 8/26/2024 | 3 | PSK |
| NW_FE_022 | 8/29/2024 | 8/26/2024 | 3 | PSK |
| NW_FE_017 | 8/29/2024 | 8/7/2024 | 22 | PSK |
| NW_FE_017 | 9/4/2024 | 8/7/2024 | 28 | PWK |
| NW_FE_022 | 9/4/2024 | 8/26/2024 | 9 | PWK |
| NW_FE_022 | 9/4/2024 | 8/26/2024 | 9 | PWK |
| NW_FE_023 | 9/4/2024 | 8/27/2024 | 8 | PWK |
| NW_FE_023 | 9/4/2024 | 8/27/2024 | 8 | PWK |
| NW_FE_024 | 9/11/2024 | 9/9/2024 | 2 | PWK |
| NW_FE_024 | 9/11/2024 | 9/9/2024 | 2 | PWK |
| NW_FE_024 | 9/11/2024 | 9/9/2024 | 2 | PWK |
| NW_FE_024 | 9/11/2024 | 9/9/2024 | 2 | BTK |
| NW_FE_024 | 9/11/2024 | 9/9/2024 | 2 | BTK |
| NW_FE_024 | 9/11/2024 | 9/9/2024 | 2 | BTK |
| NW_FE_024 | 9/11/2024 | 9/9/2024 | 2 | BTK |
| NW_FE_024 | 11/6/2024 | 9/9/2024 | 58 | PSK |
| NW_FE_024 | 11/6/2024 | 9/9/2024 | 58 | PSK |
| NW_FE_024 | 11/6/2024 | 9/9/2024 | 58 | PSK |
| NW_FE_014 | 11/6/2024 | 7/31/2024 | 98 | PSK |
| NW_FE_014 | 11/6/2024 | 7/31/2024 | 98 | PSK |
| NW_FE_014 | 11/6/2024 | 7/31/2024 | 98 | PSK |
| NW_FE_015 | 11/6/2024 | 7/31/2024 | 98 | PSK |
| NW_FE_015 | 11/6/2024 | 7/31/2024 | 98 | PSK |
| NW_FE_015 | 11/6/2024 | 7/31/2024 | 98 | PSK |
| NW_FE_014 | 11/6/2024 | 7/31/2024 | 98 | PSK |
| NW_FE_014 | 11/6/2024 | 7/31/2024 | 98 | PSK |
| NW_FE_014 | 11/6/2024 | 7/31/2024 | 98 | PSK |
| NW_FE_014 | 11/6/2024 | 7/31/2024 | 98 | PSK |
| NW_FE_025 | 11/6/2024 | 10/29/2024 | 8 | PSK |
| NW_FE_025 | 11/6/2024 | 10/29/2024 | 8 | PSK |
| NW_FE_025 | 11/6/2024 | 10/29/2024 | 8 | PSK |
| NW_FE_025 | 11/6/2024 | 10/29/2024 | 8 | PSK |
| NW_FE_025 | 11/6/2024 | 10/29/2024 | 8 | PSK |
| NW_FE_025 | 11/6/2024 | 10/29/2024 | 8 | PSK |
| NW_FE_025 | 11/6/2024 | 10/29/2024 | 8 | PSK |
| NW_FE_025 | 11/6/2024 | 10/29/2024 | 8 | PSK |
| NW_FE_011 | 11/18/2024 | 7/17/2024 | 124 | PWK |
| NW_FE_011 | 11/18/2024 | 7/17/2024 | 124 | PWK |
| NW_FE_011 | 11/18/2024 | 7/17/2024 | 124 | PWK |
| NW_FE_012 | 11/18/2024 | 7/23/2024 | 118 | PWK |
| NW_FE_012 | 11/18/2024 | 7/23/2024 | 118 | PWK |

| | | | | |
|---|---|---|---|---|
| NW_FE_012 | 11/18/2024 | 7/23/2024 | 118 | PWK |
| NW_FE_012 | 11/18/2024 | 7/23/2024 | 118 | PWK |
| NW_FE_013 | 11/18/2024 | 7/31/2024 | 110 | PWK |
| NW_FE_013 | 11/18/2024 | 7/31/2024 | 110 | PWK |
| NW_FE_015 | 11/18/2024 | 7/31/2024 | 110 | PWK |
| NW_FE_015 | 11/18/2024 | 7/31/2024 | 110 | PWK |
| NW_FE_013 | 11/18/2024 | 7/31/2024 | 110 | PWK |
| NW_FE_017 | 11/18/2024 | 8/7/2024 | 103 | PWK |
| NW_FE_017 | 11/18/2024 | 8/7/2024 | 103 | PWK |
| NW_FE_017 | 11/18/2024 | 8/7/2024 | 103 | PWK |
| NW_FE_017 | 11/18/2024 | 8/7/2024 | 103 | PWK |
| NW_FE_017 | 11/18/2024 | 8/7/2024 | 103 | PWK |
| NW_FE_017 | 11/18/2024 | 8/7/2024 | 103 | PWK |
| NW_FE_017 | 11/18/2024 | 8/7/2024 | 103 | PWK |
| NW_FE_019 | 11/18/2024 | 8/14/2024 | 96 | PWK |
| NW_FE_019 | 11/18/2024 | 8/14/2024 | 96 | PWK |
| NW_FE_019 | 11/18/2024 | 8/14/2024 | 96 | PWK |
| NW_FE_021 | 11/18/2024 | 8/14/2024 | 96 | PWK |
| NW_FE_012 | 12/2/2024 | 7/23/2024 | 132 | PWK |
| NW_FE_012 | 12/2/2024 | 7/23/2024 | 132 | PWK |
| NW_FE_012 | 12/2/2024 | 7/23/2024 | 132 | PWK |
| NW_FE_013 | 12/2/2024 | 7/31/2024 | 124 | PWK |
| NW_FE_013 | 12/2/2024 | 7/31/2024 | 124 | PWK |
| NW_FE_014 | 12/2/2024 | 7/31/2024 | 124 | PWK |
| NW_FE_015 | 12/2/2024 | 7/31/2024 | 124 | PWK |
| NW_FE_015 | 12/2/2024 | 7/31/2024 | 124 | PWK |
| NW_FE_015 | 12/2/2024 | 7/31/2024 | 124 | PWK |
| NW_FE_015 | 12/2/2024 | 7/31/2024 | 124 | PWK |
| NW_FE_013 | 12/2/2024 | 7/31/2024 | 124 | PWK |
| NW_FE_017 | 12/2/2024 | 8/7/2024 | 117 | PWK |
| NW_FE_017 | 12/2/2024 | 8/7/2024 | 117 | PWK |
| NW_FE_017 | 12/2/2024 | 8/7/2024 | 117 | PWK |
| NW_FE_017 | 12/2/2024 | 8/7/2024 | 117 | PWK |
| NW_FE_017 | 12/2/2024 | 8/7/2024 | 117 | PWK |
| NW_FE_017 | 12/2/2024 | 8/7/2024 | 117 | PWK |
| NW_FE_017 | 12/2/2024 | 8/7/2024 | 117 | PWK |
| NW_FE_017 | 12/2/2024 | 8/7/2024 | 117 | PWK |
| NW_FE_007 | 12/2/2024 | 6/19/2024 | 166 | PWK |
| NW_FE_008 | 12/2/2024 | 6/28/2024 | 157 | PWK |
| NW_FE_008 | 12/2/2024 | 6/28/2024 | 157 | PWK |
| NW_FE_008 | 12/2/2024 | 6/28/2024 | 157 | PWK |
| NW_FE_026 | 12/9/2024 | 12/3/2024 | 6 | PWK |
| NW_FE_026 | 12/9/2024 | 12/3/2024 | 6 | PWK |
| NW_FE_026 | 12/9/2024 | 12/3/2024 | 6 | PWK |
| NW_FE_026 | 12/9/2024 | 12/3/2024 | 6 | PWK |
| NW_FE_026 | 12/9/2024 | 12/3/2024 | 6 | PWK |
| NW_FE_026 | 12/9/2024 | 12/3/2024 | 6 | PWK |

| | | | | |
|---|---|---|---|---|
| NW_FE_026 | 12/9/2024 | 12/3/2024 | 6 | PWK |
| NW_FE_026 | 12/9/2024 | 12/3/2024 | 6 | PWK |
| NW_FE_026 | 12/9/2024 | 12/3/2024 | 6 | PWK |
| NW_FE_026 | 12/9/2024 | 12/3/2024 | 6 | PWK |
| NW_FE_026 | 12/9/2024 | 12/3/2024 | 6 | PWK |
| NW_FE_026 | 12/9/2024 | 12/3/2024 | 6 | PWK |
| NW_FE_026 | 12/9/2024 | 12/3/2024 | 6 | PWK |
| NW_FE_026 | 12/9/2024 | 12/3/2024 | 6 | PWK |
| NW_FE_026 | 12/9/2024 | 12/3/2024 | 6 | PWK |
| NW_FE_026 | 12/9/2024 | 12/3/2024 | 6 | PWK |
| NW_FE_026 | 12/9/2024 | 12/3/2024 | 6 | PWK |
| NW_FE_026 | 12/9/2024 | 12/3/2024 | 6 | PWK |
| NW_FE_026 | 12/9/2024 | 12/3/2024 | 6 | PWK |
| NW_FE_026 | 12/9/2024 | 12/3/2024 | 6 | PWK |
| NW_FE_026 | 12/9/2024 | 12/3/2024 | 6 | PWK |
| NW_FE_026 | 12/9/2024 | 12/3/2024 | 6 | PWK |
| NW_FE_026 | 12/9/2024 | 12/3/2024 | 6 | PWK |
| NW_FE_026 | 12/9/2024 | 12/3/2024 | 6 | PWK |
| NW_FE_026 | 12/10/2024 | 12/3/2024 | 7 | PWK |
| NW_FE_026 | 12/10/2024 | 12/3/2024 | 7 | PWK |
| NW_FE_026 | 12/10/2024 | 12/3/2024 | 7 | PWK |
| NW_FE_026 | 12/10/2024 | 12/3/2024 | 7 | PWK |
| NW_FE_026 | 12/10/2024 | 12/3/2024 | 7 | PWK |
| NW_FE_026 | 12/10/2024 | 12/3/2024 | 7 | PWK |
| NW_FE_026 | 12/10/2024 | 12/3/2024 | 7 | PWK |
| NW_FE_026 | 12/10/2024 | 12/3/2024 | 7 | PWK |
| NW_FE_026 | 12/10/2024 | 12/3/2024 | 7 | PWK |
| NW_FE_026 | 12/10/2024 | 12/3/2024 | 7 | PWK |
| NW_FE_026 | 12/10/2024 | 12/3/2024 | 7 | PWK |
| NW_FE_026 | 12/10/2024 | 12/3/2024 | 7 | PWK |
| NW_FE_026 | 12/10/2024 | 12/3/2024 | 7 | PWK |
| NW_FE_026 | 12/10/2024 | 12/3/2024 | 7 | PWK |
| NW_FE_026 | 12/10/2024 | 12/3/2024 | 7 | PWK |
| NW_FE_026 | 12/10/2024 | 12/3/2024 | 7 | PWK |
| NW_FE_026 | 12/10/2024 | 12/3/2024 | 7 | PWK |
| NW_FE_026 | 12/10/2024 | 12/3/2024 | 7 | PWK |
| NW_FE_026 | 12/10/2024 | 12/3/2024 | 7 | PWK |
| NW_FE_026 | 12/10/2024 | 12/3/2024 | 7 | PWK |
| NW_FE_026 | 12/10/2024 | 12/3/2024 | 7 | PWK |
| NW_FE_018 | 12/10/2024 | 8/14/2024 | 118 | PWK |
| NW_FE_018 | 12/10/2024 | 8/14/2024 | 118 | PWK |
| NW_FE_009 | 12/10/2024 | 7/3/2024 | 160 | PWK |
| NW_FE_012 | 1/30/2025 | 7/23/2024 | 191 | PSK |
| NW_FE_020 | 1/30/2025 | 8/14/2024 | 169 | PSK |
| NW_FE_020 | 1/30/2025 | 8/14/2024 | 169 | PSK |
| NW_FE_020 | 1/30/2025 | 8/14/2024 | 169 | PSK |
| NW_FE_020 | 1/30/2025 | 8/14/2024 | 169 | PSK |

| | | | | |
|---|---|---|---|---|
| NW_FE_021 | 1/30/2025 | 8/14/2024 | 169 | PSK |
| NW_FE_021 | 1/30/2025 | 8/14/2024 | 169 | PSK |
| NW_FE_021 | 1/30/2025 | 8/14/2024 | 169 | PSK |
| NW_FE_021 | 1/30/2025 | 8/14/2024 | 169 | PSK |
| NW_FE_020 | 1/30/2025 | 8/14/2024 | 169 | PSK |
| NW_FE_020 | 1/30/2025 | 8/14/2024 | 169 | PSK |
| NW_FE_020 | 1/30/2025 | 8/14/2024 | 169 | PSK |
| NW_FE_021 | 1/30/2025 | 8/14/2024 | 169 | PSK |
| NW_FE_021 | 1/30/2025 | 8/14/2024 | 169 | PSK |
| NW_FE_021 | 1/30/2025 | 8/14/2024 | 169 | PSK |
| NW_FE_020 | 1/30/2025 | 8/14/2024 | 169 | PSK |
| NW_FE_024 | 1/30/2025 | 9/9/2024 | 143 | PSK |
| NW_FE_024 | 1/30/2025 | 9/9/2024 | 143 | PSK |
| NW_FE_024 | 1/30/2025 | 9/9/2024 | 143 | PSK |
| NW_FE_024 | 1/30/2025 | 9/9/2024 | 143 | PSK |
| NW_FE_024 | 1/30/2025 | 9/9/2024 | 143 | PSK |
| NW_FE_024 | 1/30/2025 | 9/9/2024 | 143 | PSK |
| NW_FE_024 | 1/30/2025 | 9/9/2024 | 143 | PSK |
| NW_FE_006 | 2/19/2025 | 5/8/2024 | 287 | BTK |
| NW_FE_006 | 2/19/2025 | 5/8/2024 | 287 | BTK |
| NW_FE_006 | 2/19/2025 | 5/8/2024 | 287 | BTK |
| NW_FE_006 | 2/19/2025 | 5/8/2024 | 287 | BTK |
| NW_FE_006 | 2/19/2025 | 5/8/2024 | 287 | BTK |
| NW_FE_007 | 2/19/2025 | 6/19/2024 | 245 | BTK |
| NW_FE_007 | 2/19/2025 | 6/19/2024 | 245 | BTK |
| NW_FE_007 | 2/19/2025 | 6/19/2024 | 245 | BTK |
| NW_FE_007 | 2/19/2025 | 6/19/2024 | 245 | BTK |
| NW_FE_007 | 2/19/2025 | 6/19/2024 | 245 | BTK |
| NW_FE_007 | 2/19/2025 | 6/19/2024 | 245 | BTK |
| NW_FE_016 | 7/8/2025 | 8/7/2024 | 335 | BTK |
| NW_FE_016 | 7/8/2025 | 8/7/2024 | 335 | BTK |
| NW_FE_016 | 7/8/2025 | 8/7/2024 | 335 | BTK |
| NW_FE_016 | 7/8/2025 | 8/7/2024 | 335 | BTK |
| NW_FE_016 | 7/8/2025 | 8/7/2024 | 335 | BTK |
| NW_FE_016 | 7/8/2025 | 8/7/2024 | 335 | BTK |
| NW_FE_016 | 7/8/2025 | 8/7/2024 | 335 | BTK |
| NW_FE_016 | 7/8/2025 | 8/7/2024 | 335 | BTK |
| NW_FE_016 | 7/8/2025 | 8/7/2024 | 335 | BTK |
| NW_FE_008 | 7/22/2025 | 6/28/2024 | 389 | BTK |
| NW_FE_008 | 7/22/2025 | 6/28/2024 | 389 | BTK |
| NW_FE_008 | 7/22/2025 | 6/28/2024 | 389 | BTK |
| NW_FE_008 | 7/22/2025 | 6/28/2024 | 389 | BTK |
| NW_FE_009 | 7/22/2025 | 7/3/2024 | 384 | BTK |
| NW_FE_009 | 7/22/2025 | 7/3/2024 | 384 | BTK |
| NW_FE_009 | 7/22/2025 | 7/3/2024 | 384 | BTK |
| NW_FE_009 | 7/22/2025 | 7/3/2024 | 384 | BTK |
| NW_FE_010 | 7/22/2025 | 7/10/2024 | 377 | BTK |

| | | | | |
|---|---|---|---|---|
| NW_FE_010 | 7/22/2025 | 7/10/2024 | 377 | BTK |
| NW_FE_011 | 7/22/2025 | 7/17/2024 | 370 | BTK |
| NW_FE_012 | 7/22/2025 | 7/23/2024 | 364 | BTK |
| NW_FE_014 | 7/22/2025 | 7/31/2024 | 356 | BTK |
| NW_FE_015 | 7/22/2025 | 7/31/2024 | 356 | BTK |
| NW_FE_013 | 7/22/2025 | 7/31/2024 | 356 | BTK |
| NW_FE_016 | 7/22/2025 | 8/7/2024 | 349 | BTK |
| NW_FE_016 | 7/22/2025 | 8/7/2024 | 349 | BTK |
| NW_FE_016 | 7/22/2025 | 8/7/2024 | 349 | BTK |
| NW_FE_016 | 7/22/2025 | 8/7/2024 | 349 | BTK |
| NW_FE_016 | 7/22/2025 | 8/7/2024 | 349 | BTK |
| NW_FE_016 | 7/22/2025 | 8/7/2024 | 349 | BTK |
| NW_FE_016 | 7/22/2025 | 8/7/2024 | 349 | BTK |
| NW_FE_018 | 7/22/2025 | 8/15/2024 | 341 | BTK |
| NW_FE_001 | 7/23/2025 | 12/19/2023 | 582 | BTK |
| NW_FE_006 | 7/23/2025 | 5/8/2024 | 441 | BTK |
| NW_FE_006 | 7/23/2025 | 5/8/2024 | 441 | BTK |
| NW_FE_006 | 7/23/2025 | 5/8/2024 | 441 | BTK |
| NW_FE_006 | 7/23/2025 | 5/8/2024 | 441 | BTK |
| NW_FE_006 | 7/23/2025 | 5/8/2024 | 441 | BTK |
| NW_FE_006 | 7/23/2025 | 5/8/2024 | 441 | BTK |
| NW_FE_010 | 7/23/2025 | 7/10/2024 | 378 | BTK |
| NW_FE_010 | 7/23/2025 | 7/10/2024 | 378 | BTK |
| NW_FE_010 | 7/23/2025 | 7/10/2024 | 378 | BTK |
| NW_FE_010 | 7/23/2025 | 7/10/2024 | 378 | BTK |
| NW_FE_010 | 7/23/2025 | 7/10/2024 | 378 | BTK |
| NW_FE_010 | 7/23/2025 | 8/7/2024 | 350 | BTK |
| NW_FE_017 | 7/23/2025 | 8/7/2024 | 350 | BTK |
| NW_FE_027 | 8/6/2025 | 1/15/2025 | 203 | BTK |
| NW_FE_028 | 8/6/2025 | 1/16/2025 | 202 | BTK |
| NW_FE_029 | 8/6/2025 | 2/17/2025 | 170 | BTK |
| NW_FE_027 | 8/7/2025 | 1/15/2025 | 204 | PSK |
| NW_FE_028 | 8/7/2025 | 1/16/2025 | 203 | PSK |
| NW_FE_029 | 8/7/2025 | 2/17/2025 | 171 | PSK |
| NW_FE_025 | 8/27/2025 | 10/29/2024 | 302 | PSK |
| NW_FE_025 | 8/27/2025 | 10/29/2024 | 302 | PSK |
| NW_FE_025 | 8/27/2025 | 10/29/2024 | 302 | PSK |
| NW_FE_025 | 8/27/2025 | 10/29/2024 | 302 | PSK |
| NW_FE_025 | 8/27/2025 | 10/29/2024 | 302 | PSK |
| NW_FE_025 | 8/27/2025 | 10/29/2024 | 302 | PSK |
| NW_FE_025 | 8/27/2025 | 10/29/2024 | 302 | PSK |
| NW_FE_025 | 8/27/2025 | 10/29/2024 | 302 | PSK |
| NW_FE_025 | 8/27/2025 | 10/29/2024 | 302 | PSK |
| NW_FE_025 | 8/27/2025 | 10/29/2024 | 302 | PSK |
| NW_FE_025 | 8/27/2025 | 10/29/2024 | 302 | PSK |
| NW_FE_025 | 8/27/2025 | 10/29/2024 | 302 | PSK |
| NW_FE_025 | 8/27/2025 | 10/29/2024 | 302 | PSK |

| | | | | |
|---|---|---|---|---|
| NW_FE_025 | 8/27/2025 | 10/29/2024 | 302 | PSK |
| NW_FE_025 | 8/27/2025 | 10/29/2024 | 302 | PSK |
| NW_FE_025 | 8/27/2025 | 10/29/2024 | 302 | PSK |
| NW_FE_025 | 8/27/2025 | 10/29/2024 | 302 | PSK |
| NW_FE_025 | 8/27/2025 | 10/29/2024 | 302 | PSK |
| NW_FE_027 | 8/27/2025 | 1/15/2025 | 224 | PSK |
| NW_FE_027 | 8/27/2025 | 1/15/2025 | 224 | PSK |
| NW_FE_027 | 8/27/2025 | 1/15/2025 | 224 | PSK |
| NW_FE_027 | 8/27/2025 | 1/15/2025 | 224 | PSK |
| NW_FE_027 | 8/27/2025 | 1/15/2025 | 224 | PSK |
| NW_FE_027 | 8/28/2025 | 1/15/2025 | 225 | PSK |
| NW_FE_027 | 8/28/2025 | 1/15/2025 | 225 | PSK |
| NW_FE_027 | 8/28/2025 | 1/15/2025 | 225 | PSK |
| NW_FE_028 | 8/28/2025 | 1/16/2025 | 224 | PSK |
| NW_FE_028 | 8/28/2025 | 1/16/2025 | 224 | PSK |
| NW_FE_028 | 8/28/2025 | 1/16/2025 | 224 | PSK |
| NW_FE_028 | 8/28/2025 | 1/16/2025 | 224 | PSK |
| NW_FE_028 | 8/28/2025 | 1/16/2025 | 224 | PSK |
| NW_FE_028 | 8/28/2025 | 1/16/2025 | 224 | PSK |
| NW_FE_028 | 8/28/2025 | 1/16/2025 | 224 | PSK |
| NW_FE_028 | 8/28/2025 | 1/16/2025 | 224 | PSK |
| NW_FE_027 | 8/28/2025 | 1/15/2025 | 225 | PSK |
| NW_FE_027 | 8/28/2025 | 1/15/2025 | 225 | PSK |
| NW_FE_027 | 8/28/2025 | 1/15/2025 | 225 | PSK |
| NW_FE_027 | 8/28/2025 | 1/15/2025 | 225 | PSK |
| NW_FE_030 | 8/28/2025 | 2/17/2025 | 192 | PSK |
| NW_FE_030 | 8/28/2025 | 2/17/2025 | 192 | PSK |
| NW_FE_030 | 8/28/2025 | 2/17/2025 | 192 | PSK |
| NW_FE_030 | 8/28/2025 | 2/17/2025 | 192 | PSK |
| NW_FE_030 | 8/28/2025 | 2/17/2025 | 192 | PSK |
| NW_FE_030 | 8/28/2025 | 2/17/2025 | 192 | PSK |
| NW_FE_030 | 8/28/2025 | 2/17/2025 | 192 | PSK |
| NW_FE_030 | 8/28/2025 | 2/17/2025 | 192 | PSK |
| NW_FE_019 | 9/5/2025 | 8/14/2024 | 387 | PSK |
| NW_FE_019 | 9/5/2025 | 8/14/2024 | 387 | PSK |
| NW_FE_019 | 9/5/2025 | 8/14/2024 | 387 | PSK |
| NW_FE_023 | 9/5/2025 | 8/27/2024 | 374 | PSK |
| NW_FE_023 | 9/5/2025 | 8/27/2024 | 374 | PSK |
| NW_FE_023 | 9/5/2025 | 8/27/2024 | 374 | PSK |
| NW_FE_021 | 9/5/2025 | 8/14/2024 | 387 | PSK |
| NW_FE_029 | 9/9/2025 | 2/17/2025 | 204 | PSK |
| NW_FE_029 | 9/9/2025 | 2/17/2025 | 204 | PSK |
| NW_FE_029 | 9/9/2025 | 2/17/2025 | 204 | PSK |
| NW_FE_022 | 9/9/2025 | 8/26/2024 | 379 | PSK |
| NW_FE_024 | 9/9/2025 | 9/9/2024 | 365 | PSK |
| NW_FE_024 | 9/9/2025 | 9/9/2024 | 365 | PSK |
| NW_FE_025 | 9/9/2025 | 10/31/2024 | 313 | PSK |

| | | | | |
|---|---|---|---|---|
| NW_FE_025 | 9/9/2025 | 10/31/2024 | 313 | PSK |
| NW_FE_031 | 9/9/2025 | 3/3/2025 | 190 | PSK |
| NW_FE_031 | 9/9/2025 | 3/3/2025 | 190 | PSK |
| NW_FE_031 | 9/9/2025 | 3/3/2025 | 190 | PSK |
| NW_FE_031 | 9/9/2025 | 3/3/2025 | 190 | PSK |
| NW_FE_031 | 9/9/2025 | 3/3/2025 | 190 | PSK |
| NW_FE_031 | 9/9/2025 | 3/3/2025 | 190 | PSK |
| NW_FE_031 | 9/9/2025 | 3/3/2025 | 190 | PSK |
| NW_FE_031 | 9/9/2025 | 3/3/2025 | 190 | PSK |
| NW_FE_031 | 9/9/2025 | 3/3/2025 | 190 | PSK |
| NW_FE_031 | 9/9/2025 | 3/3/2025 | 190 | PSK |
| NW_FE_031 | 9/9/2025 | 3/3/2025 | 190 | PSK |
| NW_FE_031 | 9/9/2025 | 3/3/2025 | 190 | PSK |
| NW_FE_032 | 9/9/2025 | 4/7/2025 | 155 | PSK |
| NW_FE_032 | 9/9/2025 | 4/7/2025 | 155 | PSK |
| NW_FE_032 | 9/10/2025 | 4/7/2025 | 156 | PSK |
| NW_FE_032 | 9/10/2025 | 4/7/2025 | 156 | PSK |
| NW_FE_032 | 9/10/2025 | 4/7/2025 | 156 | PSK |
| NW_FE_033 | 9/10/2025 | 5/19/2025 | 114 | PSK |
| NW_FE_033 | 9/10/2025 | 5/19/2025 | 114 | PSK |
| NW_FE_033 | 9/10/2025 | 5/19/2025 | 114 | PSK |
| NW_FE_033 | 9/10/2025 | 5/19/2025 | 114 | PSK |
| NW_FE_033 | 9/10/2025 | 5/19/2025 | 114 | PSK |
| NW_FE_034 | 9/10/2025 | 7/1/2025 | 71 | PSK |
| NW_FE_034 | 9/10/2025 | 7/1/2025 | 71 | PSK |
| NW_FE_034 | 9/10/2025 | 7/1/2025 | 71 | PSK |
| NW_FE_034 | 9/10/2025 | 7/1/2025 | 71 | PSK |

| Sample ID |
|---|
| 20240401-PNNL-NRB-Seq1 |
| 20240401-PNNL-NRB-Seq2 |
| 20240401-PNNL-NRB-Seq3 |
| 20240401-PNNL-NRB-Seq4 |
| 20240401-PNNL-NRB-Seq5 |
| 20240401-PNNL-NRB-Seq6 |
| 20240401-PNNL-NRB-Seq7 |
| 20240418-PNNL-NRB-Seq10 |
| 20240418-PNNL-NRB-Seq11 |
| 20240418-PNNL-NRB-Seq12 |
| 20240418-PNNL-NRB-Seq13 |
| 20240418-PNNL-NRB-Seq14 |
| 20240418-PNNL-NRB-Seq15 |
| 20240418-PNNL-NRB-Seq17 |
| 20240418-PNNL-NRB-Seq18 |
| 20240418-PNNL-NRB-Seq19 |
| 20240418-PNNL-NRB-Seq20 |
| 20240418-PNNL-NRB-Seq21 |
| 20240418-PNNL-NRB-Seq22 |
| 20240418-PNNL-NRB-Seq23 |
| 20240418-PNNL-NRB-Seq25 |
| 20240418-PNNL-NRB-Seq26 |
| 20240418-PNNL-NRB-Seq27 |
| 20240418-PNNL-NRB-Seq28 |
| 20240418-PNNL-NRB-Seq29 |
| 20240418-PNNL-NRB-Seq30 |
| 20240418-PNNL-NRB-Seq31 |
| 20240418-PNNL-NRB-Seq33 |
| 20240418-PNNL-NRB-Seq34 |
| 20240418-PNNL-NRB-Seq35 |
| 20240418-PNNL-NRB-Seq36 |
| 20240418-PNNL-NRB-Seq37 |
| 20240418-PNNL-NRB-Seq38 |
| 20240418-PNNL-NRB-Seq39 |
| 20240418-PNNL-NRB-Seq9 |
| 20240513-PNNL-NRB-Seq41 |
| 20240513-PNNL-NRB-Seq42 |
| 20240513-PNNL-NRB-Seq43 |
| 20240513-PNNL-NRB-Seq44 |
| 20240513-PNNL-NRB-Seq45 |
| 20240513-PNNL-NRB-Seq46 |
| 20240513-PNNL-NRB-Seq47 |
| 20240513-PNNL-NRB-Seq48 |
| 20240513-PNNL-NRB-Seq49 |
| 20240513-PNNL-NRB-Seq50 |
| 20240513-PNNL-NRB-Seq51 |

20240513-PNNL-NRB-Seq52
20240513-PNNL-NRB-Seq53
20240718-PNNL-NRB-Seq55
20240718-PNNL-NRB-Seq56
20240718-PNNL-NRB-Seq57
20240718-PNNL-NRB-Seq58
20240718-PNNL-NRB-Seq59
20240719-PNNL-NRB-Seq61
20240719-PNNL-NRB-Seq62
20240719-PNNL-NRB-Seq63
20240719-PNNL-NRB-Seq64
20240719-PNNL-NRB-Seq65
20240724-PNNL-NRB-Seq67
20240724-PNNL-NRB-Seq68
20240724-PNNL-NRB-Seq69
20240725-PNNL-NRB-Seq71
20240725-PNNL-NRB-Seq72
20240725-PNNL-NRB-Seq73
20240805-PNNL-MPF-Seq75
20240805-PNNL-MPF-Seq76
20240805-PNNL-MPF-Seq77
20240805-PNNL-MPF-Seq78
20240805-PNNL-MPF-Seq79
20240805-PNNL-MPF-Seq80
20240805-PNNL-MPF-Seq81
20240805-PNNL-MPF-Seq82
20240805-PNNL-MPF-Seq83
20240805-PNNL-MPF-Seq84
20240805-PNNL-MPF-Seq85
20240805-PNNL-MPF-Seq86
20240806-PNNL-MPF-Seq88
20240806-PNNL-MPF-Seq89
20240806-PNNL-MPF-Seq90
20240806-PNNL-MPF-Seq91
20240806-PNNL-MPF-Seq92
20240806-PNNL-MPF-Seq93
20240806-PNNL-MPF-Seq94
20240806-PNNL-MPF-Seq95
20240806-PNNL-MPF-Seq96
20240806-PNNL-MPF-Seq97
20240806-PNNL-MPF-Seq98
20240807-PNNL-MPF-Seq100
20240807-PNNL-MPF-Seq101
20240807-PNNL-MPF-Seq102
20240807-PNNL-MPF-Seq103
20240807-PNNL-MPF-Seq104
20240807-PNNL-MPF-Seq105

20240807-PNNL-MPF-Seq106
20240807-PNNL-MPF-Seq107
20240807-PNNL-MPF-Seq108
20240828-PNNL-MPF-Seq110
20240828-PNNL-MPF-Seq111
20240828-PNNL-MPF-Seq112
20240829-PNNL-MPF-Seq114
20240829-PNNL-MPF-Seq115
20240829-PNNL-MPF-Seq116
20240829-PNNL-MPF-Seq118
20240829-PNNL-MPF-Seq119
20240829-PNNL-MPF-Seq120
20240829-PNNL-MPF-Seq121
20240829-PNNL-MPF-Seq122
20240911-PNNL-MPF-Seq124
20240911-PNNL-MPF-Seq125
20240911-PNNL-MPF-Seq126
20240911-PNNL-MPF-Seq128
20240911-PNNL-MPF-Seq129
20240911-PNNL-MPF-Seq130
20240911-PNNL-MPF-Seq131
20241106-PNNL-NRB-Seq133
20241106-PNNL-NRB-Seq134
20241106-PNNL-NRB-Seq135
20241106-PNNL-NRB-Seq136
20241106-PNNL-NRB-Seq137
20241106-PNNL-NRB-Seq138
20241106-PNNL-NRB-Seq139
20241106-PNNL-NRB-Seq140
20241106-PNNL-NRB-Seq141
20241106-PNNL-NRB-Seq142
20241106-PNNL-NRB-Seq143
20241106-PNNL-NRB-Seq144
20241106-PNNL-NRB-Seq145
20241106-PNNL-NRB-Seq146
20241106-PNNL-NRB-Seq147
20241106-PNNL-NRB-Seq148
20241106-PNNL-NRB-Seq149
20241106-PNNL-NRB-Seq150
20241106-PNNL-NRB-Seq151
20241106-PNNL-NRB-Seq152
20241106-PNNL-NRB-Seq153
20241118-PNNL-NRB-Seq155
20241118-PNNL-NRB-Seq156
20241118-PNNL-NRB-Seq157
20241118-PNNL-NRB-Seq158
20241118-PNNL-NRB-Seq159

20241118-PNNL-NRB-Seq160
20241118-PNNL-NRB-Seq161
20241118-PNNL-NRB-Seq162
20241118-PNNL-NRB-Seq163
20241118-PNNL-NRB-Seq164
20241118-PNNL-NRB-Seq165
20241118-PNNL-NRB-Seq166
20241118-PNNL-NRB-Seq167
20241118-PNNL-NRB-Seq168
20241118-PNNL-NRB-Seq169
20241118-PNNL-NRB-Seq170
20241118-PNNL-NRB-Seq171
20241118-PNNL-NRB-Seq172
20241118-PNNL-NRB-Seq173
20241118-PNNL-NRB-Seq174
20241118-PNNL-NRB-Seq175
20241118-PNNL-NRB-Seq176
20241118-PNNL-NRB-Seq177
20241202-PNNL-NRB-Seq179
20241202-PNNL-NRB-Seq180
20241202-PNNL-NRB-Seq181
20241202-PNNL-NRB-Seq182
20241202-PNNL-NRB-Seq183
20241202-PNNL-NRB-Seq184
20241202-PNNL-NRB-Seq185
20241202-PNNL-NRB-Seq186
20241202-PNNL-NRB-Seq187
20241202-PNNL-NRB-Seq188
20241202-PNNL-NRB-Seq189
20241202-PNNL-NRB-Seq190
20241202-PNNL-NRB-Seq191
20241202-PNNL-NRB-Seq192
20241202-PNNL-NRB-Seq193
20241202-PNNL-NRB-Seq194
20241202-PNNL-NRB-Seq195
20241202-PNNL-NRB-Seq196
20241202-PNNL-NRB-Seq197
20241202-PNNL-NRB-Seq198
20241202-PNNL-NRB-Seq199
20241202-PNNL-NRB-Seq200
20241202-PNNL-NRB-Seq201
20241209-PNNL-NRB-Seq203
20241209-PNNL-NRB-Seq204
20241209-PNNL-NRB-Seq205
20241209-PNNL-NRB-Seq206
20241209-PNNL-NRB-Seq207
20241209-PNNL-NRB-Seq208

20241209-PNNL-NRB-Seq209
20241209-PNNL-NRB-Seq210
20241209-PNNL-NRB-Seq211
20241209-PNNL-NRB-Seq212
20241209-PNNL-NRB-Seq213
20241209-PNNL-NRB-Seq214
20241209-PNNL-NRB-Seq215
20241209-PNNL-NRB-Seq216
20241209-PNNL-NRB-Seq217
20241209-PNNL-NRB-Seq218
20241209-PNNL-NRB-Seq219
20241209-PNNL-NRB-Seq220
20241209-PNNL-NRB-Seq221
20241209-PNNL-NRB-Seq222
20241209-PNNL-NRB-Seq223
20241209-PNNL-NRB-Seq224
20241209-PNNL-NRB-Seq225
20241209-PNNL-NRB-Seq226
20241209-PNNL-NRB-Seq227
20241209-PNNL-NRB-Seq228
20241209-PNNL-NRB-Seq229
20241209-PNNL-NRB-Seq230
20241209-PNNL-NRB-Seq231
20241209-PNNL-NRB-Seq232
20241209-PNNL-NRB-Seq233
20241209-PNNL-NRB-Seq234
20241209-PNNL-NRB-Seq235
20241209-PNNL-NRB-Seq236
20241209-PNNL-NRB-Seq237
20241209-PNNL-NRB-Seq238
20241209-PNNL-NRB-Seq239
20241209-PNNL-NRB-Seq240
20241209-PNNL-NRB-Seq241
20241209-PNNL-NRB-Seq242
20241209-PNNL-NRB-Seq243
20241209-PNNL-NRB-Seq244
20241209-PNNL-NRB-Seq245
20241209-PNNL-NRB-Seq246
20241209-PNNL-NRB-Seq247
20241209-PNNL-NRB-Seq248
20241209-PNNL-NRB-Seq249
20241209-PNNL-NRB-Seq250
20250130-PNNL-KMJ-Seq251
20250130-PNNL-KMJ-Seq252
20250130-PNNL-KMJ-Seq253
20250130-PNNL-KMJ-Seq254
20250130-PNNL-KMJ-Seq255

20250130-PNNL-KMJ-Seq256
20250130-PNNL-KMJ-Seq257
20250130-PNNL-KMJ-Seq258
20250130-PNNL-KMJ-Seq259
20250130-PNNL-KMJ-Seq260
20250130-PNNL-KMJ-Seq261
20250130-PNNL-KMJ-Seq262
20250130-PNNL-KMJ-Seq263
20250130-PNNL-KMJ-Seq264
20250130-PNNL-KMJ-Seq265
20250130-PNNL-KMJ-Seq266
20250130-PNNL-KMJ-Seq267
20250130-PNNL-KMJ-Seq268
20250130-PNNL-KMJ-Seq269
20250130-PNNL-KMJ-Seq270
20250130-PNNL-KMJ-Seq271
20250130-PNNL-KMJ-Seq272
20250130-PNNL-KMJ-Seq273
20250219-PNNL-IPM-Seq276
20250219-PNNL-IPM-Seq277
20250219-PNNL-IPM-Seq278
20250219-PNNL-IPM-Seq279
20250219-PNNL-IPM-Seq280
20250219-PNNL-IPM-Seq281
20250219-PNNL-IPM-Seq282
20250219-PNNL-IPM-Seq283
20250219-PNNL-IPM-Seq284
20250219-PNNL-IPM-Seq285
20250219-PNNL-IPM-Seq286
20250708-PNNL-OS-Seq287
20250708-PNNL-OS-Seq288
20250708-PNNL-OS-Seq289
20250708-PNNL-OS-Seq290
20250708-PNNL-OS-Seq291
20250708-PNNL-OS-Seq292
20250708-PNNL-OS-Seq293
20250708-PNNL-OS-Seq294
20250708-PNNL-OS-Seq295
20250722-PNNL-OS-Seq297
20250722-PNNL-OS-Seq298
20250722-PNNL-OS-Seq299
20250722-PNNL-OS-Seq300
20250722-PNNL-OS-Seq301
20250722-PNNL-OS-Seq302
20250722-PNNL-OS-Seq303
20250722-PNNL-OS-Seq304
20250722-PNNL-OS-Seq305

20250722-PNNL-OS-Seq306
20250722-PNNL-OS-Seq307
20250722-PNNL-OS-Seq308
20250722-PNNL-OS-Seq309
20250722-PNNL-OS-Seq310
20250722-PNNL-OS-Seq311
20250722-PNNL-OS-Seq312
20250722-PNNL-OS-Seq313
20250722-PNNL-OS-Seq314
20250722-PNNL-OS-Seq315
20250722-PNNL-OS-Seq316
20250722-PNNL-OS-Seq317
20250722-PNNL-OS-Seq318
20250722-PNNL-OS-Seq319
20250723-PNNL-OS-Seq321
20250723-PNNL-OS-Seq323
20250723-PNNL-OS-Seq324
20250723-PNNL-OS-Seq325
20250723-PNNL-OS-Seq326
20250723-PNNL-OS-Seq327
20250723-PNNL-OS-Seq328
20250723-PNNL-OS-Seq329
20250723-PNNL-OS-Seq330
20250723-PNNL-OS-Seq331
20250723-PNNL-OS-Seq332
20250723-PNNL-OS-Seq333
20250723-PNNL-OS-Seq334
20250723-PNNL-OS-Seq335
20250806-PNNL-NRB-Seq344
20250806-PNNL-NRB-Seq345
20250806-PNNL-NRB-Seq346
20250807-PNNL-NRB-Seq348
20250807-PNNL-NRB-Seq349
20250807-PNNL-NRB-Seq350
20250827-PNNL-OS-Seq352
20250827-PNNL-OS-Seq353
20250827-PNNL-OS-Seq354
20250827-PNNL-OS-Seq355
20250827-PNNL-OS-Seq356
20250827-PNNL-OS-Seq357
20250827-PNNL-OS-Seq358
20250827-PNNL-OS-Seq359
20250827-PNNL-OS-Seq360
20250827-PNNL-OS-Seq361
20250827-PNNL-OS-Seq362
20250827-PNNL-OS-Seq363
20250827-PNNL-OS-Seq364

20250827-PNNL-OS-Seq365
20250827-PNNL-OS-Seq366
20250827-PNNL-OS-Seq367
20250827-PNNL-OS-Seq368
20250827-PNNL-OS-Seq369
20250827-PNNL-OS-Seq370
20250827-PNNL-OS-Seq371
20250827-PNNL-OS-Seq372
20250827-PNNL-OS-Seq373
20250827-PNNL-OS-Seq374
20250828-PNNL-OS-Seq376
20250828-PNNL-OS-Seq377
20250828-PNNL-OS-Seq378
20250828-PNNL-OS-Seq379
20250828-PNNL-OS-Seq380
20250828-PNNL-OS-Seq381
20250828-PNNL-OS-Seq382
20250828-PNNL-OS-Seq383
20250828-PNNL-OS-Seq384
20250828-PNNL-OS-Seq385
20250828-PNNL-OS-Seq386
20250828-PNNL-OS-Seq387
20250828-PNNL-OS-Seq388
20250828-PNNL-OS-Seq389
20250828-PNNL-OS-Seq390
20250828-PNNL-OS-Seq391
20250828-PNNL-OS-Seq392
20250828-PNNL-OS-Seq393
20250828-PNNL-OS-Seq394
20250828-PNNL-OS-Seq395
20250828-PNNL-OS-Seq396
20250828-PNNL-OS-Seq397
20250828-PNNL-OS-Seq398
20250904-PNNL-OS-Seq403
20250904-PNNL-OS-Seq404
20250904-PNNL-OS-Seq405
20250904-PNNL-OS-Seq406
20250904-PNNL-OS-Seq407
20250904-PNNL-OS-Seq408
20250904-PNNL-OS-Seq409
20250909-PNNL-OS-Seq411
20250909-PNNL-OS-Seq412
20250909-PNNL-OS-Seq413
20250909-PNNL-OS-Seq414
20250909-PNNL-OS-Seq415
20250909-PNNL-OS-Seq416
20250909-PNNL-OS-Seq417

20250909-PNNL-OS-Seq418
20250909-PNNL-OS-Seq419
20250909-PNNL-OS-Seq420
20250909-PNNL-OS-Seq421
20250909-PNNL-OS-Seq422
20250909-PNNL-OS-Seq423
20250909-PNNL-OS-Seq424
20250909-PNNL-OS-Seq425
20250909-PNNL-OS-Seq426
20250909-PNNL-OS-Seq427
20250909-PNNL-OS-Seq428
20250909-PNNL-OS-Seq429
20250909-PNNL-OS-Seq430
20250909-PNNL-OS-Seq431
20250909-PNNL-OS-Seq432
20250910-PNNL-OS-Seq433
20250910-PNNL-OS-Seq435
20250910-PNNL-OS-Seq436
20250910-PNNL-OS-Seq437
20250910-PNNL-OS-Seq438
20250910-PNNL-OS-Seq439
20250910-PNNL-OS-Seq440
20250910-PNNL-OS-Seq441
20250910-PNNL-OS-Seq442
20250910-PNNL-OS-Seq443
20250910-PNNL-OS-Seq444
20250910-PNNL-OS-Seq445

| Sample Location | Tidal Stage |
| --- | --- |
| PNNL Dock | Flood |
| PNNL Dock | Flood |
| PNNL Dock | Flood |
| PNNL Dock | Flood |
| PNNL Dock | Flood |
| PNNL Dock | Flood |
| PNNL Dock | Flood |
| PNNL Dock | Ebb |
| PNNL Dock | Ebb |
| PNNL Dock | Ebb |
| PNNL Dock | Ebb |
| PNNL Dock | Ebb |
| PNNL Dock | Ebb |
| PNNL Dock | Ebb |
| PNNL Dock | Ebb |
| PNNL Dock | Ebb |
| PNNL Dock | Ebb |
| PNNL Dock | Ebb |
| PNNL Dock | Ebb |
| PNNL Dock | Ebb |
| PNNL Dock | Ebb |
| PNNL Dock | Ebb |
| PNNL Dock | Ebb |
| PNNL Dock | Ebb |
| PNNL Dock | Ebb |
| PNNL Dock | Ebb |
| PNNL Dock | Ebb |
| PNNL Dock | Ebb |
| PNNL Dock | Ebb |
| PNNL Dock | Ebb |
| PNNL Dock | Ebb |
| PNNL Dock | Ebb |
| PNNL Dock | Ebb |
| PNNL Dock | Ebb |
| PNNL Dock | Ebb |
| PNNL Dock | Flood |
| PNNL Dock | Flood |
| PNNL Dock | Flood |
| PNNL Dock | Flood |
| PNNL Dock | Flood |
| PNNL Dock | Flood |
| PNNL Dock | Flood |
| PNNL Dock | Flood |
| PNNL Dock | Flood |
| PNNL Dock | Flood |
| PNNL Dock | Flood |

| | |
|---|---|
| PNNL Dock | Flood |
| PNNL Dock | Flood |
| PNNL Dock | Slack Low |
| PNNL Dock | Slack Low |
| PNNL Dock | Slack Low |
| PNNL Dock | Flood |
| PNNL Dock | Flood |
| PNNL Dock | Ebb |
| PNNL Dock | Ebb |
| PNNL Dock | Flood |
| PNNL Dock | Slack Low |
| PNNL Dock | Slack Low |
| PNNL Dock | Ebb |
| PNNL Dock | Ebb |
| PNNL Dock | Ebb |
| PNNL Dock | Ebb |
| PNNL Dock | Ebb |
| PNNL Dock | Ebb |
| PNNL Dock | Ebb |
| PNNL Dock | Ebb |
| PNNL Dock | Ebb |
| PNNL Dock | Ebb |
| PNNL Dock | Ebb |
| PNNL Dock | Ebb |
| PNNL Dock | Flood |
| PNNL Dock | Ebb |
| PNNL Dock | Ebb |
| PNNL Dock | Ebb |
| PNNL Dock | Ebb |
| PNNL Dock | Ebb |
| PNNL Dock | Flood |
| PNNL Dock | Flood |
| PNNL Dock | Flood |
| John Wayne Marina | Flood |
| John Wayne Marina | Flood |
| John Wayne Marina | Flood |
| John Wayne Marina | Flood |
| John Wayne Marina | Flood |
| PNNL Dock | Flood |
| PNNL Dock | Flood |
| PNNL Dock | Flood |
| PNNL Dock | Flood |
| PNNL Dock | Flood |
| John Wayne Marina | Flood |
| John Wayne Marina | Flood |
| John Wayne Marina | Flood |
| John Wayne Marina | Flood |

| | |
|---|---|
| PNNL Dock | Ebb |
| PNNL Dock | Ebb |
| PNNL Dock | Ebb |
| Cline Spit | Flood |
| PNNL Dock | Slack High |
| PNNL Dock | Slack High |
| PNNL Dock | Slack High |
| PNNL Dock | Slack High |
| Cline Spit | Flood |
| Cline Spit | Flood |
| PNNL Dock | Slack High |
| PNNL Dock | Slack High |
| PNNL Dock | Slack High |
| PNNL Dock | Slack High |
| PNNL Dock | Slack High |
| PNNL Dock | Slack High |
| PNNL Dock | Slack High |
| PNNL Dock | Slack High |
| PNNL Dock | Slack High |
| PNNL Dock | Slack High |
| PNNL Dock | Slack High |
| PNNL Dock | Slack High |
| PNNL Dock | Slack High |
| PNNL Dock | Slack High |
| PNNL Dock | Flood |
| PNNL Dock | Flood |
| PNNL Dock | Flood |
| John Wayne Marina | Flood |
| John Wayne Marina | Flood |
| John Wayne Marina | Flood |
| PNNL Dock | Flood |
| PNNL Dock | Flood |
| PNNL Dock | Flood |
| PNNL Dock | Flood |
| PNNL Dock | Flood |
| PNNL Dock | Flood |
| PNNL Dock | Flood |
| PNNL Dock | Flood |
| PNNL Dock | Flood |
| PNNL Dock | Flood |
| PNNL Dock | Flood |
| PNNL Dock | Flood |
| PNNL Dock | Flood |
| PNNL Dock | Flood |
| PNNL Dock | Flood |
| PNNL Dock | Ebb |
| PNNL Dock | Ebb |

| | |
|---|---|
| PNNL Dock | Ebb |
| PNNL Dock | Ebb |
| PNNL Dock | Ebb |
| PNNL Dock | Ebb |
| John Wayne Marina | Flood |
| John Wayne Marina | Flood |
| PNNL Dock | Ebb |
| Cline Spit | Flood |
| Cline Spit | Flood |
| Cline Spit | Flood |
| Cline Spit | Flood |
| Cline Spit | Flood |
| Cline Spit | Flood |
| Cline Spit | Flood |
| Disco Bay | Flood |
| Disco Bay | Flood |
| Disco Bay | Flood |
| North of Port Townsend | Flood |
| PNNL Dock | Ebb |
| PNNL Dock | Ebb |
| PNNL Dock | Ebb |
| PNNL Dock | Ebb |
| PNNL Dock | Ebb |
| PNNL Dock | Flood |
| John Wayne Marina | Flood |
| John Wayne Marina | Flood |
| John Wayne Marina | Flood |
| John Wayne Marina | Flood |
| PNNL Dock | Ebb |
| Cline Spit | Flood |
| Cline Spit | Flood |
| Cline Spit | Flood |
| Cline Spit | Flood |
| Cline Spit | Flood |
| Cline Spit | Flood |
| Cline Spit | Flood |
| Cline Spit | Flood |
| PNNL Dock | Ebb |
| PNNL Dock | Ebb |
| PNNL Dock | Ebb |
| PNNL Dock | Ebb |
| PNNL Mesocosm Tank | n/a |
| PNNL Mesocosm Tank | n/a |
| PNNL Mesocosm Tank | n/a |
| PNNL Mesocosm Tank | n/a |
| PNNL Mesocosm Tank | n/a |
| PNNL Mesocosm Tank | n/a |

| | |
|---|---|
| PNNL Mesocosm Tank | n/a |
| PNNL Mesocosm Tank | n/a |
| PNNL Mesocosm Tank | n/a |
| PNNL Mesocosm Tank | n/a |
| PNNL Mesocosm Tank | n/a |
| PNNL Mesocosm Tank | n/a |
| PNNL Mesocosm Tank | n/a |
| PNNL Mesocosm Tank | n/a |
| PNNL Mesocosm Tank | n/a |
| PNNL Mesocosm Tank | n/a |
| PNNL Mesocosm Tank | n/a |
| PNNL Mesocosm Tank | n/a |
| PNNL Mesocosm Tank | n/a |
| PNNL Mesocosm Tank | n/a |
| PNNL Mesocosm Tank | n/a |
| PNNL Mesocosm Tank | n/a |
| PNNL Mesocosm Tank | n/a |
| PNNL Mesocosm Tank | n/a |
| PNNL Mesocosm Tank | n/a |
| PNNL Mesocosm Tank | n/a |
| PNNL Mesocosm Tank | n/a |
| PNNL Mesocosm Tank | n/a |
| PNNL Mesocosm Tank | n/a |
| PNNL Mesocosm Tank | n/a |
| PNNL Mesocosm Tank | n/a |
| PNNL Mesocosm Tank | n/a |
| PNNL Mesocosm Tank | n/a |
| PNNL Mesocosm Tank | n/a |
| PNNL Mesocosm Tank | n/a |
| PNNL Mesocosm Tank | n/a |
| PNNL Mesocosm Tank | n/a |
| PNNL Mesocosm Tank | n/a |
| PNNL Mesocosm Tank | n/a |
| PNNL Mesocosm Tank | n/a |
| PNNL Mesocosm Tank | n/a |
| PNNL Mesocosm Tank | n/a |
| PNNL Mesocosm Tank | n/a |
| PNNL Mesocosm Tank | n/a |
| PNNL Mesocosm Tank | n/a |
| PNNL Dock | Flood |
| PNNL Dock | Flood |
| PNNL Dock | Flood |
| PNNL Dock | Ebb |
| PNNL Dock | Flood |
| PNNL Dock | Flood |
| PNNL Dock | Flood |
| PNNL Dock | Flood |

| | |
|---|---|
| North of Port Townsend | Flood |
| North of Port Townsend | Flood |
| North of Port Townsend | Flood |
| North of Port Townsend | Flood |
| PNNL Dock | Flood |
| PNNL Dock | Flood |
| PNNL Dock | Flood |
| North of Port Townsend | Flood |
| North of Port Townsend | Flood |
| North of Port Townsend | Flood |
| PNNL Dock | Flood |
| PNNL Dock | Slack High |
| PNNL Dock | Slack High |
| PNNL Dock | Slack High |
| PNNL Dock | Slack High |
| PNNL Dock | Slack High |
| PNNL Dock | Slack High |
| PNNL Dock | Slack High |
| PNNL Dock | Flood |
| PNNL Dock | Flood |
| PNNL Dock | Flood |
| PNNL Dock | Flood |
| PNNL Dock | Flood |
| PNNL Dock | Ebb |
| PNNL Dock | Ebb |
| PNNL Dock | Ebb |
| PNNL Dock | Ebb |
| PNNL Dock | Ebb |
| PNNL Dock | Ebb |
| PNNL Dock | Slack high |
| PNNL Dock | Slack high |
| PNNL Dock | Slack high |
| PNNL Dock | Slack high |
| PNNL Dock | Slack high |
| PNNL Dock | Slack high |
| PNNL Dock | Slack high |
| PNNL Dock | Slack high |
| PNNL Dock | Slack high |
| PNNL Dock | Ebb |
| PNNL Dock | Ebb |
| PNNL Dock | Ebb |
| PNNL Dock | Ebb |
| PNNL Dock | Flood |
| PNNL Dock | Flood |
| PNNL Dock | Flood |
| PNNL Dock | Flood |
| PNNL Dock | Slack Low |

| | |
|---|---|
| PNNL Dock | Slack Low |
| PNNL Dock | Flood |
| PNNL Dock | Ebb |
| PNNL Dock | Flood |
| John Wayne Marina | Flood |
| PNNL Dock | Ebb |
| PNNL Dock | Slack High |
| PNNL Dock | Slack High |
| PNNL Dock | Slack High |
| PNNL Dock | Slack High |
| PNNL Dock | Slack High |
| PNNL Dock | Slack High |
| PNNL Dock | Slack High |
| PNNL Dock | Flood |
| PNNL Dock | Slack High |
| PNNL Dock | Flood |
| PNNL Dock | Flood |
| PNNL Dock | Flood |
| PNNL Dock | Flood |
| PNNL Dock | Flood |
| PNNL Dock | Flood |
| PNNL Dock | Slack Low |
| PNNL Dock | Slack Low |
| PNNL Dock | Slack Low |
| PNNL Dock | Slack Low |
| PNNL Dock | Slack Low |
| Cline Spit | Flood |
| Cline Spit | Flood |
| PNNL Dock | Ebb |
| PNNL Dock | Ebb |
| PNNL Dock | Ebb |
| PNNL Dock | Ebb |
| PNNL Dock | Ebb |
| PNNL Dock | Ebb |
| PNNL Dock | Flood |
| PNNL Dock | Flood |
| PNNL Dock | Flood |
| PNNL Dock | Flood |
| PNNL Dock | Flood |
| PNNL Dock | Flood |
| PNNL Dock | Flood |
| PNNL Dock | Flood |
| PNNL Dock | Flood |
| PNNL Dock | Flood |
| PNNL Dock | Flood |
| PNNL Dock | Flood |
| PNNL Dock | Flood |

| | |
|---|---|
| PNNL Dock | Flood |
| PNNL Dock | Flood |
| PNNL Dock | Flood |
| PNNL Dock | Flood |
| PNNL Dock | Flood |
| PNNL Dock | Ebb |
| PNNL Dock | Ebb |
| PNNL Dock | Ebb |
| PNNL Dock | Ebb |
| PNNL Dock | Ebb |
| PNNL Dock | Ebb |
| PNNL Dock | Ebb |
| PNNL Dock | Ebb |
| PNNL Dock | Ebb |
| PNNL Dock | Ebb |
| PNNL Dock | Ebb |
| PNNL Dock | Ebb |
| PNNL Dock | Ebb |
| PNNL Dock | Ebb |
| PNNL Dock | Ebb |
| PNNL Dock | Ebb |
| PNNL Dock | Ebb |
| PNNL Dock | Ebb |
| PNNL Dock | Ebb |
| PNNL Dock | Ebb |
| PNNL Dock | Flood |
| PNNL Dock | Flood |
| PNNL Dock | Flood |
| PNNL Dock | Flood |
| PNNL Dock | Flood |
| PNNL Dock | Flood |
| PNNL Dock | Flood |
| PNNL Dock | Flood |
| Disco Bay | Flood |
| Disco Bay | Flood |
| Disco Bay | Flood |
| PNNL Dock | Slack High |
| PNNL Dock | Slack High |
| PNNL Dock | Slack High |
| North of Port Townsend | Flood |
| PNNL Dock | Ebb |
| PNNL Dock | Ebb |
| PNNL Dock | Ebb |
| PNNL Dock | Slack High |
| PNNL Dock | Slack High |
| PNNL Dock | Slack High |
| PNNL Dock | Flood |

| | |
|---|---|
| PNNL Dock | Flood |
| PNNL Dock | Ebb |
| PNNL Dock | Ebb |
| PNNL Dock | Ebb |
| PNNL Dock | Ebb |
| PNNL Dock | Ebb |
| PNNL Dock | Ebb |
| PNNL Dock | Ebb |
| PNNL Dock | Ebb |
| PNNL Dock | Ebb |
| PNNL Dock | Ebb |
| PNNL Dock | Ebb |
| PNNL Dock | Ebb |
| PNNL Dock | Ebb |
| PNNL Dock | Ebb |
| PNNL Dock | Ebb |
| PNNL Dock | Ebb |
| PNNL Dock | Ebb |
| PNNL Dock | Ebb |
| PNNL Dock | Ebb |
| PNNL Dock | Ebb |
| PNNL Dock | Ebb |
| PNNL Dock | Ebb |
| PNNL Dock | Ebb |
| PNNL Dock | Ebb |
| PNNL Dock | Ebb |
| PNNL Dock | Ebb |

| Filter Fraction | Filter Material | Filtered Volume Use | NanoDrop Yield (ng/ | A260/A280 |
|---|---|---|---|---|
| Micro (> 5µm) | MCE | 3.500 | 4.600 | 1.580 |
| Pico (> 0.22µm ) | MCE | 1.000 | 2.400 | 1.560 |
| Pico (> 0.22µm ) | MCE | 1.000 | 3.800 | 1.420 |
| Micro (> 5µm) | MCE | 3.500 | 9.200 | 1.590 |
| Pico (> 0.22µm ) | MCE | 1.000 | 4.500 | 1.670 |
| Pico (> 0.22µm ) | MCE | 1.000 | 5.200 | 1.470 |
| Pico (> 0.22µm ) | MCE | 0.800 | 2.400 | 1.230 |
| Macro ( > 20µm) | Nylon | 6.250 | 49.100 | 1.850 |
| Pico (> 0.22µm ) | MCE | 1.000 | 27.600 | 1.840 |
| Pico (> 0.22µm ) | MCE | 1.000 | 25.200 | 1.810 |
| Pico (> 0.22µm ) | MCE | 0.500 | 15.500 | 1.870 |
| Pico (> 0.22µm ) | MCE | 0.500 | 9.500 | 1.750 |
| Pico (> 0.22µm ) | MCE | 0.500 | 11.800 | 1.820 |
| Macro ( > 20µm) | Nylon | 6.250 | 4.600 | 1.750 |
| Macro ( > 20µm) | Nylon | 6.250 | 5.700 | 1.610 |
| Pico (> 0.22µm ) | MCE | 1.000 | 5.500 | 1.440 |
| Pico (> 0.22µm ) | MCE | 1.000 | 4.700 | 1.750 |
| Pico (> 0.22µm ) | MCE | 0.500 | 4.000 | 1.520 |
| Pico (> 0.22µm ) | MCE | 0.500 | 3.800 | 1.300 |
| Pico (> 0.22µm ) | MCE | 0.500 | 3.600 | 1.510 |
| Macro ( > 20µm) | Nylon | 6.250 | 24.100 | 2.110 |
| Macro ( > 20µm) | Nylon | 6.250 | 15.900 | 1.900 |
| Pico (> 0.22µm ) | MCE | 1.000 | 21.300 | 1.100 |
| Pico (> 0.22µm ) | MCE | 1.000 | 12.900 | 2.000 |
| Pico (> 0.22µm ) | MCE | 0.500 | 8.700 | 1.940 |
| Pico (> 0.22µm ) | MCE | 0.500 | 9.300 | 2.110 |
| Pico (> 0.22µm ) | MCE | 0.500 | 28.100 | 1.160 |
| Macro ( > 20µm) | Nylon | 6.250 | 29.500 | 1.180 |
| Macro ( > 20µm) | Nylon | 6.250 | 4.800 | 1.780 |
| Pico (> 0.22µm ) | MCE | 1.000 | 2.700 | 2.250 |
| Pico (> 0.22µm ) | MCE | 1.000 | 3.000 | 1.830 |
| Pico (> 0.22µm ) | MCE | 0.500 | 18.600 | 1.110 |
| Pico (> 0.22µm ) | MCE | 0.500 | 2.400 | 1.590 |
| Pico (> 0.22µm ) | MCE | 0.500 | 32.600 | 1.200 |
| Macro ( > 20µm) | Nylon | 6.250 | 76.600 | 1.850 |
| Macro ( > 20µm) | Nylon | 2.750 | n/a | n/a |
| Macro ( > 20µm) | Nylon | 6.250 | n/a | n/a |
| Micro (> 5µm) | MCE | 1.500 | n/a | n/a |
| Micro (> 5µm) | MCE | 1.250 | n/a | n/a |
| Pico (> 0.22µm ) | MCE | 1.050 | n/a | n/a |
| Pico (> 0.22µm ) | MCE | 1.050 | n/a | n/a |
| Micro (> 5µm) | MCE | 2.600 | n/a | n/a |
| Micro (> 5µm) | MCE | 2.300 | n/a | n/a |
| Pico (> 0.22µm ) | MCE | 1.000 | n/a | n/a |
| Pico (> 0.22µm ) | MCE | 0.650 | n/a | n/a |
| Pico (> 0.22µm ) | MCE | 0.650 | n/a | n/a |

| | | | | |
|---|---|---|---|---|
| Pico (> 0.22µm ) | MCE | 0.650 | n/a | n/a |
| Pico (> 0.22µm ) | MCE | 0.650 | n/a | n/a |
| Pico (> 0.22µm ) | MCE | 5.000 | 29.900 | 1.760 |
| Micro (> 5µm) | MCE | 1.000 | 17.200 | 1.800 |
| Pico (> 0.22µm ) | MCE | 0.500 | 16.600 | 1.920 |
| Pico (> 0.22µm ) | MCE | 0.900 | 32.700 | 1.820 |
| Pico (> 0.22µm ) | MCE | 0.500 | 16.800 | 1.770 |
| Pico (> 0.22µm ) | MCE | 0.500 | 9.900 | 1.720 |
| Pico (> 0.22µm ) | MCE | 0.500 | 9.300 | 1.850 |
| Pico (> 0.22µm ) | MCE | 0.600 | 12.400 | 1.830 |
| Pico (> 0.22µm ) | MCE | 0.500 | 19.400 | 1.820 |
| Pico (> 0.22µm ) | MCE | 0.500 | 18.300 | 1.840 |
| Macro ( > 20µm) | Nylon | 0.400 | 11.500 | 1.720 |
| Macro ( > 20µm) | Nylon | 0.400 | 15.100 | 1.650 |
| Macro ( > 20µm) | Nylon | 0.400 | 12.500 | 1.550 |
| Macro ( > 20µm) | Nylon | 0.400 | 12.600 | 1.950 |
| Macro ( > 20µm) | Nylon | 0.400 | 12.000 | 1.990 |
| Macro ( > 20µm) | Nylon | 0.400 | 10.800 | 1.970 |
| Micro (> 5µm) | MCE | 2.500 | 7.800 | 1.930 |
| Micro (> 5µm) | MCE | 2.500 | 3.800 | 2.450 |
| Pico (> 0.22µm ) | MCE | 1.000 | 5.700 | 2.540 |
| Pico (> 0.22µm ) | MCE | 1.000 | 9.900 | 2.000 |
| Pico (> 0.22µm ) | MCE | 1.000 | 3.700 | 3.350 |
| Pico (> 0.22µm ) | MCE | 1.000 | 13.400 | 2.020 |
| Micro (> 5µm) | MCE | 1.250 | 11.800 | 2.050 |
| Pico (> 0.22µm ) | MCE | 0.750 | 6.800 | 2.090 |
| Pico (> 0.22µm ) | MCE | 0.750 | 4.000 | 2.880 |
| Pico (> 0.22µm ) | MCE | 0.750 | 4.100 | 2.520 |
| Pico (> 0.22µm ) | MCE | 0.750 | 9.700 | 1.810 |
| Pico (> 0.22µm ) | MCE | 0.750 | 8.500 | 2.180 |
| Micro (> 5µm) | MCE | 1.500 | 8.200 | 2.480 |
| Micro (> 5µm) | MCE | 1.500 | 13.100 | 1.920 |
| Micro (> 5µm) | MCE | 1.500 | 8.000 | 2.050 |
| Micro (> 5µm) | MCE | 0.500 | 18.000 | 1.600 |
| Micro (> 5µm) | MCE | 0.500 | 7.600 | 2.100 |
| Micro (> 5µm) | MCE | 0.750 | 13.100 | 1.970 |
| Micro (> 5µm) | MCE | 1.250 | 14.800 | 2.090 |
| Micro (> 5µm) | MCE | 1.250 | 20.400 | 2.000 |
| Pico (> 0.22µm ) | MCE | 1.000 | 5.700 | 2.440 |
| Pico (> 0.22µm ) | MCE | 1.000 | 6.300 | 2.010 |
| Pico (> 0.22µm ) | MCE | 1.000 | 13.000 | 1.890 |
| Micro (> 5µm) | MCE | 5.000 | 12.800 | 2.080 |
| Pico (> 0.22µm ) | MCE | 1.000 | 6.500 | 1.930 |
| Pico (> 0.22µm ) | MCE | 1.000 | 19.700 | 1.830 |
| Pico (> 0.22µm ) | MCE | 1.000 | 15.100 | 2.030 |
| Pico (> 0.22µm ) | MCE | 1.000 | 20.800 | 1.850 |
| Pico (> 0.22µm ) | MCE | 1.000 | 25.000 | 1.970 |

| | | | | |
|---|---|---|---|---|
| Micro (> 5µm) | MCE | 1.500 | 18.100 | 1.960 |
| Micro (> 5µm) | MCE | 1.750 | 16.200 | 2.060 |
| Micro (> 5µm) | MCE | 1.250 | 7.200 | 2.520 |
| Pico (> 0.22µm ) | MCE | 0.225 | 38.600 | 1.910 |
| Pico (> 0.22µm ) | aPES | 6.600 | 14.800 | 2.130 |
| Pico (> 0.22µm ) | aPES | 6.600 | 79.800 | 2.090 |
| Pico (> 0.22µm ) | aPES | 6.600 | 20.800 | 1.910 |
| Pico (> 0.22µm ) | aPES | 6.600 | 7.800 | 2.060 |
| Pico (> 0.22µm ) | MCE | 0.225 | 31.300 | 1.910 |
| Pico (> 0.22µm ) | MCE | 0.225 | 37.900 | 1.770 |
| Pico (> 0.22µm ) | aPES | 6.600 | 36.900 | 1.780 |
| Pico (> 0.22µm ) | aPES | 6.600 | 43.400 | 1.750 |
| Pico (> 0.22µm ) | aPES | 20.000 | 12.500 | 1.610 |
| Pico (> 0.22µm ) | aPES | 20.000 | 94.400 | 1.790 |
| Pico (> 0.22µm ) | aPES | 6.270 | 164.400 | 1.840 |
| Pico (> 0.22µm ) | aPES | 6.270 | 156.100 | 1.840 |
| Pico (> 0.22µm ) | aPES | 6.270 | 171.400 | 1.840 |
| Pico (> 0.22µm ) | aPES | 6.270 | 50.200 | 1.720 |
| Pico (> 0.22µm ) | aPES | 6.270 | 62.000 | 1.730 |
| Pico (> 0.22µm ) | aPES | 6.270 | 43.600 | 1.690 |
| Pico (> 0.22µm ) | aPES | 0.990 | 8.200 | 1.690 |
| Pico (> 0.22µm ) | aPES | 6.270 | 158.300 | 1.850 |
| Pico (> 0.22µm ) | aPES | 6.270 | 139.000 | 1.850 |
| Pico (> 0.22µm ) | aPES | 6.270 | 116.300 | 1.870 |
| Micro (> 5µm) | MCE | 1.500 | 9.700 | 2.030 |
| Micro (> 5µm) | MCE | 1.500 | 11.100 | 2.040 |
| Micro (> 5µm) | MCE | 1.500 | 8.400 | 1.900 |
| Micro (> 5µm) | MCE | 0.500 | 11.200 | 1.970 |
| Micro (> 5µm) | MCE | 0.500 | 8.800 | 1.950 |
| Micro (> 5µm) | MCE | 0.750 | 15.000 | 1.890 |
| Pico (> 0.22µm ) | MCE | 1.000 | 11.400 | 1.970 |
| Pico (> 0.22µm ) | MCE | 1.000 | 15.100 | 1.830 |
| Pico (> 0.22µm ) | MCE | 1.000 | 7.700 | 1.940 |
| Pico (> 0.22µm ) | MCE | 1.000 | 5.100 | 1.900 |
| Micro (> 5µm) | MCE | 2.000 | 11.100 | 1.590 |
| Micro (> 5µm) | Nylon | 12.000 | 8.300 | 1.700 |
| Pico (> 0.22µm ) | aPES | 10.000 | 24.400 | 1.840 |
| Viral (< 0.22µm) | aPES | 0.750 | 6.300 | 1.740 |
| Viral (< 0.22µm) | aPES | 2.500 | 9.600 | 1.630 |
| Viral (< 0.22µm) | aPES | 3.500 | 7.100 | 1.860 |
| Viral (< 0.22µm) | aPES | 4.250 | 10.200 | 1.730 |
| Viral (< 0.22µm) | aPES | 3.000 | 10.400 | 1.660 |
| Micro (> 5µm) | Nylon | 5.000 | 15.100 | 1.690 |
| Pico (> 0.22µm ) | MCE | 0.900 | 16.900 | 1.640 |
| Pico (> 0.22µm ) | MCE | 0.500 | 12.500 | 1.620 |
| Pico (> 0.22µm ) | MCE | 0.750 | 9.200 | 1.560 |
| Pico (> 0.22µm ) | MCE | 0.750 | 9.800 | 1.520 |

| | | | | |
|---|---|---|---|---|
| Pico (> 0.22µm ) | MCE | 0.750 | 8.700 | 1.460 |
| Pico (> 0.22µm ) | MCE | 0.750 | 8.300 | 1.420 |
| Pico (> 0.22µm ) | MCE | 1.000 | 13.300 | 1.500 |
| Pico (> 0.22µm ) | MCE | 1.000 | 12.700 | 1.480 |
| Pico (> 0.22µm ) | MCE | 1.000 | 17.900 | 1.600 |
| Pico (> 0.22µm ) | MCE | 1.000 | 15.500 | 1.580 |
| Micro (> 5µm) | MCE | 1.500 | 11.900 | 1.440 |
| Macro ( > 20µm) | Nylon | 2.000 | 7.900 | 1.320 |
| Micro (> 5µm) | Nylon | 3.000 | 11.400 | 1.430 |
| Pico (> 0.22µm ) | MCE | 0.425 | 45.100 | 1.740 |
| Pico (> 0.22µm ) | MCE | 0.425 | 49.200 | 1.730 |
| Pico (> 0.22µm ) | MCE | 0.450 | 54.700 | 1.820 |
| Pico (> 0.22µm ) | MCE | 0.450 | 48.600 | 1.790 |
| Viral (> 0.05µm) | MCE | 0.160 | 6.500 | 1.390 |
| Macro ( > 20µm) | Nylon | 7.300 | 7.100 | 1.620 |
| Micro (> 5µm) | Nylon | 3.650 | 12.800 | 1.690 |
| Pico (> 0.22µm ) | aPES | 2.500 | 73.200 | 1.850 |
| Pico (> 0.22µm ) | aPES | 3.110 | 65.500 | 1.850 |
| Micro (> 5µm) | MCE | 2.500 | 16.700 | 1.950 |
| Micro (> 5µm) | MCE | 2.500 | 17.400 | 2.040 |
| Pico (> 0.22µm ) | MCE | 0.750 | 15.400 | 1.910 |
| Pico (> 0.22µm ) | MCE | 1.000 | 17.300 | 1.930 |
| Pico (> 0.22µm ) | MCE | 1.000 | 20.400 | 1.780 |
| Pico (> 0.22µm ) | MCE | 1.250 | 17.500 | 1.860 |
| Micro (> 5µm) | MCE | 1.250 | 21.900 | 1.850 |
| Micro (> 5µm) | MCE | 1.250 | 22.400 | 1.250 |
| Pico (> 0.22µm ) | MCE | 1.000 | 24.100 | 1.840 |
| Pico (> 0.22µm ) | MCE | 1.000 | 23.800 | 1.890 |
| Micro (> 5µm) | MCE | 1.750 | 20.800 | 1.820 |
| Macro ( > 20µm) | Nylon | 8.000 | 21.600 | 1.880 |
| Macro ( > 20µm) | Nylon | 1.000 | 26.100 | 1.810 |
| Macro ( > 20µm) | Nylon | 0.500 | 22.100 | 1.900 |
| Macro ( > 20µm) | Nylon | 1.000 | 35.200 | 1.870 |
| Pico (> 0.22µm ) | MCE | 0.225 | 23.600 | 1.800 |
| Pico (> 0.22µm ) | MCE | 0.225 | 28.700 | 1.850 |
| Pico (> 0.22µm ) | MCE | 0.225 | 18.100 | 1.900 |
| Pico (> 0.22µm ) | MCE | 0.900 | 14.600 | 1.930 |
| Pico (> 0.22µm ) | MCE | 1.200 | 18.200 | 1.880 |
| Pico (> 0.22µm ) | MCE | 0.975 | 15.700 | 1.870 |
| Pico (> 0.22µm ) | MCE | 1.000 | 12.900 | 1.750 |
| Pico (> 0.22µm ) | MCE | 1.000 | 13.300 | 1.900 |
| Macro ( > 20µm) | Nylon | 12.000 | 15.360 | 1.970 |
| Macro ( > 20µm) | Nylon | 8.000 | 18.200 | 1.890 |
| Micro (> 5µm) | MCE | 3.000 | 16.000 | 1.870 |
| Micro (> 5µm) | MCE | 3.000 | 15.900 | 1.890 |
| Micro (> 5µm) | MCE | 3.500 | 16.700 | 1.870 |
| Micro (> 5µm) | MCE | 3.500 | 40.300 | 1.620 |

| | | | | |
|---|---|---|---|---|
| Micro (> 5µm) | MCE | 3.000 | 16.400 | 1.960 |
| Micro (> 5µm) | MCE | 2.500 | 18.200 | 1.820 |
| Pico (> 0.22µm ) | MCE | 2.000 | 30.900 | 1.800 |
| Pico (> 0.22µm ) | MCE | 2.000 | 39.700 | 1.730 |
| Pico (> 0.22µm ) | MCE | 2.000 | 35.100 | 1.840 |
| Pico (> 0.22µm ) | MCE | 2.000 | 11.200 | 1.840 |
| Pico (> 0.22µm ) | aPES | 2.725 | 25.000 | 1.830 |
| Pico (> 0.22µm ) | aPES | 2.725 | 26.200 | 1.830 |
| Pico (> 0.22µm ) | aPES | 2.725 | 45.600 | 1.650 |
| Pico (> 0.22µm ) | aPES | 2.725 | 30.300 | 1.770 |
| Pico (> 0.22µm ) | MCE | 1.800 | 30.400 | 1.790 |
| Pico (> 0.22µm ) | MCE | 1.800 | 26.800 | 1.810 |
| Pico (> 0.22µm ) | MCE | 1.700 | 27.200 | 1.870 |
| Pico (> 0.22µm ) | MCE | 1.800 | 25.000 | 1.780 |
| Pico (> 0.22µm ) | aPES | 2.750 | 29.400 | 1.880 |
| Pico (> 0.22µm ) | aPES | 2.750 | 28.000 | 1.790 |
| Pico (> 0.22µm ) | aPES | 2.750 | 31.300 | 1.810 |
| Pico (> 0.22µm ) | aPES | 2.750 | 32.600 | 1.780 |
| Viral (< 0.22µm) | aPES | 2.250 | 3.900 | 1.100 |
| Viral (< 0.22µm) | aPES | 2.250 | 5.000 | 1.250 |
| Viral (< 0.22µm) | aPES | 2.250 | 2.600 | 1.220 |
| Viral (< 0.22µm) | aPES | 2.250 | 3.600 | 1.390 |
| Viral (< 0.22µm) | aPES | 1.130 | 4.500 | 1.370 |
| Viral (< 0.22µm) | aPES | 1.130 | 7.100 | 1.450 |
| Viral (< 0.22µm) | aPES | 1.130 | 8.100 | 1.430 |
| Viral (< 0.22µm) | aPES | 1.130 | 8.400 | 1.390 |
| Viral (< 0.22µm) | MCE | 1.500 | 11.100 | 1.510 |
| Viral (< 0.22µm) | MCE | 1.500 | 8.600 | 1.550 |
| Viral (< 0.22µm) | aPES | 2.250 | 8.100 | 1.550 |
| Viral (< 0.22µm) | aPES | 2.250 | 10.400 | 1.530 |
| Viral (< 0.22µm) | aPES | 2.250 | 8.800 | 1.570 |
| Viral (< 0.22µm) | aPES | 2.250 | 8.600 | 1.450 |
| Viral (< 0.22µm) | aPES | 1.500 | 7.200 | 1.360 |
| Viral (< 0.22µm) | aPES | 1.500 | 7.600 | 1.460 |
| Viral (< 0.22µm) | aPES | 1.500 | 6.700 | 1.390 |
| Viral (< 0.22µm) | aPES | 1.500 | 7.200 | 1.260 |
| Viral (< 0.22µm) | MCE | 1.500 | 6.500 | 1.320 |
| Viral (< 0.22µm) | MCE | 1.500 | 4.800 | 1.190 |
| Viral (< 0.22µm) | MCE | 0.675 | 4.300 | 1.210 |
| Pico (> 0.22µm ) | MCE | 1.800 | 36.900 | 1.740 |
| Pico (> 0.22µm ) | aPES | 5.000 | 6.700 | 1.300 |
| Pico (> 0.22µm ) | MCE | 0.600 | 4.800 | 1.130 |
| Macro ( > 20µm) | Nylon | 5.000 | 5.500 | 1.970 |
| Macro ( > 20µm) | Nylon | 8.000 | 5.700 | 1.720 |
| Micro (> 5µm) | Nylon | 2.667 | 6.200 | 4.560 |
| Micro (> 5µm) | Nylon | 2.667 | 6.300 | 1.830 |
| Micro (> 5µm) | Nylon | 2.667 | 5.900 | 1.660 |

| | | | | |
|---|---|---|---|---|
| Macro ( > 20µm) | Nylon | 7.500 | 3.700 | 1.410 |
| Micro (> 5µm) | Nylon | 2.500 | 6.300 | 1.620 |
| Micro (> 5µm) | Nylon | 2.500 | 5.900 | 1.640 |
| Micro (> 5µm) | Nylon | 2.500 | 6.300 | 1.570 |
| Pico (> 0.22µm ) | aPES | 1.667 | 6.600 | 1.550 |
| Pico (> 0.22µm ) | aPES | 1.667 | 6.700 | 1.550 |
| Pico (> 0.22µm ) | aPES | 1.667 | 6.000 | 1.500 |
| Pico (> 0.22µm ) | aPES | 1.667 | 6.500 | 1.400 |
| Pico (> 0.22µm ) | aPES | 1.667 | 8.600 | 1.680 |
| Pico (> 0.22µm ) | aPES | 1.667 | 7.200 | 1.610 |
| Pico (> 0.22µm ) | aPES | 3.110 | 9.900 | 1.650 |
| Micro (> 5µm) | Nylon | 8.000 | 11.300 | 1.830 |
| Micro (> 5µm) | Nylon | 8.000 | 7.600 | 1.630 |
| Micro (> 5µm) | Nylon | 8.000 | 7.300 | 1.760 |
| Pico (> 0.22µm ) | aPES | 4.667 | 7.400 | 2.420 |
| Pico (> 0.22µm ) | aPES | 4.667 | 73.800 | 1.840 |
| Pico (> 0.22µm ) | aPES | 4.667 | 44.400 | 1.860 |
| Macro ( > 20µm) | Nylon | 20.000 | 35.000 | 1.820 |
| Pico (> 0.22µm ) | MCE | 1.000 | 8.500 | 2.050 |
| Pico (> 0.22µm ) | MCE | 0.650 | 11.300 | 2.050 |
| Pico (> 0.22µm ) | MCE | 0.650 | 11.900 | 2.120 |
| Pico (> 0.22µm ) | MCE | 0.650 | 7.800 | 2.260 |
| Pico (> 0.22µm ) | MCE | 0.650 | 8.600 | 2.130 |
| Macro ( > 20µm) | Nylon | 2.500 | 27.700 | 2.070 |
| Macro ( > 20µm) | Nylon | 2.500 | 33.000 | 2.020 |
| Macro ( > 20µm) | Nylon | 2.500 | 16.200 | 2.120 |
| Micro (> 5µm) | MCE | 3.500 | 7.000 | 2.200 |
| Micro (> 5µm) | MCE | 4.000 | 4.500 | 2.180 |
| Pico (> 0.22µm ) | MCE | 0.600 | 8.900 | 2.070 |
| Micro (> 5µm) | MCE | 1.700 | 39.800 | 1.880 |
| Micro (> 5µm) | MCE | 1.500 | 31.100 | 1.900 |
| Micro (> 5µm) | MCE | 1.500 | 23.300 | 1.970 |
| Micro (> 5µm) | MCE | 1.500 | 17.000 | 1.800 |
| Pico (> 0.22µm ) | aPES | 2.000 | 23.600 | 1.990 |
| Pico (> 0.22µm ) | aPES | 2.000 | 12.500 | 1.970 |
| Pico (> 0.22µm ) | aPES | 2.000 | 27.800 | 2.060 |
| Pico (> 0.22µm ) | aPES | 2.000 | 49.200 | 2.160 |
| Macro ( > 20µm) | Nylon | 3.000 | 16.600 | 1.840 |
| Macro ( > 20µm) | Nylon | 2.000 | 33.800 | 1.820 |
| Macro ( > 20µm) | Nylon | 2.000 | 56.100 | 1.770 |
| Micro (> 5µm) | MCE | 2.000 | 13.500 | 1.930 |
| Micro (> 5µm) | MCE | 2.000 | 17.900 | 1.910 |
| Macro ( > 20µm) | Nylon | 5.000 | 33.900 | 1.870 |
| Micro (> 5µm) | MCE | 5.000 | 10.800 | 1.650 |
| Pico (> 0.22µm ) | MCE | 1.200 | 7.200 | 1.510 |
| Pico (> 0.22µm ) | MCE | 1.200 | 5.900 | 1.490 |
| Macro ( > 20µm) | Nylon | 10.000 | 21.300 | 1.870 |

| | | | | |
|---|---|---|---|---|
| Micro (> 5µm) | MCE | 2.000 | 13.600 | 1.700 |
| Macro ( > 20µm) | Nylon | 10.000 | 45.600 | 2.010 |
| Macro ( > 20µm) | Nylon | 5.000 | 8.300 | 1.340 |
| Macro ( > 20µm) | Nylon | 11.000 | 10.600 | 1.540 |
| Macro ( > 20µm) | Nylon | 11.000 | 5.600 | 1.310 |
| Macro ( > 20µm) | Nylon | 9.000 | 10.700 | 1.460 |
| Micro (> 5µm) | MCE | 1.800 | 9.800 | 1.640 |
| Micro (> 5µm) | MCE | 1.500 | 11.100 | 1.670 |
| Macro ( > 20µm) | Nylon | 3.300 | 10.100 | 1.580 |
| Micro (> 5µm) | MCE | 1.700 | 9.100 | 1.660 |
| Micro (> 5µm) | MCE | 1.300 | 13.300 | 1.740 |
| Macro ( > 20µm) | Nylon | 3.000 | 8.000 | 1.570 |
| Macro ( > 20µm) | Nylon | 3.200 | 9.100 | 1.550 |
| Macro ( > 20µm) | Nylon | 7.000 | 18.800 | 1.850 |
| Viral (< 0.22µm) | MCE | 1.000 | 5.700 | 1.530 |
| Macro ( > 20µm) | Nylon | 6.250 | 30.700 | 1.920 |
| Micro (> 5µm) | MCE | 1.250 | Below Detection | 2.030 |
| Pico (> 0.22µm ) | MCE | 1.050 | 13.200 | 1.670 |
| Pico (> 0.22µm ) | MCE | 1.050 | 15.900 | 2.120 |
| Micro (> 5µm) | MCE | 2.750 | 73.800 | 1.200 |
| Micro (> 5µm) | MCE | 2.300 | 32.500 | 1.800 |
| Pico (> 0.22µm ) | MCE | 0.500 | 26.700 | 1.180 |
| Pico (> 0.22µm ) | MCE | 0.500 | 14.800 | 2.020 |
| Micro (> 5µm) | MCE | 1.000 | 29.200 | 1.860 |
| Pico (> 0.22µm ) | MCE | 0.500 | 20.600 | 1.910 |
| Pico (> 0.22µm ) | MCE | 0.500 | 24.100 | 1.890 |
| Micro (> 5µm) | MCE | 1.250 | 38.400 | 1.950 |
| Macro ( > 20µm) | Nylon | 8.000 | 114.600 | 1.390 |
| Viral (< 0.22µm) | aPES | 1.563 | 27.500 | 1.170 |
| Viral (< 0.22µm) | aPES | 2.375 | 50.600 | 1.190 |
| Viral (< 0.22µm) | aPES | 8.250 | 25.500 | 1.290 |
| Viral (< 0.22µm) | aPES | 1.563 | 19.800 | 2.030 |
| Viral (< 0.22µm) | aPES | 2.375 | 13.100 | 1.760 |
| Viral (< 0.22µm) | aPES | 8.250 | 10.700 | 1.470 |
| Pico (> 0.22µm ) | aPES | 10.000 | 46.300 | 1.860 |
| Pico (> 0.22µm ) | aPES | 10.000 | 41.400 | 1.850 |
| Pico (> 0.22µm ) | aPES | 10.000 | 45.900 | 1.830 |
| Pico (> 0.22µm ) | aPES | 0.750 | 8.800 | 1.550 |
| Pico (> 0.22µm ) | aPES | 0.750 | 1.900 | 2.540 |
| Pico (> 0.22µm ) | aPES | 0.750 | 1.700 | 5.880 |
| Pico (> 0.22µm ) | aPES | 2.500 | 5.600 | 2.250 |
| Pico (> 0.22µm ) | aPES | 2.500 | 4.700 | 2.140 |
| Pico (> 0.22µm ) | aPES | 2.500 | 7.000 | 2.120 |
| Pico (> 0.22µm ) | aPES | 3.500 | 10.100 | 1.990 |
| Pico (> 0.22µm ) | aPES | 3.500 | 9.400 | 1.940 |
| Pico (> 0.22µm ) | aPES | 3.500 | 12.900 | 1.910 |
| Pico (> 0.22µm ) | aPES | 4.250 | 13.200 | 1.940 |

| | | | | |
|---|---|---|---|---|
| Pico (> 0.22µm ) | aPES | 4.250 | 14.900 | 1.800 |
| Pico (> 0.22µm ) | aPES | 4.250 | 17.600 | 1.760 |
| Pico (> 0.22µm ) | aPES | 3.000 | 19.900 | 1.880 |
| Pico (> 0.22µm ) | aPES | 3.000 | 27.600 | 1.860 |
| Pico (> 0.22µm ) | aPES | 3.000 | 12.100 | 1.970 |
| Pico (> 0.22µm ) | aPES | 1.125 | 2.200 | 2.010 |
| Pico (> 0.22µm ) | aPES | 1.125 | 1.500 | 3.180 |
| Pico (> 0.22µm ) | aPES | 1.125 | 2.500 | 2.200 |
| Pico (> 0.22µm ) | aPES | 1.125 | 0.800 | 0.970 |
| Pico (> 0.22µm ) | aPES | 1.313 | 3.900 | 1.880 |
| Pico (> 0.22µm ) | aPES | 1.313 | 4.100 | 1.870 |
| Pico (> 0.22µm ) | aPES | 1.313 | 3.700 | 1.960 |
| Pico (> 0.22µm ) | aPES | 1.313 | 2.900 | 1.890 |
| Pico (> 0.22µm ) | aPES | 5.005 | 3.000 | 1.650 |
| Pico (> 0.22µm ) | aPES | 5.005 | 0.600 | 2.520 |
| Pico (> 0.22µm ) | aPES | 5.005 | 2.200 | 3.190 |
| Pico (> 0.22µm ) | aPES | 5.005 | 1.400 | 2.690 |
| Pico (> 0.22µm ) | aPES | 5.000 | 4.200 | 1.850 |
| Pico (> 0.22µm ) | aPES | 5.000 | 4.300 | 1.950 |
| Pico (> 0.22µm ) | aPES | 5.000 | 4.800 | 1.750 |
| Pico (> 0.22µm ) | aPES | 5.000 | 3.400 | 2.140 |
| Viral (< 0.22µm) | aPES | 0.700 | 1.100 | 2.110 |
| Viral (< 0.22µm) | aPES | 0.700 | 1.100 | 4.870 |
| Viral (< 0.22µm) | aPES | 0.700 | 1.300 | 1.040 |
| Viral (< 0.22µm) | aPES | 0.700 | 0.800 | 1.550 |
| Pico (> 0.22µm ) | aPES | 9.375 | 8.200 | 1.730 |
| Pico (> 0.22µm ) | aPES | 9.375 | 8.600 | 1.880 |
| Pico (> 0.22µm ) | aPES | 9.375 | 10.700 | 1.950 |
| Pico (> 0.22µm ) | aPES | 9.375 | 10.000 | 1.860 |
| Viral (< 0.22µm) | aPES | 8.750 | 5.500 | 1.990 |
| Viral (< 0.22µm) | aPES | 8.750 | 5.300 | 2.130 |
| Viral (< 0.22µm) | aPES | 8.750 | 6.800 | 1.760 |
| Viral (< 0.22µm) | aPES | 8.750 | 5.700 | 1.790 |
| Viral (< 0.22µm) | aPES | 0.167 | 1.800 | 9.950 |
| Viral (< 0.22µm) | aPES | 0.167 | 0.800 | -4.560 |
| Viral (< 0.22µm) | aPES | 0.167 | 0.700 | -1.200 |
| Viral (< 0.22µm) | aPES | 0.333 | 1.500 | 9.460 |
| Viral (< 0.22µm) | aPES | 0.333 | 1.100 | 10.440 |
| Viral (< 0.22µm) | aPES | 0.333 | 0.700 | 5.160 |
| Viral (< 0.22µm) | aPES | 0.650 | 0.900 | 2.980 |
| Pico (> 0.22µm ) | aPES | 18.750 | 36.400 | 1.910 |
| Pico (> 0.22µm ) | aPES | 9.375 | 32.300 | 1.900 |
| Pico (> 0.22µm ) | aPES | 9.375 | 28.800 | 1.960 |
| Viral (< 0.22µm) | aPES | 1.000 | 2.500 | 3.490 |
| Viral (< 0.22µm) | aPES | 7.000 | 44.800 | 1.930 |
| Viral (< 0.22µm) | aPES | 3.500 | 47.500 | 1.860 |
| Viral (< 0.22µm) | aPES | 2.000 | 1.200 | 27.710 |

| | | | | |
|---|---|---|---|---|
| Viral (< 0.22µm) | aPES | 1.000 | 1.000 | 32.240 |
| Pico (> 0.22µm ) | aPES | 13.333 | 24.200 | 1.890 |
| Pico (> 0.22µm ) | aPES | 6.667 | 15.900 | 2.040 |
| Pico (> 0.22µm ) | aPES | 13.333 | 22.000 | 1.830 |
| Pico (> 0.22µm ) | aPES | 6.667 | 21.700 | 1.910 |
| Pico (> 0.22µm ) | aPES | 13.333 | 12.300 | 1.860 |
| Pico (> 0.22µm ) | aPES | 6.667 | 7.100 | 1.920 |
| Viral (< 0.22µm) | aPES | 12.667 | 3.500 | 2.450 |
| Viral (< 0.22µm) | aPES | 6.333 | 6.800 | 1.880 |
| Viral (< 0.22µm) | aPES | 12.667 | 4.600 | 2.260 |
| Viral (< 0.22µm) | aPES | 6.333 | 5.800 | 1.590 |
| Viral (< 0.22µm) | aPES | 12.667 | 6.900 | 2.030 |
| Viral (< 0.22µm) | aPES | 6.333 | 4.100 | 2.220 |
| Pico (> 0.22µm ) | aPES | 11.667 | 4.900 | 2.010 |
| Pico (> 0.22µm ) | aPES | 5.833 | 9.200 | 2.020 |
| Pico (> 0.22µm ) | aPES | 6.200 | 6.200 | 2.070 |
| Viral (< 0.22µm) | aPES | 5.667 | 13.100 | 1.850 |
| Viral (< 0.22µm) | aPES | 6.000 | 24.700 | 1.730 |
| Pico (> 0.22µm ) | aPES | 6.333 | 50.800 | 1.850 |
| Pico (> 0.22µm ) | aPES | 6.500 | 28.600 | 1.870 |
| Viral (< 0.22µm) | aPES | 3.000 | 20.200 | 1.880 |
| Viral (< 0.22µm) | aPES | 6.167 | 34.900 | 1.890 |
| Viral (< 0.22µm) | aPES | 3.333 | 18.500 | 1.780 |
| Pico (> 0.22µm ) | aPES | 6.667 | 51.800 | 1.830 |
| Pico (> 0.22µm ) | aPES | 6.667 | 57.800 | 1.850 |
| Viral (< 0.22µm) | aPES | 4.000 | 56.100 | 1.810 |
| Viral (< 0.22µm) | aPES | 4.000 | 16.000 | 1.780 |

| A260/A230 | NanoDrop Total Amc | Qubit Kit used | Qubit Yield (ng/ul) | Qubit Total Amount ( |
|---|---|---|---|---|
| 0.150 | 432.400 | 1X dsDNA HS kit | 16.500 | 1551.000 |
| 0.050 | 230.400 | 1X dsDNA HS kit | Below Detection | Below Detection |
| 0.130 | 364.800 | 1X dsDNA HS kit | 0.590 | 56.640 |
| 0.440 | 864.800 | 1X dsDNA HS kit | 5.240 | 492.560 |
| 0.010 | 432.000 | 1X dsDNA HS kit | 7.310 | 701.760 |
| 0.020 | 499.200 | 1X dsDNA HS kit | 2.930 | 281.280 |
| 0.020 | 230.400 | 1X dsDNA HS kit | 1.510 | 144.960 |
| 2.130 | 2455.000 | 1X dsDNA HS kit | 3.230 | 161.500 |
| 0.990 | 1380.000 | 1X dsDNA HS kit | 2.020 | 101.000 |
| 0.730 | 1260.000 | 1X dsDNA HS kit | 1.990 | 99.500 |
| 0.900 | 775.000 | 1X dsDNA HS kit | 103.000 | 5150.000 |
| 0.970 | 475.000 | 1X dsDNA HS kit | 69.200 | 3460.000 |
| 0.490 | 590.000 | 1X dsDNA HS kit | 81.700 | 4085.000 |
| 0.450 | 230.000 | 1X dsDNA HS kit | 25.800 | 1290.000 |
| 0.910 | 285.000 | 1X dsDNA HS kit | 32.200 | 1610.000 |
| 0.250 | 275.000 | 1X dsDNA HS kit | 19.200 | 960.000 |
| 0.160 | 235.000 | 1X dsDNA HS kit | 21.800 | 1090.000 |
| 0.570 | 200.000 | 1X dsDNA HS kit | 11.600 | 580.000 |
| 0.170 | 190.000 | 1X dsDNA HS kit | 11.200 | 560.000 |
| 0.040 | 180.000 | 1X dsDNA HS kit | 11.500 | 575.000 |
| 0.850 | 964.000 | 1X dsDNA HS kit | 18.500 | 740.000 |
| 0.520 | 636.000 | 1X dsDNA HS kit | 11.500 | 460.000 |
| 0.560 | 852.000 | 1X dsDNA HS kit | 71.500 | 2860.000 |
| 0.360 | 516.000 | 1X dsDNA HS kit | 77.900 | 3116.000 |
| 0.240 | 348.000 | 1X dsDNA HS kit | 38.800 | 1552.000 |
| 0.270 | 372.000 | 1X dsDNA HS kit | 54.300 | 2172.000 |
| 0.560 | 1124.000 | 1X dsDNA HS kit | 48.900 | 1956.000 |
| 0.550 | 1475.000 | 1X dsDNA HS kit | 31.200 | 1560.000 |
| 0.100 | 240.000 | 1X dsDNA HS kit | 36.100 | 1805.000 |
| 0.140 | 135.000 | 1X dsDNA HS kit | 9.890 | 494.500 |
| 0.120 | 150.000 | 1X dsDNA HS kit | 10.600 | 530.000 |
| 0.560 | 930.000 | 1X dsDNA HS kit | 6.040 | 302.000 |
| 0.150 | 120.000 | 1X dsDNA HS kit | 9.840 | 492.000 |
| 0.570 | 1630.000 | 1X dsDNA HS kit | 7.330 | 366.500 |
| 1.510 | 3830.000 | 1X dsDNA HS kit | 5.000 | 250.000 |
| n/a | n/a | 1X dsDNA HS kit | 43.700 | 4195.200 |
| n/a | n/a | 1X dsDNA HS kit | 71.300 | 6844.800 |
| n/a | n/a | 1X dsDNA HS kit | 79.900 | 7670.400 |
| n/a | n/a | 1X dsDNA HS kit | 32.900 | 3158.400 |
| n/a | n/a | 1X dsDNA HS kit | 43.000 | 4128.000 |
| n/a | n/a | 1X dsDNA HS kit | 33.400 | 3206.400 |
| n/a | n/a | 1X dsDNA HS kit | 216.000 | 20736.000 |
| n/a | n/a | 1X dsDNA HS kit | 171.000 | 16416.000 |
| n/a | n/a | 1X dsDNA HS kit | 45.400 | 4358.400 |
| n/a | n/a | 1X dsDNA HS kit | 67.300 | 6460.800 |
| n/a | n/a | 1X dsDNA HS kit | 39.100 | 3753.600 |

| | | | | | |
|---|---|---|---|---|---|
| n/a | n/a | | 1X dsDNA HS kit | 44.200 | 4243.200 |
| n/a | n/a | | 1X dsDNA HS kit | 23.600 | 2265.600 |
| | 0.080 | 2870.400 | 1X dsDNA HS kit | 15.100 | 1449.600 |
| | 0.140 | 1651.200 | 1X dsDNA HS kit | 11.900 | 1142.400 |
| | 0.040 | 1314.720 | 1X dsDNA HS kit | 11.200 | 887.040 |
| | 0.230 | 3139.200 | 1X dsDNA HS kit | 24.900 | 2390.400 |
| | 0.280 | 1612.800 | 1X dsDNA HS kit | 11.300 | 1084.800 |
| | 0.070 | 891.000 | 1X dsDNA HS kit | 6.050 | 544.500 |
| | 0.050 | 874.200 | 1X dsDNA HS kit | 5.050 | 474.700 |
| | 0.130 | 1116.000 | 1X dsDNA HS kit | 7.950 | 715.500 |
| | 0.070 | 1746.000 | 1X dsDNA HS kit | 12.800 | 1152.000 |
| | 0.050 | 1647.000 | 1X dsDNA HS kit | 11.100 | 999.000 |
| | 0.070 | 1012.000 | 1X dsDNA HS kit | 6.400 | 563.200 |
| | 0.080 | 1328.800 | 1X dsDNA HS kit | 6.000 | 528.000 |
| | 0.160 | 1100.000 | 1X dsDNA HS kit | 4.180 | 367.840 |
| | 1.210 | 1184.400 | 1X dsDNA HS kit | 4.410 | 414.540 |
| | 2.110 | 1128.000 | 1X dsDNA HS kit | 2.600 | 244.400 |
| | 1.560 | 1015.200 | 1X dsDNA HS kit | 2.950 | 277.300 |
| | 0.150 | 733.200 | 1X dsDNA HS kit | 5.500 | 517.000 |
| | 0.030 | 357.200 | 1X dsDNA HS kit | 1.480 | 139.120 |
| | 0.030 | 535.800 | 1X dsDNA HS kit | 3.780 | 355.320 |
| | 0.130 | 930.600 | 1X dsDNA HS kit | 7.650 | 719.100 |
| | 0.050 | 347.800 | 1X dsDNA HS kit | 2.740 | 257.560 |
| | 0.040 | 1259.600 | 1X dsDNA HS kit | 4.860 | 456.840 |
| | 0.070 | 1109.200 | 1X dsDNA HS kit | 8.200 | 770.800 |
| | 0.040 | 639.200 | 1X dsDNA HS kit | 3.400 | 319.600 |
| | 0.060 | 376.000 | 1X dsDNA HS kit | 2.310 | 217.140 |
| | 0.050 | 385.400 | 1X dsDNA HS kit | 2.180 | 204.920 |
| | 0.100 | 911.800 | 1X dsDNA HS kit | 3.390 | 318.660 |
| | 0.070 | 799.000 | 1X dsDNA HS kit | 4.340 | 407.960 |
| | 0.020 | 770.800 | 1X dsDNA HS kit | 3.840 | 360.960 |
| | 0.140 | 1231.400 | 1X dsDNA HS kit | 9.050 | 850.700 |
| | 0.040 | 752.000 | 1X dsDNA HS kit | 4.740 | 445.560 |
| | 0.120 | 1692.000 | 1X dsDNA HS kit | 2.950 | 277.300 |
| | 0.120 | 714.400 | 1X dsDNA HS kit | 4.210 | 395.740 |
| | 0.160 | 1231.400 | 1X dsDNA HS kit | 9.550 | 897.700 |
| | 0.150 | 1391.200 | 1X dsDNA HS kit | 12.400 | 1165.600 |
| | 0.120 | 1917.600 | 1X dsDNA HS kit | 7.400 | 695.600 |
| | 0.070 | 535.800 | 1X dsDNA HS kit | 2.870 | 269.780 |
| | 0.060 | 592.200 | 1X dsDNA HS kit | 4.340 | 407.960 |
| | 0.320 | 1222.000 | 1X dsDNA HS kit | 10.300 | 968.200 |
| | 0.480 | 1203.200 | 1X dsDNA HS kit | 9.650 | 907.100 |
| | 0.320 | 611.000 | 1X dsDNA HS kit | 4.780 | 449.320 |
| | 0.120 | 1851.800 | 1X dsDNA HS kit | 13.600 | 1278.400 |
| | 0.070 | 1419.400 | 1X dsDNA HS kit | 10.200 | 958.800 |
| | 0.190 | 1955.200 | 1X dsDNA HS kit | 11.000 | 1034.000 |
| | 0.080 | 2350.000 | 1X dsDNA HS kit | 17.600 | 1654.400 |

| | | | | |
|---|---|---|---|---|
| 0.080 | 1701.400 | 1X dsDNA HS kit | 11.400 | 1071.600 |
| 0.050 | 1522.800 | 1X dsDNA HS kit | 6.850 | 643.900 |
| 0.050 | 676.800 | 1X dsDNA HS kit | 3.840 | 360.960 |
| 1.260 | 7488.400 | 1X dsDNA HS kit | 10.800 | 2095.200 |
| 2.410 | 2871.200 | 1X dsDNA HS kit | 23.400 | 4539.600 |
| 2.300 | 15481.200 | 1X dsDNA HS kit | 17.600 | 3414.400 |
| 0.170 | 1955.200 | 1X dsDNA HS kit | 11.200 | 1052.800 |
| 0.060 | 733.200 | 1X dsDNA HS kit | 3.560 | 334.640 |
| 0.260 | 2942.200 | 1X dsDNA HS kit | 2.860 | 268.840 |
| 1.020 | 3638.400 | 1X dsDNA HS kit | 25.000 | 2400.000 |
| 0.960 | 3542.400 | 1X dsDNA HS kit | 17.400 | 1670.400 |
| 1.610 | 4166.400 | 1X dsDNA HS kit | 26.500 | 2544.000 |
| 0.320 | 1200.000 | 1X dsDNA HS kit | 5.550 | 532.800 |
| 0.710 | 9062.400 | 1X dsDNA HS kit | Above Detection | Above Detection |
| 1.370 | 15453.600 | 1X dsDNA HS kit | Above Detection | Above Detection |
| 0.650 | 14673.400 | 1X dsDNA HS kit | Above Detection | Above Detection |
| 0.780 | 16111.600 | 1X dsDNA HS kit | Above Detection | Above Detection |
| 0.770 | 4718.800 | 1X dsDNA HS kit | 7.900 | 742.600 |
| 0.700 | 5828.000 | 1X dsDNA HS kit | 9.650 | 907.100 |
| 0.610 | 4098.400 | 1X dsDNA HS kit | 6.750 | 634.500 |
| 0.510 | 770.800 | 1X dsDNA HS kit | 0.457 | 42.958 |
| 0.400 | 15196.800 | 1X dsDNA HS kit | Above Detection | Above Detection |
| 0.520 | 13344.000 | 1X dsDNA HS kit | Above Detection | Above Detection |
| 0.290 | 11164.800 | 1X dsDNA HS kit | Above Detection | Above Detection |
| 0.030 | 931.200 | 1X dsDNA HS kit | 0.479 | 45.984 |
| 0.040 | 1065.600 | 1X dsDNA HS kit | 4.530 | 434.880 |
| 0.070 | 806.400 | 1X dsDNA HS kit | 3.270 | 313.920 |
| 0.020 | 1075.200 | 1X dsDNA HS kit | 4.720 | 453.120 |
| 0.040 | 844.800 | 1X dsDNA HS kit | 4.430 | 425.280 |
| 0.060 | 1440.000 | 1X dsDNA HS kit | 6.080 | 583.680 |
| 0.060 | 1094.400 | 1X dsDNA HS kit | 6.840 | 656.640 |
| 0.410 | 1449.600 | 1X dsDNA HS kit | 7.910 | 759.360 |
| 0.030 | 739.200 | 1X dsDNA HS kit | 1.880 | 180.480 |
| 0.040 | 489.600 | 1X dsDNA HS kit | 0.913 | 87.648 |
| 0.100 | 1065.600 | 1X dsDNA HS kit | 0.190 | 18.240 |
| 0.100 | 796.800 | 1X dsDNA HS kit | 0.907 | 87.072 |
| 0.100 | 2342.400 | 1X dsDNA HS kit | 15.500 | 1488.000 |
| 0.020 | 604.800 | 1X dsDNA HS kit | 0.051 | 4.896 |
| 0.030 | 921.600 | 1X dsDNA HS kit | 1.040 | 99.840 |
| 0.020 | 681.600 | 1X dsDNA HS kit | 1.400 | 134.400 |
| 0.130 | 979.200 | 1X dsDNA HS kit | 5.030 | 482.880 |
| 0.050 | 998.400 | 1X dsDNA HS kit | 4.820 | 462.720 |
| 0.230 | 1449.600 | 1X dsDNA HS kit | 7.840 | 752.640 |
| 0.590 | 1622.400 | 1X dsDNA HS kit | 8.310 | 797.760 |
| 0.050 | 1200.000 | 1X dsDNA HS kit | 5.960 | 572.160 |
| 0.420 | 883.200 | 1X dsDNA HS kit | 0.150 | 14.400 |
| 0.690 | 940.800 | 1X dsDNA HS kit | 2.760 | 264.960 |

| | | | | |
|---|---|---|---|---|
| 0.150 | 835.200 | 1X dsDNA HS kit | 1.380 | 132.480 |
| 0.340 | 796.800 | 1X dsDNA HS kit | 1.990 | 191.040 |
| 0.130 | 1276.800 | 1X dsDNA HS kit | 4.900 | 470.400 |
| 0.110 | 1219.200 | 1X dsDNA HS kit | 5.440 | 522.240 |
| 0.410 | 1718.400 | 1X dsDNA HS kit | 10.100 | 969.600 |
| 0.080 | 1488.000 | 1X dsDNA HS kit | 9.150 | 878.400 |
| 0.240 | 1142.400 | 1X dsDNA HS kit | 5.380 | 516.480 |
| 0.130 | 758.400 | 1X dsDNA HS kit | 0.838 | 80.448 |
| 0.180 | 1094.400 | 1X dsDNA HS kit | 4.180 | 401.280 |
| 1.550 | 4329.600 | 1X dsDNA HS kit | 42.700 | 4099.200 |
| 1.510 | 4723.200 | 1X dsDNA HS kit | 39.600 | 3801.600 |
| 0.460 | 5251.200 | 1X dsDNA HS kit | 46.700 | 4483.200 |
| 1.210 | 4665.600 | 1X dsDNA HS kit | 40.700 | 3907.200 |
| 0.170 | 624.000 | 1X dsDNA HS kit | 0.413 | 39.648 |
| 0.480 | 681.600 | 1X dsDNA HS kit | 4.350 | 417.600 |
| 0.380 | 1228.800 | 1X dsDNA HS kit | 8.450 | 811.200 |
| 1.860 | 7027.200 | 1X dsDNA HS kit | Above Detection | Above Detection |
| 1.680 | 6288.000 | 1X dsDNA HS kit | 38.900 | 3734.400 |
| 0.080 | 1619.900 | 1X dsDNA HS kit | 6.380 | 618.860 |
| 0.120 | 1687.800 | 1X dsDNA HS kit | 8.000 | 776.000 |
| 0.860 | 1493.800 | 1X dsDNA HS kit | 4.020 | 389.940 |
| 0.110 | 1678.100 | 1X dsDNA HS kit | 7.780 | 754.660 |
| 0.700 | 1978.800 | 1X dsDNA HS kit | 10.100 | 979.700 |
| 0.330 | 1697.500 | 1X dsDNA HS kit | 7.180 | 696.460 |
| 0.860 | 2124.300 | 1X dsDNA HS kit | 11.200 | 1086.400 |
| 0.170 | 2172.800 | 1X dsDNA HS kit | 7.780 | 754.660 |
| 1.370 | 2337.700 | 1X dsDNA HS kit | 14.500 | 1406.500 |
| 0.150 | 2308.600 | 1X dsDNA HS kit | 18.600 | 1804.200 |
| 1.140 | 2017.600 | 1X dsDNA HS kit | 8.560 | 830.320 |
| 0.840 | 2095.200 | 1X dsDNA HS kit | 10.900 | 1057.300 |
| 0.420 | 2095.200 | 1X dsDNA HS kit | 17.200 | 1668.400 |
| 0.230 | 2531.700 | 1X dsDNA HS kit | 10.600 | 1028.200 |
| 1.030 | 2143.700 | 1X dsDNA HS kit | 32.000 | 3104.000 |
| 0.950 | 3414.400 | 1X dsDNA HS kit | 6.920 | 671.240 |
| 1.450 | 2289.200 | 1X dsDNA HS kit | 24.400 | 2366.800 |
| 1.430 | 2783.900 | 1X dsDNA HS kit | 8.780 | 851.660 |
| 0.800 | 1755.700 | 1X dsDNA HS kit | 4.240 | 411.280 |
| 0.360 | 1047.600 | 1X dsDNA HS kit | 9.140 | 886.580 |
| 0.600 | 902.100 | 1X dsDNA HS kit | 6.560 | 636.320 |
| 99.000 | 1522.900 | 1X dsDNA HS kit | 0.438 | 42.486 |
| 0.140 | 1251.300 | 1X dsDNA HS kit | 1.880 | 182.360 |
| 0.050 | 1320.960 | 1X dsDNA HS kit | 8.240 | 708.640 |
| 0.050 | 1565.200 | 1X dsDNA HS kit | 12.100 | 1040.600 |
| 1.460 | 1376.000 | 1X dsDNA HS kit | 9.490 | 816.140 |
| 0.050 | 1367.400 | 1X dsDNA HS kit | 0.611 | 52.546 |
| 0.580 | 1436.200 | 1X dsDNA HS kit | 10.500 | 903.000 |
| 0.820 | 3868.800 | 1X dsDNA HS kit | 14.000 | 1344.000 |

| | | | | |
|---|---|---|---|---|
| 0.040 | 1574.400 | 1X dsDNA HS kit | 10.500 | 1008.000 |
| 0.220 | 1747.200 | 1X dsDNA HS kit | 11.100 | 1065.600 |
| 1.820 | 2966.400 | 1X dsDNA HS kit | 24.200 | 2323.200 |
| 0.400 | 3414.200 | 1X dsDNA HS kit | 22.700 | 1952.200 |
| 0.370 | 3018.600 | 1X dsDNA HS kit | 27.400 | 2356.400 |
| 0.060 | 963.200 | 1X dsDNA HS kit | 4.630 | 398.180 |
| 0.190 | 2150.000 | 1X dsDNA HS kit | 16.200 | 1393.200 |
| 1.000 | 2253.200 | 1X dsDNA HS kit | 16.900 | 1453.400 |
| 0.910 | 3921.600 | 1X dsDNA HS kit | 16.600 | 1427.600 |
| 2.140 | 2908.800 | 1X dsDNA HS kit | 21.300 | 2044.800 |
| 0.850 | 2918.400 | 1X dsDNA HS kit | 22.300 | 2140.800 |
| 0.570 | 2572.800 | 1X dsDNA HS kit | 19.000 | 1824.000 |
| 0.430 | 2611.200 | 1X dsDNA HS kit | 19.700 | 1891.200 |
| 0.650 | 2400.000 | 1X dsDNA HS kit | 17.300 | 1660.800 |
| 0.180 | 2822.400 | 1X dsDNA HS kit | 17.100 | 1641.600 |
| 0.700 | 2688.000 | 1X dsDNA HS kit | 17.300 | 1660.800 |
| 0.860 | 3004.800 | 1X dsDNA HS kit | 20.200 | 1939.200 |
| 0.900 | 3129.600 | 1X dsDNA HS kit | 19.000 | 1824.000 |
| 0.010 | 374.400 | 1X dsDNA HS kit | 1.200 | 115.200 |
| 0.010 | 480.000 | 1X dsDNA HS kit | 2.510 | 240.960 |
| 0.010 | 249.600 | 1X dsDNA HS kit | 0.794 | 76.224 |
| 0.010 | 345.600 | 1X dsDNA HS kit | 0.724 | 69.504 |
| 0.010 | 432.000 | 1X dsDNA HS kit | 0.899 | 86.304 |
| 0.480 | 610.600 | 1X dsDNA HS kit | 2.000 | 172.000 |
| 0.510 | 696.600 | 1X dsDNA HS kit | 1.600 | 137.600 |
| 0.510 | 722.400 | 1X dsDNA HS kit | 1.160 | 99.760 |
| 0.290 | 954.600 | 1X dsDNA HS kit | 5.620 | 483.320 |
| 0.530 | 739.600 | 1X dsDNA HS kit | 3.000 | 258.000 |
| 0.460 | 777.600 | 1X dsDNA HS kit | 2.750 | 264.000 |
| 0.400 | 998.400 | 1X dsDNA HS kit | 3.240 | 311.040 |
| 0.650 | 844.800 | 1X dsDNA HS kit | 3.320 | 318.720 |
| 0.190 | 825.600 | 1X dsDNA HS kit | 3.600 | 345.600 |
| 0.110 | 691.200 | 1X dsDNA HS kit | 2.220 | 213.120 |
| 0.110 | 729.600 | 1X dsDNA HS kit | 0.290 | 27.840 |
| 0.130 | 643.200 | 1X dsDNA HS kit | 1.710 | 164.160 |
| 0.460 | 691.200 | 1X dsDNA HS kit | 1.020 | 97.920 |
| 0.380 | 559.000 | 1X dsDNA HS kit | 1.030 | 88.580 |
| 0.120 | 460.800 | 1X dsDNA HS kit | BD | BD |
| 0.380 | 412.800 | 1X dsDNA HS kit | 0.734 | 70.464 |
| 0.960 | 3542.400 | 1X dsDNA HS kit | 23.100 | 2217.600 |
| 0.400 | 643.200 | 1X dsDNA HS kit | 0.388 | 37.248 |
| 0.010 | 460.800 | 1X dsDNA HS kit | 1.620 | 155.520 |
| 0.020 | 528.000 | 1X dsDNA HS kit | 1.730 | 166.080 |
| 0.070 | 547.200 | 1X dsDNA HS kit | 1.280 | 122.880 |
| 0.060 | 595.200 | 1X dsDNA HS kit | 0.081 | 7.776 |
| 0.020 | 604.800 | 1X dsDNA HS kit | 0.870 | 83.520 |
| 0.040 | 566.400 | 1X dsDNA HS kit | Below Detection | Below Detection |

| | | | | |
|---|---|---|---|---|
| 0.410 | 355.200 | 1X dsDNA HS kit | 0.178 | 17.088 |
| 0.060 | 604.800 | 1X dsDNA HS kit | Below Detection | Below Detection |
| 0.090 | 566.400 | 1X dsDNA HS kit | Below Detection | Below Detection |
| 0.080 | 604.800 | 1X dsDNA HS kit | 0.099 | 9.504 |
| 0.080 | 633.600 | 1X dsDNA HS kit | Below Detection | Below Detection |
| 0.070 | 643.200 | 1X dsDNA HS kit | Below Detection | Below Detection |
| 0.100 | 576.000 | 1X dsDNA HS kit | 0.053 | 5.088 |
| 0.100 | 624.000 | 1X dsDNA HS kit | 0.374 | 35.904 |
| 0.030 | 825.600 | 1X dsDNA HS kit | 0.260 | 24.960 |
| 0.070 | 691.200 | 1X dsDNA HS kit | 0.098 | 9.408 |
| 0.090 | 950.400 | 1X dsDNA HS kit | 4.860 | 466.560 |
| 0.080 | 1084.800 | 1X dsDNA HS kit | 5.840 | 560.640 |
| 0.060 | 729.600 | 1X dsDNA HS kit | 1.570 | 150.720 |
| 0.010 | 700.800 | 1X dsDNA HS kit | 0.181 | 17.376 |
| 0.010 | 710.400 | 1X dsDNA HS kit | 0.111 | 10.656 |
| 0.130 | 7084.800 | 1X dsDNA HS kit | 53.000 | 5088.000 |
| 0.110 | 4262.400 | 1X dsDNA HS kit | 32.300 | 3100.800 |
| 0.210 | 3360.000 | 1X dsDNA HS kit | 24.700 | 2371.200 |
| 1.300 | 1666.000 | 1X dsDNA HS kit | 10.900 | 2136.400 |
| 1.390 | 2214.800 | 1X dsDNA HS kit | 3.930 | 770.280 |
| 1.440 | 2332.400 | 1X dsDNA HS kit | 4.070 | 797.720 |
| 1.760 | 1528.800 | 1X dsDNA HS kit | 2.790 | 546.840 |
| 1.560 | 1685.600 | 1X dsDNA HS kit | 2.920 | 572.320 |
| 1.680 | 5429.200 | 1X dsDNA HS kit | 4.880 | 956.480 |
| 1.590 | 6468.000 | 1X dsDNA HS kit | 6.430 | 1260.280 |
| 1.640 | 3175.200 | 1X dsDNA HS kit | 1.820 | 356.720 |
| 0.750 | 1372.000 | 1X dsDNA HS kit | 1.570 | 307.720 |
| 0.860 | 882.000 | 1X dsDNA HS kit | 1.270 | 248.920 |
| 1.490 | 1744.400 | 1X dsDNA HS kit | 2.400 | 470.400 |
| 1.490 | 7641.600 | 1X dsDNA HS kit | 3.480 | 668.160 |
| 1.450 | 5971.200 | 1X dsDNA HS kit | 4.270 | 819.840 |
| 1.460 | 4473.600 | 1X dsDNA HS kit | 2.910 | 558.720 |
| 1.380 | 3264.000 | 1X dsDNA HS kit | 3.050 | 585.600 |
| 1.430 | 4531.200 | 1X dsDNA HS kit | 2.420 | 464.640 |
| 1.260 | 2400.000 | 1X dsDNA HS kit | 3.510 | 673.920 |
| 1.640 | 5337.600 | 1X dsDNA HS kit | 7.060 | 1355.520 |
| 2.290 | 9446.400 | 1X dsDNA HS kit | 4.440 | 852.480 |
| 1.120 | 3187.200 | 1X dsDNA HS kit | 1.660 | 318.720 |
| 1.130 | 6624.800 | 1X dsDNA HS kit | 4.730 | 927.080 |
| 1.130 | 10995.600 | 1X dsDNA HS kit | 7.280 | 1426.880 |
| 1.000 | 2646.000 | 1X dsDNA HS kit | 3.000 | 588.000 |
| 1.250 | 3508.400 | 1X dsDNA HS kit | 3.520 | 689.920 |
| 1.190 | 6644.400 | 1X dsDNA HS kit | 6.280 | 1230.880 |
| 0.970 | 2116.800 | 1X dsDNA HS kit | 2.000 | 392.000 |
| 0.700 | 1411.200 | 1X dsDNA HS kit | 2.220 | 435.120 |
| 1.030 | 1156.400 | 1X dsDNA HS kit | 2.660 | 521.360 |
| 0.390 | 4174.800 | 1X dsDNA HS kit | 0.833 | 163.268 |

| | | | | |
|---|---|---|---|---|
| 1.200 | 2665.600 | 1X dsDNA HS kit | 3.670 | 719.320 |
| 1.440 | 8937.600 | 1X dsDNA HS kit | 6.040 | 1183.840 |
| 0.560 | 1626.800 | 1X dsDNA HS kit | 0.898 | 176.008 |
| 1.020 | 2077.600 | 1X dsDNA HS kit | 0.571 | 111.916 |
| 0.480 | 1097.600 | 1X dsDNA HS kit | 0.280 | 54.880 |
| 0.670 | 2097.200 | 1X dsDNA HS kit | 0.846 | 165.816 |
| 0.700 | 1920.800 | 1X dsDNA HS kit | 2.090 | 409.640 |
| 0.870 | 2175.600 | 1X dsDNA HS kit | 2.370 | 464.520 |
| 0.490 | 1979.600 | 1X dsDNA HS kit | 1.240 | 243.040 |
| 0.570 | 1783.600 | 1X dsDNA HS kit | 1.610 | 315.560 |
| 1.050 | 2606.800 | 1X dsDNA HS kit | 2.140 | 419.440 |
| 0.590 | 1568.000 | 1X dsDNA HS kit | 1.000 | 196.000 |
| 0.560 | 1783.600 | 1X dsDNA HS kit | 0.994 | 194.824 |
| 1.090 | 3684.800 | 1X dsDNA HS kit | 2.250 | 441.000 |
| 0.410 | 1117.200 | 1X dsDNA HS kit | 0.105 | 20.580 |
| 0.430 | 6017.200 | 1X dsDNA HS kit | 4.480 | 878.080 |
| 0.100 | Below Detection | 1X dsDNA HS kit | 1.930 | 374.420 |
| 0.450 | 2587.200 | 1X dsDNA HS kit | 1.820 | 356.720 |
| 0.300 | 3116.400 | 1X dsDNA HS kit | 2.640 | 517.440 |
| 0.770 | 14464.800 | 1X dsDNA HS kit | 8.580 | 1681.680 |
| 0.630 | 6370.000 | 1X dsDNA HS kit | 9.750 | 1911.000 |
| 0.420 | 5233.200 | 1X dsDNA HS kit | 3.190 | 625.240 |
| 0.660 | 2900.800 | 1X dsDNA HS kit | 1.990 | 390.040 |
| 0.550 | 5723.200 | 1X dsDNA HS kit | 4.280 | 838.880 |
| 0.700 | 4037.600 | 1X dsDNA HS kit | 2.770 | 542.920 |
| 0.520 | 4723.600 | 1X dsDNA HS kit | 4.470 | 876.120 |
| 1.040 | 7526.400 | 1X dsDNA HS kit | 7.420 | 1454.320 |
| 0.720 | 22461.600 | 1X dsDNA HS kit | 0.220 | 43.120 |
| 1.180 | 5390.000 | 1X dsDNA HS kit | 0.085 | 16.660 |
| 0.850 | 9917.600 | 1X dsDNA HS kit | 0.099 | 19.404 |
| 1.060 | 4998.000 | 1X dsDNA HS kit | 0.134 | 26.264 |
| 0.180 | 910.800 | 1X dsDNA HS kit | 1.770 | 81.420 |
| 0.190 | 602.600 | 1X dsDNA HS kit | 5.500 | 253.000 |
| 0.220 | 492.200 | 1X dsDNA HS kit | 0.242 | 11.132 |
| 0.250 | 4444.800 | 1X dsDNA HS kit | 38.200 | 3667.200 |
| 0.450 | 3974.400 | 1X dsDNA HS kit | 32.700 | 3139.200 |
| 0.540 | 4406.400 | 1X dsDNA HS kit | 35.300 | 3388.800 |
| 0.100 | 827.200 | 1X dsDNA HS kit | Below Detection | Below Detection |
| 0.010 | 182.400 | 1X dsDNA HS kit | Below Detection | Below Detection |
| 0.010 | 163.200 | 1X dsDNA HS kit | Below Detection | Below Detection |
| 0.020 | 537.600 | 1X dsDNA HS kit | 2.730 | 262.080 |
| 0.020 | 451.200 | 1X dsDNA HS kit | 3.920 | 376.320 |
| 0.040 | 672.000 | 1X dsDNA HS kit | 5.280 | 506.880 |
| 0.040 | 969.600 | 1X dsDNA HS kit | 5.220 | 501.120 |
| 0.030 | 902.400 | 1X dsDNA HS kit | 1.260 | 120.960 |
| 0.050 | 1238.400 | 1X dsDNA HS kit | 9.000 | 864.000 |
| 0.100 | 1267.200 | 1X dsDNA HS kit | 10.300 | 988.800 |

| | | | | |
|---|---|---|---|---|
| 0.070 | 1430.400 | 1X dsDNA HS kit | 9.710 | 932.160 |
| 0.110 | 1689.600 | 1X dsDNA HS kit | 10.100 | 969.600 |
| 0.390 | 1910.400 | 1X dsDNA HS kit | 16.600 | 1593.600 |
| 0.490 | 2649.600 | 1X dsDNA HS kit | 22.200 | 2131.200 |
| 0.170 | 1161.600 | 1X dsDNA HS kit | 9.060 | 869.760 |
| 0.020 | 211.200 | 1X dsDNA HS kit | 0.971 | 93.216 |
| 0.010 | 144.000 | 1X dsDNA HS kit | 0.339 | 32.544 |
| 0.080 | 240.000 | 1X dsDNA HS kit | 1.310 | 125.760 |
| 0.010 | 76.800 | 1X dsDNA HS kit | 0.388 | 37.248 |
| 0.060 | 374.400 | 1X dsDNA HS kit | 1.860 | 178.560 |
| 0.070 | 393.600 | 1X dsDNA HS kit | 2.600 | 249.600 |
| 0.600 | 355.200 | 1X dsDNA HS kit | 2.180 | 209.280 |
| 0.060 | 272.600 | 1X dsDNA HS kit | 1.300 | 122.200 |
| 0.540 | 288.000 | 1X dsDNA HS kit | 1.760 | 168.960 |
| 0.030 | 57.600 | 1X dsDNA HS kit | 0.388 | 37.248 |
| 0.280 | 211.200 | 1X dsDNA HS kit | 0.848 | 81.408 |
| 0.040 | 134.400 | 1X dsDNA HS kit | 0.743 | 71.328 |
| 1.520 | 403.200 | 1X dsDNA HS kit | 1.920 | 184.320 |
| 0.140 | 412.800 | 1X dsDNA HS kit | 2.150 | 206.400 |
| 0.470 | 460.800 | 1X dsDNA HS kit | 3.060 | 293.760 |
| 0.190 | 326.400 | 1X dsDNA HS kit | 1.870 | 179.520 |
| 0.030 | 105.600 | 1X dsDNA HS kit | 0.249 | 23.904 |
| 0.220 | 105.600 | 1X dsDNA HS kit | 0.481 | 46.176 |
| 0.290 | 124.800 | 1X dsDNA HS kit | 0.338 | 32.448 |
| 0.080 | 76.800 | 1X dsDNA HS kit | 0.318 | 30.528 |
| 1.220 | 787.200 | 1X dsDNA HS kit | 5.660 | 543.360 |
| 0.350 | 825.600 | 1X dsDNA HS kit | 5.560 | 533.760 |
| 0.510 | 1027.200 | 1X dsDNA HS kit | 7.750 | 744.000 |
| 0.730 | 960.000 | 1X dsDNA HS kit | 7.890 | 757.440 |
| 0.300 | 528.000 | 1X dsDNA HS kit | 3.350 | 321.600 |
| 0.150 | 508.800 | 1X dsDNA HS kit | 2.370 | 227.520 |
| 0.400 | 652.800 | 1X dsDNA HS kit | 3.140 | 301.440 |
| 0.490 | 547.200 | 1X dsDNA HS kit | 2.990 | 287.040 |
| 0.010 | 172.800 | 1X dsDNA HS kit | Below Detection | Below Detection |
| 0.000 | 76.800 | 1X dsDNA HS kit | Below Detection | Below Detection |
| 0.000 | 67.200 | 1X dsDNA HS kit | Below Detection | Below Detection |
| 0.010 | 144.000 | 1X dsDNA HS kit | Below Detection | Below Detection |
| 0.010 | 105.600 | 1X dsDNA HS kit | Below Detection | Below Detection |
| 0.010 | 67.200 | 1X dsDNA HS kit | Below Detection | Below Detection |
| 0.090 | 86.400 | 1X dsDNA HS kit | Below Detection | Below Detection |
| 0.220 | 3494.400 | 1X dsDNA HS kit | 24.500 | 1127.000 |
| 0.410 | 3100.800 | 1X dsDNA HS kit | 23.500 | 2256.000 |
| 0.060 | 2764.800 | 1X dsDNA HS kit | 16.900 | 1622.400 |
| 0.010 | 240.000 | 1X dsDNA HS kit | 0.512 | 49.152 |
| 0.110 | 4300.800 | 1X dsDNA HS kit | 33.100 | 3177.600 |
| 0.200 | 4560.000 | 1X dsDNA HS kit | 34.100 | 3273.600 |
| 0.010 | 115.200 | 1X dsDNA HS kit | Below Detection | Below Detection |

| | | | | |
|---|---|---|---|---|
| 0.010 | 96.000 | 1X dsDNA HS kit | Below Detection | Below Detection |
| 0.070 | 2323.200 | 1X dsDNA HS kit | 19.300 | 1852.800 |
| 0.050 | 1526.400 | 1X dsDNA HS kit | 12.400 | 1190.400 |
| 0.220 | 2112.000 | 1X dsDNA HS kit | 17.900 | 1718.400 |
| 0.160 | 2083.200 | 1X dsDNA HS kit | 19.300 | 1852.800 |
| 0.130 | 1180.800 | 1X dsDNA HS kit | 7.480 | 718.080 |
| 0.080 | 681.600 | 1X dsDNA HS kit | 4.280 | 410.880 |
| 0.010 | 336.000 | 1X dsDNA HS kit | 1.250 | 120.000 |
| 0.150 | 652.800 | 1X dsDNA HS kit | 3.810 | 365.760 |
| 0.040 | 441.600 | 1X dsDNA HS kit | 1.770 | 169.920 |
| 0.030 | 556.800 | 1X dsDNA HS kit | 0.483 | 46.368 |
| 0.030 | 662.400 | 1X dsDNA HS kit | 2.950 | 283.200 |
| 0.010 | 393.600 | 1X dsDNA HS kit | 1.090 | 104.640 |
| 0.050 | 470.400 | 1X dsDNA HS kit | 2.990 | 287.040 |
| 0.030 | 883.200 | 1X dsDNA HS kit | 4.320 | 414.720 |
| 0.030 | 595.200 | 1X dsDNA HS kit | 2.860 | 274.560 |
| 0.380 | 1257.600 | 1X dsDNA HS kit | 12.200 | 1171.200 |
| 0.130 | 2371.200 | 1X dsDNA HS kit | 8.210 | 788.160 |
| 0.220 | 4876.800 | 1X dsDNA HS kit | 30.800 | 2956.800 |
| 0.080 | 2745.600 | 1X dsDNA HS kit | 13.500 | 1296.000 |
| 0.040 | 1939.200 | 1X dsDNA HS kit | 12.300 | 1180.800 |
| 0.050 | 3350.400 | 1X dsDNA HS kit | 24.000 | 2304.000 |
| 0.100 | 1776.000 | 1X dsDNA HS kit | 10.000 | 960.000 |
| 0.380 | 4972.800 | 1X dsDNA HS kit | 35.800 | 3436.800 |
| 0.230 | 5433.200 | 1X dsDNA HS kit | 36.400 | 3421.600 |
| 0.410 | 5273.400 | 1X dsDNA HS kit | 43.300 | 4070.200 |
| 0.110 | 1504.000 | 1X dsDNA HS kit | 10.700 | 1005.800 |

(ng)
