## Supplementary Data 4 for "Sampling Microbial Dynamics in the Salish Sea Estuary: Evaluating Methods to Capture Cyanobacteria and Cyanophage"

| Parent ID | Date of Sampling | Time in Fre | Extraction | Subsample Sequence ID |
|---|---|---|---|---|
| NW_FE_0( | 04/15/24 | 3 | PSK | 20240418-PNNL-NRB-Seq11 |
| NW_FE_0( | 04/15/24 | 3 | PSK | 20240418-PNNL-NRB-Seq13 |
| NW_FE_0( | 05/08/24 | 5 | PSK | 20240513-PNNL-NRB-Seq45 |
| NW_FE_0( | 05/08/24 | 5 | PSK | 20240513-PNNL-NRB-Seq49 |
| NW_FE_0( | 05/08/24 | 5 | PSK | 20240513-PNNL-NRB-Seq50 |
| NW_FE_01 | 07/17/24 | 1 | PSK | 20240718-PNNL-NRB-Seq58 |
| NW_FE_01 | 07/17/24 | 1 | PSK | 20240718-PNNL-NRB-Seq59 |
| NW_FE_01 | 07/23/24 | 13 | PSK | 20240805-PNNL-MPF-Seq75 |
| NW_FE_01 | 07/23/24 | 13 | PSK | 20240805-PNNL-MPF-Seq76 |
| NW_FE_01 | 07/31/24 | 5 | PSK | 20240805-PNNL-MPF-Seq77 |
| NW_FE_01 | 07/31/24 | 5 | PSK | 20240805-PNNL-MPF-Seq78 |
| NW_FE_01 | 07/31/24 | 5 | PSK | 20240805-PNNL-MPF-Seq79 |
| NW_FE_01 | 07/31/24 | 5 | PSK | 20240805-PNNL-MPF-Seq80 |
| NW_FE_01 | 07/23/24 | 13 | PSK | 20240805-PNNL-MPF-Seq82 |
| NW_FE_01 | 07/23/24 | 13 | PSK | 20240805-PNNL-MPF-Seq83 |
| NW_FE_01 | 07/23/24 | 13 | PSK | 20240805-PNNL-MPF-Seq84 |
| NW_FE_01 | 07/23/24 | 13 | PSK | 20240805-PNNL-MPF-Seq85 |
| NW_FE_01 | 07/23/24 | 13 | PSK | 20240805-PNNL-MPF-Seq86 |
| NW_FE_01 | 07/31/24 | 6 | PSK | 20240806-PNNL-MPF-Seq94 |
| NW_FE_01 | 07/31/24 | 6 | PSK | 20240806-PNNL-MPF-Seq95 |
| NW_FE_01 | 07/31/24 | 7 | PSK | 20240807-PNNL-MPF-Seq102 |
| NW_FE_01 | 07/31/24 | 7 | PSK | 20240807-PNNL-MPF-Seq103 |
| NW_FE_01 | 07/31/24 | 7 | PSK | 20240807-PNNL-MPF-Seq104 |
| NW_FE_01 | 07/31/24 | 7 | PSK | 20240807-PNNL-MPF-Seq105 |
| NW_FE_01 | 07/31/24 | 7 | PSK | 20240807-PNNL-MPF-Seq106 |
| NW_FE_01 | 07/31/24 | 7 | PSK | 20240807-PNNL-MPF-Seq107 |
| NW_FE_01 | 07/31/24 | 7 | PSK | 20240807-PNNL-MPF-Seq108 |
| NW_FE_02 | 08/26/24 | 3 | PSK | 20240829-PNNL-MPF-Seq114 |
| NW_FE_02 | 08/26/24 | 9 | PWK | 20240829-PNNL-MPF-Seq119 |
| NW_FE_02 | 08/26/24 | 9 | PWK | 20240829-PNNL-MPF-Seq120 |
| NW_FE_02 | 08/27/24 | 8 | PWK | 20240829-PNNL-MPF-Seq121 |
| NW_FE_02 | 08/27/24 | 8 | PWK | 20240829-PNNL-MPF-Seq122 |
| NW_FE_02 | 09/09/24 | 2 | PWK | 20240911-PNNL-MPF-Seq124 |
| NW_FE_02 | 09/09/24 | 2 | PWK | 20240911-PNNL-MPF-Seq125 |
| NW_FE_02 | 09/09/24 | 2 | PWK | 20240911-PNNL-MPF-Seq126 |
| NW_FE_02 | 09/09/24 | 58 | PSK | 20241106-PNNL-NRB-Seq133 |
| NW_FE_02 | 09/09/24 | 58 | PSK | 20241106-PNNL-NRB-Seq134 |
| NW_FE_02 | 09/09/24 | 58 | PSK | 20241106-PNNL-NRB-Seq135 |
| NW_FE_01 | 07/31/24 | 98 | PSK | 20241106-PNNL-NRB-Seq136 |
| NW_FE_01 | 07/31/24 | 98 | PSK | 20241106-PNNL-NRB-Seq137 |
| NW_FE_01 | 07/31/24 | 98 | PSK | 20241106-PNNL-NRB-Seq138 |
| NW_FE_01 | 07/31/24 | 98 | PSK | 20241106-PNNL-NRB-Seq142 |
| NW_FE_01 | 07/31/24 | 98 | PSK | 20241106-PNNL-NRB-Seq143 |
| NW_FE_01 | 07/31/24 | 98 | PSK | 20241106-PNNL-NRB-Seq144 |
| NW_FE_02 | 10/29/24 | 8 | PSK | 20241106-PNNL-NRB-Seq148 |
| NW_FE_01 | 08/07/24 | 103 | PWK | 20241118-PNNL-NRB-Seq168 |

| | | | | |
|---|---|---|---|---|
| NW_FE_01 | 08/07/24 | 103 | PWK | 20241118-PNNL-NRB-Seq169 |
| NW_FE_01 | 08/07/24 | 103 | PWK | 20241118-PNNL-NRB-Seq170 |
| NW_FE_01 | 08/07/24 | 103 | PWK | 20241118-PNNL-NRB-Seq171 |
| NW_FE_01 | 08/07/24 | 103 | PWK | 20241118-PNNL-NRB-Seq172 |
| NW_FE_01 | 08/14/24 | 96 | PWK | 20241118-PNNL-NRB-Seq174 |
| NW_FE_01 | 08/14/24 | 96 | PWK | 20241118-PNNL-NRB-Seq175 |
| NW_FE_01 | 08/14/24 | 96 | PWK | 20241118-PNNL-NRB-Seq176 |
| NW_FE_01 | 08/07/24 | 103 | PWK | 20241118-PNNL-NRB-Seq173 |
| NW_FE_02 | 10/29/24 | 8 | PSK | 20241106-PNNL-NRB-Seq149 |
| NW_FE_02 | 10/29/24 | 8 | PSK | 20241106-PNNL-NRB-Seq150 |
| NW_FE_02 | 10/29/24 | 8 | PSK | 20241106-PNNL-NRB-Seq151 |
| NW_FE_02 | 10/29/24 | 8 | PSK | 20241106-PNNL-NRB-Seq152 |
| NW_FE_02 | 10/29/24 | 8 | PSK | 20241106-PNNL-NRB-Seq153 |

| Sampling Location | Tidal Stage | Filter Fraction | Filter Mate | Filter Used | Total volur |
|---|---|---|---|---|---|
| PNNL Dock | Ebb | Pico (> 0.22µm ) | MCE | BF-013 | 2 |
| PNNL Dock | Ebb | Pico (> 0.22µm ) | MCE | BF-015 | 1 |
| PNNL Dock | Flood | Pico (> 0.22µm ) | MCE | BF-022 | 2.1 |
| PNNL Dock | Flood | Pico (> 0.22µm ) | MCE | BF-026 | 2 |
| PNNL Dock | Flood | Pico (> 0.22µm ) | MCE | BF-027 | 1.3 |
| PNNL Dock | Flood | Pico (> 0.22µm ) | MCE | BF-064 | 1.8 |
| PNNL Dock | Flood | Pico (> 0.22µm ) | MCE | BF-065 | 1 |
| PNNL Dock | Ebb | Micro (> 5µm) | MCE | BF-067 | 5 |
| PNNL Dock | Ebb | Micro (> 5µm) | MCE | BF-069 | 5 |
| PNNL Dock | Ebb | Pico (> 0.22µm ) | MCE | BF-075 | 2 |
| PNNL Dock | Ebb | Pico (> 0.22µm ) | MCE | BF-076 | 2 |
| PNNL Dock | Ebb | Pico (> 0.22µm ) | MCE | BF-077 | 2 |
| PNNL Dock | Ebb | Pico (> 0.22µm ) | MCE | BF-078 | 2 |
| PNNL Dock | Ebb | Pico (> 0.22µm ) | MCE | BF-070 | 1.5 |
| PNNL Dock | Ebb | Pico (> 0.22µm ) | MCE | BF-071 | 1.5 |
| PNNL Dock | Ebb | Pico (> 0.22µm ) | MCE | BF-072 | 1.5 |
| PNNL Dock | Ebb | Pico (> 0.22µm ) | MCE | BF-073 | 1.5 |
| PNNL Dock | Ebb | Pico (> 0.22µm ) | MCE | BF-074 | 1.5 |
| JWM | Flood | Micro (> 5µm) | MCE | BF-087 | 2.5 |
| JWM | Flood | Micro (> 5µm) | MCE | BF-088 | 2.5 |
| JWM | Flood | Pico (> 0.22µm ) | MCE | BF-094 | 2 |
| JWM | Flood | Pico (> 0.22µm ) | MCE | BF-095 | 2 |
| JWM | Flood | Pico (> 0.22µm ) | MCE | BF-096 | 2 |
| JWM | Flood | Pico (> 0.22µm ) | MCE | BF-097 | 2 |
| PNNL Dock | Ebb | Micro (> 5µm) | MCE | BF-099 | 3 |
| PNNL Dock | Ebb | Micro (> 5µm) | MCE | BF-100 | 3.5 |
| PNNL Dock | Ebb | Micro (> 5µm) | MCE | BF-101 | 2.5 |
| PNNL Dock | Slack High | Pico (> 0.22µm ) | aPES | BF-172 | 20 |
| PNNL Dock | Slack High | Pico (> 0.22µm ) | aPES | BF-172 | 20 |
| PNNL Dock | Slack High | Pico (> 0.22µm ) | aPES | BF-173 | 20 |
| PNNL Dock | Slack High | Pico (> 0.22µm ) | aPES | BF-178 | 20 |
| PNNL Dock | Slack High | Pico (> 0.22µm ) | aPES | BF-179 | 20 |
| PNNL Dock | Slack High | Pico (> 0.22µm ) | aPES | BF-187b | 19 |
| PNNL Dock | Slack High | Pico (> 0.22µm ) | aPES | BF-188b | 19 |
| PNNL Dock | Slack High | Pico (> 0.22µm ) | aPES | BF-189b | 19 |
| PNNL Dock | Slack High | Pico (> 0.22µm ) | aPES | BF-187c | 19 |
| PNNL Dock | Slack High | Pico (> 0.22µm ) | aPES | BF-188c | 19 |
| PNNL Dock | Slack High | Pico (> 0.22µm ) | aPES | BF-189c | 19 |
| PNNL Dock | Flood | Micro (> 5µm) | MCE | BF-081 | 3 |
| PNNL Dock | Flood | Micro (> 5µm) | MCE | BF-082 | 3 |
| PNNL Dock | Flood | Micro (> 5µm) | MCE | BF-083 | 3 |
| PNNL Dock | Flood | Pico (> 0.22µm ) | MCE | BF-090 | 2 |
| PNNL Dock | Flood | Pico (> 0.22µm ) | MCE | BF-091 | 2 |
| PNNL Dock | Flood | Pico (> 0.22µm ) | MCE | BF-092 | 2 |
| PNNL Dock | Flood | Pico (> 0.22µm ) | aPES | BF-201a | 40 |
| Cline Spit | Flood | Micro (> 5µm) | Nylon | BF-124 | 6 |

| | | | | | |
|---|---|---|---|---|---|
| Cline Spit | Flood | Pico (> 0.22µm ) | MCE | BF-125 | 0.425 |
| Cline Spit | Flood | Pico (> 0.22µm ) | MCE | BF-126 | 0.425 |
| Cline Spit | Flood | Pico (> 0.22µm ) | MCE | BF-130 | 0.45 |
| Cline Spit | Flood | Pico (> 0.22µm ) | MCE | BF-131 | 0.45 |
| Disco Bay | Flood | Macro ( > 20µm) | Nylon | BF-137 | 7.3 |
| Disco Bay | Flood | Micro (> 5µm) | Nylon | BF-138 | 7.3 |
| Disco Bay | Flood | Pico (> 0.22µm ) | aPES | BF-144 | 5 |
| Cline Spit | Flood | Viral (> 0.05µm) | MCE | BF-132 | 0.16 |
| PNNL Dock | Flood | Viral (< 0.22µm) | aPES | BF-202a | 3 |
| PNNL Dock | Flood | Viral (< 0.22µm) | aPES | BF-203a | 10 |
| PNNL Dock | Flood | Viral (< 0.22µm) | aPES | BF-204a | 14 |
| PNNL Dock | Flood | Viral (< 0.22µm) | aPES | BF-205a | 17 |
| PNNL Dock | Flood | Viral (< 0.22µm) | aPES | BF-206a | 12 |

| % Of Whol | Filtered Vc | Sequim Na | Sequim A2 | Sequim A2 | Final Extra | Sequim Na | Sequim Qu | Sequim Qu | Sequim Qu |
|---|---|---|---|---|---|---|---|---|---|
| 0.5 | 1 | 27.6 | 1.84 | 0.99 | 50 | 1380 | 1X dsDNA | 2.02 | 101 |
| 0.5 | 0.5 | 15.5 | 1.87 | 0.9 | 50 | 775 | 1X dsDNA | 103 | 5150 |
| 0.5 | 1.05 | n/a | n/a | n/a | 96 | n/a | 1X dsDNA | 43 | 4128 |
| 0.5 | 1 | n/a | n/a | n/a | 96 | n/a | 1X dsDNA | 45.4 | 4358.4 |
| 0.5 | 0.65 | n/a | n/a | n/a | 96 | n/a | 1X dsDNA | 67.3 | 6460.8 |
| 0.5 | 0.9 | 32.7 | 1.82 | 0.23 | 96 | 3139.2 | 1X dsDNA | 24.9 | 2390.4 |
| 0.5 | 0.5 | 16.8 | 1.77 | 0.28 | 96 | 1612.8 | 1X dsDNA | 11.3 | 1084.8 |
| 0.5 | 2.5 | 7.8 | 1.93 | 0.15 | 94 | 733.2 | 1X dsDNA | 5.5 | 517 |
| 0.5 | 2.5 | 3.8 | 2.45 | 0.03 | 94 | 357.2 | 1X dsDNA | 1.48 | 139.12 |
| 0.5 | 1 | 5.7 | 2.54 | 0.03 | 94 | 535.8 | 1X dsDNA | 3.78 | 355.32 |
| 0.5 | 1 | 9.9 | 2 | 0.13 | 94 | 930.6 | 1X dsDNA | 7.65 | 719.1 |
| 0.5 | 1 | 3.7 | 3.35 | 0.05 | 94 | 347.8 | 1X dsDNA | 2.74 | 257.56 |
| 0.5 | 1 | 13.4 | 2.02 | 0.04 | 94 | 1259.6 | 1X dsDNA | 4.86 | 456.84 |
| 0.5 | 0.75 | 6.8 | 2.09 | 0.04 | 94 | 639.2 | 1X dsDNA | 3.4 | 319.6 |
| 0.5 | 0.75 | 4 | 2.88 | 0.06 | 94 | 376 | 1X dsDNA | 2.31 | 217.14 |
| 0.5 | 0.75 | 4.1 | 2.52 | 0.05 | 94 | 385.4 | 1X dsDNA | 2.18 | 204.92 |
| 0.5 | 0.75 | 9.7 | 1.81 | 0.1 | 94 | 911.8 | 1X dsDNA | 3.39 | 318.66 |
| 0.5 | 0.75 | 8.5 | 2.18 | 0.07 | 94 | 799 | 1X dsDNA | 4.34 | 407.96 |
| 0.5 | 1.25 | 14.8 | 2.09 | 0.15 | 94 | 1391.2 | 1X dsDNA | 12.4 | 1165.6 |
| 0.5 | 1.25 | 20.4 | 2 | 0.12 | 94 | 1917.6 | 1X dsDNA | 7.4 | 695.6 |
| 0.5 | 1 | 19.7 | 1.83 | 0.12 | 94 | 1851.8 | 1X dsDNA | 13.6 | 1278.4 |
| 0.5 | 1 | 15.1 | 2.03 | 0.07 | 94 | 1419.4 | 1X dsDNA | 10.2 | 958.8 |
| 0.5 | 1 | 20.8 | 1.85 | 0.19 | 94 | 1955.2 | 1X dsDNA | 11 | 1034 |
| 0.5 | 1 | 25 | 1.97 | 0.08 | 94 | 2350 | 1X dsDNA | 17.6 | 1654.4 |
| 0.5 | 1.5 | 18.1 | 1.96 | 0.08 | 94 | 1701.4 | 1X dsDNA | 11.4 | 1071.6 |
| 0.5 | 1.75 | 16.2 | 2.06 | 0.05 | 94 | 1522.8 | 1X dsDNA | 6.85 | 643.9 |
| 0.5 | 1.25 | 7.2 | 2.52 | 0.05 | 94 | 676.8 | 1X dsDNA | 3.84 | 360.96 |
| 0.33 | 6.6 | 20.8 | 1.91 | 0.17 | 94 | 1955.2 | 1X dsDNA | 11.2 | 1052.8 |
| 0.33 | 6.6 | 36.9 | 1.78 | 0.96 | 96 | 3542.4 | 1X dsDNA | 17.4 | 1670.4 |
| 0.33 | 6.6 | 43.4 | 1.75 | 1.61 | 96 | 4166.4 | 1X dsDNA | 26.5 | 2544 |
| 1 | 20 | 12.5 | 1.61 | 0.32 | 96 | 1200 | 1X dsDNA | 5.55 | 532.8 |
| 1 | 20 | 94.4 | 1.79 | 0.71 | 96 | 9062.4 | 1X dsDNA | AB | AB |
| 0.33 | 6.27 | 164.4 | 1.84 | 1.37 | 94 | 15453.6 | 1X dsDNA | AB | AB |
| 0.33 | 6.27 | 156.1 | 1.84 | 0.65 | 94 | 14673.4 | 1X dsDNA | AB | AB |
| 0.33 | 6.27 | 171.4 | 1.84 | 0.78 | 94 | 16111.6 | 1X dsDNA | AB | AB |
| 0.33 | 6.27 | 158.3 | 1.85 | 0.4 | 96 | 15196.8 | 1X dsDNA | AB | AB |
| 0.33 | 6.27 | 139 | 1.85 | 0.52 | 96 | 13344 | 1X dsDNA | AB | AB |
| 0.33 | 6.27 | 116.3 | 1.87 | 0.29 | 96 | 11164.8 | 1X dsDNA | AB | AB |
| 0.5 | 1.5 | 9.7 | 2.03 | 0.03 | 96 | 931.2 | 1X dsDNA | 0.479 | 45.984 |
| 0.5 | 1.5 | 11.1 | 2.04 | 0.04 | 96 | 1065.6 | 1X dsDNA | 4.53 | 434.88 |
| 0.5 | 1.5 | 8.4 | 1.9 | 0.07 | 96 | 806.4 | 1X dsDNA | 3.27 | 313.92 |
| 0.5 | 1 | 11.4 | 1.97 | 0.06 | 96 | 1094.4 | 1X dsDNA | 6.84 | 656.64 |
| 0.5 | 1 | 15.1 | 1.83 | 0.41 | 96 | 1449.6 | 1X dsDNA | 7.91 | 759.36 |
| 0.5 | 1 | 7.7 | 1.94 | 0.03 | 96 | 739.2 | 1X dsDNA | 1.88 | 180.48 |
| 0.25 | 10 | 24.4 | 1.84 | 0.1 | 96 | 2342.4 | 1X dsDNA | 15.5 | 1488 |
| 0.5 | 3 | 11.4 | 1.43 | 0.18 | 96 | 1094.4 | 1X dsDNA | 4.18 | 401.28 |

| | | | | | | | | | |
|---|---|---|---|---|---|---|---|---|---|
| 1 | 0.425 | 45.1 | 1.74 | 1.55 | 96 | 4329.6 | 1X dsDNA | 42.7 | 4099.2 |
| 1 | 0.425 | 49.2 | 1.73 | 1.51 | 96 | 4723.2 | 1X dsDNA | 39.6 | 3801.6 |
| 1 | 0.45 | 54.7 | 1.82 | 0.46 | 96 | 5251.2 | 1X dsDNA | 46.7 | 4483.2 |
| 1 | 0.45 | 48.6 | 1.79 | 1.21 | 96 | 4665.6 | 1X dsDNA | 40.7 | 3907.2 |
| 1 | 7.3 | 7.1 | 1.62 | 0.48 | 96 | 681.6 | 1X dsDNA | 4.35 | 417.6 |
| 0.5 | 3.65 | 12.8 | 1.69 | 0.38 | 96 | 1228.8 | 1X dsDNA | 8.45 | 811.2 |
| 0.5 | 2.5 | 73.2 | 1.85 | 1.86 | 96 | 7027.2 | 1X dsDNA | AB | AB |
| 1 | 0.16 | 6.5 | 1.39 | 0.17 | 96 | 624 | 1X dsDNA | 0.413 | 39.648 |
| 0.25 | 0.75 | 6.3 | 1.74 | 0.02 | 96 | 604.8 | 1X dsDNA | 0.051 | 4.896 |
| 0.25 | 2.5 | 9.6 | 1.63 | 0.03 | 96 | 921.6 | 1X dsDNA | 1.04 | 99.84 |
| 0.25 | 3.5 | 7.1 | 1.86 | 0.02 | 96 | 681.6 | 1X dsDNA | 1.4 | 134.4 |
| 0.25 | 4.25 | 10.2 | 1.73 | 0.13 | 96 | 979.2 | 1X dsDNA | 5.03 | 482.88 |
| 0.25 | 3 | 10.4 | 1.66 | 0.05 | 96 | 998.4 | 1X dsDNA | 4.82 | 462.72 |

| Referenced in Manuscript Figure(s) |
| --- |
| Figure 11 |
| Figure 11 |
| Figure 11 |
| Figure 11 |
| Figure 11 |
| Figure 11 |
| Figure 11 |
| Figure 9 |
| Figure 9 |
| Figure 10 |
| Figure 10 |
| Figure 10 |
| Figure 10 |
| Figure 9, Figure 11 |
| Figure 9, Figure 11 |
| Figure 9, Figure 11 |
| Figure 9, Figure 11 |
| Figure 9, Figure 11 |
| Figure 9 |
| Figure 9 |
| Figure 9 |
| Figure 9 |
| Figure 9 |
| Figure 9 |
| Figure 10 |
| Figure 10 |
| Figure 10 |
| Figure 11 |
| Figure 11 |
| Figure 11 |
| Figure 11 |
| Figure 11 |
| Figure 11 |
| Figure 11 |
| Figure 11 |
| Figure 11 |
| Figure 11 |
| Figure 11 |
| Figure 10 |
| Figure 10 |
| Figure 10 |
| Figure 10 |
| Figure 10 |
| Figure 10 |
| Figure 11 |
| Figure 9 |

Figure 9
Figure 9
Figure 9
Figure 9
Figure 9
Figure 9
Figure 9
n/a
n/a
n/a
n/a
n/a
n/a
