## Supplementary figures and images for "Sampling Microbial Dynamics in the Salish Sea Estuary: Evaluating Methods to Capture Cyanobacteria and Cyanophage"

### Supplementary Figure 1

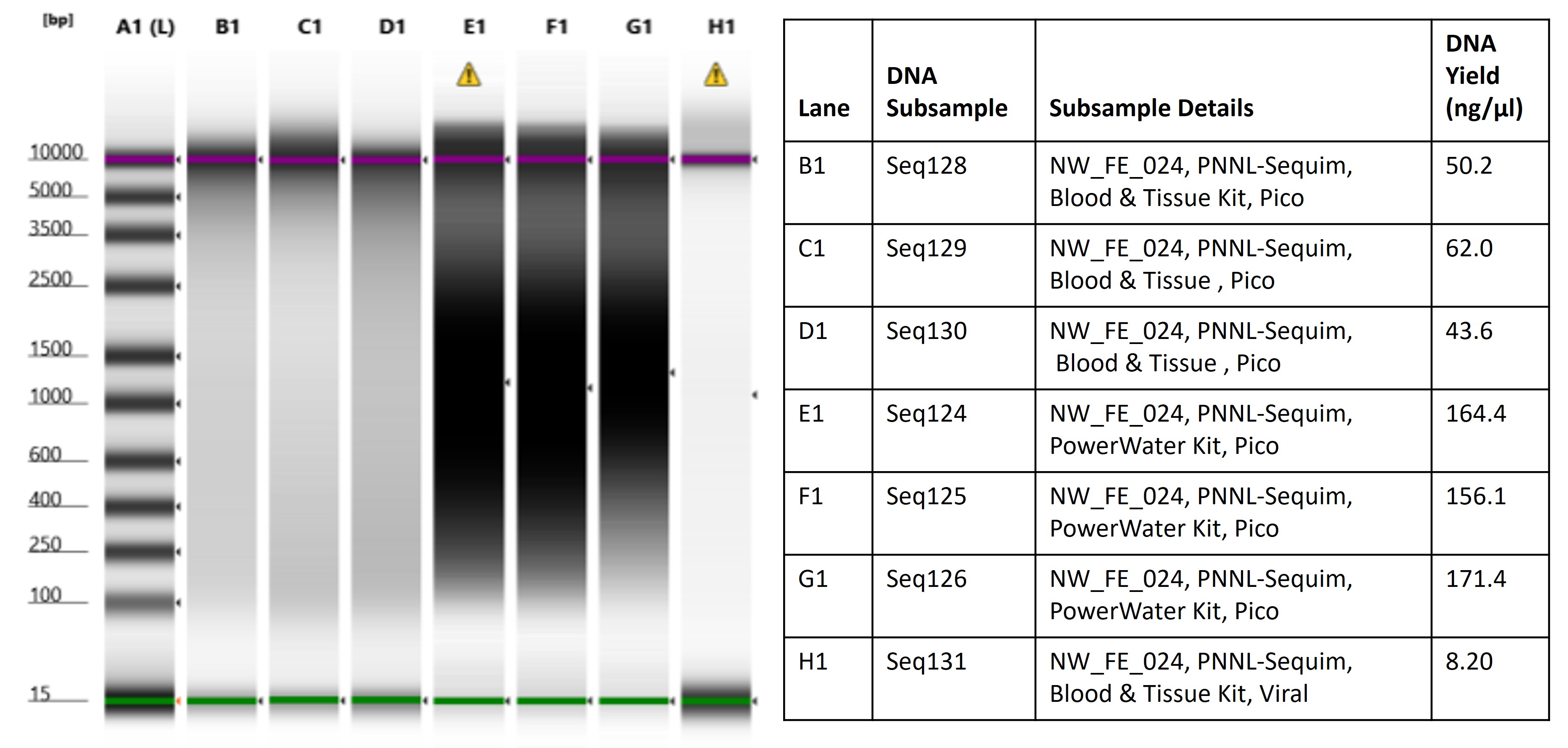
